# Microvillar cartography: a super-resolution single-molecule imaging method to map the positions of membrane proteins with respect to cellular surface topography

**DOI:** 10.1101/2020.06.11.145383

**Authors:** Shirsendu Ghosh, Ronen Alon, Andres Alcover, Gilad Haran

**Affiliations:** Department of Chemical and Biological Physics, Weizmann Institute of Science, Rehovot 76100, Israel; Department of Immunology, Weizmann Institute of Science, Rehovot 76100, Israel; Lymphocyte Cell Biology Unit, INSERM U1221, Department of Immunology, Institut Pasteur, Paris 75015

## Abstract

We introduce Microvillar Cartography (MC), a method to map proteins on cellular surfaces with respect to the membrane topography. The surfaces of many cells are not smooth, but are rather covered with various protrusions such as microvilli. These protrusions may play key roles in multiple cellular functions, due to their ability to control the distribution of specific protein assemblies on the cell surface. Thus, for example, we have shown that the T-cell receptor and several of its proximal signaling proteins reside on microvilli, while others are excluded from these projections. These results have indicated that microvilli can function as key signaling hubs for the initiation of the immune response. MC has facilitated our observations of particular surface proteins and their specialized distribution on microvillar and non-microvillar compartments. MC combines membrane topography imaging, using variable-angle total internal microscopy, with stochastic localization nanoscopy, which generates deep sub-diffraction maps of protein distribution. Since the method is based on light microscopy, it avoids some of the pitfalls inherent to electron-microscopy-based techniques, such as dehydration, carbon coating and immunogold clustering, and is amenable to future developments involving e.g. live-cell imaging. This Protocol details the procedures we developed for MC, which can be readily adopted to study a broad range of cell surface molecules and dissect their distribution within distinct surface assemblies under multiple cell activation states.

## Introduction

The surfaces of eukaryotic cells are usually rough and may carry multiple protrusions of various types. Some cells are covered with protrusions throughout their surface^1,2^. In particular, neutrophils, and lymphocytes including B- and T-cells and natural killer cells, are covered with microvilli, which are slender and flexible actin-rich protrusions, with a typical diameter of 100 nm and a length of a few hundred nanometers^2^.^3^ These membrane protrusions do not only create a structural scaffold for the formation of microclusters of specialized receptors and adhesion molecules, but also serve as a physical barrier for the diffusion of non-microvillar molecules.^4^ Microvilli should be distinguished from other dynamic protrusions that are de-novo generated by motile cells. These include thin, sheet-like actin-rich protrusions at the cell’s leading edge called lamellipodia, which play a key role in cellular migration and surface sensing ^5,6^. Additional projections include filopodia, which are critical for cell sensing of cues and directionality of migration^7^.

As opposed to these de-novo generated protrusions, microvilli are pre-existing structures. Their role in the physiology of lymphocytes has been disputed over many years. Lymphocytes travel in blood vessels, and they have to attach to endothelial cells of blood vessel walls before they penetrate into target tissues. It has been shown that adhesion molecules like L-selectin and P-selectin glycoprotein ligand-1 are concentrated on microvilli^8^, and therefore these cellular organelles have been considered as essential for the initial contact between blood borne lymphocytes and the endothelial surfaces they interact with^8^.

A different role for microvilli has been recently suggested by observations that lymphocytes use their microvilli to search for antigenic signals presented by rare antigen presenting cells (APCs)^3,9,10^. Moreover, microvilli have been shown to be involved in force-driven penetration of the dense glycocalyx barrier of APCs, which may otherwise hinder ligand/receptor interaction.^11^ This novel understanding was corroborated by our Microvillar Cartography (MC) studies, which demonstrated that T-cell receptors (TCRs) are almost exclusively positioned on microvilli in T cells prior to their encounter of antigenic signals^10^. While our experiments were conducted on fixed cells, experiments by Krummel and coworkers^3^ using light-sheet microscopy and by Sherman and coworkers^12^ using a combination of atomic force and super-resolution microscopies demonstrated the dynamic nature of microvillar involvement in contact formation with APCs. Recently, we used the MC approach to show that, in addition to TCRs, multiple membrane proteins involved in the initiation of the immune response are also enriched on microvilli^9^. We further showed that TCR localization on microvilli is mediated by ERM (ezrin, radixin and moesin) proteins, which connect membrane proteins to the cortical actin cytoskeleton. Our study has established the role of microvilli as specialized activation hubs of T cells^9^. Interestingly, recent work suggests that microvillar protrusions do not disappear following T-cell activation and do not collapse during maturation of the immunological synapse (IS). Rather, they are scattered around the cells or moved to the central supramolecular activation cluster (cSMAC) of the IS.^13^

The MC method, which can be used to study any cell type and any membrane protein, combines two optical microscopies to image the surface topography of cells and localize specific membrane proteins with respect to this topography. In this Protocol, we describe in detail the implementation of MC, from cell preparation and staining through imaging studies and statistical data analysis.

### Overview of the Microvillar Cartography procedure

MC is a novel strategy for determining at nanometer resolution the location of membrane molecules with respect to surface structures, using a synergistic combination of two types of fluorescence microcopy: variable-angle total internal reflection microscopy (VA-TIRFM) resolves the 3D topography of the cell, and stochastic localization nanoscopy (SLN) determines the position of molecules with respect to this topography. A schematic of the MC procedure is shown in the Fig. 1.

**Figure 1:**
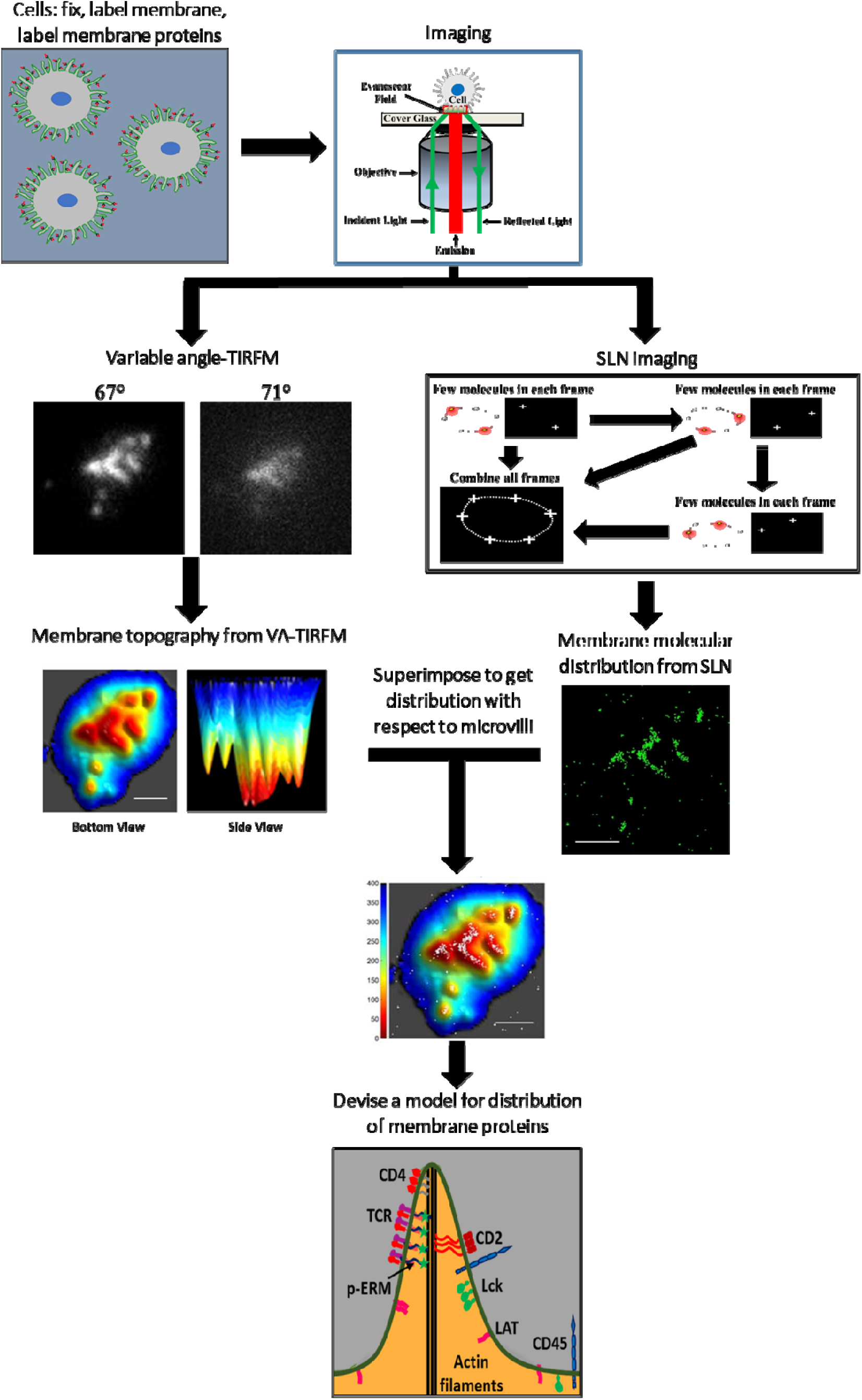
Schematic of Microvillar Cartography (MC)

VA-TIRFM ^*14-17*^ is based on the physics of TIRF illumination, which decays exponentially as a function of the distance from the glass coverslip. The penetration depth of the TIRF evanescent field (*d(θ)*) is defined as the distance where the intensity of the evanescent field is 1/e times of the incident laser intensity. The penetration depth is tunable by changing the angle of incidence, θ. With a known *d(θ)*, the relative intensity of fluorescence at each point in the image can be used to calculate its distance (*δz*) from the glass surface. The set of *δz* values can then be used to construct a 3D image of the fluorescently labelled cell membrane. It is possible to increase the statistical precision of the resulting image by combining measurements at several values of *θ*.

SLN is a form of super-resolution fluorescence microscopy in which bursts of photons from blinking labelled molecules are used to localize them with accuracy well below the diffraction limit^*18,19*^ (Fig. 1). We implement a simple technique that enables measuring molecules located on two different planes in the z direction, which we term dual-plane SLN (Fig. S3 of Ref ^10^). In this technique, a piezo-stage is used to move the sample up or down (Fig. S3*A* of Ref ^10^), and data recording is repeated, leading effectively to the collection of signals from molecules at different planes in the sample. Sectioning is achieved by rejecting out-of-focus signals of single molecules during data analysis. By combining images from VA-TIRFM and SLN, we obtain a map of the positions of membrane molecules of interest in relation to the 3D topographical distribution of T cell microvilli (Fig. 1).

While MC is general, and can be applied to any cell, we describe it here based on our studies of T-cell microvilli. The experiment starts with the labelling of specific membrane proteins using fluorescent antibodies, which is done at low temperature to prevent potential cell activation or clustering (Fig. 2A-C). To increase the resolution, we refrain from using secondary antibodies, but rather employ either commercially available or in-house labelled primary antibodies. The cells are then fixed and stained with a membrane dye such as FM143fx (Fig. 2D-E). Importantly, if the antigenic epitope for antibody labelling of the membrane protein of the interest is exposed outside the cell membrane, then labelling with antibodies is carried out before fixation (Fig. 2). This is due to the fact that the fixation procedure sometimes leads to partial masking of antigenic epitopes. For detection of intracellular proteins, fixation and permeabilization are performed prior to labelling (Fig. 3).

**Figure 2:**
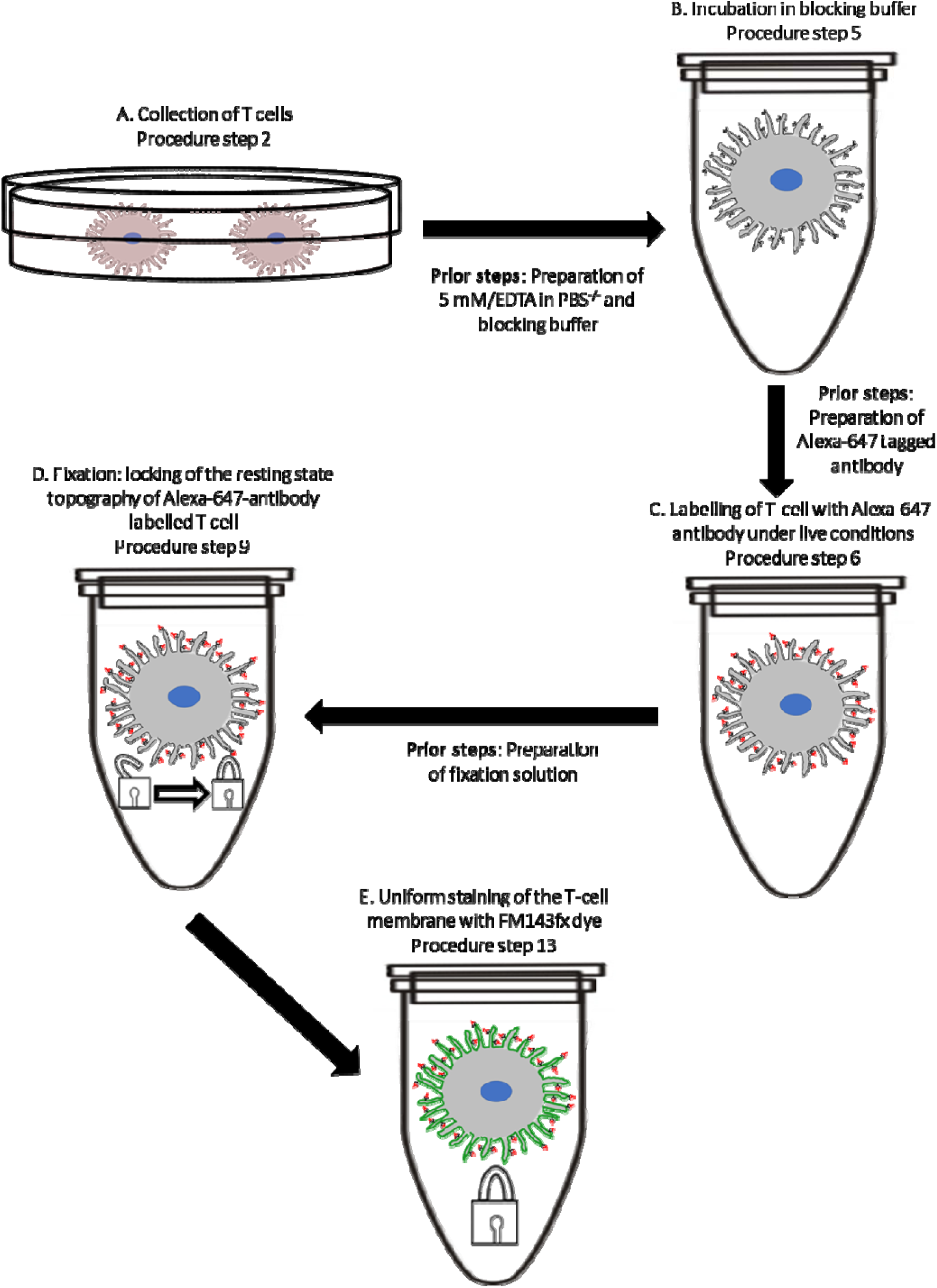
Flow chart of T-cell labelling when the membrane protein of interest is exposed outside the cell.

**Figure 3:**
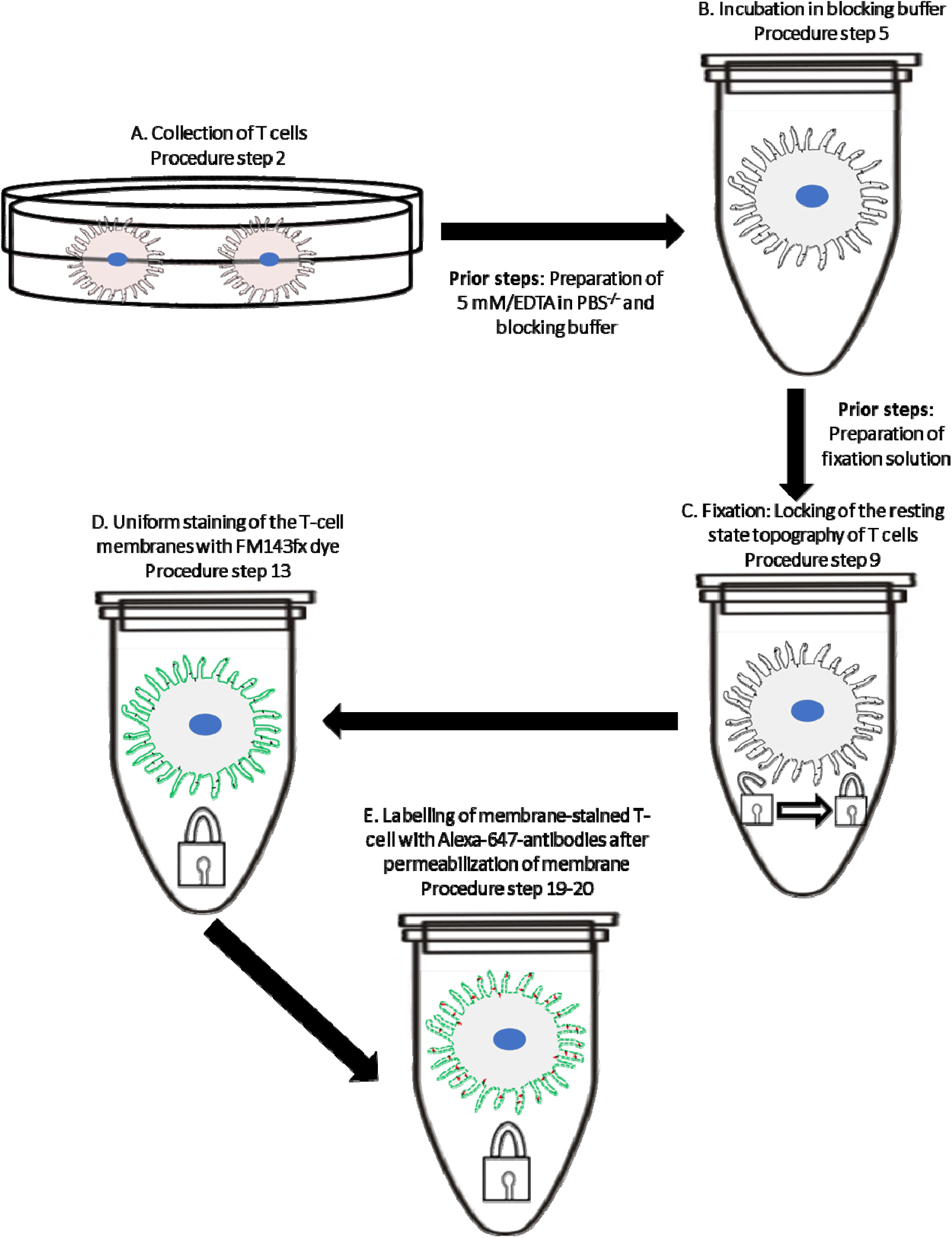
Flow chart of T-cell labelling when the membrane protein of interest is exposed only within the cell.

Using VA-TIRFM, we first resolve the finger-like structure of T cell microvilli. Based on measurements with different angles of incidence, we reconstruct an average 3D topographical map of the T cell surface. We then analyze the images and prepare maps of regions of individual microvilli and their tip positions, which we term LocTips maps. Using SLN, we localize labelled membrane proteins with a ∼10 nm accuracy at two focal planes, 0 nm and -400 nm. Molecules are assigned to one of the planes based on the size of their point-spread function (Fig. S2 of Ref. 10). The measurement at two different planes guarantees that proteins localized on membrane regions situated somewhat further from the surface due to the length of microvillar protrusions of resting T cells (∼300-400 nm, Ref. 2) would not be overlooked.

Using the two sets of images and image analysis protocols to be described in detail below, we segment the membrane area of each cell into either the microvillar region or the cell-body region, and then determine the distribution of each of the studied molecules in the two regions. We can also quantify changes in the molecular distribution as a function of distance from the central microvillar region (defined as the region that is not more than 20 nm away from the pixel with the minimum δz value). This analysis does not depend on the diffraction-limited resolution of the TIRF images in the X-Y plane, but only on the high resolution in the Z direction, which stems from the properties of the evanescent field in TIRFM. Finally, from simultaneous SLN experiments with two different labelled membrane proteins we can compute the co-localization probability (CP), which allows us to identify pairs of proteins that, due to their interaction, are situated on the average closer to each other than expected based on the cellular topography. Here some caution is in place, because two proteins located on a relatively small object like a microvillus might appear to co-localize even if they do not interact. A careful comparison to pairs of proteins that are not known to interact facilitates this kind of analysis. A detailed description of our CP analysis is, however, beyond the scope of this Protocol, and can be found in our recent publication^9^.

### Comparison to other approaches

For many years, the classical method for the localization of proteins with respect to cellular organelles at high resolution has been immunogold labelling (IL), which is based on the ability to image small gold particles by the electron microscope^20^. Antibodies labelled with gold particles are used to stain specific protein molecules in samples that are then imaged using either scanning or transmission electron microscopy. IL allows observing ultrastructural features with the electron microscope resolution while at the same time detecting the positions of specific proteins. However, it suffers from several drawbacks. First, gold particles are relatively large objects on the molecular scale, so that labelling proteins at high density is not feasible. Second, gold particles might not be able to penetrate some regions in a biological sample. Third, these particles tend to stick to each other and/or to bind non-specifically to cellular structures. Finally, electron microscopy precludes extension to live imaging. All of these drawbacks are overcome by MC. The fluorescently labelled antibodies used here are relatively small, and methods for using even smaller labelled proteins, such as antigen-binding fragments^21,22^ or nanobodies^23,24^ have been developed in recent years. Therefore, labelling at high density (which is actually a pre-requisite for high resolution in SLN) and with an excellent penetration can usually be achieved. The fluorescently labelled antibodies also suffer less from non-specific interactions, and imaging with the optical microscope paves the way to extensions to live cells, as discussed briefly below.

### Limitations and prospects

The current implementation of MC, describe in this Protocol, has two significant limitations that future work might be able to overcome.

As discussed in some detail above, the SLN component of our protocol, which is based on TIRFM, is limited to quasi-2D imaging. To ameliorate this limitation, we perform SLN in two different planes within the same sample, which allows us to better observe the distribution of molecules in relation to the membrane topography. MC can be extended in principle to bona fide 3D imaging. Several 3D SLN methods have been developed in recent years, using various point-spread function engineering techniques or light collection from multiple planes^25^. These methods are all based on epi-illumination, i.e. on a laser beam that passes through the sample. They can be readily combined with VA-TIRFM microscopy by introducing a mirror that shifts the illumination beam periodically between the center and periphery of the objective to enable epi- or total internal reflection illumination, respectively.

Another shortcoming of the current MC approach is its restriction to fixed cells, due to the relatively slow SLN procedure, which requires the collection of many thousands of frames in order to obtain a high-enough resolution. This precludes studies of dynamic processes such as the formation of the IS. There have been several attempts in recent years to push SLN technology to faster time scales^26-28^. We mention here one recent algorithm, due to Eldar and coworkers, which exploits the sparse distribution of fluorescent molecules in samples prepared for SLN and the lack of correlation between different emitters to push the time resolution by a factor of ∼100 (Ref. 29). Future incorporation of this or similar algorithms into the MC method should allow performing live-cell imaging, which may decipher, for example, the relationship between microvilli collapse and IS formation during activation.

## Experimental Design

The MC procedure encompasses broadly three stages: labelling, imaging and analysis. Figs. 2-5 summarize the experimental workflow of each stage, and a detailed description is given below.

### Staining of the cell membrane and labelling of membrane proteins

The labelling strategy should satisfy the underlying principles of the two super-resolution imaging techniques used here, VA-TIRFM and SLN. In particular, the dye molecules used in the SLN step of the procedure need to blink efficiently. Alexa Fluor-647 is well known for its unique blinking characteristics, including a high photon number yield per switching event and a low on-off duty cycle (so that the fluorophore spends only a small fraction of the time in the on state)^30^. Thus, we particularly recommend Alexa Fluor-647 tagged primary antibodies to label membrane protein molecules of interest. We strongly advise not to use secondary antibodies in the labelling procedure, because they will decrease the resolution of the image due to the extra size. In the absence of a commercially available labelled primary antibody, it is possible to label antibody proteins in house-see instructions in the sub-section ‘Antibody Labelling with Alexa Fluor-647’ of the ‘Reagent preparation’ section. We standardized the labelling procedure in such a way that it will preserve the genuine membrane topography of T cells. Briefly, we first wash the T cells with 5 mM ethylenediamine tetraacetic acid (EDTA) in a phosphate-buffered saline without calcium and magnesium (PBS^-/-^), as this helps maintaining the microvilli-dominated topography of T cells. A washing strategy similar to this one may be readily developed for other cell types. To reduce non-specific staining by antibodies, the cells are incubated in a blocking solution (1% bovine serum albumin (BSA), 5 mM EDTA, 0.05% sodium azide (NaN_3_), PBS^-/-^) on ice (Fig. 2B). If the membrane protein of interest has an antigenic epitope exposed outside the cell membrane, this protein is labelled before cell fixation (Fig. 2C). This is essential since fixation may destroy the exposed antigenic epitope. Further, a low temperature (4°C) is chosen for the labelling process in order to prevent cellular activation induced by the labelling antibody. This is particularly important when using antibodies that may affect biological function, such as anti-TCRαβ or anti-CD3, which activate T cells at 37°C and induce receptor clustering and internalization. We verified that labelling of T cells at 4°C does not lead to any artifact; see the section *‘Labelling at 4°C captures the bona fide resting state’* in Ref. *9*. We recommend performing a similar control for any cell type to be labelled at 4°C. We perform cell fixation with a specific fixation buffer (4% (wt/vol) paraformaldehyde, 0.2%–0.5% glutaraldehyde, 2% (wt/vol) sucrose, 10 mM EGTA, and 1 mM EDTA, PBS) that was shown by scanning electron microscopy (SEM)^9,10^ to retain the original membrane topography (Fig. 2D). We observe that the use of glutaraldehyde (0.2%–0.5%) during fixation is essential to retain the original membrane topography of T cell.

We recommend performing membrane staining following fixation (Fig. 2E). To obtain an unbiased representation of the cellular membrane topography from the TIRFM images, the cell membrane needs to be stained homogenously. Non-homogeneous staining of the cellular membrane would lead to inherent differences of the registered emission intensity of the dye in different regions of cell membrane. This may mask the intensity differences between dye molecules due to their axial positions, which would lead to errors in reconstruction of the 3D membrane topography. Thus, we choose to use the dye FM143fx, as it has been shown to stain cellular membranes homogeneously. (We have directly tested that FM143fx homogeneously label the T-cell membrane-see Ref ^9^ in the section ‘Cellular labelling with antibodies’ and in Fig. S1V-W.)

If the antigenic epitope of a specific membrane protein is within the cytoplasm, the cell membrane needs to be permeabilized for labelling. In this case, we prefer to invert the order introduced above and label membrane proteins *following* membrane staining (Fig. 3). This order of labelling steps is selected to prevent penetration of FM143fx molecules into the cell, which may occur if permeabilization is done before membrane staining. Penetration of the membrane dye may lead to an enormous background signal during imaging. In conclusion, our labelling procedure is designed to keep microvilli (or other membrane structures) intact, so as to capture the cells in their true resting state.

### Imaging

*TIRFM setup:* We perform the microscopy experiments using a custom-built TIRF-based setup. A detailed description of this setup follows (Fig. 4). However, any TIRF microscope that allows varying the excitation angle may be used, in principle. In our microscope, two different laser lines, at 532 nm (50 mW; Cobolt) and 642 nm (150 mW; Toptica), are coupled into a polarization-maintaining single-mode fiber by using a series of dichroic beam splitters (z408bcm, z532bcm; Chroma) and an objective lens (M-20×; N.A., 0.4; Newport). The optical fiber is pigtailed into an acousto-optic tunable filter (AOTFFnc-VIS-TN-FI; AA Opto-Electronic), which enables separately modulating each excitation beam. Free-space combination of the two laser lines is also possible. To maximize coupling efficiency, the polarization of each laser is tuned by using a polarizer (GT10-A; Thorlabs). A free-space isolator is used to block back-scattering into the laser, and is followed by a half-wave plate to rectify the polarization rotation caused by the isolator. The power of each laser is controlled by the computer. The first-order output beam from the AOTF is expanded and collimated to a diameter of 6 mm by achromatic lenses (01LAO773, 01LAO779; CVI Melles Griot). The expanded laser beams are focused at the back focal plane of the microscope objective lens (UAPON 100×OTIRF; N.A., 1.49; Olympus) by an achromatic focusing lens (f = 500 mm; LAO801; CVI Melles Griot), and total internal reflection is achieved at the sample by shifting the position of the focused beam from the center of the objective to its edge. Fluorescence emitted by the sample passes through a multiple-edge dichroic beam splitter (Di03-R405/488/532/635-t1-25×36; Semrock), which separates excitation beams from the fluorescence light, and is then coupled out from the side port of the microscope (Olympus IX71). The light is focused by a tube lens (f = 180 mm; Olympus), and then relayed with another achromatic lens (f = 100 mm; ACL0304; CASIX). The residual scattered laser light that passes through the dichroic beam splitter is blocked by notch filters (NF01-405/488/532/635 StopLine Quad-notch filter and ZET635NF; Semrock). A selective emission filter (z488-532-647m; Chroma) is also introduced within the light path. The fluorescent image is split into two areas of a single EMCCD chip (iXonEM +897 back-illuminated; Andor) by a dichroic beam splitter (640dcrx 228869; Chroma). The upper half of the EMCCD chip is dedicated to record emission below 640 nm, which we refer to as the green channel. The bottom half records emission above 640 nm, which we refer to as the red channel. Each spectrally separated image is collected with a single lens (f = 150 mm; 01LAO551; CVI Melles Griot) to refocus, and the two images are projected onto the two halves of the CCD chip. The final magnification on the EMCCD camera is 240×, resulting in a pixel size of 66.67 nm.

**Figure 4:**
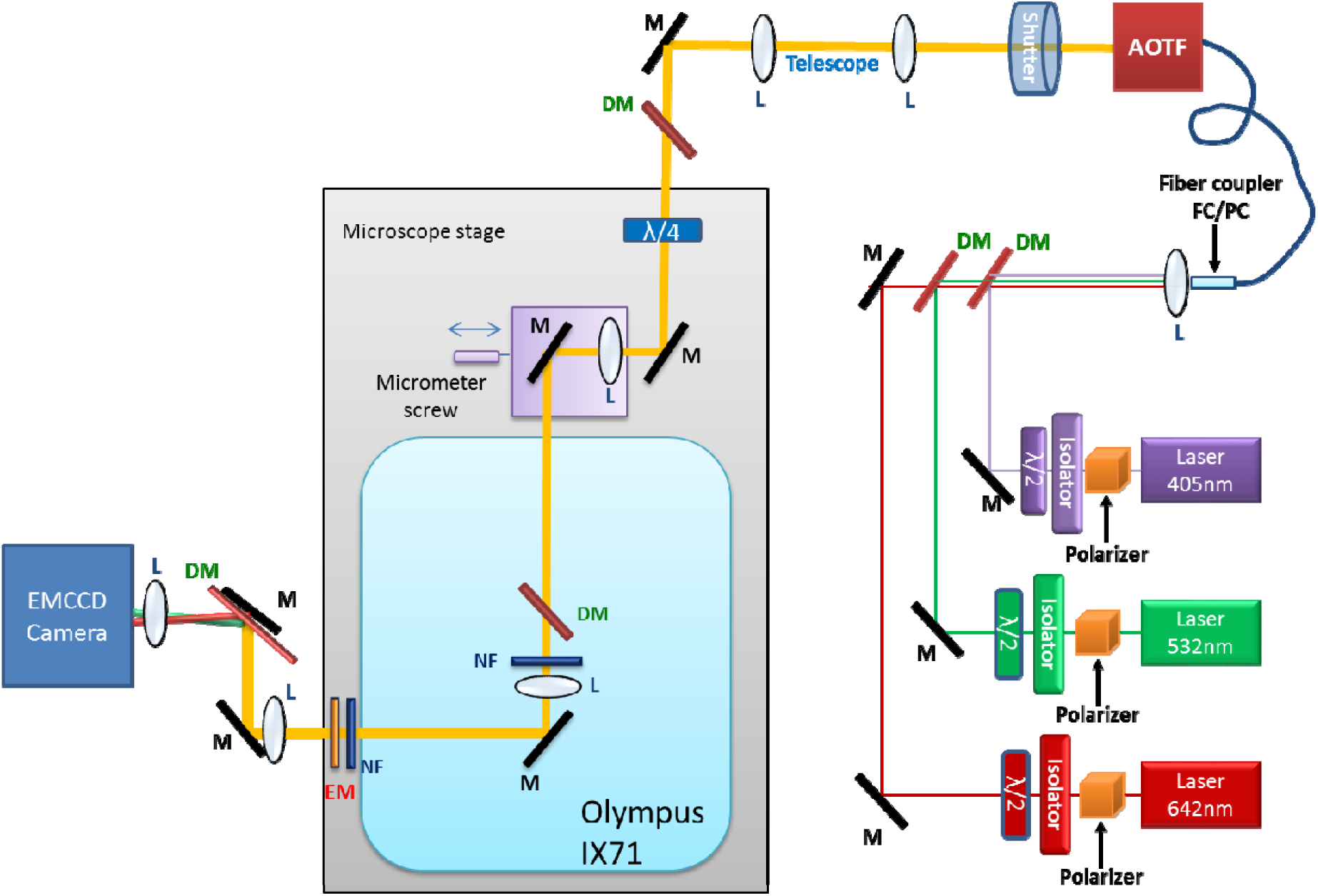
Schematic of the microscope setup. See text for details. Abbreviations used: λ/2: half-wave plate; M: mirror; DM: dichroic mirror; L: lenses; λ/4: quarter-wave plate; NF: notch filter; EM: emission filter.

We start the imaging cycle of a cell by using weak illumination of the 532-nm laser (such that the excitation power at the sample is 10-20 μw, corresponding to 6-12 W/cm^2^) to record 50 TIRF frames of the cell membrane at a series of angles of incidence (we typically use 63.0°, 64.2°, 65.5°, 66.8°, 68.2°, 69.7°, 71.2°, and 73.0°). Note that to determine the angle of incidence, the following equation is used, θ = sin −1 (D (FL × η_1_)), in which θ is the angle, D is the distance of the illumination beam from the center of the objective (which we measure by reading the scale of a linear stage used to translate the beam), FL is the focal length of the objective lens (1.8 mm in our case) and the refractive index of glass and oil (η_1_) is 1.52. Following the VA-TIRFM measurement, we perform SLN measurements at an incident angle of 66.8° using the 642-nm laser (Toptica) operating at a power of ∼60–70 mW (corresponding to ∼36–44 kW/cm^2^). The SLN images are recorded on the red (642 nm) channel of the EMCCD camera. We use a piezo stage (PI nano Z-Piezo slide scanner; PI) to move the sample up or down, and collect the SLN images at the 0 nm and −400 nm focal planes. For each focal plane we collect 30,000 camera frames, divided into 10 movies (3,000 frames in each, 15 ms per frame). We achieve the sectioning effect of dual-plane SLN by rejecting the out-of-focus images of single molecules during the localization procedure (see the section ‘dual-plane SLN’ below)^10^. The SLN imaging of membrane protein molecules at the −400 nm focal plane ensures that we are not missing any molecules due to the complex 3D topography of cell membrane. This allows us to rule out the possibility of biased detection of membrane proteins of the microvillar region compared to cell-body regions. Here we need to clarify the ambiguity between the penetration depth and the detection limit of molecules based on their distance from the glass surface upon illumination by evanescent field. The penetration depth of the evanescent field signifies the distance at which the field intensity is 1/e (∼37%) of the incident light intensity. However, depending on the incident intensity, molecules can be detected much deeper than that distance. We have verified that the intensity of the evanescent field (although decreased due to the penetration depth) is high enough to detect molecules with equal probability at the 0 nm plane and the -400 nm plane (see details in Ref. 9, particularly the ‘TIRF setup’ section and Figs. S1M– S1O). To allow us to correct images for sample drift during data collection (see below), we collect membrane dye reference images (50 frames) between the SLN movies under weak illumination of the 532-nm laser (10–20 μW, corresponding to 6-12 W/cm2). The TIRFM images are recorded in the green (532 nm) channel of the EMCCD camera.

### Analysis

*Analysis of VA-TIRFM images of cell membrane to reconstruct the 3-D topography of cell membrane:* As noted in the Overview, the intensity of fluorescence emission of a point on a uniformly stained object following excitation by an evanescent field depends on the distance of that point form the glass surface on which the object sits14-17 (Fig. 5A). This property is used to reconstruct the 3D surface topography (Fig. 5B-F). Thus, assuming that at a particular angle of incidence θ, the pixel of maximal intensity [I_max_(θ)] is the one closest to the glass surface, the relative axial distance of each point on the cell surface [with intensity I(θ)] from the glass (δz) can be calculated as follows: δz = ln(I_max_(θ)/ I(θ)/ d(θ)). Here, d(θ) is the penetration depth of the TIRF evanescent field with an angle of incidence θ, given by 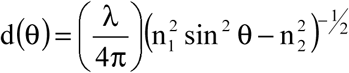, where λ is wavelength (532 nm in our case), n_1_ is the refractive index of the glass coverslip and immersion oil (1.52), and n_2_ is the refractive index of the buffer (1.35), as measured by a refractometer at 23 °C. The TIRF image taken at each angle of incidence leads to a separate topographical map.

**Figure 5:**
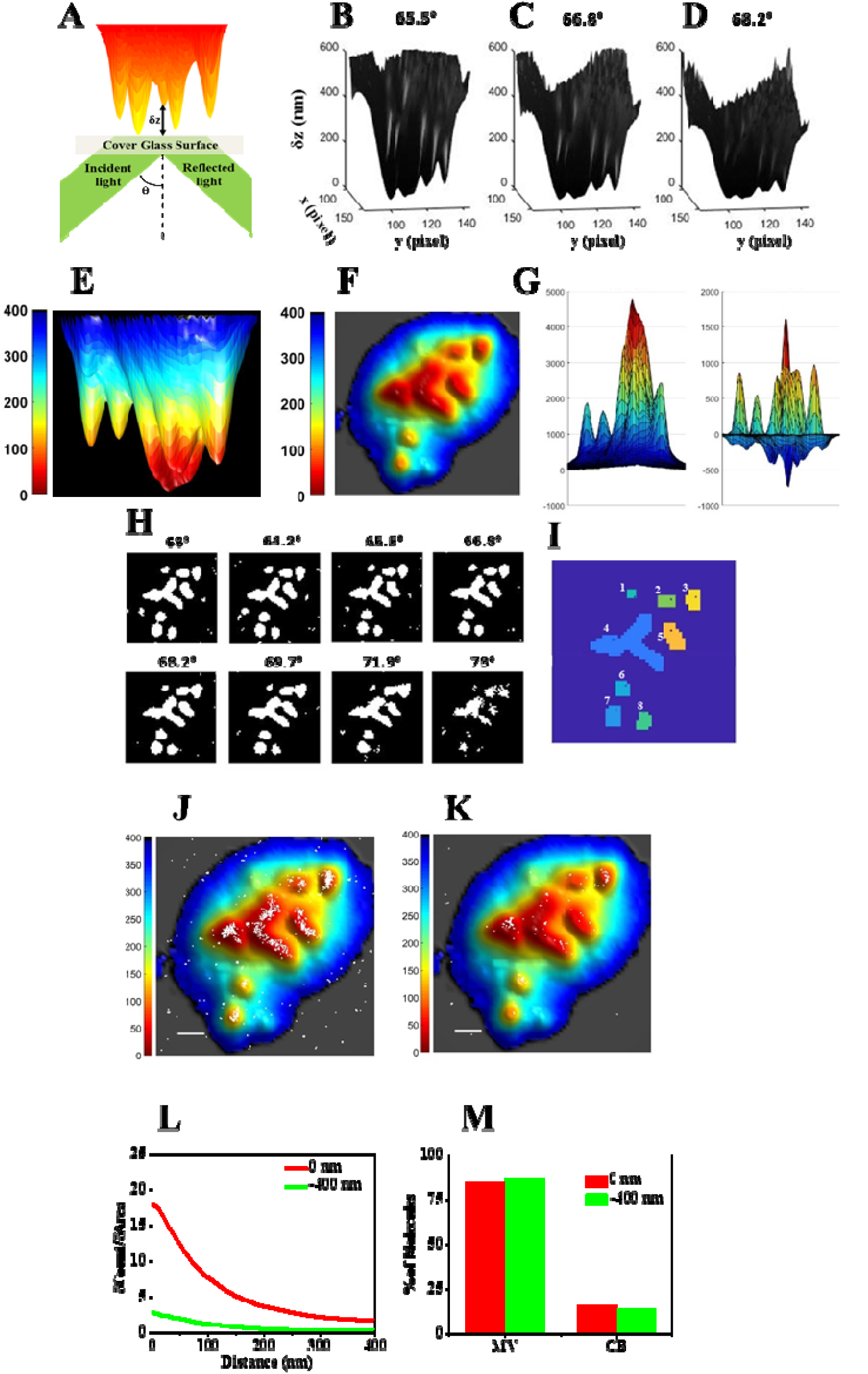
The analysis procedure. (A) A schematic of the TIRFM configuration. θ is the incident angle of light on the surface. A model membrane surface with microvilli is shown as placed on the glass surface. The distance from the glass to the surface of the object at each point is defined as δz and is obtained from analysis of measured images. (B–D) A series of 3D surface reconstruction maps of a Jurkat T cell from TIRFM images, based on calculated δz values from TIRFM measurements at angles of incidence (B) 65.5°, (C) 66.8°, and (D) 68.2° under weak illumination of a 532-nm laser. The δz map is shown from a direction perpendicular to the y–z plane. (E) A representative 3D surface reconstruction map of a T cell, calculated as the mean of the δz values of B-D. The δz values are represented by different hues with a step size of 6.25 nm. (F) A bottom-to-top 2D projection of E. (G) An example of a TIRFM image (Left) and its Laplacian of Gaussian (LoG) filter image (Right). (H) The regions above a threshold value of 5 in LoG filter images (white), calculated from a series of images acquired at variable angles of incidence (indicated above each image). (I) A LocTips map. The segmentation map of distinguishable protruding areas is determined by combining a series of segmentation analyses of VA-TIRFM measurements at incident angles from 63° to 73°. The coordinates of the pixel of minimum δz in each individual microvillus are marked (black cross). (J-K) Positions of protein molecules obtained from SLN (white dots) at the 0 nm plane (J) and at the -400 nm plane (K) are superimposed on membrane topography maps obtained from VA-TIRFM. (L) Cumulative increase of the fraction of total molecules on the cell as a function of the distance from the central microvilli region, normalized by the cumulative increase in the fraction of area (δCount/δArea) as a function of distance from microvilli. (M) Percentage of molecules on the microvillar (MV) region and the cell-body (CB) region of the membrane.

To define the regions of individual microvilli (MV) and the regions of the cell body (CB) on the topographical maps obtained with the above procedure, we use an algorithm developed in MATLAB. First, a “Laplacian of a Gaussian” (LoG) filter (Matlab; image processing tool kit), is used to process VA-TIRFM images. To flatten cell body regions while keeping the tip positions of microvilli appearing as peaks, we use the following parameters for the filter: Gaussian σ= 0.5; kernel size = 10 × 10 (Fig. 5G). Binary images are created from the LoG filter processed images by setting a threshold intensity level (we use a value of 5). This ensures the capture of most distinguishable microvilli regions. The “bwlabel” function in MATLAB is then used to segment the binary images to define individual microvilli areas. At first, the segmentation is performed on the image taken at the smallest angle of incidence and then the information from the image taken at the largest angle of incidence is combined to separate individual microvilli. This ensures inclusion of a maximum number of microvilli. More specifically, we use the function “imerode” in Matlab to treat the binary image of the smallest angle, which is then segmented using the function “bwlabel” (we set the parameter for “connected object size” in this function to 4). If the total size of an individual object is larger than 10 pixels (this mostly happens when two individual microvillar areas merge), the binary values of this structure are multiplied with the corresponding values from the image obtained at the maximum angle, resulting in a new binary map to which bwlabel is applied for segmentation. If this binary image of the object is not segmented by bwlabel, imerode is applied to the binary image from the maximum angle, and then this step is repeated up to four times. The eroded binary map is recovered by the “imdilate” function at the end. Finally, we combine the segmented areas generated from the images obtained at angles 65.5°, 66.8°, and 68.2° and the mean of the δz of each pixel is calculated and used to generate a map of the location of microvilli, the so-called ‘LocTips map’. The tip of each microvillus is defined as the pixel with the minimum δz value. We tested that our method is capable of identifying true microvilli. For that we used localization of L-selectin, as it is well known as a microvilli marker^8^. We showed that the L-selectin molecules are localized selectively on those areas of the T-cell membrane which are identified as microvilli by our analysis method (see Ref. 9 in the section ‘Microvilli can be identified using L-selectin localization’ and in Figs. S1J-L).

### Analysis of SLN movies to genarate super-resolved map of membrane proteins

Two common methods for localization of single molecules are 2D Gaussian fitting and center of mass analysis^19,31,32^ based on the point spread function model. 2D Gaussian fitting (2D-Fit) provides better precision than center of mass analysis (CM), however, the speed of 2D-Fit is much reduced, especially while attempting to fit a “bad” molecule (i.e. an impurity, a cluster of multiple molecules, an out of focus molecule, etc.) through multiple iterations. On the other hand, CM analysis is less precise but much faster since it requires one step calculation per molecule. We developed a simple algorithm, CFSTORM (Center of mass and 2d-Guassian Fit STORM) that combines the two methods: Individual emitters are identified in each frame by steps of thresholding. More specifically, a binary image is created from each frame by setting a threshold intensity level. Then the “bwlabel” function in MATLAB is used to segment the binary image to define individual emitters. Next, the data in a square of 11×11 pixels around the local maximum of each individual emitter is used in equations 1 and 2 below for calculating the center of mass coordinates (c_x_,c_y_) and the size of the pattern (s_x_, s_y_).^19^ We use a fixed threshhold for the size of molecules (s_x_, s_y_) in CM analysis. If a molecule has a larger size than the threshohold, it is discarded. We then perform 2D-Fit (equation 3) on those molecules that pass CM fitting and again discard bad molecules based on standard deviation (SD) values obtained from the fit (s_x_, s_y_). The emitters that pass the 2D-Fit step are true single molecules whose sub-pixel localization is determined from the parameters of the fit with maximum precision.

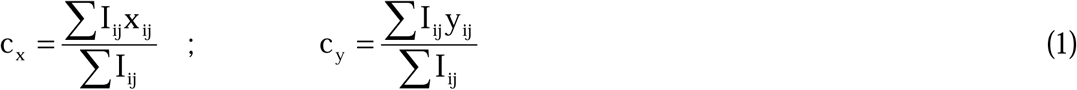

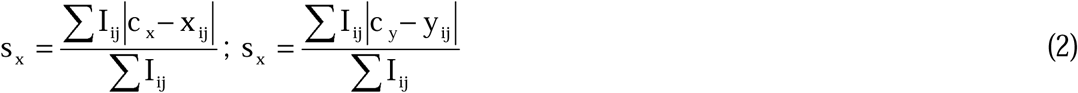

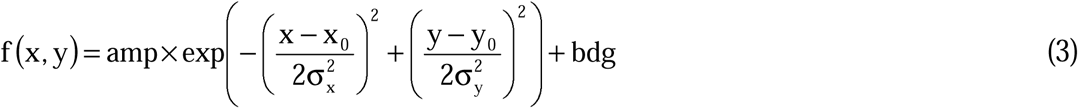

In these equations the i,j are the x-axis and y-axis indices for each pixel, x_i,j_ and y_i,j_ are the coordinates of the pixel and I_i,j_ is the intensity of the pixel. The coefficient ‘amp’ is the amplitude, ‘x_0_’, ‘y_0_’ is the center position and ‘bgd’ is a background parameter. To reconstruct an image of all protein molecules in the cell membrane, we combine the locations of single emitters determined from all SLN movies.

### Determination of localization precision

To this end, we image fluorescent beads (∼110 nm for our case) and adjust the intensity of laser in such a way that emission intensity of the beads matches the range of photon numbers emitted from a single blinking event of an Alexa Fluor 647 molecule. We measure the x and y coordinates and number of photons of each bead for sevaral frames. Based on these measurements, we estimate that the uncertainty in average localization in x–y is ∼11 nm. However, the localization precision is NOT the spatial resolution obtained in the final reconsructed image, as the latter also depends on the density of single-molecule detection events. To calculate the true image resolution, we suggest using the Fourier ring correlation analysis of images^33^, which takes into account of both localization uncertainty and labelling density.

### Dual-Plane SLN

We achieve a sectioning effect at the two measured planes (0 nm and - 400 nm) by discarding images of out-of-focus single molecules during the localization procedure. The recorded pattern of an out-of-focus molecule has a lower overall signal intensity and a higher spatial width. We use these features to reject out-of-focus molecules. For quantifying the defocusing of single-molecule images, we measure changes in the widths obtained from 2D Gaussian fits of images of individual fluorescent beads (∼110 nm in our case) adsorbed on a glass surface while moving the z position of the sample stage with a step size of 100 nm. The values of the SD of the Gaussian fits to bead images in the x and y directions at a stage position of −400 nm (that is, the glass plane is −400 nm away from the focal plane) are significantly larger than those obtained at a stage position of 0 nm (Fig. S3B in Ref. 10). This means that it is possible to collect “in-focus” molecules at 0 and −400 nm selectively by setting an appropriate threshold on the width. (We have explicitly tested that z sectioning of SLN measurements at the 0 nm and −400 nm focal planes can be achieved by setting an appropriate threshold on the width of single molecule. See the ‘Dual-Plane SLN’ section and Fig. S3C-F in Ref. 10).

### Determination of sample drift

The SLN experiment involves recording of a large number of separate movies. To correct for sample drift occurring during the measurement, we use the following movie-by-movie procedure, based on the TIRFM images of cell membrane acquired between SLN movies. We calculate the 2D cross-correlation function of each reference TIRFM image with the TIRFM image last obtained in the series. Due to the lateral drift of sample, we observe a peak in the 2D cross-correlation image that is shifted from the origin in both the x and y directions. We use the position of this peak, obtained from a 2D Gaussian fit, for correction of SLN movies.

### Determination of channel shift

To register the green (532 nm) and red (647 nm) SLN channels, we have to find the shift between the two channels. We observe that there is some leak from the 532 nm channel into the 647 nm channel when we record the membrane images of cells (this leak is not there when we take the true 647 nm images, as the 532 nm laser is then off). Thus, we cross-correlate the leaked membrane image in the 647 nm channel with the original membrane image in 532 nm channel. We use the cross-correlation to determine the shift between the two images and correct accordingly. Our Matlab code includes a procedure that corrects for both sample drift and channel shift self-consistently.

### Analysis of the distribution of membrane proteins with respect to microvilli

To analyze the distribution of each studied membrane protein, we merge the super-resolved map of membrane protein obtained from SLN movies acquired in the red channel with the membrane topography obtained from the TIRFM image acquired in the green channel after correcting for the channel shift and membrane drift effect in the red channel. We segment the membrane area of each cell into microvilli (MV) regions and non-microvilli or cell-body (CB) regions using the ‘LocTips map’ procedure described above. We then calculate the percentage of molecules on MV regions and on CB regions of each cell (See e.g. the Figs. S6A–S6D of Ref ^9^).

We also estimate the “cumulative fractional increase” of the number of molecules of a specific protein on each cell as a function of the distance from the central region of each microvillus (Fig. 5L, see also Figs. S6E–S6G of Ref. 9). To this end, we define the central region of each microvillus as the region that is not more than 20 nm from the microvillar tip, which is the pixel of the minimum δz value. The ‘boundary’ function of MATLAB is used to encapsulate this central region (See Fig. S6F of Ref. 9). This leads to a different shape of the central region of each microvillus, depending on the shape and orientation of the specific microvillus with respect to the surface. Concentric closed curves of a similar shape and increasing size (Fig. S6G of Ref. 9) are plotted, and the number of molecules in each concentric sector that forms between two closed curves is calculated. From these values, we determine the cumulative fractional increase, which is then normalized by the cumulative fractional increase of the area, to obtain the δCount/δArea plot, where δCount represents the fraction of total counts recorded within a concentric sector at a particular distance from the central microvillar region, whereas δArea represents the fractional area of that sector. The more a protein is localized to the microvillar region, the steeper is the slope of the δCount/δArea plot as a function of distance from the microvillar central region.

To decipher the distribution of a specific membrane protein in a specific cell type, we strongly advise to image at least 10 cells. The actual pattern of distribution of membrane protein may only be recognized from the average percentage of membrane protein molecules localized on the MV and CB compartments of the cells and the δCount/δArea plot averaged over all cells.

## MATERIALS

### Biological materials

1) Jurkat T cell (ATCC TIB-152) or:

2) Human T cell (isolated from both male and female healthy volunteer donors as described in Refs. 9,10)

! CAUTION Ethical approval should be obtained according to the relevant institutional and national regulations.

### Reagents

#### Labelling

3) Dulbecco’s phosphate buffered saline (PBS^-/-^), without calcium and magnesium (Biological Industries, REF: 02-023-1A)

4) Hanks □ balanced salt solution (HBSS^-/-^) without magnesium or calcium (Invitrogen #14175079)

5) Ethylenediaminetetraacetic (EDTA) acid solution (Sigma, 03690-100ML)

6) Ethylene glycol-bis(2-aminoethylether)-N,N,N′,N′-tetraacetic acid (EGTA, Sigma, E3889-10G)

7) MILLI-Q water

8) Sodium bicarbonate (NaHCO_3_,JT Baker 3506-01, CAS Number : 144-55-8)

9) Sucrose (Sigma, 84097)

10) Saponin (Sigma, 47036-50G-F)

11) Paraformaldehyde solution (16%, Electron Microscopy Sciences, Catalog number: 15710)

! CAUTION harmful for skin, proper precaution is needed.

12) Glutaraldehyde solution (25%, Electron Microscopy Sciences, Catalog number:16220)

! CAUTION harmful for skin, proper precaution is needed.

13) Alexa Fluor 647 NHS Ester (Succinimidyl Ester) (Thermo Fisher Scientific, Cat# A20106) ! CAUTION keep in dark at -20°C.

14) FM-143fx dye (Invitrogen, Catalog number: F35355)

! CAUTION keep in dark at -20°C.

15) Bovine serum albumin (BSA, Sigma, A-7030)

! CAUTION keep at 4°C.

16) Alexa Fluor 647 anti-human antibody/ LEAF purified anti-human antibody (Biolegand)

! CAUTION keep at 4°C or as specified by the supplier.

#### Imaging

17) Trizma hydrochloride (Sigma, T3253-1KG)

18) Sodium hydroxide (NaOH, Merck, CAS # 1310-73-2)

! CAUTION corrosive for skin, proper precaution measures are needed.

19) Potassium chloride (KCl, Merck, CAS # 7447-40-7)

20) Hydrochloric acid (HCl, 37%, Bio-Lab, cat no:000841020100)

! CAUTION corrosive for skin and eye, acute toxic, proper precaution is needed.

21) Glycerol (MP, cat no:800687)

22) Glucose (Merck, CAS #14431-43-7)

23) Poly-L-lysine (PLL) (0.01%; Sigma, P4707-50mL)

24) Cysteamine (Sigma, 30070-10G)

! CAUTION may absorb water, keep at 4°C in dehydrated condition. Do not use if melted.

25) Glucose Oxidase (from *Aspergillus niger*, Sigma, G7016)

! CAUTION keep at -20°C.

26) Catalase (from bovine liver, Sigma,C40-100MG)

! CAUTION keep at -20°C.

### Equipment

#### Labelling

1) Sterile pipette tips

2) Centrifuge tube

3) Micro Bio-Spin(tm) P-30 Gel Columns, (BIO-RAD #7326223)

4) Micropipettes

5) Centrifuge (Eppendorf, 5810R)

6) Syringe filters (0.22 μm, MILLEXGV, ref: SLGV033RS)

#### Imaging

7) Glass bottom culture dish (MatTek, 35 mm petri dish, 14mm micro-well, No. 1.0 cover-glass, part no.P35G1.0-14-C)

#### Reagent preparation

1) 1 M potassium chloride: Dissolve 74.55 g potassium chloride in 1L MILLI-Q water.

2) 4 M sodium hydroxide solution: Dissolve 80 g sodium hydroxide in 500 ml of MILLI-Q water

3) 1 M sodium hydroxide solution: Dissolve 20 g sodium hydroxide in 500 ml of MILLI-Q water

4) 1M trizma hydrochloride (pH 7.5-7.7): Dissolve 157.60 g trizma hydrochloride in 800 ml MILLI-Q water. Adjust pH to 7.5-7.7 with the appropriate volume of 4 M sodium hydroxide. Bring final volume to 1 liter with MILLI-Q water. Filter through a 0.22 μm filterand store at room temperature.

5) 1 M EGTA (Ethylene Glycol-Bis[β-Aminoether]N,N,′N′,N -Tetra-Acetic Acid) Solution: For a 100 mM EGTA stock solution, add 3.8 g to about 20 ml of distilled H_2_O and bring to pH 11 with 4 M sodium hydroxide and dissolve; then bring to pH 8.0 with hydrochloric acid and add H_2_O to a final volume of 100 ml. Filter through a 0.22 µM filter.

6) 40% Glucose **(w/v)**: Warm (∼60 °C) 50 ml of PBS^-/-^ and then add 40g of glucose. Stir/shake it to dissolve. Bring the final volume to 100 ml with PBS^-/-^. Filter through a 0.22 μm filter (MILLEXGV) and store at 4°C (after sealing the container properly with paraflim).

7) Cysteamine solution: Prepare 100 mM cysteamine in 125 mM trizma hydrochloride (pH 7.5-7.7).

! CAUTION should be prepared freshly.

8) Blocking solution: 1% BSA in 5 mM EDTA in PBS^-/-^.

9) Fixation solution: 4% paraformaldehyde, 0.4 % glutaraldehyde, 10 mM EGTA (ethylene glycol-bis(β-aminoethyl ether)-N,N,N’,N’-tetraacetic acid), 1mM EDTA, 2% sucrose in PBS^-/-^.

! CAUTION should be freshly prepared. Also harmful to skin, proper precaution is needed.

**CRITICAL STEP** use of glutaraldehyde is essential to retain the resting state topography of T cells during fixation.

10) Permeabilization buffer: 0.05% saponin+1 % BSA in PBS^-/-^.

! CAUTION should be freshly prepared.

11) TP 50 buffer: Add 100μL 1M trizma hydrochloride (pH 7.-7.7), 100μL 1 M potassium chloride, 1 ml glycerol and 800μL MILLI-Q water and then filter it Filter through 0.22 μm filter (MILLEXGV) and store at -20°C.

12) 50 mg/ml glucose oxidase: Dissolve 50 mg glucose oxidase in TP 50 buffer and store at - 20°C.

13) 40 mg/ml catalase: Dissolve 40 mg catalase in TP 50 buffer and store at -20°C.

14) Blinking Buffer: Add 500 μl of 100 mM cysteamine (! CAUTION should be freshly prepared.), 250 μl of 40% glucose, 10 μl of 50 mg/ml glucose oxidase, 1 μl of 40 mg/ml catalase and 239 μl of 125 mM trizma hydrochloride (pH 7.5-7.7).

! CAUTION should be freshly prepared.

15) Antibody Labelling with Alexa Fluor-647: If Alexa Fluor -647 tagged primary antibody is not available commercially then low endotoxin, azide-free (LEAF) purified commercial unlabelled antibodies may be labelled with Alex-647 in house using the following protocol. (! CAUTION the concentration of the antibody should be 1mg/ml.) The NHS ester of Alexa Fluor -647 is reacted with unlabelled antibody molecules in PBS buffer in a 10:1 molar ratio in presence of 0.1 M sodium bicarbonate buffer for 1 h at room temperature in the dark. Micro Bio-Spin column with Bio-Gel P-30 is used to remove the unlabelled dye molecules.

### Procedure

#### A. Labelling of the cell membrane and membrane proteins (Timing: 1 Day)

1) Treat the micro-well of the MakTek dish with 400 µl of 1M sodium hydroxide solution for 40 minutes. Wash it 10 times with MILLI-Q water and 5 times with PBS^-/-^ and PLL (100 µl of PLL are added for 30 min). Then, wash the well 3 times with PBS^-/-^. Place 400 µl of PBS^-/-^ within the well to keep it wet until loading the sample. Remove the PBS^-/-^ just before use.

2) Harvest 3 million T cells from human blood or from cell culture (e.g. Jurkat cells) and dilute with a 1:1 10 mM EDTA (Ethylenediaminetetraacetic acid)/ PBS^-/-^ solution. (So that the final concentration of EDTA in the solution is 5 mM.)

**CRITICAL STEP** This step is essential to keep the membrane topography of T-cells intact.

3) Spin the solution at 250G (∼1400 rpm) for 5 min and discard the supernatant.

4) Add 200 µl of the blocking solution to dissolve the pellet.

5) Incubate the solution on ice for 10 min.

#### If the membrane protein of interest has the epitope for antibody labelling exposed outside the cell membrane then step 6 should be followed. Otherwise, go directly to step 7

6) Add Alexa Fluor 647 tagged antibody to the solution such that the final concentration of the antibody should be 10-20 µg/ml and, incubate on ice for 20-40 min, depending on the antibody. (Note, if the antibody is known not to activate the cells or cluster receptors then the labelling can be done at 37°C). **?TROUBLESHOUTING**

7) Add 0.5 ml 5 mM EDTA/ PBS^-/-^ to it and spin the solution at 300 G, 5 min, 4 °C and discard the supernatant. (If the initial pellet size is small then a smaller volume of the 5 mM EDTA/ PBS^-/-^ solution may be used).

8) Repeat Step 7 once again.

9) Add 600 µl of fixation solution to dissolve the pellet and incubate the solution on ice for 2 hours.

**?TROUBLESHOUTING**

**CRITICAL STEP** The fixation procedure is optimized for Jurkat cells and may vary for other cell types.

10) Spin the solution at 800 G for 5 min, 4 °C and discard the supernatant.

11) Add 0.5ml PBS^-/-^ to dissolve the pellet, spin the solution at 800 G for 5 min, 4 °C and discard the supernatant. (If the initial pellet size is small then a smaller volume of the PBS^-/-^ solution may be used.)

12) Repeat step 11 once again.

13) Add 100 µl of 5 µg/ml FM-143FX membrane staining dye solution in HBSS^-/-^ and incubate on ice for 30 minutes.

**?TROUBLESHOUTING**

**CRITICAL STEP** The timing is important as a shorter incubation time leads to under-labelling of cell membrane, whereas a longer incubation time may lead to penetration of membrane dye inside the cytoplasm of the cells leading to increased background in imaging.

14) Add 0.5 ml PBS^-/-^, spin it at 800 G for 5 min, 4 °C and discard the supernatant.

15) Add 600 µl fixation solution to dissolve the pellet and incubate it on ice for 30 min.

16) Spin the solution at 800 G for 5 min, 4 °C and discard the supernatant.

17) Add 0.5 ml PBS^-/-^ to dissolve the pellet, spin it at 800 G for 5 min, 4 °C and discard the supernatant. (If the initial pellet size is small then a smaller volume of the PBS^-/-^ solution may be used.)

18) Repeat step 17 once again. Then, suspend the cells in 50 µl PBS^-/-^ buffer and store at 4 °C

**! CAUTION** do not freeze.

### If the membrane protein of interest has the epitope for antibody labelling buried inside the cell membrane then do not suspend the cells in PBS^-/-^ buffer and store at 4 °C after the washing step, rather follow steps 19-24

19) Add 200 µl of permeabilization buffer.

20) Add Alexa Fluor 647 tagged antibody to the solution such that the final concentration of the antibody is 10-20 µg/ml.

21) Incubate the solution at 4 °C for 4h to overnight depending on the antibody.

**?TROUBLESHOUTING**

22) Add 0.5 ml PBS^-/-^, spin at 800 G, 5 min, 4 °C and discard the supernatant. (If the initial pellet size is small then a smaller volume of the PBS^-/-^ solution may be used.)

23) Repeat Step 22 once again.

24) The cells are suspended in 50 µl PBS^-/-^ buffer and stored at 4 °C

**! CAUTION** do not freeze.

### Sample preparation for microscopy

25) On the day of imaging, 10 μl of labelled cell solution is mixed in 390 μl of the blinking buffer.

**! CAUTION** blinking buffer should be freshly prepared.

26) Place the solution of labelled cells in blinking buffer into the micro-well of the PLL coated MakTek dish. Imaging should be started after the cells settle down on the glass surface (which happens within ∼10 min).

**?TROUBLESHOUTING**

### Imaging (Timing ∼1h for imaging of 1 cell)

**! CAUTION:** Our microscope is custom build. It may be easily built following description provided above by an expert in light microscopy. The instrument needs to be properly aligned before imaging.

**! CAUTION:** We provide MATLAB code for the analysis of the results. If you plan to use this MATLAB code, you will need to make sure that files are saved in ‘.fits’ format and that file names are given in the correct format, as explained in Table S1 of the supporting file ‘Step by step guide to run analysis codes.pdf’.

#### Variable Angle-Total Internal Reflection Microscopy

27) Turn on the visible light lamp of the microscope.

28) Select a single cell on the micro-well of the MakTek dish using the eyepiece of the microscope.

29) Capture the bright-field image of the cell.

30) Turn off the visible light lamp of the microscope.

31) Turn on the 532 nm (green) laser, such that the excitation power at the sample will be 10–20 μW (corresponding to 6-12 W/cm^2^).

32) Set the angle of incident of the light to be at 66.8°.

33) Focus the laser light at glass surface.

34) Next, change the angle of incidence of the light to 63°.

CRITICAL STEP The focus should not be readjusted when changing the angle of incidence.

**?TROUBLESHOUTING**

35) Record the membrane image movie of the cell using the EMCCD camera (typical camera settings: Exposure time-0.1 s, number of frame-50, camera frame area: 256 pixel x 512 pixel for each channel).

36) Repeat steps 34-35 for angles of incidence 64.2°, 65.5°, 66.8°, 68.2°, 69.7°, 71.2°, and 73.0°.

CRITICAL STEP The focus should not be readjusted between movies.

#### Stochastic Localization Nanoscopy (SLN) imaging

37) Set the angle of incidence of the light again at 66.8°.

38) Record the membrane image only in the green channel (typical camera settings: Exposure time-0.1 s, number of frame-50, camera frame area: 256 pixel x 256 pixel in the green channel).

39) Block the 532 nm laser with the shutter.

40) Turn on/unblock the 642 nm (red) laser with high power (such that the excitation power at the sample will be 60-70 mW, corresponding to 36-44 kW/cm^2^).

41) Record the SLN movie of labelled receptor molecules in the red channel (Camera setting: Exposer time-0.015 s, number of frame-3000, Camera frame area: 256 pixel x 256 pixel in the red channel).

**?TROUBLESHOUTING**

42) Block the 642 nm laser with the shutter.

43) Unblock the 532 nm laser to illuminate the sample with the 532 nm laser light.

44) Repeat steps 38-43 10 times to acquire a total 30000 frames of SLN images of receptors.

45) Use the piezo stage to shift the imaging plane 400 nm away from the surface of the glass.

46) Record the membrane image only in the green channel (Camera settings: Exposure time-0.1 s, number of frames-50, camera arame area: 256 pixel x 256 pixel in the green channel).

47) Block the 532 nm laser with the shutter.

48) Unblock the 642 nm (red) laser with high power (such that the excitation power at the sample will be 60-70 mW, corresponding to 36-44 kW/cm^2^).

49) Record the SLN image movie of labelled receptor molecules only in the red channel (Camera setting: Exposure time-0.015 s, number of frames-3000, camera frame area: 256 pixel x 256 pixel in the red channel).

**?TROUBLESHOUTING**

50) Block the 642 nm laser with the shutter.

51) Unblock the 532 nm laser to illuminate the sample.

52) Repeat step 47-52, 10 times to acquire total 30000 frames of SLN images of receptors.

53) Bring back the imaging plane to the glass surface using the piezo stage.

54) Record the membrane image again in the green channel only (Camera settings: Exposure time-0.1 s, number of frames-50, camera frame area: 256 pixel x 256 pixel in the green channel).

### Analysis (Timing 15-30 minutes)

This section only sketches the analysis steps, which have been discussed in depth in the Experimental Design section. These analysis steps can be performed using the attached Matlab code, for which operation instructions are provided in the supplementary PDF file ‘Step by step guide to run analysis codes.pdf’.

56) Analyze the SLN movies captured with the Alexa Fluor-647 tagged membrane proteins to obtain localization of individual receptor molecules with sub-pixel resolution.

57) Analyze the VA-TIRFM movie files to reconstruct the membrane topography.

58) Correct results for sample drift during the recording of series of the SLN movies.

59) Merge positions of membrane proteins mapped by SLN imaging with the 3D topography of cell membrane, reconstructed from VA-TIRFM image of cell membrane after correcting registration shift and the effect of sample drift.

60) Determine the distribution of membrane protein with respect to microvilli and cell-body regions. Find out the cumulative fractional increase of the number of molecules of a specific protein on each cell as a function of the distance from the central region of each microvillus and also.

### Troubleshooting

Troubleshooting advice can be found in Table 1.

**Table 1.**
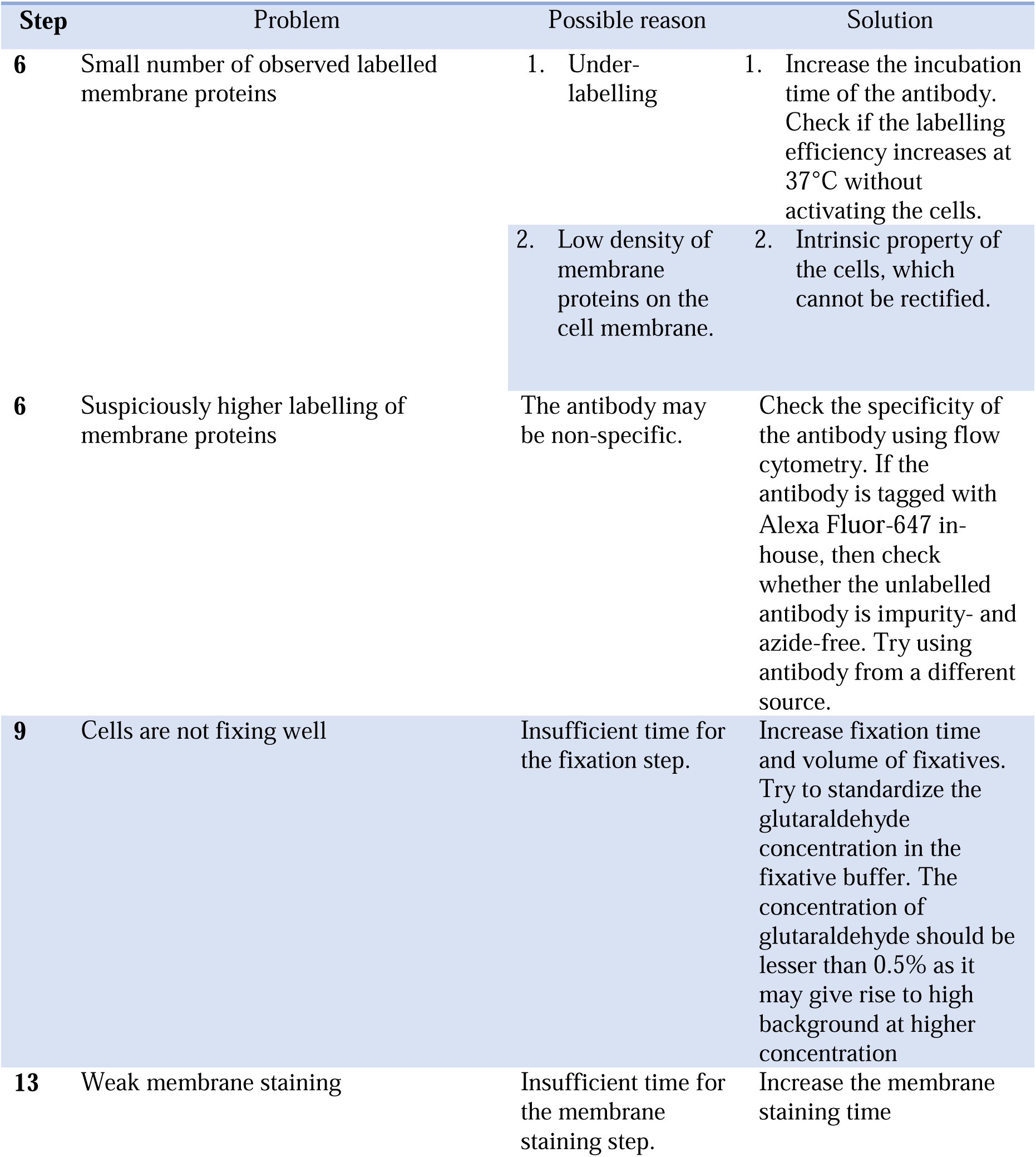
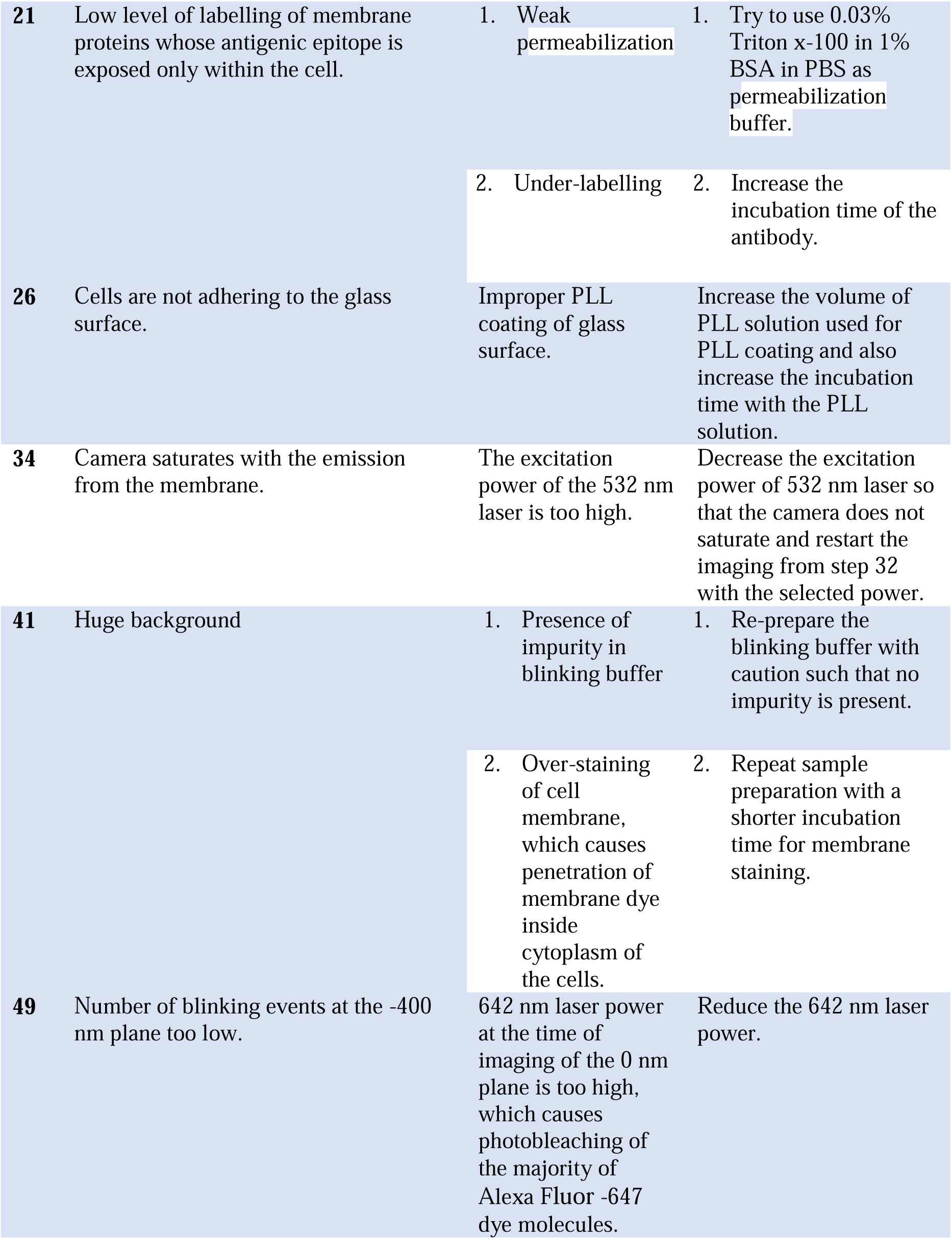
Troubleshooting table.

### Timing

Reagent preparation: 5-6 h.

Steps 1-26: Labelling of cells: 6h-1day. Steps 27-55: Imaging: 1h per cell.

Steps 56-60: Analysis: 15-30 min per cell.

## Supporting information

Analysis MATLAB codes

## Acknowledgements

We thank Dr. Sara W. Feigelson of the Weizmann Institute of Science, Israel and Dr. Yunmin Jung of La Jolla Institute for Immunology, La Jolla, CA, USA for their kind comments and suggestions. G.H. is the incumbent of the Hilda Pomeraniec Memorial Professorial Chair.

## Data availability

The datasets generated during and/or analyzed during the study are available from the corresponding author upon request.

## Code availability

The custom MATLAB code described in this study and provided as supporting information can be accessed and used by readers without restriction.

