## Supplementary Data for "Microvillar cartography: a super-resolution single-molecule imaging method to map the positions of membrane proteins with respect to cellular surface topography"

| *Table S1: Recommended names for saving image files to use our MATLAB code* | | | | |
| --- | --- | --- | --- | --- |
| *Step* | *Type of the Movie Recorded* | *Suggested Format of the Name* | *Example of Name** | *File Format* |
| *35-36* | *VA-TIRFM movie recorded at different angles* | *Samplename_VATIRFM_angle_file-number.fits* | JE2FMTCR_VATIRFM_63_01.fits** | *‘fits’* |
| *38* | *TIRFM image recorded for drift correction of SLN movies at angle 66.8° and 0nm focal plane**** | *Samplename_TIRFM_ planeheight _number.fits* | *JE2FMTCR_TIRFM_0nm_00.fits*  *To*  *JE2FMTCR_TIRFM_0nm_09.fits* | *‘fits’* |
| *41* | *SLN movies recorded at angle 66.8° and 0nm focal plane***** | *Samplename_SLN_ planeheight _number.fits* | *JE2FMTCR_SLN_0nm_00.fits*  *To*  *JE2FMTCR_SLN_0nm_09.fits* | *‘fits’* |
| *46* | *TIRFM image recorded for drift correction of SLN movies at angle 66.8° and -400 nm focal plane****** | *Samplename_TIRFM_ planeheight _number.fits* | *JE2FMTCR_TIRFM_400nm_00.fits*  *To*  *JE2FMTCR_TIRFM_400nm_09.fits* | *‘fits’* |
| *49* | *SLN movies recorded at angle 66.8° and -400 nm focal plane******* | *Samplename_SLN_ planeheight _number.fits* | *JE2FMTCR_SLN_400nm_00.fits*  *To*  *JE2FMTCR_SLN_400nm_09.fits* | *‘fits’* |
| *54* | *TIRFM image recorded at 0nm focal plane and 66.8° after recording the series of SLN movie at -400 nm plane* | *Samplename_TIRFM_ planeheight _number.fits* | *JE2FMTCR_TIRFM_400nm_10.fits* | *‘fits’* |

** Sample Name should not contain any underscore (_ ). In our case, the sample name is* ‘JE2FMTCR’.

**Angle should be rounded down to nearest integer; for example at angle 68.8, the angle should be written as 68.

****Repeated 10 times for 10 SLN movies at 0nm focal plane.*

*****Repeated 10 times to record 30000 frame in total at 0nm focal plane.*

******Repeated 10 times for 10 SLN movies at -400nm focal plane.*

*******Repeated 10 times to record 30000 frame in total at -400nm focal plane.*

**Prerequisites and environment**:

MATLAB R2017a and onward.

**Installing the program:**

1. Place all the MATLAB code files into one folder.
2. Open MATLAB and add the path of this folder to the MATLAB path.

You can do this by selecting the 'Set Path' from the 'HOME' menu of the MATLAB main window.

**Flowchart of Microvilli Cartography analysis:**

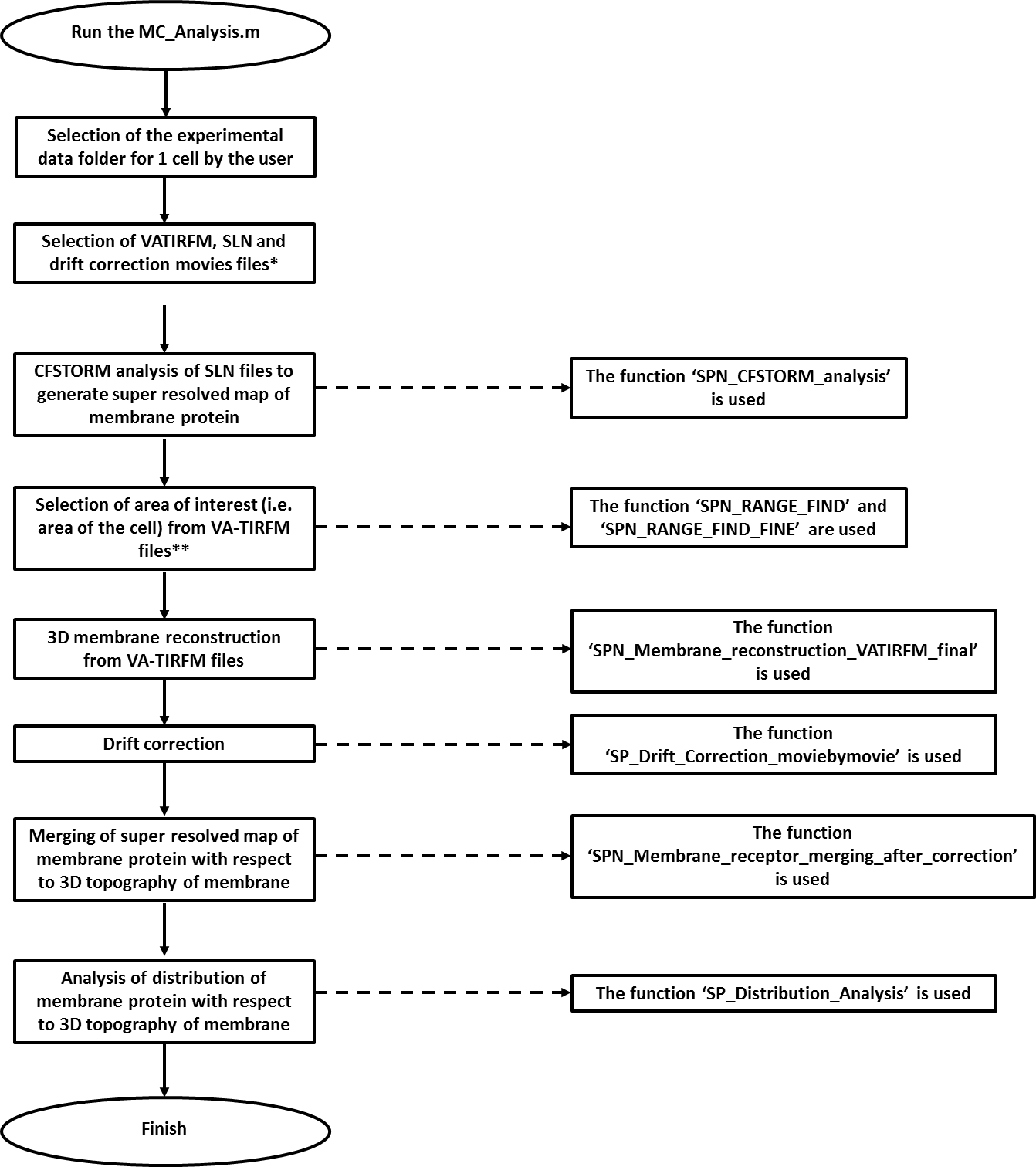

*****All the selections from this point onwards are done automatically.

******The area of interest is also selected automatically, as the rectangular area which encloses the largest connected set of pixels whose z-height is less than or equal to 400 nm according to the calculation from VA-TIRFM images.

**Practical guide to run the MATLAB code**

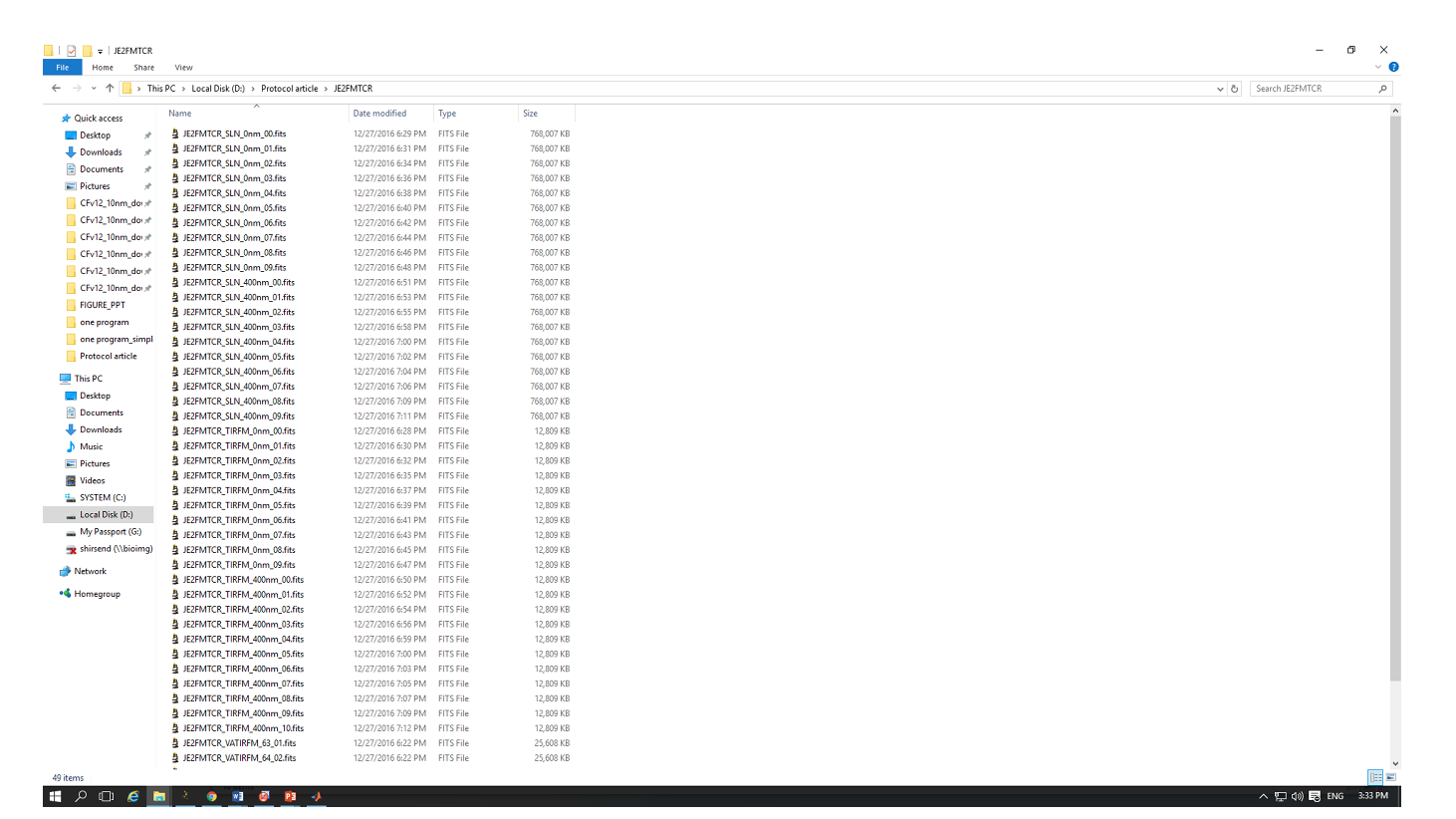
Step 1: Create a folder that contains all the movies recorded for one cell.

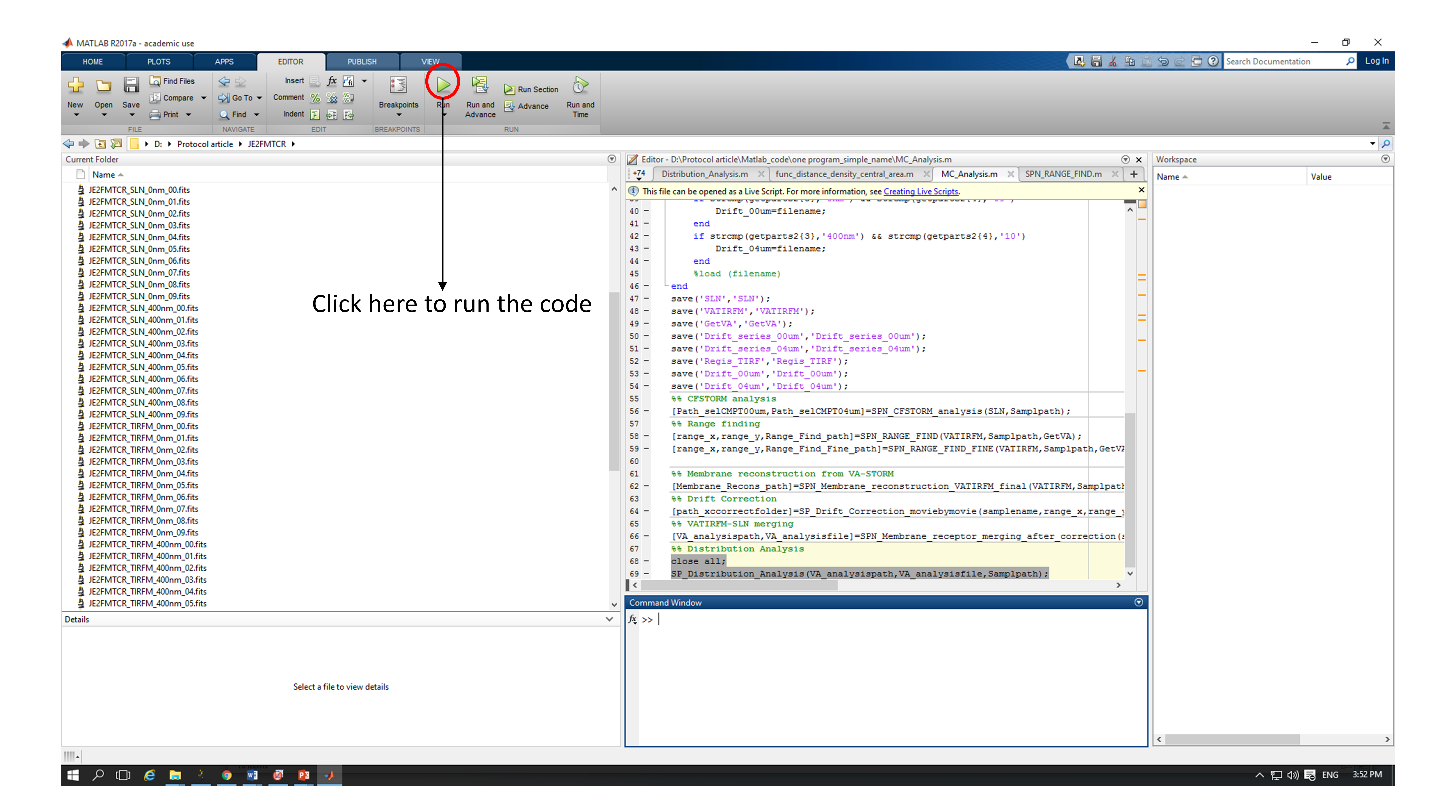
Step 2: Open the *‘****MC_Analysis.m****’* code in MATLAB and run it by clicking on the green button as shown in the figure below.

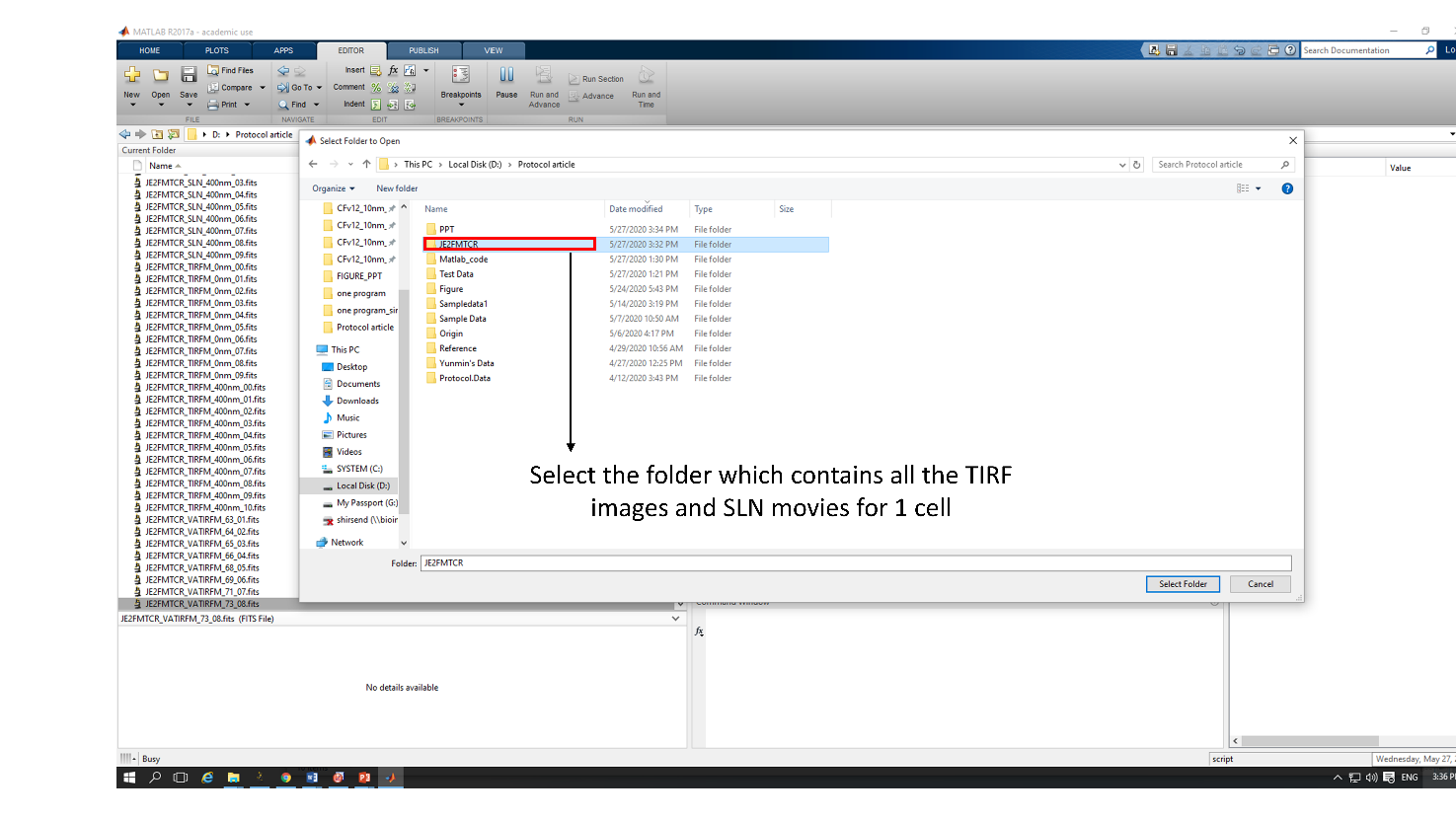
Step 3: Select the folder that contains all the files for one single cell.

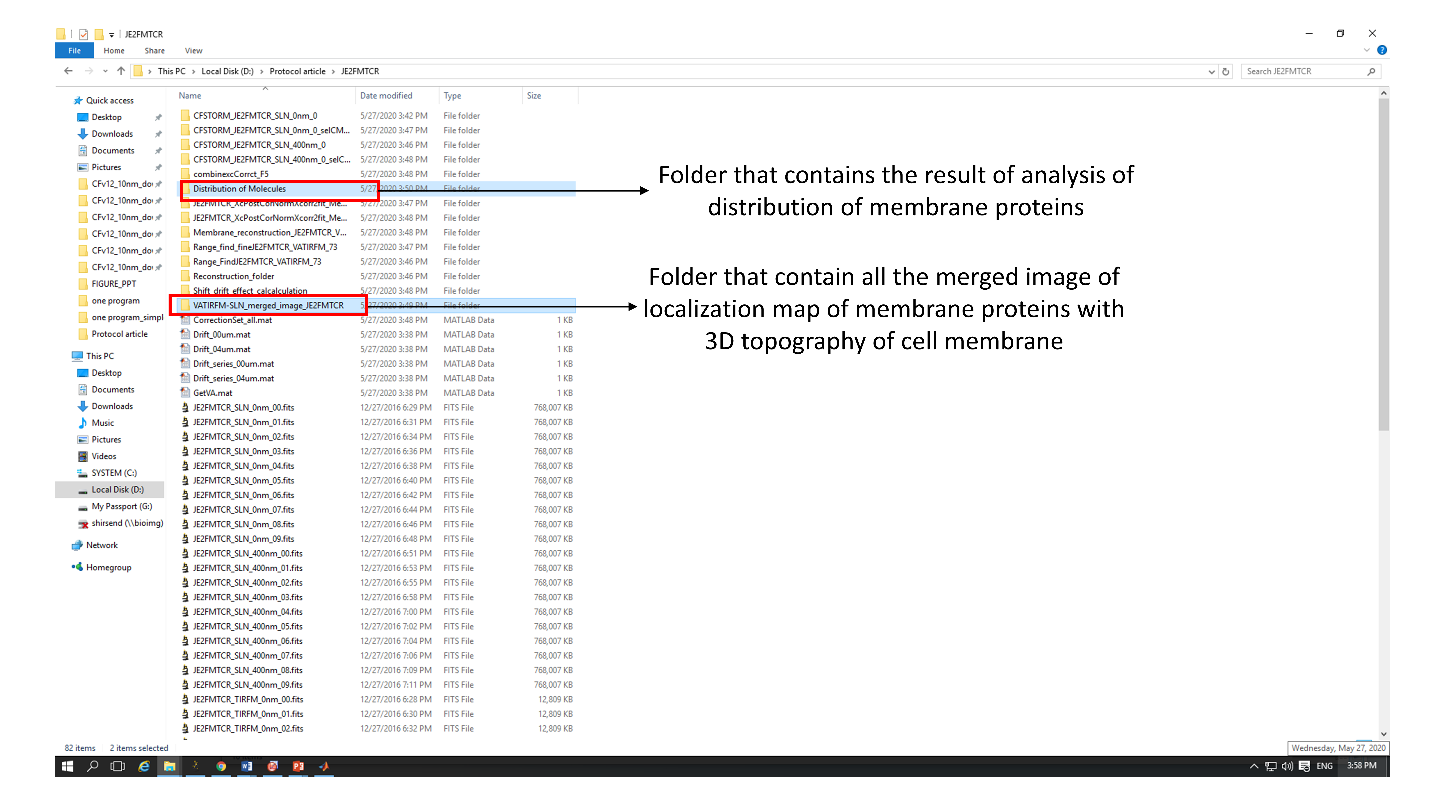
Step 4: The code will create several folders. Among them two folders contains the main results. They are **a. *‘VATIRFM-SLN_merged_image_samplename’*** and **b. ‘Distribution of Molecules’**.

The ‘VATIRFM-SLN_merged_image_samplename’ will contain all the merged image of localization map of membrane proteins with 3D topography of cell membrane (like, Fig. 5J-K).

‘Distribution of Molecules’ will contain the analysis of distribution of membrane protein with respect to membrane topography of the cells. This folder will contains two main files. One is an ‘ascii’ file whose name is *‘delta_count_delta_area’,* which contains the information about cumulative fractional increase of the number of molecules of a specific protein on each cell as a function of the distance from the central region of each microvillus. The first column of the file specifies the distance from the central microvillar region, second column represents the δCount/δArea values for the specific distance for 0nm plane and the third column represents the δCount/δArea values for -400 nm. The corresponding plot is shown in the ‘Delta_Count_Delta_Area_Plot.fig’ file.

The second file is a *‘txt’* file named *‘MV-CB-Sort.txt’*, which tabulates count of protein molecules’ in Microvilli (MV) and CellBody (CB) area and corresponding percentage of molecules in those two areas in both planes.

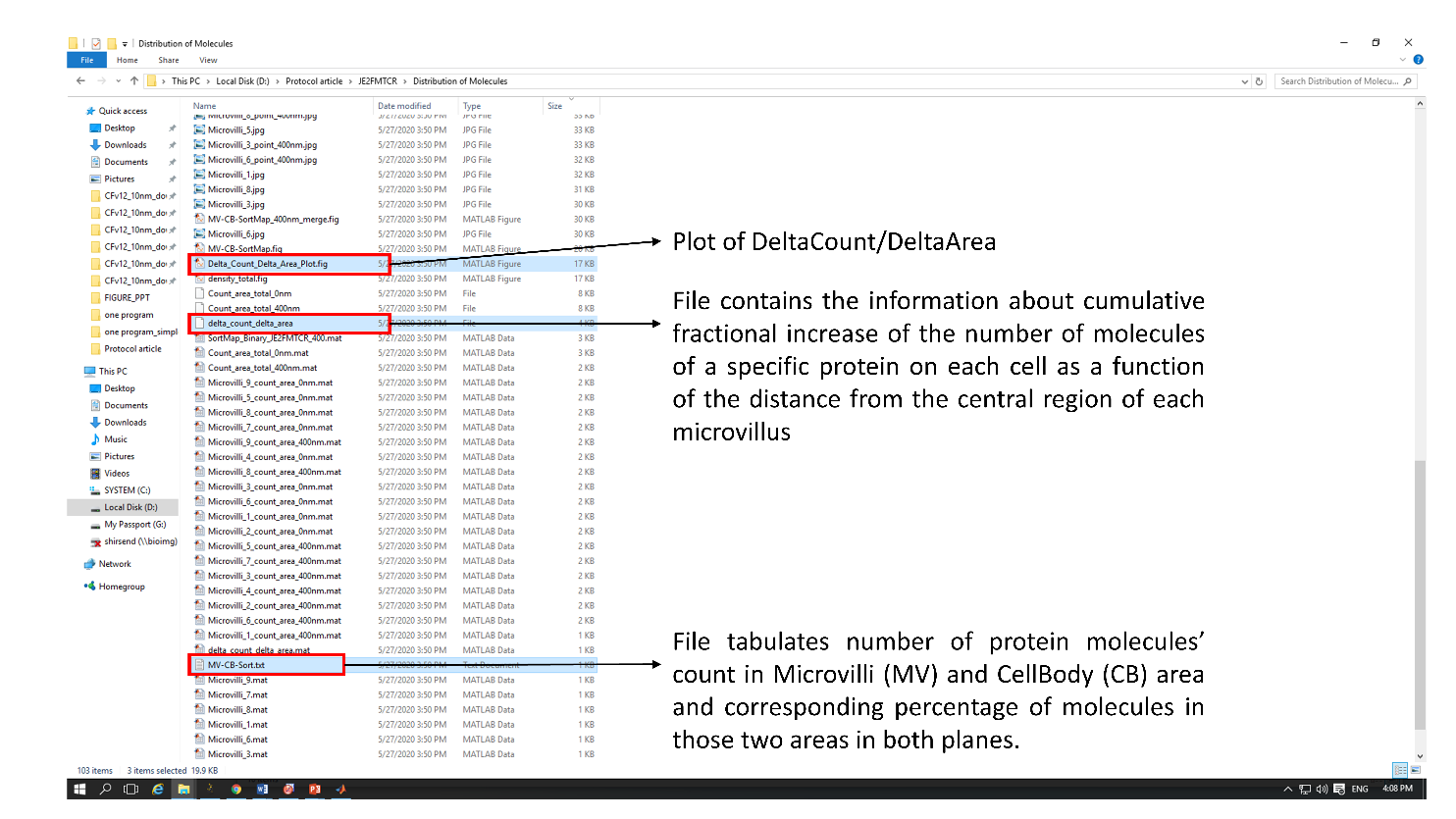
