## Supplementary material for "Microvillar cartography: a super-resolution single-molecule imaging method to map the positions of membrane proteins with respect to cellular surface topography": Analysis MATLAB codes: addscale3_um_range.m

```
function[ret]=addscale3_um_range(rangex,rangey,markersize)
% markersize=100;
% markersize=1000;
% startx=131;
% starty=157;
% onoff=1;
allws=0;
% rangex=[75 175]; 
% rangey=[90 190];
xlim(rangex);ylim(rangey);
%% marker size in namometer ; print as um scale
% markersize=1000; 
if allws==0
%markersize=markersize*1000;
startx=rangex(1)+10;
starty=rangey(1)+10;
onoff=1;
elseif allws==1;
 startx=20;
 starty=30;
 onoff=1;
end
Annotation=1;
 
%function[scalebar]=addscale(markersize,startx,starty,onoff)
%on =1;
%off=0;
hold on;
%markersize= 100; 
%markersize= 100; 
pixelsize=66.67; 

scalerange=markersize/pixelsize;
Markerx=linspace(0,scalerange);
Markerx=startx+Markerx;
Markery=ones(size(Markerx));
Markery=starty*Markery;
scalebar=line(Markerx,Markery,'Color',[0.5 0.5 0.5],'LineWidth',2); 
% set(findobj(gca,'Type','line','Color',[0 0 1]),...
%     'LineWidth',2)
  %  'Color','black',...
% %%line(XData1,YData1,'Parent',Parent1,...
%     'Color',[0.800000011920929 0.800000011920929 0.800000011920929]);
%   
%   
  
if onoff==0;
set(scalebar,'Visible','off');  %# Make it invisible
elseif onoff==1
set(scalebar,'Visible','on');   %# Make it visible
end

if Annotation==1


% Create textbox
annotation(gcf,'textbox',...
    [0.228 0.15 0.07 0.05]...   
    ,'String',{[num2str(markersize/1000) '\mum']},...
    'FontSize',15,...
    'FitBoxToText','off',...
    'LineStyle','none',...
    'LineWidth',20,...
    'Color',[0.5 0.5 0.5]); 
   % 'Color',[0.50 0.50 0.50]); %gray
    %[0.67 0.062 0.07 0.05],...
end

% annotation(figure1,'textbox',...
%     [0.622972972972973 0.0621951219512196 0.07 0.0500000000000001],...
%     'String',{'1000nm'},...
%     'FontSize',16,...
%     'FitBoxToText','off',...
%     'LineStyle','none',...
%     'LineWidth',20,...
%     'Color',[0.5 0.5 0.5]);


hold on
%end


end
```
