## Supplementary material for "Microvillar cartography: a super-resolution single-molecule imaging method to map the positions of membrane proteins with respect to cellular surface topography": Analysis MATLAB codes: CFSTORM_analysis.m

```
%% MainAnalysis program for single molecule detection from multifile
%% Analysis option setting%%
% 1=on  0=off 
clear all;close all;clc;
RecPix      = 5; % Reconstruction of CMMap in original size in nm scale : Rendering images in "nm" scale 
MeanFrames  = 1;  % sum of frame_out (integrating all raw-images)
Applyfilter = 1;
filtertype= 'MeanFilter' ;% 'MedianFilter' 'LogFilter';
kernelsize= 60;
useBgdTH=0;
MeanFrameFilter=1;
selCManalysis=1;
Reconstruction=1;

%% Open files (multiple or single) % choose single or multiple fits files
[FNlist, Path, Filterindex] = uigetfile( ...
{  '*.fits','Fits Files (*.fits)'; ...
   '*.*',  'All Files (*.*)'}, ...
   'Pick a file', ...
   'MultiSelect', 'on');
%% Parameter setting 
tic

% global parameters (doesen't change from movie to movie)
SmTH     = 400   ;    % threshold for determine single molecule for bwlabel (find islands)
MaxTH    = 20000 ;    % mimimum limit of the max value inside the island for single molecule 
MaxPNTH  = 50000 ;    %maximum limit of the Num of photon of the island (ws*ws) discard if Num of Photon is biger than MaxPNTH (maximum limit for photon number threshold) 
MinPNTH  = 1000  ;    % minimum limit of the Num of photon of the island (ws*ws) discard if Num of Photon is smaller than MaxPNTH (maximum limit for photon number threshold) 
ws       = 11    ;    % windowsize(for example 13x13, 7x7, etc,) must be odd number
sigmacm  = 1.5   ;    % sigma for single molecules for 2D fitting; % FWHM = 2.3548200*sigma of single molecules 
sigma    = 1.65  ;    % sigmax: 1.65+0.1; sigmay: 1.65
cmssd    = 0.8   ;    % sigma-standard-deviation foe center of mass
ssd      = 0.5   ;    % sigma-standard-deviation for sm                                       
mMMR     = 0     ;    % Maxpixel/MinValpixel (for center of mass)
mSNR     = 0     ;    % minimum ratio of maxint/minint in the found island
Magnify  = 240   ;    % Magnification of system : 240x 
Pixsize  = 16000 ;    % Camera CCD pixel size in nm
Pixelnm  = round((Pixsize/Magnify)*100)/100  ; %66.6700nm  
% local parameters (Get information from the individual movies)


if   iscell(FNlist)
     NumofFiles=size(FNlist,2);
else
     NumofFiles=1;
end


for ff = 1:NumofFiles

        if   iscell(FNlist)    
             FileName=FNlist{ff};
        else 
             FileName=FNlist;
             
             
             
        end

    PathName=Path;
    SaveFileName = FileName(1:end-5);  
    
   
    SaveFolderName = [SaveFileName(1:end-1) ''];% -6 for _1.txt// -5 for 1.txt
   
        
    
    fitsfilename = [PathName FileName];
    cd (PathName);
    info=fitsinfo(fitsfilename);
    FileDimention=info.PrimaryData.Size;
    FileName %%%  print filename display

    StartFrame   = 1;
    if length(FileDimention)<3; EndFrame = 1;
    else
    EndFrame     =(FileDimention(3)); 
%   EndFrame     = 1000;
    end


        if   isempty(info.PrimaryData.Keywords{30,2})
             EMgain=100;  
        else 
             EMgain=info.PrimaryData.Keywords{30,2};
        end


        if   isempty(info.PrimaryData.Keywords{34,2})
             PreAmp=5;
        else 
             PreAmp=info.PrimaryData.Keywords{34,2};
        end


    %% Data Anaysis  

    % Analysis range (start frame to end frame)

    cnt=0;
    allFrames = zeros(FileDimention(1),FileDimention(2));
    CMoutPT   = []; 
    FitoutPT  = [];
    selFitCMSigmaPT  = [];

 %% Localization Process
 
   for jj = StartFrame:EndFrame;  % FrameN=jj; % Nth Frame for analysis% #ok<ALIGN>
        %% open images from frame-by-frame    
         
         [filesData, frames_out1] = fits_shifter_single_Y_v30(PathName,FileName,jj);  % frames_out ; FrameNth value (:,:,N)
 %% Meanfilter substract 
         if Applyfilter==1
             
            frames_out2=FilterFrameOut_size(frames_out1,filtertype,kernelsize);
         end 
 %% Find Molecules based on the Center-of-mass criteria
         
            [CMout,CMnum,Fitout,Fitnum,selFitCMSigma]=findparticleFrameN(frames_out2,jj,SmTH,MaxTH,MaxPNTH,ws,sigma,ssd,sigmacm,cmssd,EMgain,PreAmp,mMMR,mSNR,MinPNTH);
        
%% create SummaryParticleTables (CMoutPT) from each frames (optional)
         if  CMnum>0  
             [CMoutPT] = ConcatResult_v30(CMoutPT,CMout,1,jj); % 1: for matrix type array 2: cell type array BUGGG need to fix for cell array 
         end

 %% create SummaryParticleTables (CMoutPT) from each frames (optional)
         if  Fitnum>0 % && selCManalysis==0 
             [FitoutPT] = ConcatResult_v30(FitoutPT,Fitout,1,jj); % 1: for matrix type array 2: cell type array
           if  selCManalysis==1 
             [selFitCMSigmaPT] = ConcatResult_v30(selFitCMSigmaPT,selFitCMSigma,1,jj); % 1: for matrix type array 2: cell type array
           end
             %FitoutPT= selFitCMSigmaPT(:,1:10); % modified on 20140323
         end 
             
 %%  create meanFrame Figure (optional)
 
         
            allFrames=allFrames+frames_out2;
        

         clear CMout Fitout selCMout
    end
%% Generate new folders 

NewfolderName=['CFSTORM','_',SaveFolderName];
NewfolderName_selCMPT=['CFSTORM','_',SaveFolderName, '_selCMPT'];
if (exist (NewfolderName)==0)
mkdir(pwd,NewfolderName)
end
Path1=pwd;
if (exist (NewfolderName_selCMPT)==0)
mkdir(pwd,NewfolderName_selCMPT)
cd(NewfolderName_selCMPT)
Path_selCMPT=pwd
cd (Path1);
end
cd (NewfolderName);   
             
%% create meanFrame Figure
            meanFrames = allFrames/jj;

            if MeanFrames == 1
                figure('Name','Meanframes')
                colormap(gray)
                h=imagesc(meanFrames);
                colormap(gray)
                set(gca,'YDir','normal')
                axis equal
                xlim([0 FileDimention(1)]);
                ylim([0 FileDimention(2)]); 
                saveas(gcf,['Mean_',SaveFileName]);%,'jpg');
                save(['Mean_',SaveFileName],'meanFrames');
               
            end
                 clear jj ;

close all        

%% Save Results
% % saveas(SumFigure,['SummaryFigure_',SaveFileName,]);%,'jpg');
% save(['CMPT_',SaveFileName],'CMoutPT');
% save(['FitPT_',SaveFileName'],'FitoutPT');    
if length(SaveFolderName)>64
filenth=FileName(end-6:end-5);% type char
SaveFileName=[FileName(1:60),filenth];
end

if ~isempty(CMoutPT)
save(['CMPT_',SaveFileName,'.ascii'],'CMoutPT','-ascii')
save(['CMPT_',SaveFileName],'CMoutPT');
end

if ~isempty(FitoutPT)
save(['FitPT_',SaveFileName,'.ascii'],'FitoutPT','-ascii')
save(['FitPT_',SaveFileName],'FitoutPT');
end

if ~isempty(FitoutPT) && selCManalysis==1
    Path_org=pwd;
save(['selCMPT_',SaveFileName,'.ascii'],'selFitCMSigmaPT','-ascii')
save(['selCMPT_',SaveFileName],'selFitCMSigmaPT');
cd (Path_selCMPT)
save(['selCMPT_',SaveFileName],'selFitCMSigmaPT');
cd (Path_org)
end
    

%% Save parameters
Parameter.SmTH                  = SmTH ;
Parameter.MaxTH                 = MaxTH;
Parameter.MaxPNTH               = MaxPNTH;
Parameter.MinPNTH               = MinPNTH;
Parameter.ws                    = ws;
Parameter.sigmax                = sigma;
Parameter.sigmay                = sigma+0.1;
Parameter.ssd                   = ssd;
Parameter.sigmacm               = sigmacm;
Parameter.cmssd                 = cmssd;
Parameter.mimRatioMaxMin        = mMMR;
Parameter.minRatioMaxphoMinpho  = mSNR;
Parameter.FileDimention         = FileDimention;
Parameter.Magnification         = Magnify;
Parameter.PixSize               = Pixsize;
Parameter.Pixelnm               = Pixelnm;
Parameter.RecPix                = RecPix;
Parameter.SaveFileFolder        = SaveFolderName;
Parameter.SaveFileName          = SaveFileName;
Parameter.FilterType            = filtertype;  
Parameter.KernelSize            = kernelsize;  
Parameter.MeanFrames            = MeanFrames;  % sum of frame_out
Parameter.ApplyFilter           = Applyfilter;
Parameter.useBgdTH              = useBgdTH;
Parameter.selCManalysis         = selCManalysis; %%%%% 
if length(SaveFolderName)<64
save(['Parameter_',SaveFolderName],'Parameter');
else 
save(['Parameter_CMFIT'],'Parameter');
end

%% Calculate means of the results
MeanCM= zeros(1,10);
STDCM= zeros(1,10);

MeanFit= zeros(1,10);
STDFit= zeros(1,10);   
% MeanselCM= zeros(1,10);   
% STDselCM= zeros(1,10);   

if size(CMoutPT,1)==1 
    MeanCM=CMoutPT; 
end

if size(FitoutPT,1)==1 
    MeanFit=FitoutPT;
end

if ~isempty (CMoutPT) && size(CMoutPT,1)>1
MeanCM=mean(CMoutPT);
STDCM=std(CMoutPT);
end

if ~isempty (FitoutPT) && selCManalysis==0
MeanFit=mean(FitoutPT);
STDFit=std(FitoutPT);  
elseif  ~isempty (FitoutPT) && selCManalysis==1
MeanFit= zeros(1,12);
STDFit= zeros(1,12);     
MeanFit=mean(selFitCMSigmaPT);
STDFit=std(selFitCMSigmaPT);  
elseif isempty (FitoutPT) && selCManalysis==1
MeanFit= zeros(1,12);
STDFit= zeros(1,12);  
end
% end
%% Save parameters


Summary.CMnum                          = size(CMoutPT,1);
Summary.mean_std_CM_sigmax             = [MeanCM(1,4) STDCM(1,4)];
Summary.mean_std_CM_sigmay             = [MeanCM(1,5) STDCM(1,5)];
Summary.mean_std_CM_ratioMaxMin        = [MeanCM(1,8) STDCM(1,8)];
Summary.mean_std_CM_numofPho           = [MeanCM(1,10) STDCM(1,10)];
if selCManalysis==0
Summary.Fitnum                         = size(FitoutPT,1);
Summary.mean_std_Fit_sigmax            = [MeanFit(1,4) STDFit(1,4)];
Summary.mean_std_Fit_sigmay            = [MeanFit(1,5) STDFit(1,5)];
Summary.mean_std_Fit_ratioAmpBac       = [MeanFit(1,8) STDFit(1,8)];
Summary.mean_std_Fit_numofPho          = [MeanFit(1,10) STDFit(1,10)];
elseif selCManalysis==1                   
Summary.Fitnum                         = size(selFitCMSigmaPT,1);
if (size(MeanFit,2)>1)
Summary.mean_std_Fit_sigmax            = [MeanFit(1,4) STDFit(1,4)];
Summary.mean_std_Fit_sigmay            = [MeanFit(1,5) STDFit(1,5)];
Summary.mean_std_Fit_ratioAmpBac       = [MeanFit(1,8) STDFit(1,8)];
Summary.mean_std_Fit_numofPho          = [MeanFit(1,10) STDFit(1,10)];
Summary.mean_std_CM2_sigmax             = [MeanFit(1,11) STDFit(1,11)];
Summary.mean_std_CM2_sigmay             = [MeanFit(1,12) STDFit(1,12)];
%Summary.mean_std_CM2_ratioMaxMin        = [MeanselCM(1,8) STDselCM(1,8)];
%Summary.mean_std_CM2_numofPho           = [MeanselCM(1,10) STDselCM(1,10)];
end
end

if length(SaveFolderName)<64 || length (SaveFileName)<64
save(['smr_',SaveFileName],'Summary');  
else 
save(['Smr',SaveFileName(1:50)],'Summary');
end

%% Reconstruction for super resolution
            Reconstruction_ver10_sigmacolor(CMoutPT,selFitCMSigmaPT,Parameter,SaveFileName); %% (CMPT, FITPT, Parameter, SaveFilename, RecPix) if you change the save filename, make it empty 
            close all; 
       
end
cd (Path);
if (exist ('Reconstruction_folder')==0)
mkdir(pwd,'Reconstruction_folder');
cd ('Reconstruction_folder');
Path_Reconstruction_folder=pwd;
cd (Path);
end
ListChildFolder= dir;
for ff= 1:size(ListChildFolder,1)
   if ListChildFolder(ff).isdir==1
   if ~isempty(strfind(ListChildFolder(ff).name,'CFSTORM'))&&isempty(strfind(ListChildFolder(ff).name,'selCMPT')) 
       ChildPath=[ListChildFolder(ff).name];
   getparts10 = strread(ChildPath,'%s','delimiter','_');
   savefilesname=[getparts10{2:end-1}];
   cd (ChildPath);
   pwd
   path = [pwd '\']; 
        Listing_par = dir([path,'*.mat']);
        if size(Listing_par,1)>0
        for nn= 1:size(Listing_par,1)
        filename_par= Listing_par(nn).name;
        if~isempty (strfind(filename_par,'Parameter'))
        Parameter=load(filename_par);
        end
        end
        end
 %       path = filepath
        Listing = dir([path,'*.mat']);
        if size(Listing,1)>0
        for nn= 1:size(Listing,1)
           
            CMoutPT={};
            FitoutPT={};
            filename= Listing(nn).name;
            getparts = strread(filename,'%s','delimiter','.');
            getparts2 = strread(getparts{1},'%s','delimiter','_');
            number=getparts2{end};
                        
            load([path filename])
            if (strcmp(getparts2{1},'CMPT')==1)
            CMoutPT(:,11)=str2num(number);
            end
            if (strcmp(getparts2{1},'FitPT')==1)
            FitoutPT(:,11)=str2num(number);
            end
            CMsetPTc(nn)={CMoutPT};
            FitsetPTc(nn)={FitoutPT};
            %Seris= load([path filename]);%,'-mat','CMoutPT'
            % Xcoor=Seris(:,2);
            % % Ycoor=Seris(:,3);
            % MovieFiles(nn)={[Xcoor Ycoor]};
        end

              
        %%% Combine results from each movies

        CMsetPTc=CMsetPTc(~cellfun('isempty',CMsetPTc));   % Remove empty cells
        CMsetPT=cell2mat(CMsetPTc');

        FitsetPTc=FitsetPTc(~cellfun('isempty',FitsetPTc));  % Revmove empty cels
        FitsetPT=cell2mat(FitsetPTc');
        path_now=pwd;
        FileDimention = Parameter.FileDimention;
Magnify       = 240   ;    % Magnification of system : 240x 
Pixsize       = 16000  ;    % Camera CCD pixel size in nm
Pixelnm       = round((Pixsize/Magnify)*100)/100  ; %66.6700nm  
Rend          = (round(RecPix/Pixelnm*100))/100;     % Rend: rendering pixels for desired size (example 1pixel is 66.67nm, 66,67/Rend=desired pixel size(nm))  

if size (CMsetPT,1)>0 
            CMfig=figure('Name','CM_Reconstruction');
            CM=CMsetPT(:,2);
            CN=CMsetPT(:,3);
            %axis equal
            plot(CN,CM,'.','MarkerSize',Rend);  %% Don't know why 
            axis equal
            xlim([0 FileDimention(2)]);
            ylim([0 FileDimention(1)]);      
end

if size (FitsetPT,1)>0 
             Fitfig=figure('Name','Fit_Reconstruction');
             FM=FitsetPT(:,2);
             FN=FitsetPT(:,3);
             %axis equal
             plot(FN,FM,'.','MarkerSize',Rend,'Color','red');   %% Don't know why 
             axis equal
             xlim([0 FileDimention(2)]);
             ylim([0 FileDimention(1)]); 
end

if size (FitsetPT,1)>0 && size(CMsetPT,1)>0 
 MerFig =figure('Name','CMFit_mergeReconstruction');
             CM=CMsetPT(:,2);
             CN=CMsetPT(:,3);
             plot(CN,CM,'.','MarkerSize',Rend); 
             axis equal
             xlim([0 FileDimention(2)]);
             ylim([0 FileDimention(1)]);              
            % xlim([1 FileDimention(1)]);
            % ylim([1 FileDimention(2)]); 
             hold on   
             FM=FitsetPT(:,2);
             FN=FitsetPT(:,3);
             %axis equal
             plot(FN,FM,'.','MarkerSize',Rend,'Color','red');   
end 
        cd (Path_Reconstruction_folder)
            save(['CMsetPT_',savefilesname,'.ascii'],'CMsetPT','-ascii')
            save(['CMsetPT_',savefilesname],'CMsetPT');
            save(['FitsetPT_',savefilesname,'.ascii'],'FitsetPT','-ascii')
            save(['FitsetPT_',savefilesname],'FitsetPT');
            saveas(CMfig,['CMrec_',savefilesname]); 
            saveas(Fitfig,['Fitrec_',savefilesname]);
            saveas(MerFig,['MerFig_',savefilesname]);
        cd (path_now);
            

     
       
        else
        clear ('filename','path') 
        
        pwd
        cd (Path) %%%%%
        %%
        end
        end
   end
   cd (Path)
end
fprintf('done\n')
toc


%% CMoutSM
% 1. frameN
% 2: xcm
% 3: ycm
% 4: sxcm
% 5: sycm
% 6: Lmax
% 7: Lmin
% 8: SNR
% 9: Qm
% 10 NumPho

%% FitoutSM
% 1. frameN
% 2: xfit
% 3: yfit
% 4: sxfit
% 5: syfit
% 6: Amp
% 7: Bgd
% 8: SNR
% 9: AveRErr; (Average Ratio of error/fittevalue ; 95% confidence)
% 10 NumPho
% option 1(nonaccumulate) and 2(accumulatedate) for fitting the ws*ws 
```
