## Supplementary material for "Microvillar cartography: a super-resolution single-molecule imaging method to map the positions of membrane proteins with respect to cellular surface topography": Analysis MATLAB codes: Combined_main_program_simplified.m

```
%% one program
clc; clear all; close all;
folder_data = uigetdir;
Samplpath = [folder_data '\'];
Listing = dir([Samplpath,'*.fits']);
cd (Samplpath)
ii=1;
jj=1;
kk=1;
ll=1;
for nn= 1:size(Listing,1)
    filename= Listing(nn).name;
    getparts1 = strread(filename,'%s','delimiter','.');
    getparts2 = strread(getparts1{1},'%s','delimiter','_');
    samplename=[getparts2{1}];
    if strcmp(getparts2{2},'SLN')
        SLN{ii}=filename;
        ii=ii+1;
    end
    if strcmp(getparts2{2},'VATIRFM')
        VATIRFM{jj}=filename;
        GetVA(jj)=str2num(getparts2{3});
        jj=jj+1;
    end
    if strcmp(getparts2{3},'0nm') && strcmp(getparts2{2},'TIRFM') 
        Drift_series_00um{kk}=filename;
        kk=kk+1;
    end
    if strcmp(getparts2{3},'400nm') && strcmp(getparts2{2},'TIRFM')
        tf=strcmp(getparts2{4},'10');
        if(tf==0)
        Drift_series_04um{ll}=filename;
        ll=ll+1;
        end
    end
    if strcmp(getparts2{3},'66')
        Regis_TIRF=filename;
    end  
    if strcmp(getparts2{3},'0nm') && strcmp(getparts2{4},'09') 
        Drift_00um=filename;
    end
    if strcmp(getparts2{3},'400nm') && strcmp(getparts2{4},'10') 
        Drift_04um=filename;
    end
    %load (filename)
end
save('SLN','SLN');
save('VATIRFM','VATIRFM');
save('GetVA','GetVA');
save('Drift_series_00um','Drift_series_00um');
save('Drift_series_04um','Drift_series_04um');
save('Regis_TIRF','Regis_TIRF');
save('Drift_00um','Drift_00um');
save('Drift_04um','Drift_04um');
%% CFSTORM analysis
[Path_selCMPT00um,Path_selCMPT04um]=SPN_CFSTORM_analysis(SLN,Samplpath);
%% Range finding
[range_x,range_y,Range_Find_path]=SPN_RANGE_FIND(VATIRFM,Samplpath,GetVA);
[range_x,range_y,Range_Find_Fine_path]=SPN_RANGE_FIND_FINE(VATIRFM,Samplpath,GetVA,range_x,range_y);

%% Membrane reconstruction from VA-STORM 
[Membrane_Recons_path]=SPN_Membrane_reconstruction_VATIRFM_final(VATIRFM,Samplpath,GetVA,range_x,range_y);
%% Drift Correction
[path_xccorrectfolder]=SP_Drift_Correction_moviebymovie(samplename,range_x,range_y,Drift_series_00um,Samplpath,Path_selCMPT00um,Drift_series_04um,Samplpath,Path_selCMPT04um);
%% VATIRFM-SLN merging
[VA_analysispath,VA_analysisfile]=SPN_Membrane_receptor_merging_after_correction(samplename,range_x,range_y,Regis_TIRF,Drift_00um,Drift_04um,Samplpath,Membrane_Recons_path,path_xccorrectfolder);
%% Distribution Analysis
close all;
SP_Distribution_Analysis(VA_analysispath,VA_analysisfile,Samplpath);
```
