## Supplementary material for "Microvillar cartography: a super-resolution single-molecule imaging method to map the positions of membrane proteins with respect to cellular surface topography": Analysis MATLAB codes: ConcatResult_v30.m

```
function [Con_out] = ConcatResult_v30(Con_in,smoutin,Method,jjj)
% create particle tables by concaternating the results from cmout or fitout

if isempty(Method); % default if matrix type
      Method=1; 
end

    switch Method
        case 1
                   Con_out = vertcat(Con_in,smoutin');

        case 2

                   Con_out{jjj} = smoutin';   
        otherwise 
            error('unexpected Method(choose--> 1:matrix; 2: Cell)')

    end

end
```
