## Supplementary material for "Microvillar cartography: a super-resolution single-molecule imaging method to map the positions of membrane proteins with respect to cellular surface topography": Analysis MATLAB codes: createfigure_subplot2type_Roi.m

```
function createfigure_subplot2type_Roi(data1,data2,whichtype,title1,title2,range_x,range_y)
switch whichtype
    case 1 % surf

% Create figure
figure1 = figure;
%hold on 
subplot1=subplot(1,2,1,'Parent',figure1);
view(subplot1,[90 0]);
grid(subplot1,'on');
hold(subplot1,'all');
surf(data1,'Parent',subplot1);
xlim([range_x]);
ylim([range_y]);
title(title1,'Fontsize',12);
% [OriZLim]=get(subplot1,'ZLim');

% zlabel('\delta z (nm)')
% Create subplot
subplot2 = subplot(1,2,2,'Parent',figure1); 
view(subplot2,[90 0]); %[-80 3]]
grid(subplot2,'on');
hold(subplot2,'all');
% Create subplot
surf(data2,'Parent',subplot2);
xlim(range_x);
ylim(range_y);
title(title2,'Fontsize',12);

% zlim(OriZLim)
% zlabel('\delta z (nm)')
hold off

    case 2
figure1 = figure;
%hold on 
subplot1=subplot(1,2,1,'Parent',figure1);
%view(subplot1,[90 0]);
grid(subplot1,'on');
%hold(subplot1,'all');
imagesc(data1,'Parent',subplot1);

set(gca,'YDir','normal')
title(title1,'Fontsize',12);
xlim(range_x);
ylim(range_y);
axis equal tight;
% [OriZLim]=get(subplot1,'ZLim');
% zlim(OriZLim)
%zlabel('\delta z (nm)')
% Create subplot
subplot2 = subplot(1,2,2,'Parent',figure1); 
%view(subplot2,[90 0]); %[-80 3]]
grid(subplot2,'on');
%hold(subplot2,'all');
% Create subplot
%colormap(prism)
imagesc(data2,'Parent',subplot2);
set(gca,'YDir','normal')

title(title2,'Fontsize',12);
xlim(range_x);
ylim(range_y);
axis equal tight;
% zlim(OriZLim)
% zlabel('\delta z (nm)')
hold off
end

end


% Create surf
```
