## Supplementary material for "Microvillar cartography: a super-resolution single-molecule imaging method to map the positions of membrane proteins with respect to cellular surface topography": Analysis MATLAB codes: createfigure_subplots6_Roi.m

```
function createfigure_subplots6_Roi(multidata,Thr,GetVA,range_x,range_y)
% z
%CREATEFIGURE(ZDATA1)
%  ZDATA1:  surface zdata

%  Auto-generated by MATLAB on 10-Oct-2012 10:58:20

% Create figure
% figure1 = figure;
% %hold on 
% subplot1=subplot(2,3,1,'Parent',figure1);
% view(subplot1,[90 0]);
% grid(subplot1,'on');
% hold(subplot1,'all');
% surf(data1,'Parent',subplot1);
% title(['LogFilter',num2str(GetVA(1))],'Fontsize',12);
% [OriZLim]=get(subplot1,'ZLim');
% zlim(OriZLim)
% %zlabel('\delta z (nm)')
% % Create subplot
% subplot2 = subplot(3,3,2,'Parent',figure1); 
% view(subplot2,[90 0]); %[-80 3]]
% grid(subplot2,'on');
% hold(subplot2,'all');
% % Create subplot
% surf(data2,'Parent',subplot2);
% title(['LogFilter',num2str(GetVA(end))],'Fontsize',12);
% zlim(OriZLim)
% % zlabel('\delta z (nm)')

if isequal(length(GetVA),size(multidata,3));
    num=length(GetVA);
else 
    error('error number of GetVA and num of delta')
end

if num==1;
    a=1;
    b=1;
elseif num==2 || num==3;
    a=1; b=num;
elseif num>3 && num<7;
    a=2;
    b=3;
elseif num>=7 && num<=9;
    a=3;
    b=3;
elseif num==10;
    a=2;
    b=5;
    
else
    error('too large stack')
end

figure1 = figure;
for k=1:num
Anglelabel=num2str(GetVA(k));
subplot3 = subplot(a,b,k,'Parent',figure1);
% grid(subplot3,'off');
hold(subplot3,'all');
imagesc(multidata(:,:,k),'Parent',subplot3);title(Anglelabel);
title(['Thr' num2str(Thr) ' ' num2str(GetVA(k))],'Fontsize',10);
colormap(gray(2))
axis equal
xlim([range_x]);%
ylim([range_y]);%
end
hold off
% saveas(gcf,'SimulationFigures')
end


% Create surf
```
