## Supplementary material for "Microvillar cartography: a super-resolution single-molecule imaging method to map the positions of membrane proteins with respect to cellular surface topography": Analysis MATLAB codes: createfigure_subplots6_surf_Roi.m

```
function createfigure_subplots6_surf_Roi(multidata,zlimit,GetVA,ws,range_x,range_y)
% 
%CREATEFIGURE(ZDATA1)
%  ZDATA1:  surface zdata

%  Auto-generated by MATLAB on 10-Oct-2012 10:58:20

% Create figure
% figure1 = figure;
% %hold on 
% subplot1=subplot(2,3,1,'Parent',figure1);
% view(subplot1,[90 0]);
% grid(subplot1,'on');
% hold(subplot1,'all');
% surf(data1,'Parent',subplot1);
% title(['LogFilter',num2str(GetVA(1))],'Fontsize',12);
% [OriZLim]=get(subplot1,'ZLim');
% zlim(OriZLim)
% %zlabel('\delta z (nm)')
% % Create subplot
% subplot2 = subplot(3,3,2,'Parent',figure1); 
% view(subplot2,[90 0]); %[-80 3]]
% grid(subplot2,'on');
% hold(subplot2,'all');
% % Create subplot
% surf(data2,'Parent',subplot2);
% title(['LogFilter',num2str(GetVA(end))],'Fontsize',12);
% zlim(OriZLim)
% % zlabel('\delta z (nm)')
ws=size(multidata,1);
[x,y] = meshgrid(1:ws);

if isequal(length(GetVA),size(multidata,3));
    num=length(GetVA);
else 
    error('error number of GetVA and num of delta')
end

if num==1;
    a=1;
    b=1;
elseif num==2 || num==3;
    a=1; b=num;
elseif num>3 && num<7;
    a=2;
    b=3;
elseif num>=7 && num<=9;
    a=3;
    b=3;
elseif num==10;
    a=2;
    b=5;
else
    error('too large stack')
    
end

figure1 = figure;
set(gcf, 'Position', [10 20 1200 800]);
for k=1:num
Anglelabel=num2str(GetVA(k));
subplot3 = subplot(a,b,k,'Parent',figure1);
hold(subplot3,'all');
%axes1 = axes('Parent',subplot3);
surf(x,y,multidata(:,:,k),'EdgeColor','none','Parent',subplot3);title(Anglelabel);
% [OriZLim]=get(subplot3,'ZLim');
 xlim([range_x]);
 ylim([range_y]);
 zlim([0 zlimit]);
 zlabel('\delta z (nm)')
title([num2str(GetVA(k))],'Fontsize',10);
colormap('gray');
view(subplot3,[0 0]);
%axis equal
end
hold off
% saveas(gcf,'SimulationFigures')
end

% figure1 = figure;
% colormap('gray');

% Create axes
% axes1 = axes('Parent',figure1);
% view(axes1,[83 12]);
% grid(axes1,'on');
% hold(axes1,'all');
% 
% % Create surf
% surf(x,y,z,'EdgeColor','none')
% axis([1 ws 1 ws 0 zheight]);title(num2str(ta(nr)),'FontSize',20);
% zlabel('\delta Z (nm)','FontSize',16);
% xlabel('x(pixel)','FontSize',16);
% ylabel('y(pixel)','FontSize',16);


% Create surf
```
