## Supplementary material for "Microvillar cartography: a super-resolution single-molecule imaging method to map the positions of membrane proteins with respect to cellular surface topography": Analysis MATLAB codes: DensityInOut_v3_final.m

```
%% Define Cell area and indexing pixels
% from TIRF images ; (using the biggest angle TIRF image)
% edit from default value for the cell body threshod from 400nm to
% CellbodyLimit using an extra input; 150828 

function[Result_DensityInOut,SortMap_Binary]=DensityInOut_v3_final(MeanSTDdelta,LocMicrovilicolor,RoiFitsetPT1,RoiFitsetPT2,range_x,range_y,samplename,CellbodyLimit)
%% Define a cell area
AA=MeanSTDdelta{1,1};

[IndCell]=find(AA<CellbodyLimit);
AA(IndCell)=-1;           
AA(AA>0)=0;
[row_MV,col_MV]=find(LocMicrovilicolor>0);
for MV_r=1:size(row_MV)
    if((col_MV(MV_r)-1)>=range_y(1))&& ((col_MV(MV_r)-1)<=range_y(2))
    LocMicrovilicolor(row_MV(MV_r),(col_MV(MV_r)-1))=LocMicrovilicolor(row_MV(MV_r),(col_MV(MV_r)-1))+2;
    end
    if((col_MV(MV_r)+1)>=range_y(1))&& ((col_MV(MV_r)+1)<=range_y(2))
    LocMicrovilicolor(row_MV(MV_r),(col_MV(MV_r)+1))=LocMicrovilicolor(row_MV(MV_r),(col_MV(MV_r)+1))+2;
    end
    if((col_MV(MV_r)-1)>=range_y(1))&& ((col_MV(MV_r)-1)<=range_y(2))&&((row_MV(MV_r)-1)>=range_x(1))&&((row_MV(MV_r)-1)>=range_x(2))
    LocMicrovilicolor((row_MV(MV_r)-1),(col_MV(MV_r)-1))=LocMicrovilicolor((row_MV(MV_r)-1),(col_MV(MV_r)-1))+2;
    end
    if((col_MV(MV_r)+1)>=range_y(1))&& ((col_MV(MV_r)+1)<=range_y(2))&&((row_MV(MV_r)+1)>=range_x(1))&&((row_MV(MV_r)+1)>=range_x(2))
    LocMicrovilicolor((row_MV(MV_r)+1),(col_MV(MV_r)+1))=LocMicrovilicolor((row_MV(MV_r)+1),(col_MV(MV_r)+1))+2;
    end
    if((col_MV(MV_r)-1)>=range_y(1))&& ((col_MV(MV_r)-1)<=range_y(2))&&((row_MV(MV_r)+1)>=range_x(1))&&((row_MV(MV_r)+1)>=range_x(2))
    LocMicrovilicolor((row_MV(MV_r)+1),(col_MV(MV_r)-1))=LocMicrovilicolor((row_MV(MV_r)+1),(col_MV(MV_r)-1))+2;
    end
    if((col_MV(MV_r)+1)>=range_y(1))&& ((col_MV(MV_r)+1)<=range_y(2))&&((row_MV(MV_r)-1)>=range_x(1))&&((row_MV(MV_r)-1)>=range_x(2))
    LocMicrovilicolor((row_MV(MV_r)-1),(col_MV(MV_r)+1))=LocMicrovilicolor((row_MV(MV_r)-1),(col_MV(MV_r)+1))+2;
    end
    if((row_MV(MV_r)-1)>=range_x(1))&&((row_MV(MV_r)-1)>=range_x(2))
    LocMicrovilicolor((row_MV(MV_r)-1),col_MV(MV_r))=LocMicrovilicolor((row_MV(MV_r)-1),col_MV(MV_r))+2;
    end
    if((row_MV(MV_r)+1)>=range_x(1))&&((row_MV(MV_r)+1)>=range_x(2))
    LocMicrovilicolor((row_MV(MV_r)+1),col_MV(MV_r))=LocMicrovilicolor((row_MV(MV_r)+1),col_MV(MV_r))+2;
    end
end
Map=LocMicrovilicolor;
[MV_ind]=find(Map > 0);

Map(MV_ind)= Map(MV_ind)+1; %% preventing removing the first microvilli
BB=AA+Map; %
%Limit DATA within ROI
Q=zeros(size(BB));
Q(range_y(1):range_y(2),range_x(1):range_x(2))=1;
BB=BB.*Q;

%% Define areas of cell body and microvill

[IndCellBody]=find(BB<0);        % Negative Values (-1) indicate cell area
[IndVill]=find(BB>0);            % Positive Values indicate Microvill Area
[IndBackground]=find(BB==0);     % 0 values indicate background Area

SortMap_Binary=BB;         
SortMap_Binary(IndVill)=1;
%% Define a cell area using cell body limit 250
%CellbodyLimit=312;
AA=MeanSTDdelta{1,1};

[IndCell]=find(AA<CellbodyLimit);
AA(IndCell)=-1;           
AA(AA>0)=0;
Map=LocMicrovilicolor;
[MV_ind]=find(Map > 0);

Map(MV_ind)= Map(MV_ind)+1; %% preventing removing the first microvilli
BB=AA+Map; %
%Limit DATA within ROI
Q=zeros(size(BB));
Q(range_y(1):range_y(2),range_x(1):range_x(2))=1;
BB=BB.*Q;

%% Define areas of cell body and microvill

[IndCellBody]=find(BB<0);        % Negative Values (-1) indicate cell area
[IndVill]=find(BB>0);            % Positive Values indicate Microvill Area
[IndBackground]=find(BB==0);     % 0 values indicate background Area

%% Pixelize localization
pix_xy1=ceil(RoiFitsetPT1(:,2:3));% 
pix_xy2=ceil(RoiFitsetPT2(:,2:3));%

%% Sorting Molecules into Villi, body, background at 00um
for kk=1:size(pix_xy1,1)
      linindex1(kk) = sub2ind(size(AA),pix_xy1(kk,1),pix_xy1(kk,2));
end

% sorting into each fraction
[CellN1]=ismember(linindex1,IndCell);      % within a cell area in ROI
[BodyN1]=ismember(linindex1,IndCellBody);  % within cellbody in a cell area
[VilliN1]=ismember(linindex1,IndVill);     % within microvill in a cell area
[BgdN1]=ismember(linindex1,IndBackground); % within backtroudn in ROI


Region_Sort1=[size(pix_xy1,1),sum(CellN1),((size(pix_xy1,1)-sum(CellN1))),sum(BodyN1),sum(VilliN1)];
 

%% Sorting Molecules into Villi, body, background at 04um 

for kk=1:size(pix_xy2,1)
       linindex2(kk) = sub2ind(size(AA),pix_xy2(kk,1),pix_xy2(kk,2));
end

[CellN2]=ismember(linindex2,IndCell);
[BodyN2]=ismember(linindex2,IndCellBody);
[VilliN2]=ismember(linindex2,IndVill);
[BgdN2]=ismember(linindex2,IndBackground);

Region_Sort2=[size(pix_xy2,1),sum(CellN2),size(pix_xy2,1)-sum(CellN2),sum(BodyN2),sum(VilliN2)];

%% percentage in a file
percent_MV_0nm=Region_Sort1(5)/Region_Sort1(2)*100;
percent_CB_0nm=Region_Sort1(4)/Region_Sort1(2)*100;
percent_MV_400nm=Region_Sort2(5)/Region_Sort2(2)*100;
percent_CB_400nm=Region_Sort2(4)/Region_Sort2(2)*100;
fid = fopen('MV-CB-Sort.txt','a');
formatSpec = '\n %s %4.2f \n';
fprintf(fid,formatSpec,'Counts in MV at 0 nm=',Region_Sort1(5),';');
fprintf(fid,formatSpec,'Counts in CB at 0 nm=',Region_Sort1(4),';');
fprintf(fid,formatSpec,'Percentage of molecules in Microvilli at 0 nm=',percent_MV_0nm,';');
fprintf(fid,formatSpec,'Percentage of molecules in CellBody at 0 nm=',percent_CB_0nm,';');
fprintf(fid,formatSpec,'Counts in MV at 400 nm=',Region_Sort2(5),';');
fprintf(fid,formatSpec,'Counts in CB at 400 nm=',Region_Sort2(4),';');
fprintf(fid,formatSpec,'Percentage of molecules in Microvilli at 400 nm=',percent_MV_400nm,';');
fprintf(fid,formatSpec,'Percentage of molecules in CellBody at 400 nm=',percent_CB_400nm),';';
fclose(fid);

%% Draw Figures

    h=pcolor(SortMap_Binary);
    xlim([range_x(1),range_x(2)])
    ylim([range_y(1),range_y(2)])
    set(h,'LineStyle','none')
    axis square
    saveas(gcf,'MV-CB-SortMap','fig');
    saveas(gcf,'MV-CB-SortMap','jpeg');
    close

    h=pcolor(SortMap_Binary);
    xlim([range_x(1),range_x(2)])
    ylim([range_y(1),range_y(2)])
    set(h,'LineStyle','none')
    axis square
    hold on 
    scatter(RoiFitsetPT1(:,3),RoiFitsetPT1(:,2),'.','w');
    title('0nm ','Fontsize',14)
    hold off
    saveas(gcf,'MV-CB-SortMap_0nm_merge','fig');
    saveas(gcf,'MV-CB-SortMap_0nm_merge','jpeg');
    close

    h=pcolor(SortMap_Binary);
    xlim([range_x(1),range_x(2)])
    ylim([range_y(1),range_y(2)])
    set(h,'LineStyle','none')
    axis square
    hold on  
    scatter(RoiFitsetPT2(:,3),RoiFitsetPT2(:,2),'.','w');
    title('-400nm ','Fontsize',14)
    hold off
    saveas(gcf,'MV-CB-SortMap_400nm_merge','fig');
    saveas(gcf,'MV-CB-SortMap_400nm_merge','jpeg');
    close


    

save(['SortMap_Binary_' samplename],'SortMap_Binary'); 

Result_DensityInOut.ROI=[range_x range_y]; 
Result_DensityInOut.CellAreaInd=IndCell;
Result_DensityInOut.VilliAreaInd=IndVill;
Result_DensityInOut.CellBodyAreaInd=IndCellBody;
Result_DensityInOut.IndBackground=IndBackground;
Result_DensityInOut.RoundedXY1= pix_xy1;
Result_DensityInOut.RoundedXY2= pix_xy2;
Result_DensityInOut.Region_Sort1=Region_Sort1;
Result_DensityInOut.Region_Sort2=Region_Sort2;
Result_DensityInOut.Map=BB;
Result_DensityInOut.Map_Binary=SortMap_Binary;

save(['DensityInOut_' samplename],'Result_DensityInOut')

end 
% Index_villi

% Index_body

%total sum of all index should be matched to the total N !!!!!!!!!!!!!

% 
% 
% pix_xy=round(RoiFitsetPT1(:,2:3));
% scatter(pix_xy(:,1),pix_xy(:,2));
% 
% =meanLocDeltaZ()
```
