## Supplementary material for "Microvillar cartography: a super-resolution single-molecule imaging method to map the positions of membrane proteins with respect to cellular surface topography": Analysis MATLAB codes: Distribution_Analysis.m

```
%%find density distribution by concentric shapes for whole shape; Created by Shirsendu on 30nd August,2017 
clear all; close all; 
date='200121';%% give expt date
%path_avg_folder= 'D:\Ronen_project\DATA\LFA1';
d=10;%%distance between ages of two shapes in nm
den_dist=41;%%density distribution will be shown upto=(den_dist-1)*10(i.e. (41-1)*10=400nm)
z_region=20; %% Region of z hight you want to chose for center area in nm
microvillino=0;
Count_area_total_0nm=zeros(den_dist,7);
Count_area_total_400nm=zeros(den_dist,7);
for sg=1:den_dist
Count_area_total_0nm(sg,1)=(sg-1)*10;
Count_area_total_400nm(sg,1)=(sg-1)*10;
end
%% Input of VA_analysis file
[matfilename, pathname] = uigetfile( ...
{  '*.mat','MAT-files (*.mat)'; ...
   '*.*',  'All Files (*.*)'}, ...
   'Pick a "VA_New_analysis_allinone file', ...
   'MultiSelect', 'off'); 

cd(pathname);
load(matfilename);
%% creation of new folder for saving file
SaveFolderName=('Distribution Of Molecules');
Tip_Analysis_Folder=SaveFolderName;


if (exist (Tip_Analysis_Folder)==0)
    mkdir(pwd,Tip_Analysis_Folder)
else(exist(Tip_Analysis_Folder)==1)
    SaveNewName=input('NewName(1)/Deletename(0)??:')
    if SaveNewName==1;
    Tip_Analysis_Folder=input('SaveFolderName?? : ','s'); 
    else
        rmdir ('Distribution Of Molecules','s');
    end
    mkdir(pwd,Tip_Analysis_Folder)
end

cd (Tip_Analysis_Folder); 
%% Finding microvilli locations
rotation=(400-z_region)/z_region;
zdist=zeros(256,256);
zdist=MeanSTDdelta{1, 1};
check=0;
row_boudary=[range_y(1) range_y(1) range_y(2) range_y(2) range_y(1)];
col_boundary=[range_x(1) range_x(2) range_x(2) range_x(1) range_x(1)];
for ii=0:rotation
    ff=(400.001-(z_region*ii));
    for aa=1:256
    for bb=1:256
        if(zdist(aa,bb)<ff)
            zdist(aa,bb)=1;
        else
            zdist(aa,bb)=0;
        end
    end
    end
[L1,num1]=bwlabel(zdist,4);
for cc=1:num1
    [row1,col1]=find(L1==cc);
    [in_boundary,on_boundary]=inpolygon(row1,col1,row_boudary,col_boundary);
    s_in_boundary=sum(in_boundary);
    s_row1=size(row1,1);
    if (s_row1==s_in_boundary)
    zarea=zeros(256,256);
    zarea(:,:)=600;
    for dd=1:size(row1)
        zarea(row1(dd),col1(dd))=MeanSTDdelta{1, 1}(row1(dd),col1(dd));
    end
    ee=min(zarea(:));
    ff1=ff-z_region;
    [rowzz,colzz]=find(zarea<ff1);
    if ee<(ff-z_region)
        check=10;
        z_check=z_region+0.001;
        if(ff==z_check)
            check=0;
        end
        if(size(rowzz)<3)
            check=0;
        end
    end
    k = boundary(row1,col1);
    if (check<10)
        t_boundary=isempty(k);
        if (t_boundary==0)
        microvillino=microvillino+1; 
        boundarysize=size(k,1);
        A=zeros(boundarysize,2);
        A(:,1)=row1(k);
        A(:,2)=col1(k);
        area_microvilli=[(polyarea(A(:,1),A(:,2)))*(66.67^2)];
         %if (microvillino~=4)%&&(microvillino~=13)&&(microvillino~=14)
         if (area_microvilli<100000)
        savename=['Microvilli_' num2str(microvillino)];
        save(savename,'A')
         end
         %end
        %density_area=func_distance_density_central_area(A,d,RoiFitsetPT1,RoiFitsetPT2,microvillino,range_x,range_y);
        %density_area_total(:,2)=density_area_total(:,2)+density_area(:,2);
        %density_area_total(:,3)=density_area_total(:,3)+density_area(:,3);
        %A=zeros(boundarysize,2);
        plot(col1(k),row1(k));
        hold on
        end
    end
    xlim([range_x(1) range_x(2)])
    ylim([range_y(1) range_y(2)])
    check=0;
    end
end
zdist=meanLocDeltaZ;
end
hold on
contour(meanLocDeltaZ)
saveas(gcf,'Area_loc_Map')
saveas(gcf,'Area_loc_Map','jpg')
hold off

%% Finding Distance vs density in each microvilli and summing up
totalpt_0nm=size(RoiFitsetPT1,1);
totalpt_400nm=size(RoiFitsetPT2,1);
cellarea_total=((range_x(2)-range_x(1))*(range_y(2)-range_y(1))*(66.67*66.67));
filetype='*.mat';
path=[pwd '\'];
Listing = dir([path,filetype]);
if size(Listing,1)>0
        for nn= 1:size(Listing,1)
            filename=(Listing(nn).name);
             A=importdata([path filename]);
             [count_area_0nm,count_area_400nm]=func_distance_density_central_area(A,d,range_x,range_y,RoiFitsetPT1,RoiFitsetPT2,filename,den_dist);
             z_0nm=sum(count_area_0nm,1);
             z_400nm=sum(count_area_400nm,1);
             if(z_0nm(2)>0)
             Count_area_total_0nm(:,2)=Count_area_total_0nm(:,2)+count_area_0nm(:,2);
             Count_area_total_0nm(:,3)=Count_area_total_0nm(:,3)+count_area_0nm(:,3);
             Count_area_total_0nm(:,5)=Count_area_total_0nm(:,5)+count_area_0nm(:,5);
             Count_area_total_0nm(:,6)=Count_area_total_0nm(:,6)+count_area_0nm(:,6);
             end
             if(z_400nm(2)>0)
             Count_area_total_400nm(:,2)=Count_area_total_400nm(:,2)+count_area_400nm(:,2);
             Count_area_total_400nm(:,3)=Count_area_total_400nm(:,3)+count_area_400nm(:,3);
             Count_area_total_400nm(:,5)=Count_area_total_400nm(:,5)+count_area_400nm(:,5);
             Count_area_total_400nm(:,6)=Count_area_total_400nm(:,6)+count_area_400nm(:,6);
             end
        end
end
Count_area_total_0nm(:,2)=Count_area_total_0nm(:,2)/(totalpt_0nm+totalpt_400nm);
Count_area_total_400nm(:,2)=Count_area_total_400nm(:,2)/(totalpt_0nm+totalpt_400nm);
Count_area_total_0nm(:,3)=Count_area_total_0nm(:,3)/cellarea_total;
Count_area_total_400nm(:,3)=Count_area_total_400nm(:,3)/cellarea_total;
Count_area_total_0nm(:,4)=Count_area_total_0nm(:,2)./Count_area_total_0nm(:,3);
Count_area_total_0nm(:,5)=(Count_area_total_0nm(:,5)/cellarea_total)*100;
Count_area_total_0nm(:,6)=(Count_area_total_0nm(:,6)/(totalpt_0nm+totalpt_400nm))*100;
Count_area_total_0nm(:,7)=(Count_area_total_0nm(:,6)./Count_area_total_0nm(:,5));
Count_area_total_400nm(:,4)=Count_area_total_400nm(:,2)./Count_area_total_400nm(:,3);
Count_area_total_400nm(:,5)=(Count_area_total_400nm(:,5)/cellarea_total)*100;
Count_area_total_400nm(:,6)=(Count_area_total_400nm(:,6)/(totalpt_0nm+totalpt_400nm))*100;
Count_area_total_400nm(:,7)=(Count_area_total_400nm(:,6)./Count_area_total_400nm(:,5));
savename1=['Count_area_total_0nm'];
save(savename1,'Count_area_total_0nm')
save(savename1,'Count_area_total_0nm','-ascii','-double','-tabs')
savename2=['Count_area_total_400nm'];
save(savename2,'Count_area_total_400nm')
save(savename2,'Count_area_total_400nm','-ascii','-double','-tabs')
delta_count_delta_area=zeros(den_dist,3);
delta_count_delta_area(:,1)=Count_area_total_0nm(:,1);
delta_count_delta_area(:,2)=Count_area_total_0nm(:,7);
delta_count_delta_area(:,3)=Count_area_total_400nm(:,7);
save('delta_count_delta_area','delta_count_delta_area');
save('delta_count_delta_area','delta_count_delta_area','-ascii','-double','-tabs')
plot(Count_area_total_0nm(:,1),Count_area_total_0nm(:,4),'r','linewidth',3)
hold on
plot(Count_area_total_400nm(:,1),Count_area_total_400nm(:,4),'g','linewidth',3)
hold on
xlabel(sprintf ('R (nm)\n Distance from Central microvilli region'))
%ylabel('Density (nm^{-2})')
ylabel('Fraction of count/Fraction of area')
saveas(gcf,['density_total'])
saveas(gcf,['density_total'],'jpg')
close
plot(Count_area_total_0nm(:,1),Count_area_total_0nm(:,7),'r','linewidth',3)
hold on
plot(Count_area_total_400nm(:,1),Count_area_total_400nm(:,7),'g','linewidth',3)
xlabel(sprintf ('R (nm)\n Distance from Central microvilli region'))
ylabel('\delta Count \div \delta Area')
saveas(gcf,['Ratio_count'])
saveas(gcf,['Ratio_count'],'jpg')
%save('density_total',density_area_total,'.mat')
close all
%% Finding microvilli vs cell body ditribution
locDisplayOption=1;
cellbodylimit=400;
samplename1=[samplename '_' num2str(cellbodylimit)];
[DensityInOut_MeanLoc,SortMap_Binary]=DensityInOut_v3_final(MeanSTDdelta,LocMicrovilicolor,RoiFitsetPT1,RoiFitsetPT2,range_x,range_y,samplename1,cellbodylimit);
save(['DensitySort_Summary_lim' num2str(cellbodylimit) '_' samplename]);
close
close all;
```
