## Supplementary material for "Microvillar cartography: a super-resolution single-molecule imaging method to map the positions of membrane proteins with respect to cellular surface topography": Analysis MATLAB codes: Drift_Correction_moviebymovie.m

```
%% Post analysis Drift correction
clc;clear all;close all
% parameter setting
samplename='JE2_FM_TCR';
x_range=[85 145];%
y_range=[95 155];%
getfocalplane='00um';
%finecorrect=0; % 0=movie by movie correction ;  1=frame by frame correction
Linfitall=0;  % 0: only missing value will be found from linear fitting 
              % 1: all value from linear fitting result from a set data
fittypename='poly2';
lastframe=1;
showmarker=1;
markersize=1000;
Channel=0; 


FileNameHead=[samplename,'_XcPostCor'];        
Do_merge =1; %% before and after merge
Refimagesave=1; 
Applyfilter=0;
filtertype= 'MeanFilter';% 'MedianFilter' 'LogFilter'; to the Average image; 
kernelsize= 60;
ApplyfiltertoCC=1;
kernelsizeCC=5;

%%% --------------------------------------------------------------------------
if ApplyfiltertoCC==0
kernelsizeCC=0;
end

    %% Open files (multiple mat or fits files
fprintf('select the TIRFM movies of cell membrane that are acquired between SLN movies captured at 00um plane \n')
[FNlist_TIRFM_00um, PathName_TIRFM_00um, Filterindex_TIRFM_00um] = uigetfile( ...
{  '*.fits','fits-files (*.fits)'
   '*.*',  'All Files (*.*)'}, ...
   'Pick a file', ...
   'MultiSelect', 'on');
fprintf('select all the mat files of CFSSTROM..._selCMPT folder for 00um plane \n')
[FitPTfile_selCMPT00um, PathName2_selCMPT00um] = uigetfile( ...
{  '*.mat','MAT-files (*.mat)'; ...
   '*.*',  'All Files (*.*)'}, ...
   'Pick a file', ...
   'MultiSelect', 'on'); 
fprintf('select the TIRFM movies of cell membrane that are acquired between SLN movies captured at 04um plane \n')
[FNlist_TIRFM_04um, PathName_TIRFM_04um, Filterindex_TIRFM_04um] = uigetfile( ...
{  '*.fits','fits-files (*.fits)'
   '*.*',  'All Files (*.*)'}, ...
   'Pick a file', ...
   'MultiSelect', 'on');
fprintf('select all the mat files of CFSSTROM..._selCMPT folder for 04um plane \n')
[FitPTfile_selCMPT04um, PathName2_selCMPT04um] = uigetfile( ...
{  '*.mat','MAT-files (*.mat)'; ...
   '*.*',  'All Files (*.*)'}, ...
   'Pick a file', ...
   'MultiSelect', 'on');
%% create combinexcCorrct_F5 folder

Motherfolder=PathName_TIRFM_00um;
NewxccorrectfolderName=['combinexcCorrct_F5'];                
                    mkdir(Motherfolder,NewxccorrectfolderName)
                    cd(Motherfolder);
                    cd(NewxccorrectfolderName)
                    path_xccorrectfolder=pwd;
               
%% find drift in 00 um movies                
    cd (PathName_TIRFM_00um);

    NumofFiles_TIRFM_00um=size(FNlist_TIRFM_00um,2);
    %Fidsfitout2=zeros(NumofFiles,14);
    %frames_out=zeros(size(FiducialFrame));
    getNthRef=zeros(1,NumofFiles_TIRFM_00um);
    
for ff = 1:NumofFiles_TIRFM_00um;

        if   ~iscell(FNlist_TIRFM_00um);    
             error('select multiple files')
        end      
%         else 
%              FileName=FNlist;
        
 FileName=FNlist_TIRFM_00um{ff};

        if strcmp(FileName(end-2:end),'mat')
           meanFrames=importdata([PathName_TIRFM_00um FileName]);
             
             
           if Channel==0
           RefImage{ff}=meanFrames ;
           elseif Channel==1
           RefImage{ff}=meanFrames(257:end,:);
           elseif Channel==2
           RefImage{ff}=meanFrames(1:256,:);    
           end

            % FileDimention=[256,256];
             
            
        elseif strcmp(FileName(end-3:end),'fits')

             fitsfilename = [PathName_TIRFM_00um FileName];
             cd (PathName_TIRFM_00um);
             info=fitsinfo(fitsfilename);
             FileDimention=info.PrimaryData.Size;
            
             allFrames = zeros(FileDimention(1),FileDimention(2));
        
             for jj=1:FileDimention(3)
             [filesData, frames_out1] = fits_shifter_single_Y_v30(PathName_TIRFM_00um,FileName,jj);  % frames_out ; FrameNth value (:,:,N)
     % Meanfilter substract 
                 if Applyfilter==1
                    frames_out2=FilterFrameOut(frames_out1,filtertype,kernelsize);
                 end 

                 if Applyfilter ==0 || MeanFrames == 1 && MeanFrameFilter==0  % sum of frame_out
                    allFrames=allFrames+frames_out1;       

                 elseif MeanFrames == 1 && MeanFrameFilter==1 % sum of frame_out
                    allFrames=allFrames+frames_out2;
                 end
             end
             

           meanFrames = allFrames/jj;
           if Channel==0
           RefImage{ff}=meanFrames ;
           elseif Channel==1
           RefImage{ff}=meanFrames(257:end,:);
           elseif Channel==2
           RefImage{ff}=meanFrames(1:256,:);    
           end
           
           
        end

        
      %% Get " File Extension "
    getparts = strread(FNlist_TIRFM_00um{ff},'%s','delimiter','.');
    getparts2= strread(getparts{1},'%s','delimiter','_');
    getN    = getparts2(end);
    getN    = str2double(getN);
    getNthRef(ff)= getN; 

end
 

if lastframe==0
    Template=RefImage{1};
elseif lastframe==1
    Template=RefImage{end};
end


for kk=1:size(RefImage,2)
RefImage_Range{kk}=RefImage{kk}(x_range(1):x_range(2),y_range(1):y_range(2));
%SimImage_Range{kk}=SimImage{kk}(x_range(1):x_range(2),y_range(1):y_range(2));
imagesc(RefImage_Range{kk})
set(gca,'YDir','normal');
axis equal tight 
        %h = imshow(inputimage,[]);
%if ==1       
%saveas(gcf,['comp','_frame',num2str(kk)],'tif');
%end

%pause
close

end

if Refimagesave==1
[h]=makeimagefile(RefImage_Range,'gray','RefImage_Range');
%[h]=makeimagefile(ccQ_all,'gray','RefImage_Range');
end


[ShiftResult]=Result_NormXcorr2_shift_v5_Range(Template,RefImage_Range,PathName_TIRFM_00um,FileNameHead,lastframe,ApplyfiltertoCC,kernelsizeCC,getfocalplane,x_range,y_range);

% if Refimagesave==1
% [h]=makeimagefile(cc_Qws,'gray','cc_Qws');
% %[h]=makeimagefile(ccQ_all,'gray','RefImage_Range');
% end

%[DriftRef]= DriftTBout(ResultXcorr,framesize1,framesize2)  

%% ReferencePT movie by movie

ReferencePT(:,1)=getNthRef;
ReferencePT(:,2)=ShiftResult(:,8);
ReferencePT(:,3)=ShiftResult(:,7);

%% Correct drift in 00 um movies
%% Edit : need to call FitsetPT; to correct;
%pause(0.5)
getNthFitset=zeros(1,size(FitPTfile_selCMPT00um,2));
for ff=1:size(FitPTfile_selCMPT00um,2)
cd (PathName2_selCMPT00um)
FitsetbeforePT{ff}=importdata(FitPTfile_selCMPT00um{ff});

      %% Get " File Extension "
    getparts = strread(FitPTfile_selCMPT00um{ff},'%s','delimiter','.');
    getparts2= strread(getparts{1},'%s','delimiter','_');
    getN    = getparts2(end);
    getN    = str2double(getN);
    getNthFitset(ff)= getN; 
end

if size(FNlist_TIRFM_00um,2)> size(FitPTfile_selCMPT00um,2)
  error('Reference point should be the same or smaller than the Fitset')   
end

%% Fill-up missin frame values 

[missingframes,misind] = setdiff(getNthFitset,getNthRef); %% A>B 


if isempty(missingframes)==1 && size(FNlist_TIRFM_00um,2)== size(FitPTfile_selCMPT00um,2);
        disp('Fiducial found from all frames')
        

elseif isempty(missingframes)==0 %% some of the fiducials were missing in a set
        fprintf(1,'Fiducial point were missing %g frames \n',length(missingframes));
        
        NewRefPT=zeros(length(getNthFitset),3);
        cnt=0;
            tf=ismember(getNthFitset,ReferencePT(:,1));
            for mm=1:size(getNthFitset,2)
                    if tf(mm)>0;
                        cnt=cnt+1;
                       NewRefPT(mm,:)=ReferencePT(cnt,:);
%                     elseif(mm)==0;
%                        NewRefPT(mm,:)=[0,0,0];
                     end 
            end
   
       
        [estPT,Refnewx,Refnewy,goodnessxy]=estPos_xcorr(NewRefPT,fittypename);

        fprintf(1,'adjrsqure_x: %g adjrsqure_y:%g \n',goodnessxy(1),goodnessxy(2));  
        
end        

%% Calculate drift xy based on the reference position
if Linfitall==0 && isempty(missingframes)==0 
   
            for mm=1:size(getNthFitset,2)
                    if tf(mm)==0;
                        cnt=cnt+1;
                       NewRefPT(mm,:)=estPT(mm,:)
%                     elseif(mm)==0;
%                        NewRefPT(mm,:)=[0,0,0];
                     end 
            end
    
    ReferencePT=NewRefPT;
    
elseif Linfitall==1;
     
    [estPT,Refnewx,Refnewy,goodnessxy]=estPos_xcorr(ReferencePT,fittypename);
     ReferencePT=estPT;
    
       fprintf(1,'adjrsqure_x: %g adjrsqure_y:%g \n',goodnessxy(1),goodnessxy(2));

%else 
 %       error('LinFitall???')   
end

%% Corrected Fiducial: Reconstruction (checking the correction of fiducial marker)
SaveName=[FitPTfile_selCMPT00um{1}(1:end-5),'_',num2str(kernelsizeCC)];
if lastframe==1
    SaveName=[SaveName,'_L'];
end

[CorrectedFitsetPT]=FitPTDriftcorrect_moviebymovie_Range(FitsetbeforePT,ReferencePT,x_range,y_range,SaveName,Do_merge,showmarker,markersize);

save(['xcCorrct_F',num2str(kernelsizeCC),'_',SaveName],'CorrectedFitsetPT');
current_path=pwd;
cd(path_xccorrectfolder)
save(['xcCorrct_F',num2str(kernelsizeCC),'_',SaveName],'CorrectedFitsetPT');
cd(current_path)
%save(['Xcorr2_Result_Fit_nm_',SaveName],'Result_Fit_nm','-ascii');


%% Find drift in 04 um movies
getfocalplane='04um';
cd (PathName_TIRFM_04um);

    NumofFiles_TIRFM_04um=size(FNlist_TIRFM_04um,2);
    %Fidsfitout2=zeros(NumofFiles,14);
    %frames_out=zeros(size(FiducialFrame));
    getNthRef=zeros(1,NumofFiles_TIRFM_04um);
    
for ff = 1:NumofFiles_TIRFM_04um

        if   ~iscell(FNlist_TIRFM_04um)    
             error('select multiple files')
        end      
%         else 
%              FileName=FNlist;
        
 FileName=FNlist_TIRFM_04um{ff};

        if strcmp(FileName(end-2:end),'mat')
           meanFrames=importdata([PathName_TIRFM_04um FileName]);
             
             
           if Channel==0
           RefImage{ff}=meanFrames ;
           elseif Channel==1
           RefImage{ff}=meanFrames(257:end,:);
           elseif Channel==2
           RefImage{ff}=meanFrames(1:256,:);    
           end

            % FileDimention=[256,256];
             
            
        elseif strcmp(FileName(end-3:end),'fits')

             fitsfilename = [PathName_TIRFM_04um FileName];
             cd (PathName_TIRFM_04um);
             info=fitsinfo(fitsfilename);
             FileDimention=info.PrimaryData.Size;
            
             allFrames = zeros(FileDimention(1),FileDimention(2));
        
             for jj=1:FileDimention(3)
             [filesData, frames_out1] = fits_shifter_single_Y_v30(PathName_TIRFM_04um,FileName,jj);  % frames_out ; FrameNth value (:,:,N)
     % Meanfilter substract 
                 if Applyfilter==1
                    frames_out2=FilterFrameOut(frames_out1,filtertype,kernelsize);
                 end 

                 if Applyfilter ==0 || MeanFrames == 1 && MeanFrameFilter==0  % sum of frame_out
                    allFrames=allFrames+frames_out1;       

                 elseif MeanFrames == 1 && MeanFrameFilter==1 % sum of frame_out
                    allFrames=allFrames+frames_out2;
                 end
             end
             

           meanFrames = allFrames/jj;
           if Channel==0
           RefImage{ff}=meanFrames ;
           elseif Channel==1
           RefImage{ff}=meanFrames(257:end,:);
           elseif Channel==2
           RefImage{ff}=meanFrames(1:256,:);    
           end
           
           
        end

        
      %% Get " File Extension "
    getparts = strread(FNlist_TIRFM_04um{ff},'%s','delimiter','.');
    getparts2= strread(getparts{1},'%s','delimiter','_');
    getN    = getparts2(end);
    getN    = str2double(getN);
    getNthRef(ff)= getN; 

end
 

if lastframe==0
    Template=RefImage{1};
elseif lastframe==1
    Template=RefImage{end};
end


for kk=1:size(RefImage,2)
RefImage_Range{kk}=RefImage{kk}(x_range(1):x_range(2),y_range(1):y_range(2));
%SimImage_Range{kk}=SimImage{kk}(x_range(1):x_range(2),y_range(1):y_range(2));
imagesc(RefImage_Range{kk})
set(gca,'YDir','normal');
axis equal tight 
        %h = imshow(inputimage,[]);
%if ==1       
%saveas(gcf,['comp','_frame',num2str(kk)],'tif');
%end

%pause
close

end

if Refimagesave==1
[h]=makeimagefile(RefImage_Range,'gray','RefImage_Range');
%[h]=makeimagefile(ccQ_all,'gray','RefImage_Range');
end


[ShiftResult]=Result_NormXcorr2_shift_v5_Range(Template,RefImage_Range,PathName_TIRFM_00um,FileNameHead,lastframe,ApplyfiltertoCC,kernelsizeCC,getfocalplane,x_range,y_range);

% if Refimagesave==1
% [h]=makeimagefile(cc_Qws,'gray','cc_Qws');
% %[h]=makeimagefile(ccQ_all,'gray','RefImage_Range');
% end

%[DriftRef]= DriftTBout(ResultXcorr,framesize1,framesize2)  

%% ReferencePT movie by movie

ReferencePT_04um(:,1)=getNthRef;
ReferencePT_04um(:,2)=ShiftResult(:,8);
ReferencePT_04um(:,3)=ShiftResult(:,7);
%% correct drift in 04um movies
%% Edit : need to call FitsetPT; to correct;
%pause(0.5)
getNthFitset=zeros(1,size(FitPTfile_selCMPT04um,2));
for ff=1:size(FitPTfile_selCMPT04um,2)
cd (PathName2_selCMPT04um)
FitsetbeforePT{ff}=importdata(FitPTfile_selCMPT04um{ff});

      %% Get " File Extension "
    getparts = strread(FitPTfile_selCMPT04um{ff},'%s','delimiter','.');
    getparts2= strread(getparts{1},'%s','delimiter','_');
    getN    = getparts2(end);
    getN    = str2double(getN);
    getNthFitset(ff)= getN; 
end

if size(FNlist_TIRFM_04um,2)> size(FitPTfile_selCMPT04um,2)
  error('Reference point should be the same or smaller than the Fitset')   
end

%% Fill-up missin frame values 

[missingframes,misind] = setdiff(getNthFitset,getNthRef); %% A>B 


if isempty(missingframes)==1 && size(FNlist_TIRFM_04um,2)== size(FitPTfile_selCMPT04um,2);
        disp('Fiducial found from all frames')
        

elseif isempty(missingframes)==0 %% some of the fiducials were missing in a set
        fprintf(1,'Fiducial point were missing %g frames \n',length(missingframes));
        
        NewRefPT=zeros(length(getNthFitset),3);
        cnt=0;
            tf=ismember(getNthFitset,ReferencePT_04um(:,1));
            for mm=1:size(getNthFitset,2)
                    if tf(mm)>0;
                        cnt=cnt+1;
                       NewRefPT(mm,:)=ReferencePT_04um(cnt,:);
%                     elseif(mm)==0;
%                        NewRefPT(mm,:)=[0,0,0];
                     end 
            end
   
       
        [estPT,Refnewx,Refnewy,goodnessxy]=estPos_xcorr(NewRefPT,fittypename);

        fprintf(1,'adjrsqure_x: %g adjrsqure_y:%g \n',goodnessxy(1),goodnessxy(2));  
        
end        

%% Calculate drift xy based on the reference position
if Linfitall==0 && isempty(missingframes)==0 
   
            for mm=1:size(getNthFitset,2)
                    if tf(mm)==0;
                        cnt=cnt+1;
                       NewRefPT(mm,:)=estPT(mm,:)
%                     elseif(mm)==0;
%                        NewRefPT(mm,:)=[0,0,0];
                     end 
            end
    
    ReferencePT_04um=NewRefPT;
    
elseif Linfitall==1;
     
    [estPT,Refnewx,Refnewy,goodnessxy]=estPos_xcorr(ReferencePT_04um,fittypename);
     ReferencePT_04um=estPT;
    
       fprintf(1,'adjrsqure_x: %g adjrsqure_y:%g \n',goodnessxy(1),goodnessxy(2));

%else 
 %       error('LinFitall???')   
end

%% Corrected Fiducial: Reconstruction (checking the correction of fiducial marker)
SaveName=[FitPTfile_selCMPT04um{1}(1:end-5),'_',num2str(kernelsizeCC)];
if lastframe==1
    SaveName=[SaveName,'_L'];
end

[CorrectedFitsetPT]=FitPTDriftcorrect_moviebymovie_Range(FitsetbeforePT,ReferencePT_04um,x_range,y_range,SaveName,Do_merge,showmarker,markersize);

save(['xcCorrct_F',num2str(kernelsizeCC),'_',SaveName],'CorrectedFitsetPT');
current_path=pwd;
cd(path_xccorrectfolder)
save(['xcCorrct_F',num2str(kernelsizeCC),'_',SaveName],'CorrectedFitsetPT');
cd(current_path)
%save(['Xcorr2_Result_Fit_nm_',SaveName],'Result_Fit_nm','-ascii');


disp('Drift correction is done. ')


cd (Motherfolder)

%% Parameter Save
CorrectionSet.ReferencePT=ReferencePT;
CorrectionSet.ReferencePT_04um=ReferencePT_04um;
%CorrectionSet.lastframe=lastframe;
CorrectionSet.showmarker=showmarker;
CorrectionSet.markersize=markersize;
CorrectionSet.Channel=Channel;
CorrectionSet.x_range=x_range;
CorrectionSet.y_range=y_range;
%CorrectionSet.RefImage_Range=RefImage_Range;
CorrectionSet.FileNameHead=FileNameHead;
%CorrectionSet.SaveName=['xcCorrect',SaveName];
CorrectionSet.Do_merge=Do_merge;
%CorrectionSet.getNthRef=getNthRef;
%CorrectionSet.getNthFitset=getNthFitset;
CorrectionSet.Applyfilter=Applyfilter;
CorrectionSet.filtertype=filtertype;
CorrectionSet.kernelsize=kernelsize;
CorrectionSet.Linfitall=Linfitall;
CorrectionSet.fittypename=fittypename;
CorrectionSet.analysistype='moviebymovie';
CorrectionSet.Refimagesave=Refimagesave;
CorrectionSet.ApplyfiltertoCC=ApplyfiltertoCC;
CorrectionSet.kernelsizeCC=kernelsizeCC;
%CorrectionSet.getfocalplane=getfocalplane;


save(['CorrectionSet_all'],'CorrectionSet');


% 
% 
% 
% 
% % CorrFidmoviePT=[DriftPT(:,1) DriftPT(:,4) DriftPT(:,5)];
% 
% % CorrFig=figure('Name','FiducialCorrection');
% %              FM=FidfitPT(:,2);
% %              FN=FidfitPT(:,3);
% %              plot(FN,FM,'.','MarkerSize',0.2);  %% Don't know why 
% %              axis equal
% %              xlim([0 size(FiducialFrame,2)]);
% %              ylim([0 size(FiducialFrame,1)]);              
% %              hold on   
% %              CorrM=CorrFidmoviePT(:,2);
% %              CorrN=CorrFidmoviePT(:,3);
% %              %axis equal
% %              plot(CorrN,CorrM,'.','MarkerSize',0.2,'Color','red');   %% Don't know why 
% % %              xlim([1 FileDimention(1)]);
% % %              ylim([1 FileDimention(2)]);  
% 
% % %% Fiducial Marker Drawing
% % [CelFidPT,CelCorrPT,MerFig]=Fidmolbymolanalysis_v4(FiducialAll,FidfitPT,CorrFidmoviePT,3,Nth);%Nth=0; molbymol name;
% % %molbymolanalysis_v4(Avefitdata,FidfitPT,CorrFidPT,sortws,Nth): default Nth=0;  molbymol name;
% % 
% % %CelFitPT is cell type of FidfitPT; 
% % SaveName=FileName(1:end-5);
% %                   
% % figuresavenamefit= ['Fid_',num2str(Nth),SaveName];
% %              saveas(CorrFig,figuresavenamefit);%,'jpg');
% % 
% %              
% %   
% % %ReferencePT=[DriftPT(:,1) DriftPT(:,2) DriftPT(:,3)];
% ReferenceMoviePT=[getNth' DriftPT(:,2) DriftPT(:,3)];
% 
% 
% 
% %% Edit : need to call FitsetPT; to correct;
% [FitPTfile, Pathname] = uigetfile( ...
% {  '*.mat','MAT-files (*.mat)'; ...
%    '*.*',  'All Files (*.*)'}, ...
%    'Pick a file', ...
%    'MultiSelect', 'on');
% 
% cd (Pathname)
% nFiles=size(FitPTfile,2);
% CorrectedFitsetPT=[];
% FitsetPT=[];
%     %% Get FitoutPT File Extension 
%         for ff=1:nFiles
% 
%         getparts = strread(FitPTfile{ff},'%s','delimiter','.');
%         getparts3= strread(getparts{1},'%s','delimiter','_');
%         getN2    = getparts3(end);
%         getN2    = str2double(getN2);
%         getNth2(ff)= getN2; 
% 
%         if ismember(getN2,getNth)
%             if   iscell(FitPTfile);    
%                  FileName3=FitPTfile{ff};
%             else 
%                  FileName3=FitPTfile;
%             end    
%             
% 
%            SaveName=FileName3(1:end-4); % matfiles
%          FitsetbeforePT=importdata([Pathname FileName3]);
%          
%          
%          
%          Reference=ReferenceMoviePT(ff,:);
% 
%         % CorrectedFitsetPTn=FitPTDriftcorrect(FitsetbeforePT,Reference,FileDimention,SaveName,0);% 0= do_mor
%          [CorrectedFitsetPTn]=FitPTDriftcorrect_movie(FitsetbeforePT,Reference,FileDimention,SaveName,0,Fx,Fy);   
%          CorrectedFitsetPT= vertcat(CorrectedFitsetPT,CorrectedFitsetPTn);
%          FitsetPT=vertcat(FitsetbeforePT,FitsetPT); 
%          
% 
% 
%         end
% 
%         end
clc
```
