## Supplementary material for "Microvillar cartography: a super-resolution single-molecule imaging method to map the positions of membrane proteins with respect to cellular surface topography": Analysis MATLAB codes: dSTORM_dualplane_Roi.m

```
function [ret]=dSTORM_dualplane_Roi(stormcolor1,stormcolor2,StormData,StormData2,StormfileName1,StormfileName2,local_scale,range_x,range_y,markersize,zMax)
%VARec = meanLocDeltaZ;
%% Merge Two figures 
% colormap('gray');
% 
% focalplanename='merge';
% 

% colormap(myColorMap)
% resultImage = imagesc(VARec);
% set(gca,'YDir','normal')
% colorbar
% caxis([0 zMax]) 
% axis equal; 
% % set(resultImage,'EdgeColor','none')
% view ([0 90])
%axis resultImage;
% figure
%set(gcf, 'Position', [10 20 2000 900]);
% hold all;
ret=figure;
%set(gcf, 'Position', [10 20 2000 900]);
%subplot(1, 2, 1);
%scatter(StormData(:,2), StormData(:,1), 3, 'o','filled','MarkerEdgeColor',stormcolor1);
%scatter(StormData(:,2), StormData(:,1), 3,'o','filled','MarkerEdgeColor',stormcolor1,'MarkerFaceColor',stormcolor1);
scatter(StormData(:,2), StormData(:,1), 3,'*','MarkerEdgeColor',stormcolor1);
title(['focal plane: 0nm'],'Fontsize',18);
%set(gca,'LooseInset',get(gca,'TightInset'))
axis equal tight

SaveName=['dSTORM_',StormfileName1(1:end-4)];
if length(SaveName)>60 
    SaveName = [SaveName(1:50),'_0nm'];% type char
end

if local_scale==1
addscale3_um_range_label(range_x,range_y,markersize,0)
    saveas(gcf,['scale' SaveName]);%,'jpg'); 
    set(gcf, 'Position', get(0, 'ScreenSize')); 
    saveas(gcf,['scale' SaveName],'jpg'); 
end
close;

ret=figure;
%set(gcf, 'Position', [10 20 2000 900]);
%subplot(1, 2, 1);
%scatter(StormData2(:,2), StormData2(:,1), 3, 'o','filled','MarkerEdgeColor',stormcolor2);
%scatter(StormData2(:,2), StormData2(:,1), 2,'o','filled','MarkerEdgeColor',stormcolor2,'MarkerFaceColor',stormcolor2);

scatter(StormData2(:,2), StormData2(:,1), 3,'*','MarkerEdgeColor',stormcolor2);
title(['focal plane: -400nm'],'Fontsize',18);
%set(gca,'LooseInset',get(gca,'TightInset'))
axis equal tight

SaveName=['dSTORM_',StormfileName2(1:end-4)];
if length(SaveName)>60 
    SaveName = [SaveName(1:50),'_-400nm'];% type char
end

if local_scale==1
addscale3_um_range_label(range_x,range_y,markersize,0)
    saveas(gcf,['scale' SaveName]);%,'jpg'); 
    set(gcf, 'Position', get(0, 'ScreenSize')); 
    saveas(gcf,['scale' SaveName],'jpg'); 
end
close;

ret=figure;
%scatter(StormData(:,2), StormData(:,1), 2,'o','filled','MarkerEdgeColor',stormcolor1,'MarkerFaceColor',stormcolor1);

scatter(StormData(:,2), StormData(:,1), 3,'*','MarkerEdgeColor',stormcolor1);
%scatter(StormData(:,2), StormData(:,1), 3, 'o','filled','MarkerEdgeColor',stormcolor1);
hold on
%scatter(StormData2(:,2), StormData2(:,1), 2,'o','filled','MarkerEdgeColor',stormcolor2,'MarkerFaceColor',stormcolor2);
scatter(StormData2(:,2), StormData2(:,1), 3,'*','MarkerEdgeColor',stormcolor2);
%scatter(StormData2(:,2), StormData2(:,1), 3, 'o','filled','MarkerEdgeColor',stormcolor2);
title(['dual focal plane'],'Fontsize',18);
%set(gca,'LooseInset',get(gca,'TightInset'))
axis equal tight

SaveName=['dual_dSTORM_',StormfileName1(1:end-4)];
if length(SaveName)>60 
    SaveName = [SaveName(1:50)];% type char
end

if local_scale==1
addscale3_um_range_label(range_x,range_y,markersize,0)
    saveas(gcf,['scale' SaveName]);%,'jpg'); 
    set(gcf, 'Position', get(0, 'ScreenSize')); 
    saveas(gcf,['scale' SaveName],'jpg'); 
end

hold off
% 
% 
% 
% 
% 
% scatter(StormData2(:,2), StormData2(:,1), 2, 'o','filled','MarkerEdgeColor',stormcolor2);
% title(['focalplane: -400nm'],'Fontsize',18);
% set(gca,'LooseInset',get(gca,'TightInset'))
% axis equal square
% if local_scale==1
% addscale3_um_range_label(range_x,range_y,markersize,0)
%     saveas(gcf,['scale' SaveName]);%,'jpg'); 
%     set(gcf, 'Position', get(0, 'ScreenSize')); 
%     saveas(gcf,['scale' SaveName],'jpg'); 
% end
% 
% SaveName=['dSTORM_',StormfileName(1:end-4)];
% % 
% if length(StormfileName)>60 
%     SaveName = [SaveName(1:60)];% type char
% end
% saveas(gcf,[SaveName]);%,'jpg');    
% %set(gcf, 'Position', get(0, 'ScreenSize'));
% saveas(gcf,[SaveName],'jpg');    
% % % 
% % if local_scale==1;
% %     
% addscale3_um_range(range_x,range_y,markersize)
%     saveas(gcf,['scale' SaveName]);%,'jpg');    
% end
% %close; 
% hold off
end
% focalplane =-400nm
% 
% figure 
% resultImage = imagesc(VARec);
% set(gca,'YDir','normal')
% myColorMap = colormap('hsv(16)'); %set the colormap used  %autumn
% myColorMap =(flipud(colormap('jet(8)')));
% colorbar
% caxis([0 600]) 
% axis equal; 
% set(resultImage,'EdgeColor','none')
% view ([0 90])
% axis resultImage;
% hold all;
% if dual focalplane==1
% scatter(postData2shift(:,2), postData2shift(:,1), 2, '+','MarkerEdgeColor',[0 0 0]);
% set(gca,'LooseInset',get(gca,'TightInset'))
% title(['focal plane: -400nm'],'Fontsize',12);
% 
% [focalplane]= GetPartName(sourceDataName2,'um');
% 
% SaveName=sourceDataName2;
% 
% if length(sourceDataName2)>60 
%     SaveName = [sourceDataName2(1:50),'_',focalplane];% type char
% end
% 
% saveas(gcf,[SaveName]);%,'jpg');    
% 
% if local_scale==1;
% addscale3_um_range(range_x,range_y,markersize);
%    saveas(gcf,['scale_' SaveName]);%,'jpg');    
% close; 
% 
% %% dual focal plane merge
% 
% figure;
% resultImage = imagesc(VARec);
% set(gca,'YDir','normal')
% 
% myColorMap = colormap('hsv(16)'); %set the colormap used  %autumn
% % myColorMap =(flipud(colormap('jet(8)')));
% colorbar
% caxis([0 600]) 
% axis equal; 
% % set(resultImage,'EdgeColor','none')
% % view ([0 90])
% %axis resultImage;
% 
% hold all;
% 
% %  colormap('gray');
% % scatter(postData(:,2), postData(:,1), 1, postData(:,3),'MarkerEdgeColor',[1 0 0]);
% %scatter(postData1shift(:,2), postData1shift(:,1), 2, '+','MarkerEdgeColor',[0 0.5 0.5]);
% scatter(postData1shift(:,2), postData1shift(:,1), 2, '+','MarkerEdgeColor',[0 0 0]);
% %hold all;
% scatter(postData2shift(:,2), postData2shift(:,1), 2, '+','MarkerEdgeColor',[0 0.5 0.5]);
% title(['dual focal plane'],'Fontsize',12);
% set(gca,'LooseInset',get(gca,'TightInset'))
% 
% 
% SaveName=sourceDataName;
% 
% if length(sourceDataName)>60 
%     SaveName = [sourceDataName(1:50),'_',focalplane];% type char
% end
% 
% saveas(gcf,['Dual_' SaveName]);%,'jpg');    
% 
% if local_scale==1;
%     
% addscale3_um_range(range_x,range_y,markersize);
% 
%    saveas(gcf,['Dual_' SaveName]);%,'jpg');    
% 
% end
```
