## Supplementary material for "Microvillar cartography: a super-resolution single-molecule imaging method to map the positions of membrane proteins with respect to cellular surface topography": Analysis MATLAB codes: estPos_xcorr.m

```
function[estPT,fitx,fity,goondessxy]=estPos_xcorr(FidfitPT,fittype) 

% LIBNAME           DESCRIPTION
% 'poly1'           Linear polynomial curve
% 'poly11'          Linear polynomial surface
% 'poly2'           Quadratic polynomial curve
% 'linearinterp'    Piecewise linear interpolation
% 'cubicinterp'     Piecewise cubic interpolation
% 'smoothingspline' Smoothing spline (curve)
% 'lowess'          Local linear regression (surface)

    x = 1:size(FidfitPT,1);
    x1=x';
    x2=x';
    y1 = FidfitPT(:,2); % coordinate x
    y2 = FidfitPT(:,3); % coordinate y
    
    
    [indy1]=find(y1==0);   
    y1(indy1)=[];
    x1(indy1)=[];
    
    [indy2]=find(y2==0);
    y2(indy2)=[];
    x2(indy2)=[];
    
    [fitx,gofx] = fit(x1,y1,fittype)%poly1
    figure
    scatter(x1,y1,'b','*') ;hold on;
    plot(x,fitx(x),'r','LineWidth',2)

    saveas(gcf,['Fitx_',fittype],'fig');hold off
    saveas(gcf,['Fitx_',fittype],'bmp');
%    saveas(gcf,['FidFitx' num2str(Nth)],'jpg');
                % close 
    
    
    
    [fity,gofy] = fit(x2,y2,fittype)%poly1
    figure
    scatter(x2,y2,'c','*') ;hold on;
    plot(x,fity(x),'r','LineWidth',2);
   
    saveas(gcf,['Fitx_',fittype],'fig'); hold off
    saveas(gcf,['Fitx_',fittype],'bmp');
%      y1fit = Lfitx.p1*x+Lfitx.p2;
%      y2fit = Lfity.p1*x+Lfity.p2;
     
     estPT=FidfitPT;
     estPT(:,2)=fitx(x);
     estPT(:,3)=fity(x);
    % Figure show %% for all
%     figure 
%      scatter(x1,y1,'b','*') ;hold on;
%      plot(x,y1fit,'r-.');hold off
%     figure
%      scatter(x2,y2,'c','*'); hold on; 
%      plot(x,y2fit,'r-.');hold off;
     
%      Ps=[fitx.p1, fitx.p2, fity.p1, fity.p2];% Linear fit parameter
     goondessxy=[gofx.adjrsquare,gofy.adjrsquare];
% output structure: 
% fitcurve = 
% 
%      Linear model Poly1:
%      fitcurve(x) = p1*x + p2
%      Coefficients (with 95% confidence bounds):
%        p1 =           5  (5, 5)
%        p2 =          12  (12, 12)
% 
% goodness = 
% 
%            sse: 1.0623e-024
%        rsquare: 1
%            dfe: 98
%     adjrsquare: 1
%           rmse: 1.0411e-013


%     scatter(x,y1,'b','*') 
%     P1 = polyfit(x,y1,1);
%     P2 = polyfit(x,y2,1);
%     y1fit = P1(1)*x+P1(2);
%     y2fit = P2(1)*x+P2(2);
%    figure;
%     scatter(x,y1,'b','*');hold on
%     plot(x,y1fit,'r-.');hold off
%    figure;
%     scatter(x,y2,'b','*');hold on
%     plot(x,y2fit,'g-.');hold off
   % fitobject = fit(x,y,fitType)
  
end
```
