## Supplementary material for "Microvillar cartography: a super-resolution single-molecule imaging method to map the positions of membrane proteins with respect to cellular surface topography": Analysis MATLAB codes: FilterFrameOut_size.m

```
function[C] = FilterFrameOut_size(A,filtertype,filtersize)
B=zeros(size(A,2),size(A,1));
C=zeros(size(A,2),size(A,1));

%A=frames_out;


switch filtertype

%% For mean, the region of mocrivill determied after i used LoG filter 
% edges = imfilter(AA,h,'replicate'); AA=images
% h = fspecial('log', 14, 0.5); filtertype: log, 14= kernel size (14),
% gaussian sigma
%h = fspecial('log', hsize, sigma)  returns a rotationally symmetric Laplacian of Gaussian filter of size hsize with standard deviation sigma (positive). hsize can be a vector specifying the number of rows and columns in h, or it can be a scalar, in which case h is a square matrix. The default value for hsize is [5 5] and 0.5 for sigma.
% median filter
    case 'MedianFilter'
B = medfilt2(A, [filtersize filtersize]);
C=A-B;
% figure
% imagesc(A)
% figure
% imagesc(B)
% C=A-B;
% figure
% imagesc(C)

    case 'MeanFilter'
h = fspecial('average', [filtersize filtersize]);
B = imfilter(A,h,'replicate'); 
% figure
% imagesc(B)
C=A-B;
% figure
% imagesc(C)

    case 'DiscFilter'
h = fspecial('disc', filtersize/2);
B = imfilter(A,h,'replicate'); 
C=A-B;
% figure
% imagesc(B)
% C=A-B;
% figure
% imagesc(C)


     case'LogFilter'
h = fspecial('log', 10, 0.5);
B = imfilter(A,h,'replicate'); 
C=B;
% figure
% imagesc(D)       
% figure
% imagesc(B)
% C=A-B;
% figure
% imagesc(C)
case'GaussFilter'
h = fspecial('gaussian', [filtersize filtersize], 0.5) 
B = imfilter(A,h,'replicate'); 


end

end


% B = medfilt2(A, [25 25])
% C=A-B;
% figure
% imagesc(A)
% figure
% imagesc(B)
% C=A-B;
% figure
% % imagesc(C)
% 
% h = fspecial('log', 14, 0.5);
% D = imfilter(A,h,'replicate'); 
% figure
% imagesc(D)
```
