## Supplementary material for "Microvillar cartography: a super-resolution single-molecule imaging method to map the positions of membrane proteins with respect to cellular surface topography": Analysis MATLAB codes: findmicrovilli_LoGFilter_angle15_Roi.m

```
 %%%  Find general microvilli regions (larger extend)
 %%% 15: save map
 %%% 14: change figure format; subplot
%%%  Threshold(for LogFilter threshold),N(numberoffiles),GetVA(micrometer position),SaveFileName,IgStack
%%%  LogfilterIgstack,meanLocMicrovilli]

function[LogfilterIgstack,LocMicrovili,LocTips]=findmicrovilli_LoGFilter_angle15_Roi(Threshold,N,GetVA,SaveFileName,IgStack,range_x,range_y)
h = fspecial('log',10, 0.5);
    for k=1:N
       LogfilterIgstack(:,:,k) = imfilter(IgStack(:,:,k),h,'replicate');

%     Medges=mean(edges,3);
%     figure
    %     surf(Medges)
       Microvilli(:,:,k)=1*(LogfilterIgstack(:,:,k)>Threshold);
       
    end
    save(['MV_Binary_' num2str(Threshold) SaveFileName num2str(GetVA(1)) num2str(GetVA(k))],'Microvilli');
%% create figure of Logfilter
       title1=['TIRF ',num2str(GetVA(1))];
       title2=['TIRF ',num2str(GetVA(1))];
       createfigure_subplot2type_Roi(IgStack(:,:,1),LogfilterIgstack(:,:,1),1,title1,title2,range_x,range_y);
       saveas(gcf,['LoGimage' SaveFileName num2str(GetVA(k))],'fig'); 
       saveas(gcf,['LoGimage' SaveFileName num2str(GetVA(k))],'jpg'); 
%       close;
%% create figure of Logfilter with Threshold       
       createfigure_subplots6_Roi(Microvilli,Threshold,GetVA,range_x,range_y);
%        pause
%        pcolor(LogfilterIgstack(:,:,k));title(['LoGimage Thr' num2str(Threshold)]); 
%        colormap(gray(2))
%        axis equal
       
       saveas(gcf,['LoGThr' num2str(Threshold) SaveFileName num2str(GetVA(k))],'fig');
       saveas(gcf,['LoGThr' num2str(Threshold) SaveFileName num2str(GetVA(k))],'jpg'); 

%     close;
%   
    %Medges=mean(LogfilterIgstack,3);
  %  MeanMicrovilli=mean(Microvilli,3); %logical
  % MeanLocMicrovilli=1*(MeanMicrovilli>0);
    %meanLocMicrovilli=1*(MeanMicrovilli==1); 
%   
BWminangle=Microvilli(:,:,1);
BWmaxangle=Microvilli(:,:,end);

% se90 = strel('line', 2, 90);%%se2 = strel('line',10,45)     % length 10, angle 45 degrees
% se0 = strel('line', 2, 0);
% BWsErode = imerode(BWs, [se90 se0]);
% figure, imshow(BWsErode), title('dilated gradient mask');
% 
% BWdfill = imfill(BWsErode, 'holes');
% figure, imshow(BWdfill);
% title('binary image with filled holes');

%BWnobord=BWminangle;
se90 = strel('line', 2, 90);%!!
se0 = strel('line', 2, 0);
% se90 = strel('diamond', 2, 90);
% se0 = strel('diamond', 2, 0);


% se90_1 = strel('line', 1, 90);
% se0_1 = strel('line', 1, 0);

%se = strel('ball',15,2);
%se = strel('disk',1); 
se = strel('diamond',2); 
%BWerode = imerode(BWnobord,seD);
%BWerode = imerode(BWminangle,[se90 se0]);!!
BWerode = imerode(BWminangle,se);


%BWmaxerode=imerode(BWmaxangle,[se90 se0]);
[LL,Lnum]=bwlabel(BWerode,4);
figure;
imagesc(LL);

xlim([range_x]);
ylim([range_y]);
set(gca,'YDir','normal')
axis equal tight ;
title('LocErode 2','FontSize',20);%!!
%title('Erode RollingBall','FontSize',20);%!!
%pause

LocTips=zeros(size(LL));

for kk=1:Lnum
                QL = 1*(LL==kk);  
               Qarea=find(QL>0);             
% se_n = 1;
% Factorial = 1;
% while nFactorial < 1e100
%     sen = sen + 1;
%     nFactorial = nFactorial * n;
   if length(Qarea)>= 10;
                   se_90 = strel('line', 2, 90);
                    se_0 = strel('line', 2, 0);
                    seD = strel('diamond',2);
                   BWmaxerode=imerode(BWmaxangle,[se_90 se_0]);
                   BWmaxerode2=imerode(BWmaxerode,[se_90 se_0]);
                   BWmaxerode3=imerode(BWmaxerode2,seD);
                   BWmaxerode4=imerode(BWmaxerode3,seD);
                   QL1=QL.*BWmaxangle;
                   Qarea1=find(QL1>0);
                   QL2=QL1*BWmaxerode;
                   Qarea2=find(QL2>0);
                   QL3=QL2*BWmaxerode2;
                   Qarea3=find(QL3>0);
                   QL4=QL3*BWmaxerode3;
                   Qarea4=find(QL4>0);
                   QL5=QL4*BWmaxerode4;
                   Qarea5=find(QL5>0);
      if ((size(Qarea1,1)>0)&&(size(Qarea2,1)>0)&&(size(Qarea3,1)>0)&&(size(Qarea4,1)>0)&&(size(Qarea5,1)>0))             
                   QL=QL.*BWmaxangle; 
                   %Qarea2=find(QL>0);
                   [~,num2]=bwlabel(QL,4);
                   %Qarea=find(QL>0);
                   
                   %if (size(Qarea,1)==0)|
                    %  QL = 1*(LL==kk); 
                   %end
           if num2==1 %&& (size(Qarea,1)>0)
                    %se_90 = strel('line', 2, 90);
                    %se_0 = strel('line', 2, 0);
                    %BWmaxerode=imerode(BWmaxangle,[se_90 se_0]);                   
                    QL = QL.*BWmaxerode ; %% if both values are 1, it is 1 otherwise 0 
                    [~,num3]=bwlabel(QL,4);
                  if num3==1 
                        %BWmaxerode2=imerode(BWmaxerode,[se_90 se_0]);
                        QL = QL.*BWmaxerode2;
                        [Lq,num4]=bwlabel(QL,4);
                       % Lqarea=1*Lq(Lq>0);
                    if num4==1
                        %seD = strel('diamond',2);
                        %BWmaxerode3=imerode(BWmaxerode2,seD);
                        QL = QL.*BWmaxerode3;
                        [~,num5]=bwlabel(QL,4);
                       if num5==1
                        %BWmaxerode4=imerode(BWmaxerode3,seD);
                        QL = QL.*BWmaxerode4;
                       end
                    end
               
                 end
           end   
     end
   end
%  pcolor(QL);hold on
  % dividing two tips 

        
        
   
               

  LocTips=LocTips+QL;
         clear Qarea QL
end

% figure;pcolor(LocTips);title('LocTips','FontSize',20);
%   colormap(gray(2))
%   axis equal tight
%   saveas(gcf,['LocTips' SaveFileName],'jpg'); 
% 
[TipL,finaln]=bwlabel(LocTips,4);
% LocMicrovilicolor=imdilate(TipL,[se0 se90]);
% LocMicrovili=imdilate(LocTips,[se0 se90]);
seD = strel('diamond',2);
LocMicrovilicolor=imdilate(TipL,[seD]);
LocMicrovili=imdilate(LocTips,[seD]);


save(['LocTipMap' SaveFileName],'TipL');
save(['LocTipMapDilate' SaveFileName],'LocMicrovilicolor');


figure;imagesc(LocMicrovili);title('LocMicrovilli','FontSize',20);

  colormap(gray(2))
  set(gca,'YDir','normal')
  axis equal tight
  
saveas(gcf,['LocMicrovilli_map' SaveFileName],'jpg');


title1=['LocTips n=',num2str(finaln)];
title2=['LocMicrovilli n=',num2str(finaln)];
createfigure_subplot2type_Roi(TipL,LocMicrovilicolor,2,title1,title2,range_x,range_y)
saveas(gcf,['LocMicrovilli' SaveFileName],'jpg');
saveas(gcf,['LocMicrovilli' SaveFileName]); 
%close all
% BWerode = imerode(BWerode,seD);
% se3 = strel('diamond',2);   nhood = getnhood(se3);   imagesc(nhood); axis xy; colormap('gray');

% % figure, imshow(BWfinal), title('segmented image');
% imshow (BWerode)

%LocTips=1*(LocTips>0);% 

% se90 = strel('line', 2, 90);
% se0 = strel('line', 2, 0);
% BWsdil = imdilate(BWerode, [se90 se0]);
% % figure, imshow(BWsdil), title('dilated gradient mask');


%AAA=1*(MeanMicrovilli~=1); 
%    AAA=1*(MeanMicrovilli>(1/(N))); %%%%%%%%%%%%%%%%%%%%
%    AAAA=meanLocMicrovilli-AAA;
    
%    AAAAA=1*(AAAA>0);
%    meanLocMicrovilli=AAAAA+meanLocMicrovilli;
%     BB=(IgStack(:,:,1).*AAAA);
%     LogBB = imfilter(imshow,h,'replicate');
%     
    % meanLocMicrovilli=1*(Medges>Threshold);
%     figure

end
 
```
