## Supplementary material for "Microvillar cartography: a super-resolution single-molecule imaging method to map the positions of membrane proteins with respect to cellular surface topography": Analysis MATLAB codes: findparticleFrameN.m

```
% correction for different sigma x and y 
function [CMoutSM,CMNumSM,FitoutSM,FitnumSM,selFitCMSigma]=findparticleFrameN(data,frameN,smTH,maxTH,maxPNTH,ws,sigma,ssd,sigmacm,cmssd,EMgain,PreAmp,mMMR,mSNR,minPNTH)
%            [CMout,CMnum]=findparticleFrameN_CM11(frames_out,FrameN,SmTH,BgdTH,MaxPNTH,ws,sigma,cmssd,EMgain,PreAmp,mSNR);(frames_out,FrameN,MinTH,MaxPNTH,ws,sigma,cmssd,EMgain,PreAmp,mSNR)
%changes: data1 (original frames_out; data2(imfilter applied frames_out);
%remove BgdTH; use fit data for original ; remove error 0.1
%remove -Lmin for CMfit
% correction made:  fitout(9,cnt)=  AveRErr;%%% correction made from mse to AveRErr ; which was done before Jan 2014. 
%% CMoutSM
% 1. frameN
% 2: xcm
% 3: ycm
% 4: sxcm
% 5: sycm
% 6: Lmax
% 7: Lmin
% 8: Lmax/Lmin
% 9: Qm
% 10 NumPho

%% FitoutSM
% 1. frameN
% 2: xfit
% 3: yfit
% 4: sxfit
% 5: syfit
% 6: Amp
% 7: Bgd
% 8: SNR
% 9: sme; AveRErr; (Average Ratio of error/fittevalue ; 95% confidence)
% 10 NumPho
% option 1(nonaccumulate) and 2(accumulatedate) for fitting the ws*ws 
% data1= before filter
% data2= after filter

%% selFitCMSigma
% 1. frameN
% 2: xfit
% 3: yfit
% 4: sxfit
% 5: syfit
% 6: Amp
% 7: Bgd
% 8: SNR
% 9: sme; AveRErr; (Average Ratio of error/fittevalue ; 95% confidence)
% 10 NumPho
% 11: sxcm
% 12: sycm
% option 1(nonaccumulate) and 2(accumulatedate) for fitting the ws*ws 
% data1= before filter
% data2= after filter

CMoutSM(1:10,1) =0;%centerofmass
CMNumSM=0;
FitoutSM(1:10,1) =0;%centerofmass
FitnumSM=0;
cnt=0;
fitcnt=0;
selFitCMSigma(1:12,1) =0;%centerofmass
%local maximum pixel coordinate and value
% wx0 = sigma;
% wy0 = sigma;
pn= (ws-1)/2;% pixel number from the center ex.ws=11, pn=4

 %% SEnsitivity of Andor iXon according to the A/D rate from the performance Sheet
if  PreAmp == 5
    Sensitivity = 11.65;
elseif PreAmp==1;
    Sensitivity = 64.2;
elseif PreAmp==2.4;
    Sensitivity = 25.54;
elseif PreAmp==5.2
    Sensitivity = 10.96; % Added for TIRF room setup
end
%Sensitivity

%% finds all the islands where the hypothesis is bigger than a certain threshold value
islands=1*(data>=smTH);

if  max(max(islands))>0
    [L,num] = bwlabel(islands,8); % creates a logical matrix at the same size as islands where each island is numbered. 8 is default
else 
    return  
end

if num>0        % num is the total number of islands.
    
% fprintf(1,'found %g islands from bwlabel\n',num); 

%% in each island finds the local maxima which will be used in the fitting as the starting point
    for n = 1:num    
         n                                   % finds all the particles inside the matrix and returns their indices in variable loc
         Q = data.*(L==n);                % sizeQ=size of filedimensionxfiledimention
         Lmax=max(max(Q))%;                % find max pixel in the island           
         [xmax,ymax] = find(Q==Lmax,1); 
           
       if xmax > pn && ymax > pn && xmax < size(Q,1)-pn &&  ymax < size(Q,2)-pn %%%% Center of Mass range

           
         Q(xmax-pn:xmax+pn,ymax-pn:ymax+pn)=1; % extanded a logical matrix as a size of ws (13x13)
         Q = data.*Q; 
          % [nn mm]=size(Q);% input original data on the recreated matrix

           Lmin=min(Q(find(Q>0))); 
                       
           
           if (Lmax < maxTH) % !!!!!!!!!!!!!!&& (Lmax/Lmin > mMMR)  %since the filter can make Lmin as negative value; modified on 140323              
              xcm=0; ycm=0; Mt=0;                    % xcm: x center of mass; ycm: y center of mass; Qm: total mass of the island %%%%% why?? sigmay is bigger than sigmax
% 
Q1=Q(xmax-pn:xmax+pn,ymax-pn:ymax+pn);
Q2=Q1;%-Lmin;;
[rc,cc] = ndgrid(1:size(Q2,1),1:size(Q2,2)); %!!!!!!!!!!
Mt = sum(Q2(:));
c1 = sum(Q2(:).* rc(:)) / Mt;
c2 = sum(Q2(:).* cc(:)) / Mt;

%xcm=c1
%ycm=c2

% b =beta,
 xcm = xmax + c1 - pn -1; % original x coordinate %%%%%%%%%%
 ycm = ymax + c2 - pn -1; % original y coordinate %%%%%%%%%%               


%               MtsubBac=Mt-(Lmin*(ws^2)); 
               NumPho = Sensitivity*Mt/EMgain/0.95%;
%% Lower Threshold
               
               if NumPho < maxPNTH && NumPho > minPNTH
%% calculate for the sigma based on the center of mass
                    % sx=0; sy=0;  % sx= sigmacmx, sy=sigmacmy                    
S1 = sum(Q2(:).* abs(c1-rc(:))) / Mt;
S2 = sum(Q2(:).* abs(c2-cc(:))) / Mt;                   

                   
            %    XYloc(round(xcm),round(ycm))=n;
                  cmout(1:10,n)=0;
                  cmout(1,n)=frameN; 
                  cmout(2,n)=xcm;
                  cmout(3,n)=ycm;
                  cmout(4,n)=S1;
                  cmout(5,n)=S2; 
                  cmout(6,n)=Lmax;
                  cmout(7,n)=Lmin; 
                  cmout(8,n)=Lmax/Lmin; 
                  cmout(9,n)=Mt;
                  cmout(10,n)=NumPho;
                      

             
                    if  cmout(4,n)>(sigmacm-cmssd) && cmout(5,n)>(sigmacm-cmssd) && cmout(4,n)<(sigmacm+cmssd) && cmout(5,n)<(sigmacm+cmssd) 
            %% Report CM result
                        cnt=cnt+1; % counter
                        CMoutSM(:,cnt)=cmout(:,n);
%                         fprintf(1,'found %g OK /islands from center of mass \n',cnt); 
                        CMNumSM=cnt;

                        %% 2D -Gaussian Fitting Function            

                        
                       % Q1=Q(xmax-pn:xmax+pn,ymax-pn:ymax+pn); %(for cut the ws*ws of single mole)
                        B =reshape(Q1,[ws^2,1]);
                        b0 = [(Lmax-Lmin) ws/2+1 ws/2+1 sigma sigma Lmin ];
                        [X, Y] = meshgrid(1:ws,1:ws);
                        xy_dat = [X(:) Y(:)];    

                        [b,r,J] = nlinfit(xy_dat, B, @mygauss2D_v19, b0); % b=[ampl, y0, x0, sigmax0, sigmay0, background]
                        
                         if b(2)>0 && b(2)< ws && b(3)>0 && b(3)< ws
                            


                                     b(2) = xmax + b(2) - pn -1; % original x coordinate
                                     b(3) = ymax + b(3) - pn -1; % original y coordinate                
                                                                     % r= residual
                    %%Fit confidence from estimated Error in Fitting                                                              % J=Jacobian
                                delta = Modnlparci_ver19(b,r,J);
                                AveRErr=mean(delta./b');                   % delta(for each b value) = se * tinv(1-alpha/2,v); alpha: default if 0.05 (95% confidence)
                                                                           % AveRErr=mean of ratio of Error/Fitresults in 95% confidence of b   
                                opts.Algorithm = 'Levenberg-Marquardt';
                                opts.Display = 'Off';
                                opts.Lower = [-Inf -Inf -Inf -Inf -Inf];
                                opts.Robust = 'Bisquare';


                                fitout(1,cnt)=cmout(1,n);
                                fitout(2,cnt)=b(2);%!!!!
                                fitout(3,cnt)=b(3);%!!!!
                                fitout(4,cnt)=b(4);
                                fitout(5,cnt)=b(5);  
                                fitout(6,cnt)=b(1); 
                                fitout(7,cnt)=b(6);
                                fitout(8,cnt)=(b(1)/b(6));
                                fitout(9,cnt)=AveRErr;%%% correction made from mse to AveRErr ; which was done before Jan 2014. 
                                fitout(10,cnt)=cmout(10,n);

                                
                                        if fitout(4,cnt)<(sigma + ssd +0.1)&& fitout(5,cnt)<(sigma + ssd)&&fitout(4,cnt)>(sigma - ssd +0.1)&& fitout(5,cnt)>(sigma - ssd)%&& fitout(9,cnt) < 0.1 
                                           %fitout(8,cnt)>= mSNR &&

                                            fitcnt=fitcnt+1;
                                            FitoutSM(:,fitcnt)=fitout(:,cnt);
                                             selFitCMSigma(:,fitcnt) =[fitout(1:5,cnt);cmout(6:8,n);fitout(8,cnt);fitout(10,cnt);cmout(4,n);cmout(5,n)]; 
                                        end

                        end
                    end
               end
           end
       end
    end
%   fprintf(1,'found %g/%g/%g molecules from 2Dfitting/cm/bw\n',fitcnt,cnt,num);      

     FitnumSM=fitcnt;

% figure
% h=pcolor(XYloc);
% set(h,'EdgeColor','none')

end 
end

    %            % fitting from the original matrix (SLOW)      
%             elseif Opt==0 || isempty(Opt)
%                 
%                 Q1=Q;
%                 Q1(Q==0)= Lmin; %(for cut the ws*ws of single mole)!!!!
%                 B =reshape(Q1,[nn*mm,1]);
%                 %size(B);
%                 b0 = [(Lmax-Lmin) cmout(2,n) cmout(3,n) sigma sigma Lmin];
%                 [X, Y] = meshgrid(1:nn,1:mm);
%                 xy_dat = [X(:) Y(:)];    
%             end
```
