## Supplementary material for "Microvillar cartography: a super-resolution single-molecule imaging method to map the positions of membrane proteins with respect to cellular surface topography": Analysis MATLAB codes: fit2dgaussian_4YM.m

```
function [b,ci] = fit2dgaussian_4YM(data,loc,ws)

% Inputs:
%   loc -   a list of central (x,y) pixels. 
%   ws -    the size of the region of interest
%   data -  the full matrix in which the 2d gaussian is embeded
%   Note -  this fits the data into a symmetric 2d gaussian, which might
%           not be the case in the cross correlation of the images.

% Outputs:
%   b -     5 parameters: [Intensity x y sigma bkd]
%   ci -    upper and lower limit of the estimate

n=1; % assuming you are only interested in the central peak
[X, Y] = meshgrid(1:ws, 1:ws); xy_data = [X(:) Y(:)];
A = double(data(loc(1,n)-(ws-1)/2:loc(1,n)+(ws-1)/2,loc(2,n)-(ws-1)/2:loc(2,n)+(ws-1)/2));
estimates = fit2dguasswbg(A);
[b,r1,J] = nlinfit(xy_data, reshape(A,[ws^2,1]), @mygauss3, estimates);
ci = nlparci(b,r1,J);
b(2)=loc(1,n)+b(2)-ceil(ws/2);  b(3)=loc(2,n)+b(3)-ceil(ws/2);

end
```
