## Supplementary material for "Microvillar cartography: a super-resolution single-molecule imaging method to map the positions of membrane proteins with respect to cellular surface topography": Analysis MATLAB codes: fit2dguasswbg.m

```
function [estimates,fval,model,FittedCurve] = fit2dguasswbg(data, firstguess)
% Call fminsearch with a random starting point.

switch nargin
    case 1
        start_point = [max(data(:)),ceil(size(data,1)/2),ceil(size(data,1)/2),0.703,mean(data(:))];
    case 2
        start_point=firstguess;
end

start_point=double(start_point);
[X,Y]=meshgrid(1:size(data,1),1:size(data,1));
xy_data=[X(:),Y(:)];

model = @guass2dfun;
[estimates,fval] = fminsearch(model, start_point);
[v,FittedCurve] = guass2dfun(estimates);
FittedCurve = reshape(FittedCurve,[size(data,1) size(data,2)]);

% guass2dfun accepts 2d gaussian parameters as inputs, and outputs sse,
% the sum of squares error for the measured and the FittedCurve. 
% FMINSEARCH only needs sse, but we want to plot the FittedCurve at the end.
    function [sse, FittedCurve] = guass2dfun(params)
        A = params(1);
        x0 = params(2);
        y0 = params(3);
        sig = params(4);
        bg = params(5);
        FittedCurve = A .* exp(-((xy_data(:,1)-x0).^2 + (xy_data(:,2)-y0).^2)./(2*sig^2)) + bg;
        ErrorVector = FittedCurve - reshape(data,[size(data,1)^2,1]);
        sse = sum(ErrorVector .^ 2);
    end
end
```
