## Supplementary material for "Microvillar cartography: a super-resolution single-molecule imaging method to map the positions of membrane proteins with respect to cellular surface topography": Analysis MATLAB codes: FitPTDriftcorrect_moviebymovie_Range.m

```
%Function to compare before and after correction; 
% use it for the all fiducial markers as well; 
% input ReferencePT contains correction factors for every frames 
% this is for fine tunning case; 
% FitPT; ReferecePT should be cell type; 

function [CorrectedFitsetPT]=FitPTDriftcorrect_moviebymovie_Range(FitPT,ReferencePT,x_range,y_range,SaveName,Do_merge,showmarker,markersize)
%function [CorrectedFitsetPT]=FitPTDriftcorrect_moviebymovie_Range(FitsetPT,ReferencePT,x_range,y_range,SaveName,Do_merge,showmarker,markersize)
if size (FitPT,2) ~= size(ReferencePT,1) 
error('Reference Point should be same size as sets of FitPT')
end

if ~iscell(FitPT) 
error('FitPT should be a cell type')
end


movienum= size(ReferencePT,1);

%end
% FitSingleMol=zeros(1,10);
% CMSingleMol=zeros(1,10);

for kk=1:movienum
%=find(FitsetPT{kk}(:,1);
CorrectedFitPT{kk}=FitPT{kk};
CorrectedFitPT{kk}(:,2)=FitPT{kk}(:,2)-ReferencePT(kk,2);
CorrectedFitPT{kk}(:,3)=FitPT{kk}(:,3)-ReferencePT(kk,3);
CorrectedFitPT{kk}(:,end+1)=kk; % write movie number

end

% ConFITmol = FitsetPT;   
% ConCorrFitmol = CorrectedFitsetPT; 
%   

%FitsetPT=;
FitsetPT=cell2mat(FitPT');
CorrectedFitsetPT=cell2mat(CorrectedFitPT');
%CorrectedFitsetPT=cell2mat(FitPT');

if Do_merge==1
    MerFig = figure('Name','Before-After');
             FM=FitsetPT(:,2);
             FN=FitsetPT(:,3);
             CM=CorrectedFitsetPT(:,2);
             CN=CorrectedFitsetPT(:,3);
           
             plot(FN,FM,'.','MarkerSize',0.07,'Color','green');  
             hold on
        
             plot(CN,CM,'.','MarkerSize',0.07,'Color','red'); 
             axis equal
             xlim([x_range(1) x_range(2)]);
             ylim([y_range(1) y_range(2)]);       
             saveas(gcf,['xcBef_Aft' SaveName],'fig');
             %saveas(gcf,['xcBef_Aft' SaveName],'jpg');
             saveas(gcf,['xcBef_Aft' SaveName],'tif');
             if showmarker==1
              %  markersize=1000;
                 addscale3_um_range_label(x_range,y_range,markersize,1)
             end
             
             
             hold off
%            view([90 90]);
%              xlim([mxm mxp]);
%              ylim([mym myp]);
%              view([90 90]);


%            [scalebar]=addscale_gray(100,mxm+0.5,mym+0.5,1,1);

end 
             figure;
             CM=CorrectedFitsetPT(:,2);
             CN=CorrectedFitsetPT(:,3);
             plot(CN,CM,'.','MarkerSize',0.07,'Color','red');   
             axis equal
             xlim([x_range(1) x_range(2)]);
             ylim([y_range(1) y_range(2)]); 
             
             saveas(gcf,['xcCorrectFit' SaveName],'fig');
             saveas(gcf,['xcCorrectFit' SaveName],'tif');

             if showmarker==1
              %  markersize=1000;
                 addscale3_um_range_label(x_range,y_range,markersize,1)
             end
%             close 
   
             %saveas(gcf,['Fiducial' num2str(Nth)],'fig');
%              saveas(gcf,['Fiducial' num2str(Nth)],'jpg');
            % close 
             


% close
%            
end
```
