## Supplementary material for "Microvillar cartography: a super-resolution single-molecule imaging method to map the positions of membrane proteins with respect to cellular surface topography": Analysis MATLAB codes: fits2frames_Y.m

```
function frames_out = fits2frames_Y(fid_in, frames2take_in, method, cut_in)
% Get Frames_out(data of Nth Frame) based on Method
% located inside fits_shifter_single
% Gabi made 
%syntax:
%  frames_out=spe2frames(fid, frames2take, method, cut)
%
% INPUT
% fid_in         - a scalar (file  identifier) / a valid file name (path inculded) / [] {if empty an open file dialog box will be opend
%                  or a {n,2} cell array were:  elements of the 1st column are the file names strings or file identifiers
%                                           and elements of the 2nd column are the [1,2] shift vector companasating for small mouvments between files
%
%               NOTE:  fits2frames will close, at the end of the run, all the file stated by a string file name but not the file stated by file identifiers
%
% frames2take    - [N,1] array : The list of frame numbers to be read from the file and processed.
%                                Frames will be procesed according to the order stated.
%                                If empty all frames will be processed.
%                                If all frames are negative all frames will be processed accept abs(frames2take).
%                                If zero is stated a zero padded frame will be produced and put into the output.
%                                Non-negative and negative frame numbers cannot be givven in the same frames2take vector.
%                                If fid_in is a cell array with length q frames2take shuld be a cell array of vectors of frames2take
%
% Method (optional) - the type of data processing wanted
%       * by default: method gives frames with fast method
% data wanted can be:
%                       1. Fast      - read all frames between frames2take(1) and frames2take(end) as a block and output a 3D array of frames
%                       2. Framers   - read all frames stated in frames2take frame by frame and creates a 3D array of frames
%                       3. MinMax    - calcualte the min and max reading in the all the frames stated in frames2take
%                       3. Max       - calcualte the maximum projection of the frames stated in frames2take
%                       4. Min       - calcualte the minimun projection of the frames stated in frames2take
%                       5. Sum       - sums up the pixels frame by frame
%                       4. Mean, Avg - calcualte the average of all the frames stated in frames2take
%                       6. Std       - calcualte the std value of each pixel in the file along the time trajectory
%                       7. Stat      - calculate the Mean the Std the Min and Max projections and send them as an output to a cell array
%                       8. Descend   - sort the intensty trajectory (i.e. 3rd dimension) of each pixel in descending order producing a sorted 3D array
%                                      and output a 3D array of the sorted imiges in descending order in the output only elements between n-th and m-th order
%                                      (i.e. elemnts between the n-th element smaller than the maximum and m-th element  smaller than the maximum)
%                                      m and n (high_cut and low_cut respectivily) high_cut and low_cut are delived by the user by stating {1,2} cell array
%                                      with the string 'Descend' at the 1st element and a [1,2] vector of [high_cut, low_cut] at the 2nd element.
%                       9. Fetch    - creates an experiment structure array with the molecules trajectory
%                                     in order for this method to work this metohd shuld be states as a cell array: {'Fetch',CetresList,RegionSize}
%                                     were:
%                                     CentresList - a [n,2] list of coordinate indices loacting the center pixel of each molecule
%                                     RegionSize  - a [1,2] the number of pixels before and after the center pixel to be taken into the region
%                                                   (i.e. RegionSize=[2,2] will lead to [5,5, FrameNum] regions) if not stated or empty defoult RegionSize=[2,2]
%
%
%                                     output will have the following fields:
%                                     frames_out.Molecules(q).Molecule.CenterIndices    : [1,2] center indices
%                                     frames_out.Molecules(q).Molecule.RegionIndices    : [1,4] region indices
%                                     frames_out.Molecules(q).Molecule.Region           : [RegionSize(1),RegionSize(2),FrameNum] region trajectory
%                                     frames_out.Molecules(q).Molecule.Max              : [1, FrameNum]                          max value trajectory
%                                     frames_out.Molecules(q).Molecule.Center           : [1, FrameNum]                          center vlaue trjaectory
%                                     frames_out.Molecules(q).Molecule.DataSourceNumber : 1  - this scalar connects the molecule data to the data file they were
%                                                                                              extracted from (the index of frames_out.DataSource(ind).DataSource)
%                                                                                              In tihs case all molecules come from the same file hance
%                                                                                              there is only one DataSource
%
%                                     frames_out.DataSource(1).DataSource    :  Informetion about the data file from which molecules were extracted
%                                                                               DataSource containes the info if the last fits file opend fits file
%                                                                               info.FitsFiles  - fid_in vector
%                                                                               info.Frame2Take - frames2take_in (inculding the shift vectors)
%                                                                               info.Image      - a 3D array of mean images taken from each file opend
%                                                                               info.Cut        - the cut vector
%
%                                     NOTE: 1) This Fecth method ignors frames2take and will take all the frames in the file
%                                           2) If a cut vector is provided CentresList should be in the coordinate systme after the cut
%
% cut_in (optional) - [1x4] array (from the word cutout) : the indices that form the edges of the region that form the hole frame
%                 that will be read by the program and saved to the *.mat file (sometimes specified as region of analesis-roa)
%                                   cut(1)-----cut(3)
%                                    |          |
%                                    |          |
%                                   cut(2)-----cut(4)   thus: frame(cut(1):cut(2), cut(3):cut(4))
%       * by default: cut=cut=[1,s_frame(1),1,s_frmae(2)]
% OUTPUT
% frames_OUT - a 3D array of the frames specified by 'frames2read'


switch nargin
    case 0
        fid_in=[];
        frames2take=[];
        method=[];
        cut_in=[];

    case 1
        frames2take=[];
        method=[];
        cut_in=[];
        %cash=[];
    case 2
        method=[];
        cut_in=[];
        %cash=[];
    case 3
        cut_in=[];
        %cash=[];
        %case 4
        %cash=[];
end

if isempty(method); method='FAST'  ; end

if isempty(fid_in)
    [FileName,PathName] = uigetfile({'*.fits'; '*.fit'; '*.*'},'Please select a movie to open');
    fid_in=[PathName,FileName];
end


if ~iscell(fid_in); fid_in={fid_in}; end

fid=[];
for q_files = 1:size(fid_in,1)
    if ~ischar(fid_in{q_files,1});
        fid(q_files) = fid_in{q_files,1};
    else
        fid(q_files) = fopen(fid_in{q_files,1},'r','ieee-be');
    end
end
% MILA (NOTE only): check if fid_in is cell array with shift_vector in
                                                        % second coloumn
if size(fid_in,2)==1
    shift_vect=cell(size(fid_in,1),1);
else
    shift_vect=fid_in(:,2);
end


%% Methods
if iscell(method)
    method_parameters=method(2:end);
    method=method{1};
end

switch upper(method)
    case 'FAST' % creates a 3D array of frames with frames: frames2take(1):frames2take(2)
        % this is the deafult method
        % this method is much more time efficient than 'FRAMES'
        frames_out=[];
        for q_files=1:size(fid,1)
            [info,frames2take,cut]=arrange_IO_data(fid(q_files),frames2take_in,cut_in,q_files);
            DataStart    =info.PrimaryData.Offset;
            DataType     =info.PrimaryData.DataType;
            FrameLength  =info.PrimaryData.Size(1);
            FrameWidth   =info.PrimaryData.Size(2);
            FrameNum     =info.PrimaryData.Size(3);
            BitesInFrame =prod(info.PrimaryData.Size)*precisionSize(DataType)/FrameNum;
            bscale       =info.PrimaryData.Slope;
            bzero        =info.PrimaryData.Intercept;

            if frames2take(1)>frames2take(end);  error('the ''FAST'' opening method must get frames in an ascending order'); end

            StartFrame=frames2take(1);
            num_frames2take  =1+frames2take(end)-frames2take(1);

            fseek(fid, DataStart+BitesInFrame*(StartFrame-1), 'bof');
            frames2put=fread(fid,FrameLength*FrameWidth*num_frames2take,['*',DataType]);
            frames2put=permute(reshape(frames2put,FrameLength,FrameWidth,[]),[2,1,3]);
            frames2put=frames2put(cut(1):cut(2),cut(3):cut(4),:);
            frames2put=shift_mat(frames2put, shift_vect{q_files});
            if bzero~=0 || bscale~=1
                frames2put=bscale*double(frames2put)+bzero;
            end

            frames_out=cat(3,frames_out,frames2put);
        end %q_files

    case 'FRAMES'  % creates a 3D array of frames with the frames stated in frames2take
        frames_out=[];
        for q_files=1:size(fid,1)
            [info,frames2take,cut]=arrange_IO_data(fid(q_files),frames2take_in,cut_in,q_files);
            DataStart    =info.PrimaryData.Offset;
            DataType     =info.PrimaryData.DataType;
            FrameLength  =info.PrimaryData.Size(1);
            FrameWidth   =info.PrimaryData.Size(2);
            FrameNum     =info.PrimaryData.Size(3);
            BitesInFrame =prod(info.PrimaryData.Size)*precisionSize(DataType)/FrameNum;
            bscale       =info.PrimaryData.Slope;
            bzero        =info.PrimaryData.Intercept;

            num_frames2take=numel(frames2take);
            zero_frame_OUT=zeros(1+cut(2)-cut(1),1+cut(4)-cut(3),DataType);

            frames_out_q=repmat(zero_frame_OUT, [1,1,num_frames2take]);
            for q=1:num_frames2take   %
                if frames2take(q)==0
                else
                    fseek(fid, DataStart+BitesInFrame*(frames2take(q)-1), 'bof');
                    frame2put=(fread(fid,[FrameLength,FrameWidth],['*',DataType])).';
                    frames_out_q(:,:,q)=frame2put(cut(1):cut(2), cut(3):cut(4));
                end


                frames_out_q=shift_mat(frames_out_q, shift_vect{q_files});
                if bzero~=0 || bscale~=1
                    frames_out_q=bscale*double(frames_out_q)+bzero;
                end

                frames_out=cat(3,frames_out,frames_out_q);
            end % q
        end % q_files

    case 'MINMAX'
        Mn=[];
        Mx=[];
        for q_files=1:size(fid,1)
            [info,frames2take,cut]=arrange_IO_data(fid(q_files),frames2take_in,cut_in,q_files);
            DataStart    =info.PrimaryData.Offset;
            DataType     =info.PrimaryData.DataType;
            FrameLength  =info.PrimaryData.Size(1);
            FrameWidth   =info.PrimaryData.Size(2);
            FrameNum     =info.PrimaryData.Size(3);
            BitesInFrame =prod(info.PrimaryData.Size)*precisionSize(DataType)/FrameNum;
            bscale       =info.PrimaryData.Slope;
            bzero        =info.PrimaryData.Intercept;

            num_frames2take=numel(frames2take);
            for q=1:num_frames2take   %
                if frames2take(q)==0
                else
                    fseek(fid, DataStart+BitesInFrame*(frames2take(q)-1), 'bof');
                    frame2put=(fread(fid,[FrameLength,FrameWidth],[DataType,'=>double'])).';
                    frame2put=frame2put(cut(1):cut(2), cut(3):cut(4));
                    frames2put=shift_mat(frames2put, shift_vect{q_files});
                    if bzero~=0 || bscale~=1
                        frames2put=bscale*frames2put+bzero;
                    end
                    Mn=min([Mn; frame2put(:)]);
                    Mx=max([Mx; frame2put(:)]);
                end
            end % num_frames2take
            frames_out=[Mn,Mx];
        end % q_files

    case 'MAX'  % the max projection along the frames
        frames_out=-inf;
        for q_files=1:size(fid,1)
            [info,frames2take,cut]=arrange_IO_data(fid(q_files),frames2take_in,cut_in,q_files);
            DataStart    =info.PrimaryData.Offset;
            DataType     =info.PrimaryData.DataType;
            FrameLength  =info.PrimaryData.Size(1);
            FrameWidth   =info.PrimaryData.Size(2);
            FrameNum     =info.PrimaryData.Size(3);
            BitesInFrame =prod(info.PrimaryData.Size)*precisionSize(DataType)/FrameNum;
            bscale       =info.PrimaryData.Slope;
            bzero        =info.PrimaryData.Intercept;

            num_frames2take=numel(frames2take);
            for q=1:num_frames2take   %
                if frames2take(q)==0
                else
                    fseek(fid, DataStart+BitesInFrame*(frames2take(q)-1), 'bof');
                    frame2put=(fread(fid,[FrameLength,FrameWidth],['*',DataType])).';
                    frame2put=frame2put(cut(1):cut(2), cut(3):cut(4));
                    frame2put=shift_mat(frame2put, shift_vect{q_files});
                end

                if bzero~=0 || bscale~=1
                    frame2put=bscale*double(frame2put)+bzero;
                end
                frames_out=max(frames_out,frame2put);
            end
        end % q_files

    case 'MIN'  % the min projection along the frames
        frames_out=inf;
        for q_files=1:size(fid,1)
            [info,frames2take,cut]=arrange_IO_data(fid(q_files),frames2take_in,cut_in,q_files);
            DataStart    =info.PrimaryData.Offset;
            DataType     =info.PrimaryData.DataType;
            FrameLength  =info.PrimaryData.Size(1);
            FrameWidth   =info.PrimaryData.Size(2);
            FrameNum     =info.PrimaryData.Size(3);
            BitesInFrame =prod(info.PrimaryData.Size)*precisionSize(DataType)/FrameNum;
            bscale       =info.PrimaryData.Slope;
            bzero        =info.PrimaryData.Intercept;

            num_frames2take=numel(frames2take);
            for q=1:num_frames2take   %
                if frames2take(q)==0
                else
                    fseek(fid, DataStart+BitesInFrame*(frames2take(q)-1), 'bof');
                    frame2put=(fread(fid,[FrameLength,FrameWidth],['*',DataType])).';
                    frame2put=frame2put(cut(1):cut(2), cut(3):cut(4));
                    frame2put=shift_mat(frame2put, shift_vect{q_files});
                end

                if bzero~=0 || bscale~=1
                    frame2put=bscale*double(frame2put)+bzero;
                end
                frames_out=min(frames_out,frame2put);
            end
        end % q_files

    case 'SUM'  % the sum of the frames
        frames_out=0;
        for q_files=1:size(fid,1)
            [info,frames2take,cut]=arrange_IO_data(fid(q_files),frames2take_in,cut_in,q_files);
            DataStart    =info.PrimaryData.Offset;
            DataType     =info.PrimaryData.DataType;
            FrameLength  =info.PrimaryData.Size(1);
            FrameWidth   =info.PrimaryData.Size(2);
            FrameNum     =info.PrimaryData.Size(3);
            BitesInFrame =prod(info.PrimaryData.Size)*precisionSize(DataType)/FrameNum;
            bscale       =info.PrimaryData.Slope;
            bzero        =info.PrimaryData.Intercept;

            num_frames2take=numel(frames2take);
            for q=1:num_frames2take   %
                if frames2take(q)==0
                else
                    fseek(fid, DataStart+BitesInFrame*(frames2take(q)-1), 'bof');
                    frame2put=(fread(fid,[FrameLength,FrameWidth],[DataType,'=>double'])).';
                    frame2put=frame2put(cut(1):cut(2), cut(3):cut(4));
                    frame2put=shift_mat(frame2put, shift_vect{q_files});
                end

                if bzero~=0 || bscale~=1
                    frame2put=bscale*frame2put+bzero;
                end
                frames_out=frames_out+frame2put;
            end
        end % q_files

    case {'MEAN','AVG','AVERAGE'}
        frames_out=0;
        frames_total=0;
        for q_files=1:size(fid,1)
            [info,frames2take,cut]=arrange_IO_data(fid(q_files),frames2take_in,cut_in,q_files);
            DataStart    =info.PrimaryData.Offset;
            DataType     =info.PrimaryData.DataType;
            FrameLength  =info.PrimaryData.Size(1);
            FrameWidth   =info.PrimaryData.Size(2);
            FrameNum     =info.PrimaryData.Size(3);
            BitesInFrame =prod(info.PrimaryData.Size)*precisionSize(DataType)/FrameNum;
            bscale       =info.PrimaryData.Slope;
            bzero        =info.PrimaryData.Intercept;

            num_frames2take=numel(frames2take);
            for q=1:num_frames2take   %
                if frames2take(q)==0
                else
                    fseek(fid, DataStart+BitesInFrame*(frames2take(q)-1), 'bof');
                    frame2put=(fread(fid,[FrameLength,FrameWidth],[DataType,'=>double'])).';
                    frame2put=frame2put(cut(1):cut(2), cut(3):cut(4));
                    frame2put=shift_mat(frame2put, shift_vect{q_files});
                end

                if bzero~=0 || bscale~=1
                    frame2put=bscale*frame2put+bzero;
                end
                %frames_total=frames_total+num_frames2take;
                frames_out=frames_out+frame2put;
            end
            frames_total=frames_total+num_frames2take;
        end % q_files
        frames_out=frames_out/frames_total;

    case 'STD' % the standard deviation of the all the frames
        frames_out_sum=0;
        frames_out_sum_sq=0;
        frames_total=0;
        for q_files=1:size(fid,1)
            [info,frames2take,cut]=arrange_IO_data(fid(q_files),frames2take_in,cut_in,q_files);
            DataStart    =info.PrimaryData.Offset;
            DataType     =info.PrimaryData.DataType;
            FrameLength  =info.PrimaryData.Size(1);
            FrameWidth   =info.PrimaryData.Size(2);
            FrameNum     =info.PrimaryData.Size(3);
            BitesInFrame =prod(info.PrimaryData.Size)*precisionSize(DataType)/FrameNum;
            bscale       =info.PrimaryData.Slope;
            bzero        =info.PrimaryData.Intercept;

            num_frames2take=numel(frames2take);
            for q=1:num_frames2take   %
                if frames2take(q)==0
                else
                    fseek(fid, DataStart+BitesInFrame*(frames2take(q)-1), 'bof');
                    frame2put=(fread(fid,[FrameLength,FrameWidth],[DataType,'=>double'])).';
                    frame2put=frame2put(cut(1):cut(2), cut(3):cut(4));
                    frame2put=shift_mat(frame2put, shift_vect{q_files});
                end

                if bzero~=0 || bscale~=1
                    frame2put=bscale*frame2put+bzero;
                end
                frames_total=frames_total+num_frames2take;
                frames_out_sum=frames_out_sum+frame2put;
                frames_out_sum_sq = frames_out_sum_sq + frame2put.^2;
            end
        end % q_files
        frames_out = sqrt(frames_out_sum_sq/frames_total -(frames_out_sum/frames_total).^2);

    case 'STAT' % the standard deviation of the all the frames
        frames_out_MN    = inf;
        frames_out_MX    =-inf;
        frames_out_sum   = 0;
        frames_out_sum_sq= 0;
        frames_total     = 0;
        for q_files=1:size(fid,1)
            [info,frames2take,cut]=arrange_IO_data(fid(q_files),frames2take_in,cut_in,q_files);
            DataStart    =info.PrimaryData.Offset;
            DataType     =info.PrimaryData.DataType;
            FrameLength  =info.PrimaryData.Size(1);
            FrameWidth   =info.PrimaryData.Size(2);
            FrameNum     =info.PrimaryData.Size(3);
            BitesInFrame =prod(info.PrimaryData.Size)*precisionSize(DataType)/FrameNum;
            bscale       =info.PrimaryData.Slope;
            bzero        =info.PrimaryData.Intercept;

            num_frames2take=numel(frames2take);
            for q=1:num_frames2take   %
                if frames2take(q)==0
                else
                    fseek(fid, DataStart+BitesInFrame*(frames2take(q)-1), 'bof');
                    frame2put=(fread(fid,[FrameLength,FrameWidth],[DataType,'=>double'])).';
                    frame2put=frame2put(cut(1):cut(2), cut(3):cut(4));
                    frame2put=shift_mat(frame2put, shift_vect{q_files});
                end

                if bzero~=0 || bscale~=1
                    frame2put=bscale*frame2put+bzero;
                end
                frames_total=frames_total+num_frames2take;
                frames_out_sum=frames_out_sum+frame2put;
                frames_out_sum_sq = frames_out_sum_sq + frame2put.^2;
                frames_out_MX=max(frames_out_MX,frame2put);
                frames_out_MN=min(frames_out_MN,frame2put);
            end
        end % q_files
        frames_out = { 'AVG', 'STD', 'MIN', 'MAX';
            frames_out_sum/frames_total, ...
            sqrt(frames_out_sum_sq/frames_total -(frames_out_sum/frames_total).^2), frames_out_MN, frames_out_MX };

        %     case 'DESCEND'
        %         high_cut=method_parameters{1}(1);
        %         low_cut =method_parameters{1}(2);
        %         frames_out=repmat(zero_frame_OUT, [1,1,2*low_cut]);
        %         h = waitbar(0,'Please wait...opening fits file');
        %         for q=1:num_frames2take   %
        %
        %             if frames2take(q)==0
        %             else
        %                 fseek(fid, DataStart+BitesInFrame*(frames2take(q)-1), 'bof');
        %                 frame2put=(fread(fid,[FrameLength,FrameWidth],['*',DataType])).';
        %                 frames_out(:,:,low_cut+1+mod(q-1,low_cut))=frame2put(cut(1):cut(2), cut(3):cut(4));
        %                 if mod(q-1,low_cut)+1 == low_cut || q==num_frames2take
        %                     frames_out=sort(frames_out,3, 'Descend' );
        %                     waitbar(q/num_frames2take,h);
        %                 end
        %             end
        %         end
        %
        %         if bzero~=0 || bscale~=1
        %             frames_out=bscale*frames_out+bzero;
        %         end
        %
        %         frames_out=mean(frames_out(:,:,high_cut:low_cut),3);
        %         close(h);

    case 'FETCH' % taking the data of [m,n,all_frames_in_file] around the list of points
        CentersList=method_parameters{1};
        num_of_regions=size(CentersList,1);

        if length(method_parameters)>=2
            RegionSize=method_parameters{2};
        else
            RegionSize=[];
        end

        if isempty(RegionSize)
            RegionSize=[2,2];
        elseif all(size(RegionSize))==1
            RegionSize=[RegionSize,RegionSize];
        end

        sum_seg=0;
        frames_total=0;
        image_all_files=[];
        
        for q_files=1:numel(fid)
            
            [info,frames2take,cut]=arrange_IO_data(fid(q_files),[],cut_in,q_files);
            DataStart    =info.PrimaryData.Offset;
            DataType     =info.PrimaryData.DataType;
            FrameLength  =info.PrimaryData.Size(1);
            FrameWidth   =info.PrimaryData.Size(2);
            FrameNum     =info.PrimaryData.Size(3);
            BitesInFrame =prod(info.PrimaryData.Size)*precisionSize(DataType)/FrameNum;
            bscale       =info.PrimaryData.Slope;
            bzero        =info.PrimaryData.Intercept;

            if q_files==1
                sz_frame=[1+cut(2)-cut(1),1+cut(4)-cut(3)];

                RegionsList=[CentersList(:,1)-RegionSize(1),CentersList(:,1)+RegionSize(1), CentersList(:,2)-RegionSize(2),CentersList(:,2)+RegionSize(2)];
                good_regions=~( any(RegionsList<1,2) | (RegionsList(:,1) > sz_frame(1)) | (RegionsList(:,2) > sz_frame(1)) | (RegionsList(:,3) > sz_frame(2)) | (RegionsList(:,4) > sz_frame(2)) );
                %PositionList=[fliplr(CentersList)-(RegionSize+.5), repmat(RegionSize*2+1,[size(CentersList,1),2])];

                ind=find(good_regions);
                num_of_good_regions=length(ind);
                %if num_of_good_regions~=size(CentersList,1);
                 %   warning('Regions lost due to shalow bounderies');
                %end

                CentersList=CentersList(ind,:);
                RegionsList=RegionsList(ind,:);
            end % q_files==1

            seg_size=min(200,FrameNum-1);    %  defines the maximal number of frames in a segment
            %total_seg_num=size([1:seg_size:FrameNum],2); % the number of all segments;
            %data_all={};     %  each line in 'data_all' contains the data of a region in 'good_regions'
         
            for seg_start=1:seg_size:FrameNum       %    this loop runs through all the segments
                seg_end=min(FrameNum,(seg_start+seg_size-1));   % calculatinting the number of frames in the current segment
                %   i.e. seg_size or the remaning frames at the end of the spe file

                num_frames2take  =1+seg_end-seg_start;
                fseek(fid(q_files), DataStart+BitesInFrame*(seg_start-1), 'bof'); %MILA: added index to fid
                seg_frames=fread(fid(q_files),FrameLength*FrameWidth*num_frames2take,['*',DataType]);
                seg_frames=permute(reshape(seg_frames,FrameLength,FrameWidth,[]),[2,1,3]);
                seg_frames=seg_frames(cut(1):cut(2),cut(3):cut(4),:);
                seg_frames=shift_mat(seg_frames, shift_vect{q_files});

                if bzero~=0 || bscale~=1
                    seg_frames=bscale*double(seg_frames)+bzero;
                end

                for q=1:num_of_good_regions      % this loop passes through all the 'good_regions' and collects the data
                    if seg_start==1 && q_files==1  % 'molecules' is initialized at the first segment of the first file
                        seg_region_tra=seg_frames(RegionsList(q,1):RegionsList(q,2), RegionsList(q,3):RegionsList(q,4), :);
                        molecules(q).Molecule.Region=seg_region_tra;
                        molecules(q).Molecule.Max=(squeeze(max((max(seg_region_tra)))))';
                        molecules(q).Molecule.Center=(squeeze((seg_region_tra(RegionSize(1)+1,RegionSize(2)+1,:))))';
                        molecules(q).Molecule.CenterIndices=CentersList(q,:);
                        molecules(q).Molecule.RegionIndices=RegionsList(q,:);
                        molecules(q).Molecule.DataSourceNumber=1;

                    else         %  append the data in 'data_all' when seg>1
                        seg_region_tra=seg_frames(RegionsList(q,1):RegionsList(q,2), RegionsList(q,3):RegionsList(q,4), :);
                        seg_max_tra=(squeeze(max((max(seg_region_tra)))))';
                        seg_center_tra=(squeeze((seg_region_tra(RegionSize(1)+1,RegionSize(2)+1,:))))';

                        molecules(q).Molecule.Center =[molecules(q).Molecule.Center, seg_center_tra];
                        molecules(q).Molecule.Max    =[molecules(q).Molecule.Max, seg_max_tra];
                        molecules(q).Molecule.Region =cat(3, molecules(q).Molecule.Region, seg_region_tra);
                    end
                end % q (good regions)
                sum_seg=sum_seg+sum(seg_frames,3);
            end % segments
            image_all_files=cat(3,image_all_files,frames_total/FrameNum);
           %%%%######################### should insert possibility to join several files
            end % q_files

        info.FitsFiles =fid_in;
        info.Frame2Take=frames2take_in;
        info.Image     =image_all_files;
        info.Cut       =cut_in;

        frames_out.Molecules=molecules;
        frames_out.DataSource(1).DataSource=info;

    otherwise
        error([method,' is an unknown method']);
end % switch method

for q_files=1:size(fid_in,1)    % closing all the files that were stated by the user as file name (i.e file that were open in this run of the function)
    if ischar(fid_in{q_files})
        fclose(fid(q_files));
    end
end


function [info,frames2take_out,cut_out]=arrange_IO_data(fid, frames2take_in, cut_in,q_files)
% this function cheks the validity of frames2take and cut
% it also takes care of the defoult vlaues of frames2take and cut in case they are empty
% the function also sed as output the info data of the fits file

if iscell(fid)
    fid=fid{q_files};
elseif numel(fid)>1
    fid=fid(q_files);
end


if isempty(frames2take_in) && iscell(frames2take_in) % GAF 20-DEC-2009
    frames2take=[];
elseif iscell(frames2take_in)
    frames2take=frames2take_in{q_files};
else
    frames2take=frames2take_in;
end

if isempty(cut_in) && iscell(cut_in) % GAF 20-DEC-2009
    cut=[];
elseif iscell(cut_in)
    cut=cut_in{q_files};
else
    cut=cut_in;
end

info = fitsinfo_internal(fid); % fitsinfo_internal is a copy pase of the original fitsinfo.m MATLAB function chnged so it will work with a fid of a preopend file

info.PrimaryData.Size=[info.PrimaryData.Size(1:end),1];
FrameLength  =info.PrimaryData.Size(1);
FrameWidth   =info.PrimaryData.Size(2);
FrameNum     =info.PrimaryData.Size(3);
if numel(info.PrimaryData.Size)>=5
    warinig('This function is written for [x,y,t] moives')
end

if isempty(frames2take)            ; frames2take=(1:FrameNum)            ; end
if isempty(cut)                    ; cut=[1,FrameWidth,1,FrameLength]    ; end % the transpose here is due to the fact all the data in the file is ordered colomn wise

% Input data validity check
if any(frames2take>FrameNum); error('frames asked by the user are larger than the number of frames in the file'); end
if any( isnan(frames2take)) || any( floor(frames2take)~=frames2take); error('Frame numbers must be proper indices !'); end
if all(frames2take<0)
    FramesNot2Take=-frames2take;
    frames2take=(1:FrameNum);
    for q=FramesNot2Take
        ind=(frames2take==q);
        frames2take(ind)=[];
    end
elseif any(frames2take<0)
    error('Negative frame numbers cannot be givvenn with positive frame numbers !')
end

if any(cut)<=0 || any(floor(frames2take)~=frames2take) || any(~isfinite(cut)); error('All ''cut'' values must be proper indices'); end
if length(cut(:))~=4; error('''cut'' must be 4 elements array of indices'); end
if cut(1)>cut(2) || cut(3)>cut(4) ; error('''cut'' values must be ordered as follows: cut(2)>=cut(1) and cut(4)>=cut(3)'); end
if any(cut(1:2))>FrameLength || any(cut(3:4))>FrameWidth; error('Indices in ''cut'' are out of the frame size'); end

frames2take_out=frames2take;
cut_out=cut;


function frame_out=shift_mat(frame_in,shift_vect)
%
% frame_out=shif_mat(frame_in,shift_vect)
%
% this function get a 2D or a 3D matrix frame_in and shifts it according to the shift vector shift_vect
% the OUTPUT matrix frame_out has the same dimensisons as frame_in with borders padded

padding=zeros(1,1,class(frame_in));

if ~isempty(shift_vect)
    sz=size(frame_in);
    sz=[sz,1];
    frame_out=frame_in;

    % Vertical shift
    pad_mat=repmat(padding,[abs(shift_vect(1)),sz(2),sz(3)]);
    if shift_vect(1)>0
        frame_out=[pad_mat ; frame_out(1:end-shift_vect(1),:,:)];
    elseif shift_vect(1)<0
        frame_out=[frame_out(abs(shift_vect(1))+1:end,:,:) ; pad_mat];
    end


    % Horizontal shift

    % Vertical shift
    pad_mat=repmat(padding,[sz(1),abs(shift_vect(2)),sz(3)]);
    if shift_vect(2)>0
        frame_out=[pad_mat , frame_out(:,1:end-shift_vect(2),:)];
    elseif shift_vect(2)<0
        frame_out=[frame_out(:,abs(shift_vect(2))+1:end,:), pad_mat];
    end

else %shift_vect
    frame_out=frame_in;
end


function sz = precisionSize(type)
% return the precision size, in bytes.
switch type
    case {'uint8' 'char' 'bit8'}
        sz = 1;
    case {'int16' 'bit16'}
        sz = 2;
    case {'int32' 'single' 'bit32'}
        sz = 4;
    case {'double' 'bit64'}
        sz = 8;
    otherwise
        error('MATLAB:fitsread:invalidPrecisionSize', ...
            'Incorrect precision value in FITS file (%s)', type);
end

%% The next part is a cut and paste from fitsinfo.m
% the function was changed so it will work with fid and not a file name

function info = fitsinfo_internal(fid)
%FITSINFO  Information about FITS file
%
%   INFO = FITSINFO(FILENAME) returns a structure whose fields contain
%   information about the contents a FITS (Flexible Image Transport System)
%   file.  FILENAME is a string that specifies the name of the FITS file.
%
%   The INFO structure will contain the following fields:
%
%      Filename      A string containing the name of the file
%
%      FileSize      An integer indicating the size of the file in bytes
%
%      FileModDate   A string containing the modification date of the file
%
%      Contents      A cell array containing a list of extensions in the
%                    file in the order that they occur
%
%      PrimaryData   A structure containing information about the primary
%                    data in the FITS file
%
%   The PrimaryData structure will contain the following fields:
%
%      DataType      Precision of the data
%
%      Size          Array containing sizes of each dimension
%
%      DataSize      Size in bytes of primary data
%
%      MissingDataValue  Value used to represent undefined data
%
%      Intercept     Value used along with Slope to calculate true pixel
%                    values from the array pixel values using the equation:
%                    actual_value = Slope*array_value + Intercept
%
%      Slope         Value used along with Intercept to calculate true pixel
%                    values from the array pixel values using the equation:
%                    actual_value = Slope*array_value + Intercept
%
%      Offset        Number of bytes from beginning of file to location of
%                    first data value
%
%      Keywords      A number-of-keywords x 3 cell array which contain all
%                    the Keywords, Values and Comments of the header in each
%                    column.
%
%
%   A FITS file may have any number of extensions.  One or more of the
%   following fields may also be present in the INFO structure:
%
%      AsciiTable  An array of structures containing information
%                  about the Ascii Table extensions in this file
%
%      BinaryTable An array of structures containing information
%                  about any Binary Table extensions in this file
%
%      Image       An array of structures containing information
%                  about any Image extensions in this file
%
%      Unknown     An array of structures containing information
%                  about any non standard extensions in this file.
%
%   The AsciiTable structure will contain the following fields:
%
%      Rows         The number of rows in the table
%
%      RowSize      The number of characters in each row
%
%      NFields      The number of fields in each row
%
%      FieldFormat  A 1 x NFields cell containing formats in which each
%                   field is encoded.  The formats are FORTRAN-77 format codes.
%
%      FieldPrecision A 1 x NFields cell containing precision of the data in
%                     each field
%
%      FieldWidth   A 1 x NFields array containing the number of characters
%                   in each field
%
%      FieldPos     A 1 x NFields array of numbers representing the
%                   starting column for each field
%
%      DataSize     Size in bytes of the data in the ASCII Table
%
%      MissingDataValue  A 1 x NFields array of numbers used to represent
%                        undefined data in each field
%
%      Intercept    A 1 x NFields array of numbers used along with Slope to
%                   calculate actual data values from the array data values
%                   using the equation: actual_value = Slope*array_value+
%                   Intercept
%
%      Slope        A 1 x NFields array of numbers used with Intercept to
%                   calculate true data values from the array data values
%                   using the equation: actual_value = Slope*array_value+
%                   Intercept
%
%      Offset       Number of bytes from beginning of file to location of
%                   first data value in the table
%
%      Keywords     A number-of-keywords x 3 cell array containing all the
%                   Keywords, Values and Comments in the ASCII table header
%
%   The BinaryTable structure will contain the following fields:
%
%      Rows        The number of rows in the table
%
%      RowSize     The number of bytes in each row
%
%      NFields     The number of fields in each row
%
%      FieldFormat  A 1 x NFields cell containing the data type of the data
%                   in each field. The data type is represented by a
%                   FITS binary table format code.
%
%      FieldPrecision A 1 x NFields cell containing precision of the data in
%                     each field
%
%      FieldSize    A 1 x NFields array containing the number of values in
%                   the Nth field
%
%      DataSize     Size in bytes of data in the Binary Table.  Value
%                   includes any data past the main table
%
%      MissingDataValue  An 1 x NFields array of numbers used to represent
%                        undefined data in each field
%
%      Intercept    A 1 x NFields array of numbers used along with Slope to
%                   calculate actual data values from the array data values
%                   using the equation: actual_value = Slope*array_value+
%                   Intercept
%
%      Slope        A 1 x NFields array of numbers used with Intercept to
%                   calculate true data values from the array data values
%                   using the equation: actual_value = Slope*array_value+
%                   Intercept
%
%      Offset       Number of bytes from beginning of file to location of
%                   first data value
%
%      ExtensionSize    The size in bytes of any data past the main table
%
%      ExtensionOffset  The number of bytes from the beginning of the file to
%                       the location of the data past the main table
%
%      Keywords      A number-of-keywords x 3 cell array containing all the
%                    Keywords, Values and Comments in the Binary table header
%
%   The Image structure will contain the following fields:
%
%      DataType      Precision of the data
%
%      Size          Array containing sizes of each dimension
%
%      DataSize      Size in bytes of data in the Image extension
%
%      MissingDataValue  Value used to represent undefined data
%
%      Intercept     Value used along with Slope to calculate true pixel
%                    values from the array pixel values using the equation:
%                    actual_value = Slope*array_value + Intercept
%
%      Slope         Value used along with Intercept to calculate true pixel
%                    values from the array pixel values using the equation:
%                    actual_value = Slope*array_value + Intercept
%
%      Offset        Number of bytes from beginning of file to location of
%                    first data value
%
%      Keywords      A number-of-keywords x 3 cell array containing all the
%                    Keywords, Values and Comments in the Image header
%
%   The Unknown structure will contain the following fields:
%
%      DataType      Precision of the data
%
%      Size          Array containing sizes of each dimension
%
%      DataSize      Size in bytes of data in the extension
%
%      Intercept     Value used along with Slope to calculate true data
%                    values from the  array data values using the equation:
%                    actual_value = Slope*array_value + Intercept
%
%      Slope         Value used along with Intercept to calculate true data
%                    values from the array data values using the equation:
%                    actual_value = Slope*array_value + Intercept
%
%      MissingDataValue  Value used to represent undefined data
%
%      Offset        Number of bytes from beginning of file to location of
%                    first data value
%
%      Keywords      A number-of-keywords x 3 cell array containing all the
%                    Keywords, Values and Comments in the extension header
%
%   See also FITSREAD.

%   Copyright 2001-2006 The MathWorks, Inc.
%   $Revision: 1.1.6.7 $  $Date: 2007/11/26 20:36:09 $


info = buildDefaultInfo;
info = getFileSystemInfo(info, fid);

% Read primary header.
% All FITS files are required to have a primary header.
info.PrimaryData = readheaderunit(fid, info.FileSize);
info.Contents{1} = 'Primary';

if (~isStandardFile(info))

    msg1 = sprintf('File %s is not a standard FITS file.', filename);
    msg2 = sprintf('FITSINFO will attempt to determine the contents.');

    warning('MATLAB:fitsinfo:nonstandardFile', '%s\n%s', msg1, msg2);

end

% Skip the primary data

if (hasExtensions(info))

    success = skipHduData(fid, computeDataSize(info.PrimaryData));

    if (success)

        info = readExtensions(fid, info);

    end

end


function info = readheaderunit(fid, fileSize)
%Read a header unit from a FITS file. File curser is placed at the end of
%the header (an integral number of 2880 bytes).
%
% INFO header information
% HEADERTYPE Type of data this is describing

%Read first card image to determine the type of header unit.
[keyword, value, comment] = readCardImage(fid);

try

    headerType = getHeaderType(keyword, value);

catch

    fclose(fid);
    rethrow(lasterror)

end

info = createBlankHeaderInfo(headerType);
info(1).Keywords = {keyword, value, comment};

keywordNum = 1;
endFound = false;
while ((~atEOF(fid, fileSize)) && (~endFound))

    keywordNum = keywordNum + 1;

    [keyword, value, comment] = readCardImage(fid);

    switch headerType
        case 'primary'

            info = updateInfoForPrimary(info, keyword, value);

        case 'ascii'

            info = updateInfoForTable(info, keyword, value);

        case 'binary'

            info = updateInfoForBintable(info, keyword, value);

        case 'image'

            info = updateInfoForImage(info, keyword, value);

        otherwise

            info = updateInfoForGeneric(info, keyword, value);

    end

    info.Keywords(keywordNum,:) = {keyword, value, comment};

    if (strcmpi(keyword, 'end'))
        endFound = true;
    end

end

if (atEOF(fid,fileSize) && ~endFound)

    msg = sprintf('%s\n%s\n%s', ...
        'End of file found before the end of the header was reached.', ...
        'File may be an invalid FITS file or corrupt.', ...
        'Output structure may not contain complete file information.');
    warning('MATLAB:fitsinfo:earlyEOF', msg)

end

info.DataSize = getDataSize(info, headerType);

%Move file position to integral number of 2880 bytes
status = moveToNextHDU(fid);

info.Offset = ftell(fid);
if status == -1
    warning('MATLAB:fitsinfo:seekFailed', 'Seek failed because of incorrect header information.\nFile may be an invalid FITS file or corrupt.\nOutput structure may not contain complete file information.');
end
%End READHEADERUNIT

function [keyword,value,comment] = readCardImage(fid)
[card, readcount] = fread(fid,80,'uchar=>char');
card = card';
if feof(fid)
    return;
elseif readcount ~= 80
    fclose(fid);
    error('MATLAB:fitsinfo:cardbytes', 'FITS card image was not 80 bytes. FITS file may be invalid or corrupt.');
end

try
    [keyword,value,comment] = parsecard(card);
catch
    fclose(fid);
    error('MATLAB:fitsinfo:cardFormat', 'Unexpected FITS card image format. FITS file may be invalid or corrupt.');
end
%End READCARDIMAGE


function [keyword, value, comment] = parsecard(card)
%Extract the keyword, value and comment from a FITS card image

% See section 5 and appendix A of the FITS spec for details on card images.

keyword = getKeyword(card);
[value, comment]   = getValueAndComment(card);


function info = updateInfoForPrimary(info, keyword, value)

switch lower(keyword)
    case 'bitpix'
        info(1).DataType =  bitdepth2precision(value);
    case 'naxis'
        info(1).Intercept = 0;
        info(1).Slope = 1;
    case 'extend'
    case 'bzero'
        info(1).Intercept = value;
    case 'bscale'
        info(1).Slope = value;
    case 'blank'
        info(1).MissingDataValue = value;
    case 'end'
    otherwise
        %Take care of NAXISN keywords
        if findstr('naxis',lower(keyword))
            dim = sscanf(keyword(6:end),'%f');
            info(1).Size(dim) = value;
            if length(info(1).Size)==1
                info(1).Size(2) = 1;
            end
        end
end
%END UPDATEINFOFORPRIMARY

function info = updateInfoForTable(info, keyword, value)

switch lower(keyword)
    case 'end'
    case 'naxis1'
        info.RowSize = value;
    case 'naxis2'
        info.Rows = value;
    case 'tfields'
        info.NFields = value;
        info.Slope = ones(1,value);
        info.Intercept = zeros(1,value);
        info.MissingDataValue = cell(1,value);
    otherwise
        %Take care of indexed keywords
        if findstr('tform',lower(keyword))
            % Format is a FORTRAN F-77 code. The code is a characer followed by w.d,
            % the width(w) and implicit decimal place (d).
            % Valid format values in ascii table extensions
            %
            %  A - Character
            %  I - Decimal integer
            %  F - Single precision real
            %  E - Single precision real, exponential notation
            %  D - Double precision real, exponential notation
            idx = sscanf(keyword(6:end),'%s');
            switch(sscanf(value,' %c',1))
                case 'A'
                    format = 'Char';
                case 'I'
                    format = 'Integer';
                case {'E','F'}
                    format = 'Single';
                case 'D'
                    format = 'Double';
                otherwise
                    format = 'Unknown';
            end

            idxNum = sscanf(idx, '%f');

            info.FieldFormat{idxNum} = value;
            info.FieldPrecision{idxNum} = format;
            width = sscanf(value,' %*c%f');
            info.FieldWidth(idxNum) = width;
        elseif findstr('tbcol',lower(keyword))
            idx = sscanf(keyword(6:end),'%s');
            info.FieldPos(sscanf(idx, '%f')) = value;
        elseif findstr('tscal',lower(keyword));
            tscale_idx = sscanf(keyword(6:end),'%i');
            info.Slope(tscale_idx) = value;
        elseif findstr('tzero',lower(keyword));
            tzero_idx = sscanf(keyword(6:end),'%i');
            info.Intercept(tzero_idx) = value;
        elseif findstr('tnull',lower(keyword));
            tnull_idx = sscanf(keyword(6:end),'%i');
            info.MissingDataValue{tnull_idx} = value;
        end
end
%END UPDATEINFOFORTABLE

function info = updateInfoForBintable(info, keyword, value)

switch lower(keyword)
    case 'end'
    case 'naxis1'
        info.RowSize = value;
    case 'naxis2'
        info.Rows = value;
    case 'tfields'
        info.NFields = value;
        info.Slope = ones(1,value);
        info.Intercept = zeros(1,value);
        info.MissingDataValue = cell(1,value);
    case 'pcount'
        info.ExtensionSize = value;
    otherwise
        if findstr('tscal',lower(keyword));
            tscale_idx = sscanf(keyword(6:end),'%i');
            info.Slope(tscale_idx) = value;
        elseif findstr('tzero',lower(keyword));
            tzero_idx = sscanf(keyword(6:end),'%i');
            info.Intercept(tzero_idx) = value;
        elseif findstr('tnull',lower(keyword));
            tnull_idx = sscanf(keyword(6:end),'%i');
            info.MissingDataValue{tnull_idx} = value;
        elseif findstr('tform',lower(keyword))
            idx = sscanf(keyword(6:end),'%s');
            repeat = sscanf(value,'%f',1);
            if isempty(repeat)
                repeat = 1;
            end
            format = sscanf(value,' %*i%c',1);
            if isempty(format)
                format = sscanf(value,' %c',1);
            end
            % The value for tformN is a format defined in the FITS standard.  The
            % form is rTa, where r is the repeat, T is the precision and a is a
            % number undefined by the standard.
            %
            % Binary Table Format valid T (precision) characters and # bytes
            %
            %  L - Logical             1
            %  X - Bit                 *
            %  B - Unsigned Byte       1
            %  I - 16-bit integer      2
            %  J - 32-bit integer      4
            %  A - Character           1
            %  E - Single              4
            %  D - Double              8
            %  C - Complex Single      8
            %  M - Complex Double      16
            %  P - Array Descriptor    8
            switch format
                case 'L'
                    format = 'char';
                case 'X'
                    precisionWidth = 8 * ceil(double(repeat) / 8);
                    format = sprintf('bit%d', precisionWidth);
                    if repeat~=0
                        repeat = 1;
                    end
                case 'B'
                    format = 'uint8';
                case 'I'
                    format = 'int16';
                case 'J'
                    format = 'int32';
                case 'A'
                    format = 'char';
                case 'E'
                    format = 'single';
                case 'D'
                    format = 'double';
                case 'C'
                    format = 'single complex';
                case 'M'
                    format = 'double complex';
                case 'P'
                    format = 'int32';
                    if repeat~=0
                        repeat = 2;
                    end
            end

            idxNum = sscanf(idx, '%f');

            info.FieldFormat{idxNum} = value;
            info.FieldPrecision{idxNum} = format;
            info.FieldSize(idxNum) = repeat;
        end
end
%END UPDATEINFOFORBINTABLE

function info = updateInfoForImage(info, keyword, value)

switch lower(keyword)
    case 'end'
    case 'bitpix'
        info.DataType =  bitdepth2precision(value);
    case 'naxis'
        info.Intercept = 0;
        info.Slope = 1;
    case 'blank'
        info.MissingDataValue= value;
    case 'bzero'
        info.Intercept = value;
    case 'bscale'
        info.Slope = value;
    otherwise
        %NAXISN keywords
        if findstr('naxis',lower(keyword))
            dim = sscanf(keyword(6:end),'%f');
            info(1).Size(dim) = value;
        end
end
%END UPDATEINFOFORIMAGE

function info = updateInfoForGeneric(info, keyword, value)

switch lower(keyword)
    case 'bitpix'
        info.DataType =  bitdepth2precision(value);
    case 'naxis'
        info.Intercept = 0;
        info.Slope = 1;
        % case 'extend'         % This code doesn't appear to do anything,
        %  extensions = 1;      % but I'm keeping it here just 'cuz.
    case 'bzero'
        info.Intercept = value;
    case 'bscale'
        info.Slope = value;
    case 'blank'
        info.MissingDataValue = value;
    case 'pcount'
        info.PCOUNT = value;
    case 'gcount'
        info.GCOUNT = value;
    case 'end'
    otherwise
        %Take care of NAXISN keywords
        if findstr('naxis',lower(keyword))
            dim = sscanf(keyword(6:end),'%f');
            info.Size(dim) = value;
        end
end
%END UPDATEINFOFORGENERIC

function precision = bitdepth2precision(value)
switch value
    case 8
        precision = 'uint8';
    case 16
        precision = 'int16';
    case 32
        precision = 'int32';
    case -32
        precision = 'single';
    case -64
        precision = 'double';
end
%END BITDEPTH2PRECISION

function bitdepth = precision2bitdepth(precision)
switch precision
    case 'uint8'
        bitdepth = 8;
    case 'int16'
        bitdepth = 16;
    case 'int32'
        bitdepth = 32;
    case 'single'
        bitdepth = 32;
    case 'double'
        bitdepth = 64;
end
%END PRECISION2BITDEPTH

function v = atEOF(fid,fileSize)
% Sometimes FEOF does not recognize that the file identifier is at the end of
% the file.  So, the end of the file has been reached if FEOF returns true or if
% the file identifyer is at the end of the file.
v = (feof(fid) == 1) || (ftell(fid) == fileSize);


function success = skipHduData(fid,datasize)

if (datasize ~= 0)

    % Data must be multiple of 2880 bytes, so skip to the end of the
    % last block.
    if (rem(datasize,2880) == 0)

        bytes_to_seek = datasize;

    else

        bytes_to_seek = datasize + (2880 - rem(datasize,2880));

    end

    status = fseek(fid,bytes_to_seek,0);

    % It's possible that the dataset didn't have enough data or that the
    % file wasn't properly padded to fill the 2,880 byte block.
    % Processing should stop with a warning.  Other failures should
    % cause an error.
    if (status == -1)

        [err, id] = ferror(fid);

        str1 = 'Seek failed because of incorrect header information.';
        str2 = 'File may be invalid or corrupt.  Output structure may not contain complete file information.';

        OFFSETBEYONDEOF = -27;  % From bnfile/fiofcn.cpp
        if (id == OFFSETBEYONDEOF)

            warning('MATLAB:fitsinfo:seekFailed', '%s\n%s', str1, str2)

        else

            error('MATLAB:fitsinfo:seekFailed', '%s\n%s', str1, str2)

        end

        success = false;

    else

        success = true;

    end

else

    % No data to skip.
    success = true;

end


function info = buildDefaultInfo
%buildDefaultInfo   Create a structure with the default info fields.

info.Filename = '';
info.FileModDate = '';
info.FileSize = [];
info.Contents = {};
info.PrimaryData = [];


function info = getFileSystemInfo(info, fid)
%getPhysicalFileInfo   Get file system information about the FITS file.

% Get the full filename.
info.Filename = fopen(fid);

% Get info about the file from DIR.
d = dir(info.Filename);
info.FileModDate = d.date;
info.FileSize = d.bytes;


function headerInfo = createBlankHeaderInfo(headerType)

switch (headerType)
    case 'primary';

        fields = {'DataType', []
            'Size', []
            'DataSize', []
            'MissingDataValue', []
            'Intercept', []
            'Slope', []
            'Offset', []
            'Keywords', {}};

    case 'ascii';

        fields = {'Rows', []
            'RowSize', []
            'NFields', []
            'FieldFormat', {}
            'FieldPrecision', ''
            'FieldWidth', []
            'FieldPos', []
            'DataSize', []
            'MissingDataValue', {}
            'Intercept', []
            'Slope', []
            'Offset', []
            'Keywords', {}};

    case 'binary';

        fields = {'Rows', []
            'RowSize', []
            'NFields', []
            'FieldFormat', {}
            'FieldPrecision', ''
            'FieldSize', []
            'DataSize', []
            'MissingDataValue', {}
            'Intercept', []
            'Slope', []
            'Offset', []
            'ExtensionSize', []
            'ExtensionOffset', []
            'Keywords', {}};

    case 'image';

        fields = {'DataType', []
            'Size', []
            'DataSize', []
            'Offset', []
            'MissingDataValue', {}
            'Intercept', []
            'Slope', []
            'Keywords', {}};

    case 'unknown';

        fields = {'DataType', []
            'Size', []
            'DataSize', []
            'PCOUNT', []
            'GCOUNT', []
            'Offset', []
            'MissingDataValue', []
            'Intercept', []
            'Slope', []
            'Keywords', {}};

end

fields = fields';
headerInfo = struct(fields{:});


function dataSize = getDataSize(info, headerType)

switch (headerType)
    case {'primary', 'image'}

        if (~isempty(info.Size))
            dataSize = prod(info.Size) * precision2bitdepth(info.DataType)/8;
        else
            dataSize = 0;
        end

    case {'ascii', 'binary'}

        dataSize = info.RowSize * info.Rows;

    case 'unknown'

        if (~isempty(info.Size))

            dataSize = (precision2bitdepth(info.DataType) * info.GCOUNT * ...
                (info.PCOUNT + prod(info.Size))) / 8;

        end

end


function info = readExtensions(fid, info)

asciicount = 1;
binarycount = 1;
imagecount = 1;
unknowncount = 1;

while ~atEOF(fid,info.FileSize)

    if (~strcmpi(readCardImage(fid), 'xtension'))

        status = fseek(fid, 2880 - 80, 0); % Seek one fits block minus
        % one card image just read.
        if status == -1

            msg = sprintf('%s\n%s\n%s', ...
                'Seek failed because of incorrect header information.', ...
                'File may be invalid or corrupt.', ...
                'Output structure may not contain complete file information.');
            warning('MATLAB:fitsinfo:seekFailed', msg);

            return;

        end
        continue;

    else

        % Rewind one card image
        fseek(fid,-80,0);
    end

    extensionInfo = readheaderunit(fid,info.FileSize);
    if isempty(extensionInfo)
        %No extension
        fclose(fid);
        return;
    end
    switch (getExtensionType(extensionInfo))
        case 'ascii'
            info.AsciiTable(asciicount) = extensionInfo;
            info.Contents{end+1} = 'ASCII Table';
            asciicount = asciicount+1;
        case 'binary'
            info.BinaryTable(binarycount) = extensionInfo;
            info.Contents{end+1} = 'Binary Table';
            info.BinaryTable(binarycount).ExtensionOffset = info.BinaryTable(binarycount).Offset+ info.BinaryTable(binarycount).RowSize*info.BinaryTable(binarycount).Rows;
            binarycount = binarycount+1;
        case 'image'
            info.Image(imagecount) = extensionInfo;
            info.Contents{end+1} = 'Image';
            imagecount = imagecount+1;
        case 'unknown'
            info.Unknown(unknowncount) = extensionInfo;
            info.Contents{end+1} = 'Unknown';
            unknowncount = unknowncount+1;
    end % End switch headerType

    success = skipHduData(fid, computeDataSize(extensionInfo));
    if (~success)

        return

    end

end % End while ~atEOF(....)


function isCompliant = isStandardFile(info)

isCompliant = any(strcmp('SIMPLE', info.PrimaryData.Keywords(:,1)));


function extensions = hasExtensions(info)

idx = strcmpi('extend', info.PrimaryData.Keywords(:,1));

if (any(idx))

    if (strcmpi('t', info.PrimaryData.Keywords(find(idx), 2)))

        extensions = true;

    else

        extensions = false;

    end

else

    extensions = false;

end


function bytes = computeDataSize(extensionInfo)

if (isequal(getExtensionType(extensionInfo), 'binary'))

    bytes = extensionInfo.DataSize + extensionInfo.ExtensionSize;

else

    bytes = extensionInfo.DataSize;

end


function headerType = getExtensionType(extensionInfo)

headerType = getHeaderType(extensionInfo.Keywords{1,1}, ...
    extensionInfo.Keywords{1,2});


function headerType = getHeaderType(extensionKeyword, extensionValue)

switch (lower(extensionKeyword))
    case 'simple'

        headerType = 'primary';

    case 'xtension'

        switch (lower(extensionValue))
            case 'bintable'

                headerType = 'binary';

            case 'image'

                headerType = 'image';

            case 'table'

                headerType = 'ascii';

            otherwise

                headerType = 'unknown';

        end

    otherwise

        error('MATLAB:fitsinfo:unrecognizedExtension', ...
            'Cannot determine extension from keyword "%s".', ...
            extensionKeyword)

end


function keyword = getKeyword(cardImage)

%"The keyword shall be a left justified, 8-character, blank-filled, ASCII
% string with no embedded blanks."

keyword = cardImage(1:8);
keyword = keyword(~isspace(keyword));


function [value, comment] = getValueAndComment(cardImage)

% Isolate the value from a possible comment, which will be preceded by a
% slash and might contain quotes (i.e., quotes can appear in comments and
% slashes can occur in quoted values).
%
% If the value is a character string, then the value is within the first
% set of matched single quotes -- double single quotes ('') don't count.
% The comment begins after the first slash after the value.

% If no value indicator, then the card image is a commentary card image
% and a value is not present.
if ((~isequal(cardImage(9:10), '= ')) || (isCOMMENT(cardImage)))

    value = '';
    comment = cardImage(9:end);

elseif (isCharacterString(cardImage))

    [value, comment] = splitCharacterCard(cardImage);
    value = cleanupCharacterValue(value);

elseif (isTIMFILE(cardImage))

    [value, comment] = splitTIMFILECard(cardImage);

else

    [value, comment] = splitNumericCard(cardImage);
    value = cleanupNumericValue(value);
    value = convertNumericValue(value);

end

% Get rid of comment's "/" marker.  Trailing whitespace is not mentioned
% in the standard so leave it (unless it's the whole comment).
comment = stripCommentMarker(comment);
comment = cleanupCommentWhitespace(comment);


function [value, comment] = splitCharacterCard(cardImage)

quoteChar = char(39);  % 39 == '

% Look for the quotes denoting the value.
quoteIdx = find(cardImage == quoteChar);

% Temporarily set the endQuote to empty, in case the end quote is
% never found.
endQuote = [];

% The first "lone" closing single quote is the value's closing quote.
for p = 2:2:numel(quoteIdx)

    if ((p == numel(quoteIdx)) || ...
            (p == numel(quoteIdx) - 1))

        % If it's the last even-numbered quote, this is the end.
        endQuote = quoteIdx(p);

    elseif (quoteIdx(p + 1) ~= quoteIdx(p) + 1)

        % If the next odd quote is not adjacent, this is the end.
        endQuote = quoteIdx(p);
        break

    end

end

% Make sure a closing quote was found.
if (isempty(endQuote))

    warning('MATLAB:fitsinfo:missingEndQuote', ...
        'The card image was missing an ending quote.')

    endQuote = numel(cardImage);

end

% Assign the value.
value = cardImage(quoteIdx(1):endQuote);

% Look for the comment.
slashIdx = find(cardImage == '/');

if (~isempty(slashIdx))

    % The first slash beyond the value's end quote starts the comment.
    commentStartIdx = min(slashIdx(slashIdx > endQuote));

    if (~isempty(commentStartIdx))

        comment = cardImage(commentStartIdx:end);

    else

        comment = '';

    end

else

    comment = '';

end


function [value, comment] = splitNumericCard(cardImage)

% It's possible to have slashes in keywords.  Start just after keyword.
slashIdx = find(cardImage(9:end) == '/') + 8;

if (~isempty(slashIdx))

    commentStartIdx = min(slashIdx);

    value   = cardImage(11:(commentStartIdx - 1));
    comment = cardImage(commentStartIdx:end);

else

    value   = cardImage(11:end);
    comment = '';

end


function comment = stripCommentMarker(comment)

if (isempty(comment))

    return

elseif (isequal(comment(1), '/'))

    comment(1) = '';

end


function value = cleanupNumericValue(value)

if (isempty(value))

    return

else

    % Remove blanks from other formats.
    value = sscanf(value, ' %s');

end


function value = cleanupCharacterValue(value)

if (isempty(value))

    return

else

    % Remove opening and closing single quote from character values.
    if (numel(value) >= 2)
        value([1 end]) = '';
    end

    % Remove trailing blanks.
    value = deblank(value);

    % Convert paired single quotes to one single quote.
    quoteChar = char(39);  % 39 == '
    value = strrep(value, [quoteChar quoteChar], quoteChar);

end


function valueNum = convertNumericValue(valueStr)

if (isempty(valueStr))

    valueNum = [];
    return

elseif (strfind(valueStr, '('))

    % Complex values.
    numberParts = sscanf(valueStr, ' ( %f , %f ) ');

    if (numel(numberParts) == 2)

        valueNum = complex(numberParts(1), numberParts(2));

    else

        valueNum = [];

    end

else

    switch (valueStr)
        case {'T', 'F'}

            valueNum = valueStr;

        otherwise

            valueNum = sscanf(valueStr, '%f');

            if (isempty(valueNum))

                warning('MATLAB:fitsinfo:unknownFormat', ...
                    ['Unable to determine format of the card value.\n', ...
                    'Returning the value as a string.'])

                valueNum = valueStr;

            end


    end

end


function tf = isCharacterString(cardImage)

% A value is a character string if the value begins with a quote.

valueAndComment = strtrim(cardImage(11:end));

if (isempty(valueAndComment))

    tf = false;

else

    quoteChar = char(39);  % 39 == '
    tf = isequal(valueAndComment(1), quoteChar);

end


function cardHasTIMFILE = isTIMFILE(cardImage)

cardHasTIMFILE = isequal(cardImage(1:7), 'TIMFILE');


function [value, comment] = splitTIMFILECard(cardImage)

% This keyword appears to contain a path string erroneously written without
% quotes.

keywordLength = 9;
valueAndComment = cardImage((keywordLength + 1):end);

slashIdx = find(valueAndComment == '/');

if (isempty(slashIdx))

    value = valueAndComment;
    comment = '';
    return

end

% This keyword must have a value, so the first slash cannot start a comment
% and be the first nonspace element of the card after the keyword.  It
% suffices to find the first " /" after the value and use that as the
% comment.

% Find the whitespace boundaries around the value.
spaceIdx = find(isspace(valueAndComment));
letterIdx = find(~isspace(valueAndComment));

valueStart = min(letterIdx);
valueEnd = min(spaceIdx(spaceIdx > valueStart)) - 1;

value = valueAndComment(valueStart:valueEnd);

% Find the optional comment.
commentStart = min(slashIdx(slashIdx > valueEnd));
if (isempty(commentStart))

    comment = '';

else

    comment = valueAndComment(commentStart:end);

end


function commentOut = cleanupCommentWhitespace(commentIn)

% The standard doesn't say how to treat whitespace before and after
% comments, so we will leave it.  The one exception is if the entire
% comment is empty.  This will likely only happen if there wasn't a value
% or comment in the card (i.e., everything after the keyword was empty).

if (all(commentIn == ' '))  % Compare to space.

    commentOut = '';

else

    commentOut = commentIn;

end


function status = moveToNextHDU(fid)

curr_pos = ftell(fid);

if rem(curr_pos, 2880) == 0
    bytes_to_seek = 0;
else
    bytes_to_seek = 2880 - rem(curr_pos, 2880);
end

status = fseek(fid, bytes_to_seek, 'cof');


function cardHasCOMMENT = isCOMMENT(cardImage)

cardHasCOMMENT = isequal(cardImage(1:7), 'COMMENT');
```
