## Supplementary material for "Microvillar cartography: a super-resolution single-molecule imaging method to map the positions of membrane proteins with respect to cellular surface topography": Analysis MATLAB codes: fits_shifter_single_Y_v30.m

```
function [filesData, frames_out] = fits_shifter_single_Y_v30(PathName,FileName,FrameN)
% Read Frame-Nth data 
%FrameN=Frame Nth   
    fid_in=[PathName,FileName];
    filesData = fid_in;
    frames_out = fits2frames_Y(fid_in, FrameN , 'Max', []); %frames_out=spe2frames(fid, frames2take, method, cut), eg. 1=first frame
    frames_out = double (frames_out); % change single to double
end
```
