## Supplementary material for "Microvillar cartography: a super-resolution single-molecule imaging method to map the positions of membrane proteins with respect to cellular surface topography": Analysis MATLAB codes: FitsetRoi_range.m

```
function[RoifitsetPT]=FitsetRoi_range(data,range_x,range_y)
%test
%data=FitsetPT;
% xcnt=135;
% ycnt=135;
% roiws=101;

B=[];
%pn=(roiws-1)/2;

ct=0;
A=data;
% if size(A,2)~=2
%     error ('use FitsetRoi.m')
% end
% xmin=xcnt-pn;
% xmax=xcnt+pn;
% ymin=ycnt-pn;
% ymax=ycnt+pn;
xmin=range_x(1);
xmax=range_x(2);
ymin=range_y(1);
ymax=range_y(2);


for k=1:size(data,1)
    if A(k,3)>xmin && A(k,3)<xmax && A(k,2)>ymin && A(k,2)<ymax
        ct=ct+1;
       B(ct,:)=A(k,:);
    end
end
RoifitsetPT=B;

% for k=1:size(data,1)
%     if A(k,2)>ymin && A(k,2)<ymax && A(k,3)>xmin && A(k,3)<xmax
%         ct=ct+1;
%        B(ct,:)=A(k,:);
%     end
% end
%RoifitsetPT=B;

% test figure
% figure
% subplot(1,2,1)
% scatter(A(:,3),A(:,2),'.r');
% axis square
% subplot(1,2,2)
% scatter(B(:,3),B(:,2),'.b');
% axis square


end
```
