## Supplementary material for "Microvillar cartography: a super-resolution single-molecule imaging method to map the positions of membrane proteins with respect to cellular surface topography": Analysis MATLAB codes: func_consentric_shape_formation_area.m

```
%%find concentric shapes, total area of previous and new shapes, area
%%difference between two shapes; Created by Shirsendu on 22nd August,2017 
function[finalpt1,area_previous,area_final,area_diff]=func_consentric_shape_formation_area(A,d)
%call2=1
%% d=distane between ages in nm; %%find no of edges
%A=[164.683418207005,146.350082495875;167.333335711130,149;162.683418207005,153.649917504125;159.350082495875,153.649917504125;159.350082495875,146.316581792995;163.649917504125,142.016746784745;163.649917504125,146.316581792995;163.683418207005,146.350082495875;164.683418207005,146.350082495875];
%d=10;
%for sgg=1:40
noofedges=(size(A,1)-1);
%plot(A(:,1),A(:,2))
%hold on
%%find coeffecients of the straightline of edges of edges
coeffecient_edges=zeros(noofedges,2);
middle_point=zeros(noofedges,2);
finalpt=zeros(1,2);
final_check_cutting_point=zeros(1,2);
final_right_parallel_line_coeffecient_edges=zeros(1,2);
parallel_line_coeffecient_edges=zeros(noofedges,4);
perpendicular_line_coeffecient_edges=zeros(noofedges,2);
right_parallel_line_coeffecient_edges=zeros(noofedges,2);
cutting_point=zeros(noofedges,4);

for aa=1:noofedges
    coeffecient_edges(aa,1)= ((A(aa,2)-A(aa+1,2))/(A(aa,1)-A(aa+1,1)));
    if ((abs(coeffecient_edges(aa,1))==Inf))
        coeffecient_edges(aa,1)=Inf;
    end
    if(coeffecient_edges(aa,1)~=Inf)
    coeffecient_edges(aa,2)=A(aa,2)-(coeffecient_edges(aa,1)*A(aa,1));
    else
      coeffecient_edges(aa,2)=A(aa,1);  
    end
end
%%find coeffecients of the parallel line of edges
for aa=1:noofedges
   if(coeffecient_edges(aa,1)~=inf)
   dist=(d/66.67)*sqrt((coeffecient_edges(aa,1)*coeffecient_edges(aa,1))+1);
   parallel_line_coeffecient_edges(aa,:)= [coeffecient_edges(aa,1),(coeffecient_edges(aa,2)-(dist)),coeffecient_edges(aa,1),(coeffecient_edges(aa,2)+(dist))];
   else
   parallel_line_coeffecient_edges(aa,:)=[inf,(coeffecient_edges(aa,2)-(d/66.67)),inf,(coeffecient_edges(aa,2)+(d/66.67))];     
   end
end
%%% find the middle point of edges
for aa=1:noofedges
middle_point(aa,:)=[((A(aa,1)+A(aa+1,1))/2) ((A(aa,2)+A(aa+1,2))/2)];
%scatter(middle_point(aa,1),middle_point(aa,2))
%hold on
end

%%%find coefficient of perpendicular lines passing through the middle point of
%%%edges
for aa=1:noofedges
 if(coeffecient_edges(aa,1)~=0)&&(coeffecient_edges(aa,1)~=inf)  
perpendicular_line_coeffecient_edges(aa,1)=-1/coeffecient_edges(aa,1);
perpendicular_line_coeffecient_edges(aa,2)=middle_point(aa,2)-(perpendicular_line_coeffecient_edges(aa,1)*middle_point(aa,1));
 end
 if(coeffecient_edges(aa,1)==0)
     perpendicular_line_coeffecient_edges(aa,1)=inf;
     perpendicular_line_coeffecient_edges(aa,2)=middle_point(aa,1);
 end
 if(coeffecient_edges(aa,1)==inf)
     perpendicular_line_coeffecient_edges(aa,1)=0;
     perpendicular_line_coeffecient_edges(aa,2)=middle_point(aa,2);
 end
 
%   u=0:.1:256;
%   v=(perpendicular_line_coeffecient_edges(aa,1)*u)+perpendicular_line_coeffecient_edges(aa,2);
   %plot(u,v)
%   hold on
end
%for aa=1:noofedges
% if(parallel_line_coeffecient_edges(aa,1)~=inf)   
%  u=-5:1:5;
%   v=(parallel_line_coeffecient_edges(aa,1)*u)+parallel_line_coeffecient_edges(aa,2);
%   plot(u,v)
%   hold on
%  u=-5:1:5;
%   v=(parallel_line_coeffecient_edges(aa,3)*u)+parallel_line_coeffecient_edges(aa,4);
%   plot(u,v)
%   hold on
% else
%    v=-5:1:5;
%    v=v';
%    vsize=size(v,1);
%    u=zeros(vsize,1);
%    u(:,1)=parallel_line_coeffecient_edges(aa,2);
%    plot(u,v)
%   hold on
%    u=zeros(vsize,1);
%    u(:,1)=parallel_line_coeffecient_edges(aa,4);
%   plot(u,v)
%   hold on
% end
% end
%%% find the cutting point of perpendicular lines and parallel lines and
%%% right parallel
for aa=1:noofedges
    if(perpendicular_line_coeffecient_edges(aa,1)~=inf)&&(abs(perpendicular_line_coeffecient_edges(aa,1))>=10000)
     perpendicular_line_coeffecient_edges(aa,1)=inf;
     perpendicular_line_coeffecient_edges(aa,2)=middle_point(aa,1);
    end
if(perpendicular_line_coeffecient_edges(aa,1)~=0)&&(perpendicular_line_coeffecient_edges(aa,1)~=inf)
cutting_point(aa,1)=[(perpendicular_line_coeffecient_edges(aa,2)-parallel_line_coeffecient_edges(aa,2))/(parallel_line_coeffecient_edges(aa,1)-perpendicular_line_coeffecient_edges(aa,1))];
cutting_point(aa,2)=[(cutting_point(aa,1)*perpendicular_line_coeffecient_edges(aa,1))+perpendicular_line_coeffecient_edges(aa,2)];
cutting_point(aa,3)=[(perpendicular_line_coeffecient_edges(aa,2)-parallel_line_coeffecient_edges(aa,4))/(parallel_line_coeffecient_edges(aa,3)-perpendicular_line_coeffecient_edges(aa,1))];
cutting_point(aa,4)=[(cutting_point(aa,3)*perpendicular_line_coeffecient_edges(aa,1))+perpendicular_line_coeffecient_edges(aa,2)];
end
if(perpendicular_line_coeffecient_edges(aa,1)==0)
   cutting_point(aa,1)= parallel_line_coeffecient_edges(aa,2);
   cutting_point(aa,2)= perpendicular_line_coeffecient_edges(aa,2);
   cutting_point(aa,3)= parallel_line_coeffecient_edges(aa,4);
   cutting_point(aa,4)= perpendicular_line_coeffecient_edges(aa,2);
end
if(perpendicular_line_coeffecient_edges(aa,1)==Inf)
   cutting_point(aa,1)= perpendicular_line_coeffecient_edges(aa,2);
   cutting_point(aa,2)= parallel_line_coeffecient_edges(aa,2);
   cutting_point(aa,3)= perpendicular_line_coeffecient_edges(aa,2);
   cutting_point(aa,4)= parallel_line_coeffecient_edges(aa,4);
end
%scatter(cutting_point(aa,1),cutting_point(aa,2))
%hold on
%scatter(cutting_point(aa,3),cutting_point(aa,4))
%hold on
if(cutting_point(aa,1)<middle_point(aa,1))
xup=cutting_point(aa,1):0.0001:middle_point(aa,1);
yup=[(xup*perpendicular_line_coeffecient_edges(aa,1))+perpendicular_line_coeffecient_edges(aa,2)];
end
if(cutting_point(aa,1)>middle_point(aa,1))
xup=middle_point(aa,1):0.0001:cutting_point(aa,1);
yup=[(xup*perpendicular_line_coeffecient_edges(aa,1))+perpendicular_line_coeffecient_edges(aa,2)];
end
if(cutting_point(aa,1)==middle_point(aa,1))
    if(cutting_point(aa,2)>middle_point(aa,2))
    yup=middle_point(aa,2):0.0001:cutting_point(aa,2);
    yup=yup';
    checkxup=size(yup,1);
    xup=zeros(checkxup,1);
    xup(:,1)=cutting_point(aa,1);
    end
    if(cutting_point(aa,2)<middle_point(aa,2))
    yup=cutting_point(aa,2):0.0001:middle_point(aa,2);
    yup=yup';
    checkxup=size(yup,1);
    xup=zeros(checkxup,1);
    xup(:,1)=cutting_point(aa,1);
    end
end
%plot(xup,yup);
%hold on
if(cutting_point(aa,3)<middle_point(aa,1))
xdown=cutting_point(aa,3):.0001:middle_point(aa,1);
ydown=[(xdown*perpendicular_line_coeffecient_edges(aa,1))+perpendicular_line_coeffecient_edges(aa,2)];
end
if(cutting_point(aa,3)>middle_point(aa,1))
xdown=middle_point(aa,1):.0001:cutting_point(aa,3);
ydown=[(xdown*perpendicular_line_coeffecient_edges(aa,1))+perpendicular_line_coeffecient_edges(aa,2)];
end
if(cutting_point(aa,3)==middle_point(aa,1))
    if(cutting_point(aa,4)>middle_point(aa,2))
    ydown=middle_point(aa,2):0.0001:cutting_point(aa,4);
    ydown=ydown';
    checkxdown=size(ydown,1);
    xdown=zeros(checkxdown,1);
    xdown(:,1)=cutting_point(aa,3);
    end
    if(cutting_point(aa,4)<middle_point(aa,2))
    ydown=cutting_point(aa,4):0.0001:middle_point(aa,2);
    ydown=ydown';
    checkxdown=size(ydown,1);
    xdown=zeros(checkxup,1);
    xdown(:,1)=cutting_point(aa,4);
    end
end
%plot(xdown,ydown)
%hold on
[inup,onup] = inpolygon(xup,yup,A(:,1),A(:,2));
[indown,ondown] = inpolygon(xdown,ydown,A(:,1),A(:,2));
zup=sum(inup);
zdown=sum(indown);
if zup<zdown
    right_parallel_line_coeffecient_edges(aa,1)=parallel_line_coeffecient_edges(aa,1);
    right_parallel_line_coeffecient_edges(aa,2)=parallel_line_coeffecient_edges(aa,2);
end
if zup>zdown
    right_parallel_line_coeffecient_edges(aa,1)=parallel_line_coeffecient_edges(aa,3);
    right_parallel_line_coeffecient_edges(aa,2)=parallel_line_coeffecient_edges(aa,4);
end
end
%scatter(middle_point(:,1),middle_point(:,2))
%hold on
%scatter(cutting_point(:,1),cutting_point(:,2))
%hold on
%scatter(cutting_point(:,3),cutting_point(:,4))
%hold on
%plotrange_1=min(A);
%plotrange_2=max(A);
%plot_xrange=[plotrange_1(1) plotrange_2(1)];
%plot_yrange=[plotrange_1(2) plotrange_2(2)];
%for aa=1:noofedges
%   if(right_parallel_line_coeffecient_edges(aa,1)~=inf) 
%    u=plot_xrange(1):.1:plot_xrange(2);
%    v=(right_parallel_line_coeffecient_edges(aa,1)*u)+right_parallel_line_coeffecient_edges(aa,2);
%    plot(u,v)
%    hold on
%    else
%    v=plot_yrange(1):.1:plot_yrange(2);
%    v=v';
%    u=zeros((size(v,1)),1);
%    u(:,1)=right_parallel_line_coeffecient_edges(aa,2);
%    plot(u,v)
%    hold on
%    end
%end
%    u=-3:.1:3;
%   v=(parallel_line_coeffecient_edges(aa,1)*u)+parallel_line_coeffecient_edges(aa,2);
%   plot(u,v)
%   hold on
%  u=-3:.1:3;
%   v=(parallel_line_coeffecient_edges(aa,3)*u)+parallel_line_coeffecient_edges(aa,4);
%   plot(u,v)
%   hold on
%    end
%    end
final_point_count=0;
for aa=1:noofedges
    if aa<noofedges
      if (right_parallel_line_coeffecient_edges(aa,1)~=right_parallel_line_coeffecient_edges((aa+1),1))&&(right_parallel_line_coeffecient_edges(aa,2)~=right_parallel_line_coeffecient_edges((aa+1),2))
      final_point_count=final_point_count+1;
      final_right_parallel_line_coeffecient_edges(final_point_count,1)=right_parallel_line_coeffecient_edges(aa,1);
      final_right_parallel_line_coeffecient_edges(final_point_count,2)=right_parallel_line_coeffecient_edges(aa,2);
      if(right_parallel_line_coeffecient_edges(aa,1)~=inf)&&(right_parallel_line_coeffecient_edges((aa+1),1)~=inf)   
      finalpt(final_point_count,1)=[(right_parallel_line_coeffecient_edges(aa+1,2)-right_parallel_line_coeffecient_edges(aa,2))/(right_parallel_line_coeffecient_edges(aa,1)-right_parallel_line_coeffecient_edges((aa+1),1))];
      finalpt(final_point_count,2)=[(finalpt(final_point_count,1)*right_parallel_line_coeffecient_edges(aa,1))+right_parallel_line_coeffecient_edges(aa,2)];
      end
      if(right_parallel_line_coeffecient_edges(aa,1)==inf)&&(right_parallel_line_coeffecient_edges((aa+1),1)~=inf)   
      finalpt(final_point_count,1)=right_parallel_line_coeffecient_edges(aa,2);
      finalpt(final_point_count,2)=[(finalpt(final_point_count,1)*right_parallel_line_coeffecient_edges((aa+1),1))+right_parallel_line_coeffecient_edges((aa+1),2)];
      end
      if(right_parallel_line_coeffecient_edges(aa,1)~=inf)&&(right_parallel_line_coeffecient_edges((aa+1),1)==inf)   
      finalpt(final_point_count,1)=right_parallel_line_coeffecient_edges((aa+1),2);
      finalpt(final_point_count,2)=[(finalpt(final_point_count,1)*right_parallel_line_coeffecient_edges(aa,1))+right_parallel_line_coeffecient_edges(aa,2)];
      end
      end
    end
    if aa==noofedges
      if (right_parallel_line_coeffecient_edges(aa,1)~=right_parallel_line_coeffecient_edges(1,1))&&(right_parallel_line_coeffecient_edges(aa,2)~=right_parallel_line_coeffecient_edges(1,2))
      final_point_count=final_point_count+1;
      final_right_parallel_line_coeffecient_edges(final_point_count,1)=right_parallel_line_coeffecient_edges(aa,1);
      final_right_parallel_line_coeffecient_edges(final_point_count,2)=right_parallel_line_coeffecient_edges(aa,2);
      if(right_parallel_line_coeffecient_edges(aa,1)~=inf)&&(right_parallel_line_coeffecient_edges(1,1)~=inf) 
      finalpt(final_point_count,1)=[(right_parallel_line_coeffecient_edges(1,2)-right_parallel_line_coeffecient_edges(aa,2))/(right_parallel_line_coeffecient_edges(aa,1)-right_parallel_line_coeffecient_edges(1,1))];
      finalpt(final_point_count,2)=[(finalpt(final_point_count,1)*right_parallel_line_coeffecient_edges(aa,1))+right_parallel_line_coeffecient_edges(aa,2)];
      end
      if(right_parallel_line_coeffecient_edges(aa,1)==inf)&&(right_parallel_line_coeffecient_edges(1,1)~=inf)   
      finalpt(final_point_count,1)=right_parallel_line_coeffecient_edges(aa,2);
      finalpt(final_point_count,2)=[(finalpt(final_point_count,1)*right_parallel_line_coeffecient_edges(1,1))+right_parallel_line_coeffecient_edges(1,2)];
      end
      if(right_parallel_line_coeffecient_edges(aa,1)~=inf)&&(right_parallel_line_coeffecient_edges(1,1)==inf)   
      finalpt(final_point_count,1)=right_parallel_line_coeffecient_edges(1,2);
      finalpt(final_point_count,2)=[(finalpt(final_point_count,1)*right_parallel_line_coeffecient_edges(aa,1))+right_parallel_line_coeffecient_edges(aa,2)];
      end
      end
    end
end
finalpt((final_point_count+1),:)=finalpt(1,:);
%plot(finalpt(:,1),finalpt(:,2));
%hold on
%scatter(finalpt(:,1),finalpt(:,2));
hold on
finalpt1=finalpt;
finalpt_size=size(finalpt,1);
finalpt1(finalpt_size,:)=[];
if(finalpt_size>4)
%%checking for crossing
cc_check=0;
bb=1;
while bb<(final_point_count+1)
    cc_check=cc_check+1;
    if(bb<(final_point_count-1))
if (final_right_parallel_line_coeffecient_edges(bb,1)~=final_right_parallel_line_coeffecient_edges((bb+2),1))
    if(final_right_parallel_line_coeffecient_edges((bb+2),1)~=Inf)&&(final_right_parallel_line_coeffecient_edges(bb,1)~=Inf)
    final_check_cutting_point(bb,1)=[(final_right_parallel_line_coeffecient_edges((bb+2),2)-final_right_parallel_line_coeffecient_edges(bb,2))/(final_right_parallel_line_coeffecient_edges(bb,1)-final_right_parallel_line_coeffecient_edges((bb+2),1))];
    final_check_cutting_point(bb,2)=(final_check_cutting_point(bb,1)*final_right_parallel_line_coeffecient_edges(bb,1))+final_right_parallel_line_coeffecient_edges(bb,2);
    end
    if(final_right_parallel_line_coeffecient_edges((bb+2),1)==Inf)
    final_check_cutting_point(bb,1)=final_right_parallel_line_coeffecient_edges((bb+2),2);
    final_check_cutting_point(bb,2)=(final_check_cutting_point(bb,1)*final_right_parallel_line_coeffecient_edges(bb,1))+final_right_parallel_line_coeffecient_edges(bb,2);
    end
    if(final_right_parallel_line_coeffecient_edges(bb,1)==Inf)
    final_check_cutting_point(bb,1)=final_right_parallel_line_coeffecient_edges(bb,2);
    final_check_cutting_point(bb,2)=(final_check_cutting_point(bb,1)*final_right_parallel_line_coeffecient_edges((bb+2),1))+final_right_parallel_line_coeffecient_edges((bb+2),2);
    end
    [in,on] = inpolygon(final_check_cutting_point(bb,1),final_check_cutting_point(bb,2),finalpt(:,1),finalpt(:,2));
    if (on==1)
        %SSG=1
        %cc_check
        %bb
       if(cc_check>1)
        x1=[finalpt1((cc_check-1),1) finalpt1((cc_check-1),2);finalpt1((cc_check),1) finalpt1((cc_check),2)];
       else
        x1=[finalpt1((final_point_count),1) finalpt1((final_point_count),2);finalpt1((cc_check),1) finalpt1((cc_check),2)];   
       end
        if(cc_check<=final_point_count-2)
            y1=[finalpt1((cc_check+1),1) finalpt1((cc_check+1),2);finalpt1((cc_check+2),1) finalpt1((cc_check+2),2)];
        end
        if(cc_check==final_point_count-1)
            y1=[finalpt1((cc_check+1),1) finalpt1((cc_check+1),2);finalpt1(1,1) finalpt1(1,2)];
        end
        if(cc_check==final_point_count)
            y1=[finalpt1(1,1) finalpt1(1,2);finalpt1(2,1) finalpt1(2,2)];
        end
        z1=[finalpt1(cc_check,1) finalpt1(cc_check,2);final_check_cutting_point(bb,1) final_check_cutting_point(bb,2)];
        if(cc_check>1)
            z2=[finalpt1((cc_check-1),1) finalpt1((cc_check-1),2);final_check_cutting_point(bb,1) final_check_cutting_point(bb,2)];
        else
            z2=[finalpt1((final_point_count),1) finalpt1((final_point_count),2);final_check_cutting_point(bb,1) final_check_cutting_point(bb,2)];
        end
        if(cc_check<final_point_count-1)
        z3=[finalpt1((cc_check+2),1) finalpt1((cc_check+2),2);final_check_cutting_point(bb,1) final_check_cutting_point(bb,2)];
        end
        if(cc_check==final_point_count-1)
        z3=[finalpt1(1,1) finalpt1(1,2);final_check_cutting_point(bb,1) final_check_cutting_point(bb,2)];
        end
        if(cc_check==final_point_count)
        z3=[finalpt1(2,1) finalpt1(2,2);final_check_cutting_point(bb,1) final_check_cutting_point(bb,2)];
        end
        if(cc_check<final_point_count)
            z4=[finalpt1((cc_check+1),1) finalpt1((cc_check+1),2);final_check_cutting_point(bb,1) final_check_cutting_point(bb,2)];
        else
            z4=[finalpt1(1,1) finalpt1(1,2);final_check_cutting_point(bb,1) final_check_cutting_point(bb,2)];
        end
        dist_x1=pdist(x1,'euclidean');
        dist_y1=pdist(y1,'euclidean');
        dist_z1=pdist(z1,'euclidean');
        dist_z2=pdist(z2,'euclidean');
        dist_z3=pdist(z3,'euclidean');
        dist_z4=pdist(z4,'euclidean');
        if (dist_x1>=dist_z1)&&(dist_x1>=dist_z2)&&(dist_y1>=dist_z3)&&(dist_y1>=dist_z4)
            %SSG=2
       finalpt1(cc_check,:)=[];
       finalpt1(cc_check,:)=[final_check_cutting_point(bb,1) final_check_cutting_point(bb,2)];
       bb=bb+1;
       final_point_count=final_point_count-1;
        end
    end
end
    end
    if(bb==(final_point_count-1))
if (final_right_parallel_line_coeffecient_edges(bb,1)~=final_right_parallel_line_coeffecient_edges(1,1))
   if(final_right_parallel_line_coeffecient_edges(1,1)~=Inf)&&(final_right_parallel_line_coeffecient_edges(bb,1)~=Inf)
    final_check_cutting_point(bb,1)=[(final_right_parallel_line_coeffecient_edges(1,2)-final_right_parallel_line_coeffecient_edges(bb,2))/(final_right_parallel_line_coeffecient_edges(bb,1)-final_right_parallel_line_coeffecient_edges(1,1))];
    final_check_cutting_point(bb,2)=(final_check_cutting_point(bb,1)*final_right_parallel_line_coeffecient_edges(bb,1))+final_right_parallel_line_coeffecient_edges(bb,2);
    [in,on] = inpolygon(final_check_cutting_point(bb,1),final_check_cutting_point(bb,2),finalpt(:,1),finalpt(:,2));
   end
    if(final_right_parallel_line_coeffecient_edges(1,1)==Inf)
    final_check_cutting_point(bb,1)=final_right_parallel_line_coeffecient_edges(1,2);
    final_check_cutting_point(bb,2)=(final_check_cutting_point(bb,1)*final_right_parallel_line_coeffecient_edges(bb,1))+final_right_parallel_line_coeffecient_edges(bb,2);
    end
    if(final_right_parallel_line_coeffecient_edges(bb,1)==Inf)
    final_check_cutting_point(bb,1)=final_right_parallel_line_coeffecient_edges(bb,2);
    final_check_cutting_point(bb,2)=(final_check_cutting_point(bb,1)*final_right_parallel_line_coeffecient_edges(1,1))+final_right_parallel_line_coeffecient_edges(1,2);
    end
    if (on==1)
        %SSG=3
        if(cc_check>1)
        x1=[finalpt1((cc_check-1),1) finalpt1((cc_check-1),2);finalpt1((cc_check),1) finalpt1((cc_check),2)];
       else
        x1=[finalpt1(final_point_count,1) finalpt1(final_point_count,2);finalpt1((cc_check),1) finalpt1((cc_check),2)];   
       end
        if(cc_check<=final_point_count-2)
            y1=[finalpt1((cc_check+1),1) finalpt1((cc_check+1),2);finalpt1((cc_check+2),1) finalpt1((cc_check+2),2)];
        end
        if(cc_check==final_point_count-1)
            y1=[finalpt1((cc_check+1),1) finalpt1((cc_check+1),2);finalpt1(1,1) finalpt1(1,2)];
        end
        if(cc_check==final_point_count)
            y1=[finalpt1(1,1) finalpt1(1,2);finalpt1(2,1) finalpt1(2,2)];
        end
        z1=[finalpt1(cc_check,1) finalpt1(cc_check,2);final_check_cutting_point(bb,1) final_check_cutting_point(bb,2)];
        if(cc_check>1)
            z2=[finalpt1((cc_check-1),1) finalpt1((cc_check-1),2);final_check_cutting_point(bb,1) final_check_cutting_point(bb,2)];
        else
            z2=[finalpt1((final_point_count),1) finalpt1((final_point_count),2);final_check_cutting_point(bb,1) final_check_cutting_point(bb,2)];
        end
        if(cc_check<final_point_count-1)
        z3=[finalpt1((cc_check+2),1) finalpt1((cc_check+2),2);final_check_cutting_point(bb,1) final_check_cutting_point(bb,2)];
        end
        if(cc_check==final_point_count-1)
        z3=[finalpt1(1,1) finalpt1(1,2);final_check_cutting_point(bb,1) final_check_cutting_point(bb,2)];
        end
        if(cc_check==final_point_count)
        z3=[finalpt1(2,1) finalpt1(2,2);final_check_cutting_point(bb,1) final_check_cutting_point(bb,2)];
        end
        if(cc_check<final_point_count)
            z4=[finalpt1((cc_check+1),1) finalpt1((cc_check+1),2);final_check_cutting_point(bb,1) final_check_cutting_point(bb,2)];
        else
            z4=[finalpt1(1,1) finalpt1(1,2);final_check_cutting_point(bb,1) final_check_cutting_point(bb,2)];
        end
        dist_x1=pdist(x1,'euclidean');
        dist_y1=pdist(y1,'euclidean');
        dist_z1=pdist(z1,'euclidean');
        dist_z2=pdist(z2,'euclidean');
        dist_z3=pdist(z3,'euclidean');
        dist_z4=pdist(z4,'euclidean');
        if (dist_x1>=dist_z1)&&(dist_x1>=dist_z2)&&(dist_y1>=dist_z3)&&(dist_y1>=dist_z4)
         %   SSG=4
       finalpt1(cc_check,:)=[];
       finalpt1(cc_check,:)=[final_check_cutting_point(bb,1) final_check_cutting_point(bb,2)];
       bb=bb+1;
       final_point_count=final_point_count-1;
       end
    end
end
    end
    if(bb==final_point_count)
if (final_right_parallel_line_coeffecient_edges(bb,1)~=final_right_parallel_line_coeffecient_edges(2,1))
    final_check_cutting_point(bb,1)=[(final_right_parallel_line_coeffecient_edges(2,2)-final_right_parallel_line_coeffecient_edges(bb,2))/(final_right_parallel_line_coeffecient_edges(bb,1)-final_right_parallel_line_coeffecient_edges(2,1))];
    final_check_cutting_point(bb,2)=(final_check_cutting_point(bb,1)*final_right_parallel_line_coeffecient_edges(bb,1))+final_right_parallel_line_coeffecient_edges(bb,2);
    [in,on] = inpolygon(final_check_cutting_point(bb,1),final_check_cutting_point(bb,2),finalpt(:,1),finalpt(:,2));
    if (on==1)
        %SSG=5
        if(cc_check>1)
        x1=[finalpt1((cc_check-1),1) finalpt1((cc_check-1),2);finalpt1((cc_check),1) finalpt1((cc_check),2)];
       else
        x1=[finalpt1(final_point_count,1) finalpt1(final_point_count,2);finalpt1((cc_check),1) finalpt1((cc_check),2)];   
       end
        if(cc_check<=final_point_count-2)
            y1=[finalpt1((cc_check+1),1) finalpt1((cc_check+1),2);finalpt1((cc_check+2),1) finalpt1((cc_check+2),2)];
        end
        if(cc_check==final_point_count-1)
            y1=[finalpt1((cc_check+1),1) finalpt1((cc_check+1),2);finalpt1(1,1) finalpt1(1,2)];
        end
        if(cc_check==final_point_count)
            y1=[finalpt1(1,1) finalpt1(1,2);finalpt1(2,1) finalpt1(2,2)];
        end
        z1=[finalpt1(cc_check,1) finalpt1(cc_check,2);final_check_cutting_point(bb,1) final_check_cutting_point(bb,2)];
        if(cc_check>1)
            z2=[finalpt1((cc_check-1),1) finalpt1((cc_check-1),2);final_check_cutting_point(bb,1) final_check_cutting_point(bb,2)];
        else
            z2=[finalpt1((final_point_count),1) finalpt1((final_point_count),2);final_check_cutting_point(bb,1) final_check_cutting_point(bb,2)];
        end
        if(cc_check<final_point_count-1)
        z3=[finalpt1((cc_check+2),1) finalpt1((cc_check+2),2);final_check_cutting_point(bb,1) final_check_cutting_point(bb,2)];
        end
        if(cc_check==final_point_count-1)
        z3=[finalpt1(1,1) finalpt1(1,2);final_check_cutting_point(bb,1) final_check_cutting_point(bb,2)];
        end
        if(cc_check==final_point_count)
        z3=[finalpt1(2,1) finalpt1(2,2);final_check_cutting_point(bb,1) final_check_cutting_point(bb,2)];
        end
        if(cc_check<final_point_count)
            z4=[finalpt1((cc_check+1),1) finalpt1((cc_check+1),2);final_check_cutting_point(bb,1) final_check_cutting_point(bb,2)];
        else
            z4=[finalpt1(1,1) finalpt1(1,2);final_check_cutting_point(bb,1) final_check_cutting_point(bb,2)];
        end
        dist_x1=pdist(x1,'euclidean');
        dist_y1=pdist(y1,'euclidean');
        dist_z1=pdist(z1,'euclidean');
        dist_z2=pdist(z2,'euclidean');
        dist_z3=pdist(z3,'euclidean');
        dist_z4=pdist(z4,'euclidean');
        if (dist_x1>=dist_z1)&&(dist_x1>=dist_z2)&&(dist_y1>=dist_z3)&&(dist_y1>=dist_z4)
         %   SSG=6
       finalpt1(cc_check,:)=[final_check_cutting_point(bb,1) final_check_cutting_point(bb,2)];
       finalpt1(1,:)=[];
        end
    end
end
    end
    bb=bb+1;
end
end
finalpt1_size=size(finalpt1,1)+1;
finalpt1(finalpt1_size,:)=finalpt1(1,:);
plot(finalpt1(:,1),finalpt1(:,2));
%hold on
%scatter(finalpt1(:,1),finalpt1(:,2));
%xlim([(plot_xrange(1)-5) (plot_xrange(2)+5)])
%ylim([(plot_yrange(1)-5) (plot_yrange(2)+5)])
area_previous=[(polyarea(A(:,1),A(:,2)))*(66.67^2)];
area_final=[(polyarea(finalpt(:,1),finalpt(:,2)))*(66.67^2)];
area_diff=area_final-area_previous;
%A=finalpt1;
%end
end
%%%%%%%
 
```
