## Supplementary material for "Microvillar cartography: a super-resolution single-molecule imaging method to map the positions of membrane proteins with respect to cellular surface topography": Analysis MATLAB codes: func_distance_density_central_area.m

```
%%find density vs distance plot of each microvilli by calculating the distance from central region of microvill; Created by Shirsendu on 27th August,2017 
function[count_area_0nm,count_area_400nm]=func_distance_density_central_area(A,d,range_x,range_y,RoiFitsetPT1,RoiFitsetPT2,filename,den_dist)
%[matfilename, pathname] = uigetfile( ...
%{  '*.mat','MAT-files (*.mat)'; ...
%   '*.*',  'All Files (*.*)'}, ...
%   'Pick a "VA_New_analysis_allinone file', ...
%   'MultiSelect', 'off');  

%cd(pathname);
%load(matfilename);
%[matfilename, pathname] = uigetfile( ...
%{  '*.mat','MAT-files (*.mat)'; ...
%   '*.*',  'All Files (*.*)'}, ...
%   'Pick a "VA_New_analysis_allinone file', ...
%   'MultiSelect', 'off'); 
%cd(pathname);
%load(matfilename);
%%inputing filename and microvill no etc.
getparts = strread(filename,'%s','delimiter','.');
getparts2= strread(getparts{1},'%s','delimiter','_');
getName  = getparts2(1);
getno    = getparts2(2);
microvillino=getno{1};
inpoint_0nm=zeros(size(RoiFitsetPT1,1),(den_dist+1));
inpoint_400nm=zeros(size(RoiFitsetPT2,1),(den_dist+1));
%d=10;
%count_area=zeros(23,3);
%% creating Count_area and finding the are of initial microvill shape 
count_area_0nm=zeros(den_dist,6);
count_area_400nm=zeros(den_dist,6);
%%find area and density of central region in 0 and 400nm
area_A=[(polyarea(A(:,1),A(:,2)))*(66.67^2)];
%scatter(RoiFitsetPT1(:,3),RoiFitsetPT1(:,2))
%hold on
%scatter(RoiFitsetPT2(:,3),RoiFitsetPT2(:,2))
[in_A_0nm,on_A_0nm] = inpolygon(RoiFitsetPT1(:,3),RoiFitsetPT1(:,2),A(:,2),A(:,1));
inpoint_0nm(:,1)=in_A_0nm;
points_A_0nm=sum(in_A_0nm);
count_area_0nm(1,1)=0;
count_area_0nm(1,2)=points_A_0nm;
count_area_0nm(1,3)=area_A;
count_area_0nm(1,4)=count_area_0nm(1,2)/count_area_0nm(1,3);
count_area_0nm(1,5)=area_A;
count_area_0nm(1,6)=points_A_0nm;
[in_A_400nm,on_A_400nm] = inpolygon(RoiFitsetPT2(:,3),RoiFitsetPT2(:,2),A(:,2),A(:,1));
inpoint_400nm(:,1)=in_A_400nm;
points_A_400nm=sum(in_A_400nm);
count_area_400nm(1,1)=0;
count_area_400nm(1,2)=points_A_400nm;
count_area_400nm(1,3)=area_A;
count_area_400nm(1,4)=count_area_400nm(1,2)/count_area_400nm(1,3);
count_area_400nm(1,5)=area_A;
count_area_400nm(1,6)=points_A_400nm;
B=zeros((size(A,1)),2);
B(:,1)=A(:,2);
B(:,2)=A(:,1);
plot(A(:,2),A(:,1))
xlim([range_x(1) range_x(2)])
ylim([range_y(1) range_y(2)])
hold on
%% finding concentric shapes 10 nm aparts and density changes with respect to distance
for i=2:den_dist
    %i
    [finalpt1,area_previous,area_final,area_diff]=func_consentric_shape_formation_area(B,d);
    area_B_diff=area_diff;
    [in_B_0nm,on_B_0nm] = inpolygon(RoiFitsetPT1(:,3),RoiFitsetPT1(:,2),B(:,1),B(:,2));
    [in_finalpt_0nm,on_finalpt_0nm] = inpolygon(RoiFitsetPT1(:,3),RoiFitsetPT1(:,2),finalpt1(:,1),finalpt1(:,2));
    inpoint_0nm(:,i)=in_finalpt_0nm;
    points_B_0nm_only=sum(in_finalpt_0nm)-sum(in_B_0nm);
    count_area_0nm(i,1)=((i-1)*10);
    count_area_0nm(i,2)=points_B_0nm_only;
    count_area_0nm(i,3)=area_B_diff;
    count_area_0nm(i,4)=count_area_0nm(i,2)/count_area_0nm(i,3);
    count_area_0nm(i,5)=area_final;
    count_area_0nm(i,6)=sum(in_finalpt_0nm);
    [in_B_400nm,on_B_400nm] = inpolygon(RoiFitsetPT2(:,3),RoiFitsetPT2(:,2),B(:,1),B(:,2));
    [in_finalpt_400nm,on_finalpt_400nm] = inpolygon(RoiFitsetPT2(:,3),RoiFitsetPT2(:,2),finalpt1(:,1),finalpt1(:,2));
    points_B_400nm_only=sum(in_finalpt_400nm)-sum(in_B_400nm);
    count_area_400nm(i,1)=((i-1)*10);
    count_area_400nm(i,2)=points_B_400nm_only;
    count_area_400nm(i,3)=area_B_diff;
    count_area_400nm(i,4)=count_area_400nm(i,2)/count_area_400nm(i,3);
    count_area_400nm(i,5)=area_final;
    count_area_400nm(i,6)=sum(in_finalpt_400nm);
    inpoint_400nm(:,i)=in_finalpt_400nm;
    plot(finalpt1(:,1),finalpt1(:,2))
    hold on
    B=finalpt1;
end
inpoint_0nm(:,(den_dist+1))=1;
inpoint_400nm(:,(den_dist+1))=1;
j=(den_dist+1);
savename=['Microvilli_' microvillino];
saveas(gcf,[savename])
saveas(gcf,[savename],'jpg')
hold on
while j>1
  inpoint_0nm(:,j)=inpoint_0nm(:,j)-inpoint_0nm(:,(j-1));  
  inpoint_400nm(:,j)=inpoint_400nm(:,j)-inpoint_400nm(:,(j-1));
  j=j-1;
end
xp_0nm=RoiFitsetPT1(:,3);
xp_400nm=RoiFitsetPT2(:,3);
yp_0nm=RoiFitsetPT1(:,2);
yp_400nm=RoiFitsetPT2(:,2);
k=den_dist;
while k>1
   l1=find(inpoint_0nm(:,k)>0);
   plot(xp_0nm(l1),yp_0nm(l1),'r+','markersize',.1) % points inside specific area
   hold on
   l2=find(inpoint_0nm(:,(k-1))>0);
   plot(xp_0nm(l2),yp_0nm(l2),'g.','markersize',.2) 
   hold on
    k=k-2;
end
hold on
l=find(inpoint_0nm(:,1)>0);
plot(xp_0nm(l),yp_0nm(l),'b*','markersize',.1) % points inside
hold on
l3=find(inpoint_0nm(:,(den_dist+1))>0);
plot(xp_0nm(l3),yp_0nm(l3),'k*','markersize',.1) % points outside
hold on
savename1=['Microvilli_' microvillino '_point_0nm'];
saveas(gcf,[savename1])
saveas(gcf,[savename1],'jpg')
hold off
openfig(savename)
k=den_dist;
while k>1
   ll1=find(inpoint_400nm(:,k)>0);
   plot(xp_400nm(ll1),yp_400nm(ll1),'r+','markersize',.1) % points inside specific area
   hold on
   ll2=find(inpoint_400nm(:,(k-1))>0);
   plot(xp_400nm(ll2),yp_400nm(ll2),'g.','markersize',.2) 
   hold on
    k=k-2;
end
hold on
ll=find(inpoint_400nm(:,1)>0);
plot(xp_400nm(ll),yp_400nm(ll),'b*','markersize',.1) % points inside
hold on
ll3=find(inpoint_400nm(:,(den_dist+1))>0);
plot(xp_400nm(ll3),yp_400nm(ll3),'k*','markersize',.1) % points outside
hold on
savename1=['Microvilli_' microvillino '_point_400nm'];
saveas(gcf,[savename1])
saveas(gcf,[savename1],'jpg')
hold off
%bar(density_area_0nm(:,1),density_area_0nm(:,2),'r')
%hold on
%bar(density_area_400nm(:,1),density_area_400nm(:,2),'g')
%hold off
%saveas(gcf,[savename1])
%saveas(gcf,[savename1],'jpg')
%count_area(:,1:2)=count_area_0nm;
%count_area(:,3)=count_area_400nm(:,2);
savename2=['Microvilli_' microvillino '_count_area_400nm'];
save(savename2,'count_area_400nm')
savename3=['Microvilli_' microvillino '_count_area_0nm'];
save(savename3,'count_area_0nm')
end
```
