## Supplementary material for "Microvillar cartography: a super-resolution single-molecule imaging method to map the positions of membrane proteins with respect to cellular surface topography": Analysis MATLAB codes: makeimagefile.m

```
function[h]=makeimagefile(FramesMat,colortype,savename)
if iscell(FramesMat)
    nframes =size(FramesMat,2);
 
elseif length(size(FramesMat))<3
    nframes=1;
   
elseif ~isempty(size(FramesMat),3)
     nframes=size(FramesMat,3);
end  
   
    for k=1:nframes
    
        
        if iscell(FramesMat)
          Frame=FramesMat{k};

        elseif length(size(FramesMat))<3
           Frame=FramesMat;
        
        elseif ~isempty(size(FramesMat),3)
           Frame=FramesMat(:,:,k);
        end  

     
       if max(Frame(:))<1

        figure
        colormap(colortype)
        h=imagesc(Frame);
        set(gca,'YDir','normal');
        axis equal tight 
        %h = imshow(inputimage,[]);
        saveas(h,[savename,'_frame',num2str(k)],'tif')
       else 
           
        
    figure
    colormap(colortype)
     Frame=flipud(Frame);
     h=(imshow(Frame,[]));
%        set(h,'YDir','normal');
        imagetest = getimage(h);
        cmap = colormap(colortype);
       
        if max(imagetest(:))<65535
        imagetest=uint16(imagetest);
        else 
        imagetest=imagetest/256;
        imagetest=uint16(imagetest);
        end
        %imwrite(imagetest,cmap,['Image_frame',num2str(k),'.tif']);
    
        imwrite(imagetest,cmap,[savename,'_frame',num2str(k),'.tif'],'Compression','none')%,...
%'WriteMode','append')
       end
       
 %    pause
    close
    end
end
```
