## Supplementary material for "Microvillar cartography: a super-resolution single-molecule imaging method to map the positions of membrane proteins with respect to cellular surface topography": Analysis MATLAB codes: mean_std_microvilli_fast_Roi.m

```
      %%%input: N= number of files for each angle
%%%delta(window size of stacked movie),LocMicrovilli(location of "microvilli" region,SaveFileName,axisxnm
%%%output: mean of DeltaZ and standard deviation of DeltaZ

function [meanLocDeltaZ,stdLocDeltaZ]= mean_std_microvilli_fast_Roi(N,delta,LocMicrovilli,SaveFileName,axispixel,zhighet,ws,range_x,range_y)
%% options

clear meanLocDeltaZ stdLocDeltaZ

makeThreshold=0;

if makeThreshold==0;
%% calculate meanLocDeltaZ and stdLocDeltaZ
    for kk=1:N
    LocDelta(:,:,kk)=delta(:,:,kk).*LocMicrovilli;
    %imshow(LocDelta(:,:,kk));
    end
    meanLocDeltaZ=mean(LocDelta,3);
    stdLocDeltaZ=std(LocDelta,0,3);
    index=(find(meanLocDeltaZ==0));
    meanLocDeltaZ(index)=NaN;
    stdLocDeltaZ(index)=NaN;

%     figure
%     surf(axisxnm,axisxnm,meanLocDeltaZ);
%     figure
%     surf(axisxnm,axisxnm,stdLocDeltaZ);
%     figure
%     surfc_YM(axisxnm,axisxnm,meanLocDeltaZ,meanLocDeltaZ,stdLocDeltaZ)

else
    
end
%%  Create figure of mean of deltaZ
    figure;
    colormap('gray');

    % Create axes
    axes1 = axes('Parent',gcf);
    view(axes1,[83 12]);
    grid(axes1,'on');
    hold(axes1,'all');

        
        surf(axispixel,axispixel,meanLocDeltaZ,'EdgeColor','none')
        axis([0 ws 0 ws 0 zhighet]);
        xlim(range_x);ylim(range_y);
             set(gca,'FontSize',14)

             title( ['Mean (\delta)Z'],'FontSize',16)
             xlabel('x(pixel)','FontSize',16)
             ylabel('y(pixel)','FontSize',16)
             zlabel('z(nm) Mean','FontSize',16)
        
               
        save(['AveZnm_' SaveFileName,'.mat'],'meanLocDeltaZ','-mat');
        saveas(gcf,['ave3Znm_' SaveFileName],'jpg'); 
        saveas(gcf,['ave3Znm_' SaveFileName],'fig'); 

%        pause 
        
%% Create figure of mean of STD of delta Z 
    figure;
    axes1 = axes('Parent',gcf);
    view(axes1,[83 12]);
    grid(axes1,'on');
    hold(axes1,'all');
    colormap('gray');

    surf(axispixel,axispixel,stdLocDeltaZ,'EdgeColor','none')
    axis([1 ws 1 ws 0 zhighet]);
    xlim(range_x);ylim(range_y);
        set(gca,'FontSize',14)
             title( ['STD (\delta)Z'],'FontSize',16)
             xlabel('x(pixel)','FontSize',16)
             ylabel('y(pixel)','FontSize',16)
             zlabel('z (nm) STD','FontSize',16)

        
        save(['std3Znm_' SaveFileName,'.mat'],'stdLocDeltaZ','-mat');
        saveas(gcf,['std3Znm_' SaveFileName],'jpg'); 
% %% create mean and std figure (surfc_YM)    
%     figure
%     colormap('Jet');
% 
%     axes1 = axes('Parent',gcf);
%     view(axes1,[83 12]);
%    %% surfc_YM(axispixel,axispixel,meanLocDeltaZ,meanLocDeltaZ,stdLocDeltaZ) %% surf=AveZi; contour=STDZi
%     axis([1 ws 1 ws 0 zhighet]);
%     xlim(range_x);ylim(range_y);
%     colorbar
%              set(gca,'FontSize',14)
%              title( ['Mean and STD of(\delta)Z'],'FontSize',16)
%              xlabel('x(pixel)','FontSize',16)
%              ylabel('y(pixel)','FontSize',16)
%              zlabel('z(nm) Mean','FontSize',16)
% 
%     
%     saveas(gcf,['AveStd_' SaveFileName],'jpg'); 
%     saveas(gcf,['AveStd_' SaveFileName],'fig'); 
% %    pause
%% create histogram figure (STD of z)
    figure
    hist (stdLocDeltaZ(:),20);
    set(gca,'FontSize',14)
    xlabel('STD (nm)','FontSize',16), ylabel('Frequency','FontSize',16)
   
    saveas(gcf,['histSTD_' SaveFileName],'jpg'); 

end
    % Saving 3D 
%save(['ave3Z_' SaveFileName,'.ascii'],'AveZi','-ascii');
```
