## Supplementary material for "Microvillar cartography: a super-resolution single-molecule imaging method to map the positions of membrane proteins with respect to cellular surface topography": Analysis MATLAB codes: Membrane_receptor_merging_after_correction.m

```
clear all; close all;
samplename='JE2_FM_TCR';
x_range=[85 145];%
y_range=[95 155];%
ch_TIRFM=[257 512];
ch_SLN=[1 256];
fprintf('select the TIRFM movies of cell membrane that is acquired at 66.8 degree \n')
[FileName_TIRF899, PathName_TIRF899] = uigetfile( ...
{  '*.fits','Fits Files (*.fits)'; ...
   '*.mat','MAT-files (*.mat)'
   '*.*',  'All Files (*.*)'}, ...
   'Pick Channel 1 (532nm)', ...
   'MultiSelect', 'off');
fprintf('select the TIRFM movies of cell membrane that is acquired at last in the 00um SLN series \n')
[FileName_TIRF00umlast, PathName_TIRF00umlast] = uigetfile( ...
{  '*.fits','Fits Files (*.fits)'; ...
   '*.mat','MAT-files (*.mat)'
   '*.*',  'All Files (*.*)'}, ...
   'Pick Channel 1 (532nm)', ...
   'MultiSelect', 'off');
fprintf('select the TIRFM movies of cell membrane that is acquired at last in the 04um SLN series \n')
[FileName_TIRF04umlast, PathName_TIRF04umlast] = uigetfile( ...
{  '*.fits','Fits Files (*.fits)'; ...
   '*.mat','MAT-files (*.mat)'
   '*.*',  'All Files (*.*)'}, ...
   'Pick Channel 1 (532nm)', ...
   'MultiSelect', 'off');
DisplayFrameN=1;
Movie1_Ch=0;%
Movie2_Ch=1;%
allframe=1; 
FileNameHead_RS=[FileName_TIRF899(1:end-5),num2str(x_range(1)),'_',num2str(x_range(2)),'_',num2str(y_range(1)),'_',num2str(y_range(2))];
FileNameHead_DE00um=[FileName_TIRF00umlast(1:end-5),num2str(x_range(1)),'_',num2str(x_range(2)),'_',num2str(y_range(1)),'_',num2str(y_range(2))];
FileNameHead_DE04um=[FileName_TIRF04umlast(1:end-5),num2str(x_range(1)),'_',num2str(x_range(2)),'_',num2str(y_range(1)),'_',num2str(y_range(2))];

ws=9;

ApplyfiltertoCC=1;
kernelsizeCC=5;
getfocalplane='nan';
BxAcheck=0; %% for check AxB (default) and BxA 
OptionNewpeak=1;

%% find first and last frame of all file
StartFrame   = 1;
fitsfilename_TIRF899 = [PathName_TIRF899 FileName_TIRF899];
   % cd (PathName2);
    info_TIRF899=fitsinfo(fitsfilename_TIRF899);
    FileDimention_TIRF899=info_TIRF899.PrimaryData.Size;
    if length(FileDimention_TIRF899)<3; EndFrame_TIRF899 = 1;
    else
    EndFrame_TIRF899     =(FileDimention_TIRF899(3)); 
    end
    
    fitsfilename_TIRF00umlast = [PathName_TIRF00umlast FileName_TIRF00umlast];
   % cd (PathName2);
    info_TIRF00umlast=fitsinfo(fitsfilename_TIRF00umlast);
    FileDimention_TIRF00umlast=info_TIRF00umlast.PrimaryData.Size;
    if length(FileDimention_TIRF00umlast)<3; EndFrame_TIRF00umlast = 1;
    else
    EndFrame_TIRF00umlast     =(FileDimention_TIRF00umlast(3)); 
    end
    
    fitsfilename_TIRF04umlast = [PathName_TIRF04umlast FileName_TIRF04umlast];
   % cd (PathName2);
    info_TIRF04umlast=fitsinfo(fitsfilename_TIRF04umlast);
    FileDimention_TIRF04umlast=info_TIRF04umlast.PrimaryData.Size;
    if length(FileDimention_TIRF04umlast)<3; EndFrame_TIRF04umlast = 1;
    else
    EndFrame_TIRF04umlast     =(FileDimention_TIRF04umlast(3)); 
    end
%% find ref image for Registration shift
for jj = StartFrame:EndFrame_TIRF899
           [filesData, FrameB] = fits_shifter_single_Y_v30(PathName_TIRF899,FileName_TIRF899,jj);  % frames_out ; FrameNth value (:,:,N)
     
               FrameB=FrameB(ch_SLN(1):ch_SLN(2),1:end);         
               
    RefImage_Range_RS{jj}=FrameB(y_range(1):y_range(2),x_range(1):x_range(2));
 end
%% find templet image for registration shift
  for jj = StartFrame:EndFrame_TIRF899
               %FrameN=jj; %% 
               % frames_out => FrameNth value (:,:,N)

                   [filesData, FrameA] = fits_shifter_single_Y_v30(PathName_TIRF899,FileName_TIRF899,jj);  % frames_out ; FrameNth value (:,:,N)
     
               FrameA=FrameA(ch_TIRFM(1):ch_TIRFM(2),1:end);  


      FrameATemplate_RS{jj}=FrameA; 
  end
  %% templet image for drift effect at 00um 
     for jj = StartFrame:EndFrame_TIRF00umlast
               

                   [filesData, FrameA] = fits_shifter_single_Y_v30(PathName_TIRF00umlast ,FileName_TIRF00umlast ,jj);  % frames_out ; FrameNth value (:,:,N)

                   
      FrameATemplate_de00um{jj}=FrameA; 
     end
     %% templet image for drift effect at 04um 
     for jj = StartFrame:EndFrame_TIRF04umlast
               

                   [filesData, FrameA] = fits_shifter_single_Y_v30(PathName_TIRF04umlast ,FileName_TIRF04umlast ,jj);  % frames_out ; FrameNth value (:,:,N)

                   
      FrameATemplate_de04um{jj}=FrameA; 
     end
  %% find ref image for drift effect at 00 um
  for jj = StartFrame:EndFrame_TIRF899
               %FrameN=jj; %% 
               % frames_out => FrameNth value (:,:,N)

                   [filesData, FrameB] = fits_shifter_single_Y_v30(PathName_TIRF899,FileName_TIRF899,jj);  % frames_out ; FrameNth value (:,:,N)
     
               FrameB=FrameB(ch_TIRFM(1):ch_TIRFM(2),1:end);  


      RefImage_Range_de00um{jj}=FrameB(y_range(1):y_range(2),x_range(1):x_range(2));
  end  
   %% find ref image for drift effect at 00 um
  for jj = StartFrame:EndFrame_TIRF899
               %FrameN=jj; %% 
               % frames_out => FrameNth value (:,:,N)

                   [filesData, FrameB] = fits_shifter_single_Y_v30(PathName_TIRF899,FileName_TIRF899,jj);  % frames_out ; FrameNth value (:,:,N)
     
               FrameB=FrameB(ch_TIRFM(1):ch_TIRFM(2),1:end);  


      RefImage_Range_de04um{jj}=FrameB(y_range(1):y_range(2),x_range(1):x_range(2));
  end     
     
    
     %% create new folder 
     NewfolderName=['Shift_drift_effect_calcalculation']; 
     if (exist (NewfolderName)==0)
    mkdir(PathName_TIRF899,NewfolderName)
    else (exist(NewfolderName)==1)
    SaveNewName=input('NewName(1)/Deletename(0)??:')
    if SaveNewName==1;
    NewfolderName=input('SaveFolderName?? : ','s'); 
    else
        rmdir ('Shift_drift_effect_calcalculation','s');
    end
    mkdir(PathName_TIRF899,NewfolderName)
     end
      %              mkdir(PathName_TIRF899,NewfolderName)
                    cd(PathName_TIRF899);
                    cd(NewfolderName);
                    
%% Registration shift
FileNameHead1_TIRF899=[FileName_TIRF899(1:end-5),num2str(x_range(1)),'_',num2str(x_range(2)),'_',num2str(y_range(1)),'_',num2str(y_range(2))];
[ShiftResult]=Result_NormXcorr2_shift_v5_Range_ChannelComp(FrameATemplate_RS,RefImage_Range_RS,PathName_TIRF899,FileNameHead1_TIRF899,0,ApplyfiltertoCC,kernelsizeCC,getfocalplane,x_range,y_range,ws);
save('ResultXcoor_TIRF899_ShiftResult','ShiftResult');
[meanFrameResult]=Result_NormXcorr2_shift_v6_Range_ChannelComp_2MeanMovie(FrameATemplate_RS,RefImage_Range_RS,PathName_TIRF899,FileNameHead1_TIRF899,ApplyfiltertoCC,kernelsizeCC,getfocalplane,x_range,y_range,ws,OptionNewpeak);
save('ResultXcoor_TIRF899_meanFrameResult','meanFrameResult');
chx=-meanFrameResult(7);
chy=-meanFrameResult(8);
save('chx','chx');
save('chy','chy');
close all
%% effect of drift on 00um plane
cd(PathName_TIRF899);
cd(NewfolderName);
FileNameHead1_TIRF00umlast=[FileName_TIRF00umlast(1:end-5),num2str(x_range(1)),'_',num2str(x_range(2)),'_',num2str(y_range(1)),'_',num2str(y_range(2))];
[ShiftResult]=Result_NormXcorr2_shift_v5_Range_ChannelComp(FrameATemplate_de00um,RefImage_Range_de00um,PathName_TIRF899,FileNameHead1_TIRF00umlast,0,ApplyfiltertoCC,kernelsizeCC,getfocalplane,x_range,y_range,ws);
save('ResultXcoor_TIRF00umlast_ShiftResult','ShiftResult');
[meanFrameResult]=Result_NormXcorr2_shift_v6_Range_ChannelComp_2MeanMovie(FrameATemplate_de00um,RefImage_Range_de00um,PathName_TIRF899,FileNameHead1_TIRF00umlast,ApplyfiltertoCC,kernelsizeCC,getfocalplane,x_range,y_range,ws,OptionNewpeak);
save('ResultXcoor_TIRF00umlast_meanFrameResult','meanFrameResult');
shiftx=meanFrameResult(7);
shifty=meanFrameResult(8);
save('shiftx','shiftx');
save('shifty','shifty');
close all
%% effect of drift on 04um plane
cd(PathName_TIRF899);
cd(NewfolderName);
FileNameHead1_TIRF04umlast=[FileName_TIRF04umlast(1:end-5),num2str(x_range(1)),'_',num2str(x_range(2)),'_',num2str(y_range(1)),'_',num2str(y_range(2))];
[ShiftResult]=Result_NormXcorr2_shift_v5_Range_ChannelComp(FrameATemplate_de04um,RefImage_Range_de04um,PathName_TIRF899,FileNameHead1_TIRF04umlast,0,ApplyfiltertoCC,kernelsizeCC,getfocalplane,x_range,y_range,ws);
save('ResultXcoor_TIRF04umlast_ShiftResult','ShiftResult');
[meanFrameResult]=Result_NormXcorr2_shift_v6_Range_ChannelComp_2MeanMovie(FrameATemplate_de04um,RefImage_Range_de04um,PathName_TIRF899,FileNameHead1_TIRF04umlast,ApplyfiltertoCC,kernelsizeCC,getfocalplane,x_range,y_range,ws,OptionNewpeak);
save('ResultXcoor_TIRF04umlast_meanFrameResult','meanFrameResult');
shift2x=meanFrameResult(7);
shift2y=meanFrameResult(8);
save('shift2x','shift2x');
save('shift2y','shift2y');     
close all     
     
     


%% VATIRFM-SLN merging
%% option for chanel 
minoption=0; %% min option ==0 min minoption==1 cm

range_x=x_range;
range_y=y_range;

usexcoorFitset=1; %!!!!!!!!
channelcorrelction=1;
markersize =1000;

if channelcorrelction==1
shiftx=shiftx+chx; %% plus shift to right // mimusshifst to left : 0.7 (-0.2)
shifty=shifty+chy; %% plus shift up // minus shift down -->shift to down: -0.4 (0.4) 
shift2x=shift2x+chx; %% plus shift to right // mimusshifst to left : 0.7 (-0.2)
shift2y=shift2y+chy;%
end

zMax=600;
%minoption=1; % option ==0 (minvalue of pixel); option==1 (cntroid of the tip)
resolvenm = 10; % nm: for tip by tip analysis it was 25
maxsteppix= 7; % for tip by tip analysis %pixel max (7*2*66.67nm)
zSTDlimit=50; %% I don't want to use;use 0 
dualfocalplane=1; %% choose two Fitset Table (for 00 um to -04um merge together) 
%focalplane=-4;
local_scale=1;
%VAtype=3; %% meanLocTip=1(local tip region; 'AveZnm....'); at specific angle=2(z) ;MeanSTDdelta(celltype) total reconstruction from several angle
datatype=3; %% 1=FitsetPT; 2=CMsetPT ; 3=PallOKPT;
tipws=7; 
limitNumofMols=10; %% ignore Tip region molecules if total num of mols are smaller than limitNumofMols  
% allws=1;
% xcnt=125;%% desired center pixel x-coordinate; usually center of mass of object 
% ycnt=125;%% desired center pixel y-coordinate; usually center of mass of object
% ws=101;%% must be odd number


%% Analysis 1 _all region
%% open VA Reconstruction files 
data1=[];
StormData2 =[];
MergeFigure = 1;
fprintf('select the Membrane_reconstruction folder \n')
folder1 = uigetdir; % check the help for uigetdir to see how to specify a starting path, which makes your life easier
VApath = [folder1 '\'];
Listing = dir([VApath,'*.mat']);
cd (VApath)

for nn= 1:size(Listing,1)
    filename= Listing(nn).name;
    load (filename)
end
fprintf('select the combinexcCorrct_F5 folder \n')
folder2 = uigetdir; 
StormPath = [folder2 '\'];
StormList = dir([StormPath,'*.mat']);

cd(StormPath)
if usexcoorFitset==1
    
for ff= 1:size(StormList ,1)
   
 stormDatafile=[StormList(ff).name];

if  ~isempty(strfind(stormDatafile,'0um'))...
        && ~isempty(strfind(stormDatafile,samplename)) ...
        && ~isempty(strfind(stormDatafile,'xcCorrct'))
    
StormFileName1=stormDatafile;
StormData1=importdata(StormFileName1);

%data = open([stormDataPath, stormDatafile{1}]);
end

if  ~isempty(strfind(stormDatafile,'4um'))...
        && ~isempty(strfind(stormDatafile,samplename))...
         && ~isempty(strfind(stormDatafile,'xcCorrct')) 
StormFileName2=stormDatafile;
StormData2 = importdata(StormFileName2);
end


end


Storm2Path=StormPath;
if isempty(StormData2)
  
    folder3 = uigetdir; 
Storm2Path = [folder3 '\'];
Storm2List = dir([Storm2Path,'*.mat']);

cd(Storm2Path)

for ff= 1:size(Storm2List ,1)
   
 stormDatafile=[Storm2List(ff).name];

if  ~isempty(strfind(stormDatafile,'4um'))...
        && ~isempty(strfind(stormDatafile,samplename))...
      %   && ~isempty(strfind(stormDatafile,'xcCorrct')) 
StormFileName2=stormDatafile;
StormData2 = importdata(StormFileName2);
end
end
    
end

elseif usexcoorFitset==0
    
    for ff= 1:size(StormList ,1)
   
 stormDatafile=[StormList(ff).name];

if  ~isempty(strfind(stormDatafile,'0um'))...
        && ~isempty(strfind(stormDatafile,samplename))% ...
     %   && ~isempty(strfind(stormDatafile,'xcCorrct'))
    
StormFileName1=stormDatafile;
StormData1=importdata(StormFileName1);

%data = open([stormDataPath, stormDatafile{1}]);
end

if  ~isempty(strfind(stormDatafile,'4um'))...
        && ~isempty(strfind(stormDatafile,samplename))%...
        % && ~isempty(strfind(stormDatafile,'xcCorrct')) 
StormFileName2=stormDatafile;
StormData2 = importdata(StormFileName2);
end


    end
end 
cd(StormPath)

%% save Roi Fitset

%% RoiFitset analysis
SaveFolderName=(['RoiFitset_xcCorrect']);
if channelcorrelction==1
SaveFolderName=(['Ch_RoiFitset_xcCorrect']);
end

if shiftx~=0 || shiftx~=0
    RoifolderName=([SaveFolderName,'_1x',num2str(shiftx),'_1y',num2str(shifty),'_2x_',num2str(shift2x),'_2y_',num2str(shift2y)]);
else
    RoifolderName=SaveFolderName;
end

if (exist (RoifolderName)==0);
        mkdir(StormPath,RoifolderName)
end
    
cd (RoifolderName);  


if ~isempty(StormData1)
postData1shift(:,1)=StormData1(:,2)+shifty;
postData1shift(:,2)=StormData1(:,3)+shiftx;
StormData1(:,2)=StormData1(:,2)+shifty;
StormData1(:,3)=StormData1(:,3)+shiftx;
[RoiFitsetPT1]=FitsetRoi_range(StormData1,range_x,range_y);
%FitsetFileName=(StormFileName1(1:end-4));
save(['Roixc_',StormFileName1(1:end-4)],'RoiFitsetPT1');

end
 
if ~isempty(StormData2)
postData2shift(:,1)=StormData2(:,2)+shift2y;
postData2shift(:,2)=StormData2(:,3)+shift2x;
StormData2(:,2)=StormData2(:,2)+shift2y;
StormData2(:,3)=StormData2(:,3)+shift2x;
[RoiFitsetPT2]=FitsetRoi_range(StormData2,range_x,range_y);
%FitsetFileName=(StormFileName2(1:end-4));
save(['Roixc_',StormFileName2(1:end-4)],'RoiFitsetPT2');
end


%% make VASTORM merge new folder
cd (VApath);  

SaveFolderName=(['VASTORM_SLN_merge_Results_']);


if shiftx~=0 || shiftx~=0
%MergeFolder=([samplename,'_',SaveFolderName,'_shiftx_',num2str(shiftx),'_shifty_',num2str(shifty),'_',num2str(range_x(1)),num2str(range_x(2)),num2str(range_y(1)),num2str(range_y(2))]);
MergeFolder=([samplename,'_',SaveFolderName,'_',num2str(range_x(1)),'_',num2str(range_x(2)),'_',num2str(range_y(1)),'_',num2str(range_y(2))]);

else
MergeFolder=SaveFolderName;
end

if (exist (MergeFolder)==0);
    mkdir(pwd,MergeFolder)
end


cd (MergeFolder);  
%% mean VA figure

myColorMap = colormap('colorcube(16)');% 
%savefilename=[samplename];
[resultImage]=VA_Rec_2D_color(samplename,myColorMap,MeanSTDdelta{1},local_scale,range_x,range_y,markersize,zMax);
close


%% All cell area; 

myColorMap = colormap('gray(16)');% 
%myColorMap =(flipud(myColorMap));set the colormap used  %autumn
meanLocDeltaZ(isnan(meanLocDeltaZ)) = zMax ;

savefilename=['TipLoc_',StormFileName1];
%focalplanename='0nm';
stormcolor= [1 0 0];
[resultImage]=VA_STORM_merge(stormcolor,myColorMap,meanLocDeltaZ,postData1shift,savefilename,0,local_scale,range_x,range_y,markersize,zMax);
%pause;
close;


savefilename=['TipLoc_',StormFileName2];
%focalplanename='-400nm';
stormcolor= [0 0 1];
[resultImage]=VA_STORM_merge(stormcolor,myColorMap,meanLocDeltaZ,postData2shift,savefilename,-4,local_scale,range_x,range_y,markersize,zMax);
close;

myColorMap = colormap('hsv(16)'); %set the colormap used  %autumn
%myColorMap =(flipud(myColorMap));0

%focalplanename='0nm';
savefilename=['Mean_',StormFileName1];
stormcolor= [0 0 0];
[resultImage]=VA_STORM_merge(stormcolor,myColorMap,MeanSTDdelta{1},postData1shift,savefilename,0,local_scale,range_x,range_y,markersize,zMax);
close;

savefilename=['Mean_',StormFileName2];
stormcolor= [0 0 0];
%focalplanename='-400nm';
[resultImage]=VA_STORM_merge(stormcolor,myColorMap,MeanSTDdelta{1},postData2shift,savefilename,-4,local_scale,range_x,range_y,markersize,zMax);
close
% hold on 


%% dual plane merge figures; 
savefilename=['TipLoc_',StormFileName1];
stormcolor1= [1 0 0];
stormcolor2= [0 0 1];
myColorMap = colormap('gray(16)');
[resultImage]=VA_STORM_dual(stormcolor1,stormcolor2,myColorMap,meanLocDeltaZ,postData1shift,postData2shift,savefilename,local_scale,range_x,range_y,markersize,zMax);
% [resultImage]=VA_STORM_merge(stormcolor,myColorMap,MeanSTDdelta{1},postData2shift,savefilename,-4,local_scale,range_x,range_y,markersize,zMax);
% hold on 
close;
 
savefilename=['Mean_',StormFileName1];
stormcolor1= [0 0 0];
stormcolor2= [0 0 1];
myColorMap = colormap('hsv(16)');
[resultImage]=VA_STORM_dual(stormcolor1,stormcolor2,myColorMap,MeanSTDdelta{1},postData1shift,postData2shift,savefilename,local_scale,range_x,range_y,markersize,zMax);
% [resultImage]=VA_STORM_merge(stormcolor,myColorMap,MeanSTDdelta{1},postData2shift,savefilename,-4,local_scale,range_x,range_y,markersize,zMax);
% hold on 
close;

%% 2Plot_dSTORM
%cd (VAPath);  

savefilename=['2plot_',StormFileName1];
stormcolor1= [1 0 0];
stormcolor2= [0 0 1]; % olive
% myColorMap = colormap('gray(16)');
[resultImage]=dSTORM_2plot(stormcolor1,stormcolor2,postData1shift,postData2shift,savefilename,1,range_x,range_y,markersize,zMax);
close;

%savefilename=[StormFileName1];
stormcolor1= [1 0 0];
stormcolor2= [0 0 1]; % olive (0 0.5 0)
% myColorMap = colormap('gray(16)');

[resultImage]=dSTORM_dualplane_Roi(stormcolor1,stormcolor2,postData1shift,postData2shift,StormFileName1,StormFileName2,1,range_x,range_y,markersize,zMax);
close;

%% 2colorplot_dstorm
%colortype='cool';
myColorMap = colormap('cool(16)');
miniPN=3000; maxiPN=8000;
savefilename=['Cool_',StormFileName1];
[ColorFigure]=NumPho_2plot_color(myColorMap,StormData1,StormData2,savefilename,miniPN,maxiPN,range_x,range_y,markersize);
%miniPN,maxiPN
close 

myColorMap = colormap('jet(16)');
miniPN=1500; maxiPN=10000;
savefilename=['jet_',StormFileName1];
[ColorFigure]=NumPho_2plot_color(myColorMap,StormData1,StormData2,savefilename,miniPN,maxiPN,range_x,range_y,markersize);
close

%% Tip Col map label and LocTipTable_Sort_by Z (Ascending order)
myColorMap=colormap('pink(16)');
MyMarkerColor=[1 1 1];
savefilename=['RoiLocTipMap_',samplename,'_',samplename,'_',num2str(range_x(1)),'_',num2str(range_x(2)),'_',num2str(range_y(1)),'_',num2str(range_y(2))];
[ret,ROILocTipsTable]=VAtipRIO_sort_Map(savefilename,meanLocDeltaZ,myColorMap,MyMarkerColor,LocTipsTable,range_x,range_y,markersize,minoption,zMax);
close all

%% Tip Col map label and LocTipTable_Sort_by Z (Ascending order)
myColorMap=colormap('flag(16)');
MyMarkerColor=[0 0 0];
%minoption=1; % option ==1 (centerofmass); option==1 (min value of locTip)
savefilename=['conLocTipMap_',samplename,'_',num2str(range_x(1)),'_',num2str(range_x(2)),'_',num2str(range_y(1)),'_',num2str(range_y(2))];
[ret,ROILocTipsTable]=VAtipRIO_contour_Map(savefilename,meanLocDeltaZ,myColorMap,MyMarkerColor,LocTipsTable,range_x,range_y,markersize,minoption,zMax);
close all

if zSTDlimit>0
%% Tip Col map label and LocTip by Tip table_sort by Z(limitzSTD)
[zSTD_LocTipsTable]=zSTDlimit_sort(ROILocTipsTable,zSTDlimit);
myColorMap=colormap('pink(16)');
MyMarkerColor=[1 1 1];
% minoption=1;
savefilename=['zSTDlim_',num2str(zSTDlimit),'nm_LocTipMap_',samplename,'_',num2str(range_x(1)),'_',num2str(range_x(2)),'_',num2str(range_y(1)),'_',num2str(range_y(2))];
[ret,zSTD_ROILocTipsTable]=VAtipRIO_sort_Map(savefilename,meanLocDeltaZ,myColorMap,MyMarkerColor,zSTD_LocTipsTable,range_x,range_y,markersize,minoption,zMax);
close all

%% Tip Col map label and LocTipTable_Sort_by Z (Ascending order)
myColorMap=colormap('pink(16)');
MyMarkerColor=[0 0 0];
%minoption=1; % option ==1 (centerofmass); option==1 (min value of locTip)
savefilename=['zSTDlim_',num2str(zSTDlimit),'nm_conLocTipMap_',samplename,'_',num2str(range_x(1)),'_',num2str(range_x(2)),'_',num2str(range_y(1)),'_',num2str(range_y(2))];
[ret,zSTD_ROILocTipsTable]=VAtipRIO_contour_Map(savefilename,meanLocDeltaZ,myColorMap,MyMarkerColor,zSTD_LocTipsTable,range_x,range_y,markersize,minoption,zMax);
close all
end

Mergepath = [folder1 '\' MergeFolder '\'];
cd (Mergepath); 

save('VA_analysis_allinone');

AllinOneSetting.StormFileName1=StormFileName1
AllinOneSetting.StormFileName2=StormFileName2;
AllinOneSetting.VApath=VApath;
AllinOneSetting.usexcoorFitset=usexcoorFitset
AllinOneSetting.StormPath=StormPath;
if usexcoorFitset==1
    
AllinOneSetting.StormPath2=Storm2Path;
elseif usexcoorFitset==0
AllinOneSetting.StormPath2=StormPath;
end

AllinOneSetting.Mergepath=Mergepath;
AllinOneSetting.minoption=minoption;
AllinOneSetting.samplename=samplename;
AllinOneSetting.range_x=range_x;
AllinOneSetting.range_y=range_y;
AllinOneSetting.markersize=markersize;
AllinOneSetting.shiftx=shiftx;
AllinOneSetting.shifty=shifty;
AllinOneSetting.shift2x=shift2x;
AllinOneSetting.shift2y=shift2y;
AllinOneSetting.channelcorrelction=channelcorrelction;
AllinOneSetting.chx=chx; %% 
AllinOneSetting.chy=chy; %%
AllinOneSetting.zMax=zMax;
AllinOneSetting.resolvenm=resolvenm ;
AllinOneSetting.maxsteppix=maxsteppix;
AllinOneSetting.zSTDlimit=zSTDlimit;
AllinOneSetting.dualfocalplane=dualfocalplane;
AllinOneSetting.tipws=tipws;
AllinOneSetting.limitNumofMols=limitNumofMols;

save([samplename,'_AllinOneSetting'],'AllinOneSetting');


%% creating VATIRFM-SLN merge image
load('VA_analysis_allinone.mat')
scalebarsize=500;
range=64;
zdist=400;
stormresolution=15;
zMax=400;
SaveFolderName=(['VATIRFM-SLN_merged_image_' samplename]); % '_' num2str(midColor) '_' num2str(zdist)]);


if (exist (SaveFolderName)==0)
    mkdir(VApath,SaveFolderName)
end
cd(VApath)
cd(SaveFolderName)

[aa]=find(meanDeltaZ>zMax);
meanDeltaZnobgd=meanDeltaZ;
meanDeltaZnobgd(aa)=zMax;
meanDeltaZ=meanDeltaZnobgd;
whitebg('white');

NewColorMap=flipud(colormap(jet(range)));NewColorMap(range,:)=0;

close
myColorMap=NewColorMap;
%myColorMap=scaledcolormap_shirsendu(NewColorMap,range,zdist,midColor,zMax);%(name_colormap,flippedoption,range,zdist,midColor)
SaveName=['Hot_W_',SaveFileName,'00um'];
StormColor1=[1 1 1];
Merge_createfigure_Surfshade_STORM_3Dlike(meanDeltaZ,RoiFitsetPT1,zMax,range_x,range_y,SaveName,myColorMap,StormColor1,scalebarsize,stormresolution);
close
StormColor1=[1 1 1];
SaveName=['Hot_W_',SaveFileName,'04um'];
Merge_createfigure_Surfshade_STORM_3Dlike(meanDeltaZ,RoiFitsetPT2,zMax,range_x,range_y,SaveName,myColorMap,StormColor1,scalebarsize,stormresolution);
close
%myColorMap=scaledcolormap_shirsendu(NewColorMap,range,zdist,midColor,zMax);%(name_colormap,flippedoption,range,zdist,midColor)
SaveName=['Hot_B_',SaveFileName,'00um'];
StormColor1=[0 0 0];
Merge_createfigure_Surfshade_STORM_3Dlike(meanDeltaZ,RoiFitsetPT1,zMax,range_x,range_y,SaveName,myColorMap,StormColor1,scalebarsize,stormresolution);
close
StormColor1=[0 0 0];
SaveName=['Hot_B_',SaveFileName,'04um'];
Merge_createfigure_Surfshade_STORM_3Dlike(meanDeltaZ,RoiFitsetPT2,zMax,range_x,range_y,SaveName,myColorMap,StormColor1,scalebarsize,stormresolution);

     
     
     
 
```
