## Supplementary material for "Microvillar cartography: a super-resolution single-molecule imaging method to map the positions of membrane proteins with respect to cellular surface topography": Analysis MATLAB codes: Membrane_reconstruction_VATIRFM.m

```
clc;close all;clear all;

ch=1; % ch=0 all ; ch=1 ;  ch=2;

range_x=[85 145];
range_y=[95 155];

markersize =1000; 

subbac=1;

%if subbac==0 && subbacmin==0;
subbacmaxangle=0; %% for sub from window edgevalues of the largest angle

TipCentroid=0; %% for option for not fixed 
Threshold=5; %(threshold for LogFiltered Intensity) 
ws=256;

%% variable settings
% Center coordinate (where should be the center)
% xcenter=125;%% desired center pixel x-coordinate; usually center of mass of object 
% ycenter=125 ;%% desired center pixel y-coordinate; usually center of mass of object
% ws=101;%% must be odd number
%minoption=1;
%samplename=['EFF5'];
%zMax=450;

wl=532;%(nm) wavelength 
n1=1.52;%glass or immersion oli
n2=1.35;%PBS % BK=1.35
% xynmscale=0;
zMax=600;
pixel=66.67;

%% angle Table 
ta = [63.03 64.23 65.48 66.79 68.18 69.66 71.25 72.98];%    

%% open fits files or tif files
[FNlist, Path, Filterindex] = uigetfile( ...
{  '*.fits','Fits Files (*.fits)'; ...
   '*.tif','fit Files (*.tif)'; ...
   '*.*',  'All Files (*.*)'}, ...
   'Pick a file', ...
   'MultiSelect', 'on');

cd (Path)
Motherfolder = Path; 


[IgStack,FileName,NumofFiles,GetVA,meanbac]=OpenVAfigures_Roi(FNlist,Path,ws,subbac,subbacmaxangle,ch,range_x,range_y);%% 1.6 gaussian filter sigmasize:1.6 and size 5x5 (sigma*3)

FileName;

getparts = strread(FileName,'%s','delimiter','_');

SavefolderName=strcat(getparts{1},'_',getparts{2},'_',getparts{3});


NewfolderName=['Membrane_reconstruction_' SavefolderName];
if (exist (NewfolderName)==0)
    mkdir(pwd,NewfolderName)
else (exist(NewfolderName)==1)
    SaveNewName=input('NewName??(Y==>1,N==>0):')
    if SaveNewName==1;
    NewfolderName=input('SaveFolderName?? : ','s'); 
    end
mkdir(pwd,NewfolderName)
end
cd (NewfolderName);  


N=NumofFiles;


%% Get penetration Depth(D) and T at each angle;
cnt=0;
for fa=1:N
    %% penetration depth
    D =(wl/(4*pi))*(n1^2*(sind(ta(fa)))^2-n2^2)^(-1/2);
%D2=(wl/(4*pi*n1))/((sind(ta))^2-(n2/n1)^2)^(0.5);same as above
% penetration depth %% sind==> sin
% InvD=(1/D); 
    %% T = transmission factor (linearly polaized --> incorrect!!!)
    T=4*((cosd(ta(fa)))^2)/(1-((n2/n1)^2));%% cosd==> cos
cnt=cnt+1; % counter
%X=1/D;

Tn(cnt)=T ;
Dn(cnt)=D ;
xn(cnt)=1/D;

%Q=IgStack(:,:,cnt);
%size(Q)

end 


%% When I(a)=Imax(a) when d=0;
for nr=1:N %% nr =number of files
        if   iscell(FNlist);     
             FileName=FNlist{nr};
        else 
             FileName=FNlist;
        end
%         
        FileNamePart= ['fix' FileName(1:end-5)];
        

        

Q(:,:)=IgStack(:,:,nr);

Lmax=max((Q(:)));            
LmaxSeries(nr)=Lmax;

delta(:,:,nr)=Dn(nr)*log((Lmax)./Q);%%%%;
if isreal (delta)
    delta=delta;
else 
    delta=real(delta);
end
%end

%if allws==1
%ws=size(Q,2);
%end

%% 3D meshgrid

% xy scale 
[x,y] = meshgrid(1:ws);
z=delta(:,:,nr);

% Create figure
figure1 = figure;
colormap('gray');

% Create axes
axes1 = axes('Parent',figure1);
view(axes1,[83 12]);
grid(axes1,'on');
hold(axes1,'all');

% Create surf
surf(x,y,z,'EdgeColor','none')
axis([1 ws 1 ws 0 zMax]);title(num2str(ta(nr)),'FontSize',20);
%addscale3_um_range_label(range_x,range_y,markersize,0);
xlim(range_x);
ylim(range_y);
zlabel('\delta Z (nm)','FontSize',16);
xlabel('x(pixel)','FontSize',16);
ylabel('y(pixel)','FontSize',16);
shading interp;
light;
lighting phong;
% set(gca,'FontSize',14)
% Saving 3D 
save(['RelZ_' FileNamePart,'.ascii'],'z','-ascii');
save(['RelZ_' FileNamePart,'.mat'],'z','-mat');
saveas(gcf,['3DRelZ_' FileNamePart],'jpg'); 
saveas(gcf,['3DRelZ_' FileNamePart],'fig'); 
% axis equal
%pause
close


%% 2D (x,z)

% xscale change
%axisxnm=round([1:ws]*pixel);
axispixel=([1:ws]);
figure;
plot(axispixel,delta(1:end,1:end,nr));%plot(y,z);
axis([1 ws 0 zMax]); 
xlim([range_x]);
% ylim([range_y]);
title(['xz' num2str(ta(nr))],'FontSize',20);
xlabel('x(pixel)','FontSize',16)
ylabel('y(pixel)','FontSize',16)
%set(gca,'FontSize',14)
%axis equal %%%
saveas(gcf,['2DRelZx_' FileNamePart],'jpg'); 
%saveas(gcf,['2DRelZx_' FileNamePart],'tif'); 
close;
%% 2D (y,z)
figure;
plot(axispixel,delta(1:end,1:end,nr)');%plot(y,z);
axis([1 ws 0 zMax]);
xlim([range_y]);
%ylim([range_y]);
%addscale3_um_range_label(range_x,range_y,markersize,0);
title(['yz' num2str(ta(nr))],'FontSize',16);
saveas(gcf,['2DRelZy_' FileNamePart],'jpg'); 
%saveas(gcf,['2DRelZy_' FileNamePart],'tif'); 
set(gca,'FontSize',14)
clear FileNamePart z ;
%pause
close all;
Zicell{nr}=delta(:,:,nr);

% Get maximum Z 
            Qz=delta(:,:,nr);
            Qz(2:ws-1,2:ws-1)=0;% extanded a logical matrix as a size of ws (13x13)
            ind=find(Qz>0);
            Zmax=mean(Qz(ind)); %% BGD signal (highest value of z)
            ZmaxExp(nr)=Zmax*(1/exp(1))
            
% Get background threshold bac everage*3
            Qz=delta(:,:,nr);
            Qz(2:ws-1,2:ws-1)=0;% extanded a logical matrix as a size of ws (13x13)
            ind=find(Qz>0);
            Zmaxmean=mean(Qz(ind));
            Zstd=std(Qz(ind));
            ZThd(nr)=Zmaxmean-(4*Zstd);       
            BgdzValue(nr)=Zmaxmean; 
[xm,ym]=find(IgStack(:,:,nr)==Lmax);
indexmaxx(nr)=xm;
indexmaxy(nr)=ym;            
%    contour(Qb,'DisplayName','Qb');figure(gcf)
 
end 
clear Q Qz;
%% Penetration Depth and Angle Table for micrometer reading

micrometer= GetVA;
mmGetVA=(micrometer-800).*(0.0254); %% 1 inch= 25.4mm

PenetrationDepthTable=[GetVA;mmGetVA;ta;Dn;ZmaxExp;ZThd]';

%% Save Files
% cd (Path)
%create savefilename ;
     
        SaveFileName= ['fix' FileName(1:end-13)];
             
%save Matrix files  
save(['PDtable_' SaveFileName,'.ascii'],'PenetrationDepthTable','-ascii');
save(['PDtable_' SaveFileName],'PenetrationDepthTable');
save(['RelZall_' SaveFileName],'Zicell');

%% Test Maximum pixel delta zero;
%y= ln(I(ta)/T(ta)); x= 1/D(ta)
%Indexmax=[round(mean(indexmaxx)),round(mean(indexmaxy))];
Indexmax=[(indexmaxx(end-1)),(indexmaxy(end-1))];
for ff=1:N
    XX=(IgStack(Indexmax(1),Indexmax(2),ff))/Tn(ff);
yyn(ff)= log(XX);
xnyyn(ff)=xn(ff)*yyn(ff);
end  

ddt=N*sum(xn.^2)-((sum(xn))^2);
aa= 1/ddt*(sum(xn.^2)*sum(yyn)-sum(xn)*sum(xnyyn));
bb= 1/ddt*((N*sum(xnyyn))-(sum(xn).*sum(yyn)));
close all;
%% Mean and STD of recontructed images from ZiCell
%[AveZi,STDZi]= mean_std_VATIRFm(Zicell,ZmaxExp*2,SaveFileName); 
%% Find the region of mivrovilli (Find microvill based on Ig)
%[LogfilterIgstack,meanLocMicrovilli]=findmicrovilli_LoGFilter(Threshold,N,GetVA,SaveFileName,IgStack);
%[LogfilterIgstack,BWsdil,LocmicrovilliAngle]=findmicrovilli_LoGFilter_angle(Threshold,N,GetVA,SaveFileName,IgStack);
[LogfilterIgstack,LocMicrovili,LocTips]=findmicrovilli_LoGFilter_angle15_Roi(Threshold,N,GetVA,SaveFileName,IgStack,range_x,range_y);
%pause
close all

for kk=1:3
deltapart(:,:,kk)=delta(:,:,kk+2);
end

%% Get mean and STD of deltaZ (height of microvilli)at the microvilli regionn mean only at the angle 898-900; 

[meanLocDeltaZ,stdLocDeltaZ]= mean_std_microvilli_fast_Roi(size(deltapart,3),deltapart,LocMicrovili,SaveFileName,axispixel,zMax,ws,range_x,range_y); 
close all
%% Find location of microvilli Tips (x,y,x)
% inside of function, choose the option for xy coordinate from minimun or centroid of
% delta z of each microvilli
%[LocTipsTable,xyLocMinZ]= microvilliTipLoc_dilate_angle(LocTips,stdLocDeltaZ,meanLocDeltaZ,axisxnm,SaveFileName,TipCentroid);


[LocTipsTable,xyLocMinZ,dilateLocMicrovilli2]= microvilliTipLoc_dilate_angle14_Roi(LocTips,stdLocDeltaZ,meanLocDeltaZ,axispixel,SaveFileName,TipCentroid,range_x,range_y);
close all
% AA=1*(dilateLocMicrovilli2>0);
% 
% isequal(LocMicrovili,AA)
%[LocTipsTable]= microvilliTipLoc(meanLocMicrovilli,meanLocDeltaZ);

%% compare total region and Tip region
%mean all vs tip

meanDeltaZ=mean(deltapart,3);
stdDeltaZ=std(deltapart,0,3);
MeanSTDdelta{1}=meanDeltaZ;
MeanSTDdelta{2}=stdDeltaZ;
%save(['MeanSTDdelta_' SaveFileName],MeanSTDdelta);

%% Dualplot with ViewAngle

title1=['mean 898-900 \delta Z'];
title2=['meanTip 898-900 \delta Z'];
View_range=[90,0];
createfigure_subplot2type1_shade_Roi(meanDeltaZ,meanLocDeltaZ,1,title1,title2,range_x,range_y,View_range);
saveas(gcf,['A_Mean_898-900' SaveFileName],'fig');
saveas(gcf,['A_Mean_898-900' SaveFileName],'jpg');
close

title1=['mean 898-900 \delta Z'];
title2=['meanTip 898-900 \delta Z'];
View_range=[45,-45];
createfigure_subplot2type1_shade_Roi(meanDeltaZ,meanLocDeltaZ,1,title1,title2,range_x,range_y,View_range);
saveas(gcf,['B_Mean_898-900' SaveFileName],'fig');
saveas(gcf,['B_Mean_898-900' SaveFileName],'jpg');
close

title1=['mean 898-900 \delta Z'];
title2=['meanTip 898-900 \delta Z'];
View_range=[135,-45];
createfigure_subplot2type1_shade_Roi(meanDeltaZ,meanLocDeltaZ,1,title1,title2,range_x,range_y,View_range);
saveas(gcf,['C_Mean_898-900 ' SaveFileName],'fig');
saveas(gcf,['C_Mean_898-900' SaveFileName],'jpg');
close
% %std all vs tip
% title1=['std\delta Z'];
% title2=['std \delta Z'];
% createfigure_subplot2type1_Roi(stdDeltaZ,stdLocDeltaZ,2,title1,title2,range_x,range_y)
% saveas(gcf,['Std_' SaveFileName],'fig');
% saveas(gcf,['Std_' SaveFileName],'jpg');
%hist all vs tip
% hist(stdDeltaZ(:),10);hold on freeze
% hist(stdLocDeltaZ(:),10);
close
% cd(Motherfolder)

%% compare each angle and Total area
%% create figure of Logfilter with Threshold       
getVAPart=[GetVA(3),GetVA(4),GetVA(5)];
createfigure_subplots6_surf_Roi(deltapart,zMax,getVAPart,ws,range_x,range_y)
saveas(gcf,['RecAng_898-900' SaveFileName num2str(GetVA(3)) num2str(GetVA(5))],'fig'); %% show anly between 898-900
saveas(gcf,['RecAng_898-900' SaveFileName num2str(GetVA(3)) num2str(GetVA(5))],'jpg'); %% show anly between 898-900
% 
close 
createfigure_subplots6_surf_Roi_shade(deltapart,zMax,getVAPart,ws,range_x,range_y)
%        pause
%        pcolor(LogfilterIgstack(:,:,k));title(['LoGimage Thr' num2str(Threshold)]); 
%        colormap(gray(2))
%        axis equal
saveas(gcf,['RecAng_898-900_shade' SaveFileName num2str(GetVA(3)) num2str(GetVA(5))],'fig'); %% show anly between 898-900
saveas(gcf,['RecAng_898-900_shade' SaveFileName num2str(GetVA(3)) num2str(GetVA(5))],'jpg'); %% show anly between 898-900
%       close;

%% save parameter

ParameterSheet.subbac=subbac;
ParameterSheet.subbacmaxangle=subbacmaxangle; %% for sub from window edgevalues of the largest angle
ParameterSheet.ch=ch; % ch=0 all ; ch=1 ;  ch=2;
ParameterSheet.TipCentroid=TipCentroid; %% for option for not fixed 
ParameterSheet.Threshold=Threshold; %(threshold for LogFiltered Intensity) 
ParameterSheet.range_x=range_x;%% desired center pixel x-coordinate; usually center of mass of object 
ParameterSheet.range_y=range_y;%% desired center pixel y-coordinate; usually center of mass of object
ParameterSheet.ws=ws;%% must be odd number
ParameterSheet.wl=wl;
ParameterSheet.n1=n1;
ParameterSheet.n2=n2;
ParameterSheet.zheight=zMax;
ParameterSheet.ta=ta;
ParameterSheet.GetVA=GetVA;
ParameterSheet.files=FNlist;
ParameterSheet.indexmaxx=indexmaxx;
ParameterSheet.indexmaxy=indexmaxy;
ParameterSheet.indexmax=Indexmax;
ParameterSheet.LmaxSeries=LmaxSeries;
ParameterSheet.maxcount=reshape([IgStack(Indexmax(1),Indexmax(2),:)],1,size(IgStack,3));
ParameterSheet.meanbac=meanbac;
ParameterSheet.Tn=Tn;
ParameterSheet.Dn=Dn;
ParameterSheet.InverseDn=xn;
ParameterSheet.PDtable=PenetrationDepthTable;
ParameterSheet.BgdzValue=BgdzValue; 
ParameterSheet.meanbac=meanbac; 
ParameterSheet.getVAPart=getVAPart; 
%mean background count
%% Save All parameter and mean relative delta results for all; 

save(['ParatmeterSheet' SaveFileName],'ParameterSheet');
MeanSTDdelta{3}=ParameterSheet;
MeanSTDdelta{4}=Zicell;
save(['MeanSTDdelta_' SaveFileName],'MeanSTDdelta');
save('setting');


% 131105
%ver23: save locmap matrix (loctip and dilated) careful to use for the file
%list order! 
% ver 22: "n3", water gap model includes 
%ver21: add shading light effect; logThr figure ylim fixed
%ver20; bgd fix bug
%%ver19:before ver18 
%% ver18; save parameters and mean values but bug when the subbac=0; find the solution
% ver17: add subbac option (choose subbac value from largest angle)
% ver16, multi angle add figure 
%ver14: substract mean background of the original IgStack==> 
%check without sub back improve the detection of microvilli
%mean : 115; 
% check Ta==> ver13 ; Ta, when N2=1.515
%%ver13: type option: membrane(0) actin filament(1)
         % didn't make it yet%%ver12:bwlabel,4; 
%%ver11: find microvillitips from biggest and smallest angle
%%ver10 : find microvillitips by first(lowest angle!)
%%Things to improve: microvilliTipLoc==> Watershade! 
%% For mean, the region of mocrivill determied after i used LoG filter 
% edges = imfilter(AA,h,'replicate'); AA=images
% h = fspecial('log', 14, 0.5); filtertype: log, 14= kernel size (14),
% gaussian sigma
%h = fspecial('log', hsize, sigma)  returns a rotationally symmetric Laplacian of Gaussian filter of size hsize with standard deviation sigma (positive). hsize can be a vector specifying the number of rows and columns in h, or it can be a scalar, in which case h is a square matrix. The default value for hsize is [5 5] and 0.5 for sigma.
%% Main VA Reconstructio n Program (with known angle)
%% calculate "z" from incident angle(assumed by fitting;used by MainVA_calibration_withR)
%% Find tip region of microvilli (tip position x,y,z)
%% gaussian filter option (choose between 0 or 1)
% input: xcenter;ycenter;Penetration Depth Table;fits files; ws; wavelength; n1; n2 
% output %: PDtable =[GetVA;mmGetVA;PDfit;ta;Dn;] penetration depth table
%        %: Zicell{nr}=delta(:,:,nr); Relative z-distance(nm)when Imax is z=0;                                      intensity (Imax:z=0)
%        %: absolute distance from the glass surface -b= slop (need to confirm)   
%        %: figure 2D-RelZx 2D-RelZy & 3D with jpg format;
%        %: figure scale have been changed as nm scale
% bug    % careful for using ws 
% choose option: allws =0 (ws)  or allws=1 (xy size of original movie file frame)
% ver4: center of mass
% choose option: centercoor=1;
% choose multiangle=0;
%% Option settings
%% Figur _creation
range=64;
zMax=400;
[aa]=find(meanDeltaZ>zMax);
meanDeltaZnobgd=meanDeltaZ;
meanDeltaZnobgd(aa)=zMax;
meanDeltaZ=meanDeltaZnobgd;
 whitebg('white');
%%%%%%%%%%%%%%%%%%%%%%%%%%%%%%%%%%

    NewColorMap=flipud(colormap(jet(range)));NewColorMap(range,:)=0;

close
myColorMap=NewColorMap;
figure
set(gcf, 'Position', [ 1  1  1000  950]);
colormap(myColorMap)
surf(meanDeltaZ,'SpecularStrength',0.2,'FaceLighting','phong',...
    'FaceColor','interp',...
    'EdgeColor','none');
axis([range_x range_y 0 zMax]);
view([0 90])
caxis([0, zMax])
axis square
set(gca,'YTickLabel',[]);
set(gca,'XTickLabel',[]);
camlight('headlight')
Li=light;
lightangle(Li,-80,50);
Li2=light;
lightangle(Li2,160,120);
lighting gouraud;
saveas(gcf,['membrane_bottom_view' SavefolderName],'fig');
close
whitebg('black');
myColorMap=NewColorMap;
%myColorMap=scaledcolormap_shirsendu_new(NewColorMap,range,zdist,midColor,zMax);%(name_colormap,flippedoption,range,zdist,midColor)
colormap(myColorMap);
%colormap gray(128);
surf(meanDeltaZ,'EdgeColor','none');
hold on
View_range=[90,0];
view(View_range);
zlim([0 400]);
xlim(range_x);
ylim(range_y);
grid('off');
axis off
camlight('headlight')
Li=light;
lightangle(Li,-80,50);
Li2=light;
lightangle(Li2,0,280);
lighting gouraud;
set(gcf,'color','black')
saveas(gcf,['membrane_side_view' SavefolderName],'fig');
```
