## Supplementary material for "Microvillar cartography: a super-resolution single-molecule imaging method to map the positions of membrane proteins with respect to cellular surface topography": Analysis MATLAB codes: Merge_createfigure_Surfshade_STORM_3Dlike.m

```
function [ret]=Merge_createfigure_Surfshade_STORM_3Dlike(meanDeltaZ,RoiFitsetPT1,zMax,range_x,range_y,SaveFileName,myColorMap,StormColor,markersize,stormresolution)
%% make
figure
set(gcf, 'Position', [ 1  1  1000  950]);
%set(gcf,'Position',get(0,'Screensize'))%enlarge image to full screen\
colormap(myColorMap)
%surf(meanDeltaZ,'EdgeColor','none')
surf(meanDeltaZ,'SpecularStrength',0.2,'FaceLighting','phong',...
    'FaceColor','interp',...
    'EdgeColor','none');
axis([range_x range_y 0 zMax]);
view([0 90])
caxis([0, zMax])
%light;
%lighting phong
%shading interp 

axis square
set(gca,'YTickLabel',[]);
set(gca,'XTickLabel',[]);
%(gray) %colormap(flipud(colormap))%colormap(colorcube(42))
%light;
camlight('headlight')
Li=light;
lightangle(Li,-80,50);
Li2=light;
lightangle(Li2,160,120);
lighting gouraud;
% colormap(myColorMap)
%colormap(flipud(colormap)) % ####################
%% make Frame images
 F = getframe; % image(F.cdata)
 Zimage=F.cdata;% colormap(F.colormap)
% 
% imwrite(Zimage,[SaveFileName,'3Dshade_','.tif'],'Compression','none')%,...
% 
% close


%figure;
%% AutoRescaling of STORM data %
ROIPix_Y=range_y(2)-range_y(1)+1;
ROIPix_X=range_x(2)-range_x(1)+1;
X_Rezero=((RoiFitsetPT1(:,3)-range_x(1))+1)*(size(Zimage,1)/ROIPix_X);  %% nm scale
Y_Rezero=((RoiFitsetPT1(:,2)-range_y(1))+1)*(size(Zimage,2)/ROIPix_Y); 

%% merge the 3D like VA and STORM data
Z = flipdim(Zimage,1);

resultImage = image(Z, 'CDataMapping', 'scaled');
set(gcf, 'Position', [ 1  1  1000  950]);
axis xy;
axis image;
hold on;
colorbar('location','westoutside','fontsize',24')

caxis([0 zMax])
hold on
if stormresolution ==5
scatter(X_Rezero, Y_Rezero, 1, 'o','MarkerEdgeColor',StormColor,'MarkerFaceColor',StormColor);
elseif stormresolution ==15
scatter(X_Rezero, Y_Rezero, 10, 'o','MarkerEdgeColor',StormColor,'MarkerFaceColor',StormColor);%% 15nm
elseif stormresolution ==20
scatter(X_Rezero, Y_Rezero, 3,'o','filled','MarkerEdgeColor',StormColor,'MarkerFaceColor',StormColor);    
end    

%% add scale
hold on;
%markersize= 1000; 
pixelsize=66.67; 
startx=size(Zimage,2)/10;
starty=size(Zimage,2)/10;

scalerange=markersize/pixelsize*(size(Zimage,2)/ROIPix_Y);
Markerx=linspace(0,scalerange);
Markerx=startx+Markerx;
Markery=ones(size(Markerx));
Markery=starty*Markery;
scalebar=line(Markerx,Markery,'Color',[1 1 1],'LineWidth',4); 

set(gca,'XTickLabel',[]);
set(gca,'YTickLabel',[]);
set(gca,'YTick',[]);
set(gca,'XTick',[]);
set(gca,'box','off'); %you take off the box ticks around
set(gca,'YColor','w')
set(gca,'XColor','w')

saveas(gcf,['Light_VA_STORM_' SaveFileName]);%,'jpg'); 
saveas(gcf,['Light_VA_STORM_' SaveFileName],'jpg');
F = getframe;
%GetF=F.cdata;
%colormap(F.colormap)
imwrite(F.cdata,['Light_VA_STORM',SaveFileName,'.tif'],'Compression','none')%,...


%set(gcf,'Position',get(0,'Screensize'))%enlarge image to full screen
% %pause 
% F = getframe;
% GetF=F.cdata;
% % colormap(F.colormap)
% imwrite(F.cdata,['LiVA_STORM_',SaveFileName,'.tif'],'Compression','none')%,...


%close all

% how to make plot data to figures
% uiopen('F:\BestExample\TCR_R\140304_R5\New1R5_TCR_FM143fx_ch12_ex532_896-903\CFver12_R5_TCR_90_170_110_190\scaledual_dSTORM_XC_140304_R5_TCR_selCMPTset_.fig',1)
% 
% set(gca,'XTickLabel',[]);
% set(gca,'YTickLabel',[]);
% set(gca,'YTick',[]);
% set(gca,'XTick',[]);
% set(gca,'box','off'); %you take off the box ticks around
% set(gca,'YColor','w')
% set(gca,'XColor','w')
% set(gcf,'Position',get(0,'Screensize'))%enlarge image to full screen
% F = getframe;
% % image(F.cdata)
% Zimage=F.cdata;
% % colormap(F.colormap)
% imwrite(Zimage,[SaveFileName,'STORM_pix','.tif'],'Compression','none')%,...


end


% 
% 
% % 
% %CREATEFIGURE(ZDATA1)
% %  ZDATA1:  surface zdata
% 
% %  Auto-generated by MATLAB on 10-Oct-2012 10:58:20
% 
% % % Create figure
% % % figure1 = figure;
% % % %hold on 
% % % subplot1=subplot(2,3,1,'Parent',figure1);
% % % view(subplot1,[90 0]);
% % % grid(subplot1,'on');
% % % hold(subplot1,'all');
% % % surf(data1,'Parent',subplot1);
% % % title(['LogFilter',num2str(GetVA(1))],'Fontsize',12);
% % % [OriZLim]=get(subplot1,'ZLim');
% % % zlim(OriZLim)
% % % %zlabel('\delta z (nm)')
% % % % Create subplot
% % % subplot2 = subplot(3,3,2,'Parent',figure1); 
% % % view(subplot2,[90 0]); %[-80 3]]
% % % grid(subplot2,'on');
% % % hold(subplot2,'all');
% % % % Create subplot
% % % surf(data2,'Parent',subplot2);
% % % title(['LogFilter',num2str(GetVA(end))],'Fontsize',12);
% % % zlim(OriZLim)
% % % % zlabel('\delta z (nm)')
% % ws=size(multidata,1);
% % [x,y] = meshgrid(1:ws);
% % 
% % if isequal(length(GetVA),size(multidata,3));
% %     num=length(GetVA);
% % else 
% %     error('error number of GetVA and num of delta')
% % end
% % 
% % if num==1;
% %     a=1;
% %     b=1;
% % elseif num==2 || num==3;
% %     a=1; b=num;
% % elseif num>3 && num<7;
% %     a=2;
% %     b=3;
% % elseif num>=7 && num<=9;
% %     a=3;
% %     b=3;
% % elseif num==10;
% %     a=2;
% %     b=5;
% % else
% %     error('too large stack')
% %     
% % end
% % 
% % figure1 = figure;
% % %set(gcf, 'Position', [10 20 1200 400]);
% % for k=1:num
% % Anglelabel=num2str(GetVA(k));
% % subplot3 = subplot(a,b,k,'Parent',figure1);
% % hold(subplot3,'all');
% % %axes1 = axes('Parent',subplot3);
% % surf(x,y,multidata(:,:,k),'EdgeColor','none','Parent',subplot3);title(Anglelabel);
% % % [OriZLim]=get(subplot3,'ZLim');
% %  xlim([range_x]);
% %  ylim([range_y]);
% %  zlim([0 zlimit]);
% %  zlabel('\delta z (nm)')
% % title([num2str(GetVA(k))],'Fontsize',10);
% % colormap('gray');
% % %view(subplot3,[0 0]);
% % light;
% % lighting phong
% % view(subplot3,[10 -10]);
% % 
% % %axis equal
% % end
% % hold off
% % % saveas(gcf,'SimulationFigures')
% % end
% % Create surf
% 
% 
```
