## Supplementary material for "Microvillar cartography: a super-resolution single-molecule imaging method to map the positions of membrane proteins with respect to cellular surface topography": Analysis MATLAB codes: microvilliTipLoc_dilate_angle14_Roi.m

```
 %% Find Tip Regions (center of mass,pixel position)
%% ver13; save name change
%%% input:% meanLocMicrovilli(meanLocationofMicrovill-->logical matrix),
          % meanLocDeltaZ (mean Delta Z at microvilli region)
%%% LocTipsTable (X,Y,Z)


function[LocTipsTable,xyLocMinZ,LocMicrovilli2,num]= microvilliTipLoc_dilate_angle14_Roi(LocTips,stdLocDeltaZ,meanLocDeltaZ,axispixel,SaveFileName,centroid,range_x,range_y)
%LocTip

if isempty(stdLocDeltaZ)
    stdLocDeltaZ=zeros(size(meanLocDeltaZ));
end
centerofmass=centroid;%% option for xyz coordinate from center of mass or min "dZ"
      if centerofmass==0
          SaveFileName=['min' SaveFileName];
      else
          SaveFileName=['ctr' SaveFileName];
      end
      
clear Q1 Q LocTipsTable

if  max(max(LocTips))>0
    [L,num] = bwlabel(LocTips,4); % creates a logical matrix at the same size as islands where each island is numbered. 8 is default
else 
    return  
end

if num>0
    % num is the total number of islands.
fprintf(1,'found %g microvilli from bwlabel\n',num); 

%% in each island finds the local maxima which will be used in the fitting as the starting point
xyLocMinZ=zeros(size(LocTips));
% dilateL=zeros(size(LocTips,1));

    for n = 1:num   
               Q = 1*(L==n);  
               se90 = strel('line', 2, 90);
               se0 = strel('line', 2, 0);
               BWsdil = imdilate(Q, [se90 se0]);
              % figure, imshow(BWsdil), title('dilated gradient mask');
               
% finds all the particles inside the matrix and returns their indices in variable loc
          Q1 = meanLocDeltaZ.*BWsdil ;
%                surf(axisxnm,axisxnm,Q1,'EdgeColor','none')
%                hold on 
               Q1(Q1==0)=nan; %% remove other region(change 0 to non)
     
               [Tipminx,Tipminy]=find(Q1==min(Q1(:)));
               meanTipZvalue=Q1(Tipminx(1),Tipminy(1));
               STDvalue=stdLocDeltaZ(Tipminx(1),Tipminy(1));
               % Q2=1./(Q1);    %% invert Q1  
               Q1(isnan(Q1))=0; %% change non to 0
               area=find(Q1>0);
               areaQ1=length(area);                    
  
% sizeQ=size of filedimensionxfiledimention
              %[ind]=find(Q1==0);
              %Q1(index)=NaN;n
                           
          %     Q2 = 1./(double(Q1)./double(max(Q1(:)))); normalize;
               %% normalized and inverse
               % xcm: x center of mass; ycm: y center of mass; Qm: total mass of the island %%%%% why?? sigmay is bigger than sigmax

        [rc,cc] = ndgrid(1:size(Q1,1),1:size(Q1,2)); %!!!!!!!!!!
        Mt = sum(Q1(:));
        c1 = sum(Q1(:).* rc(:)) / Mt;
        c2 = sum(Q1(:).* cc(:)) / Mt;

        %xcm=c1
        %ycm=c2


        S1 = sum(Q1(:).* abs(c1-rc(:))) / Mt;
        S2 = sum(Q1(:).* abs(c2-cc(:))) / Mt;        

      %  fwhm= 


                    %    XYloc(round(xcm),round(ycm))=n;
                          LocTipsTable(1:10,n)=0;
                          LocTipsTable(1,n)=n; 
                          LocTipsTable(2,n)=Tipminx(1);
                          LocTipsTable(3,n)=Tipminy(1);
                          LocTipsTable(4,n)=meanTipZvalue;
                          LocTipsTable(5,n)=STDvalue;                
                          LocTipsTable(6,n)=c1;
                          LocTipsTable(7,n)=c2; 
                          LocTipsTable(8,n)=S1;
                          LocTipsTable(9,n)=S2; 
                          LocTipsTable(10,n)=areaQ1;
                          %LocTipsTable(9,n)=fwhm()

        BWsdil=BWsdil.*n;
         % xy coordinate from minpixel value 
        if centerofmass==0   
        xyLocMinZ(Tipminx,Tipminy)=num+1; 

        else
        xyLocMinZ(round(c1),round(c2))=num+1;    

        end
        dilateLocMicrovilli(:,:,n)= BWsdil;
        dilateLocMicrovilli=sum(dilateLocMicrovilli,3); %% sum is the problem ==> make it strange peak
        clear Q1 Q BWsdil

    end


end

%% create tip xy position matrix 
figure
% colormap(Jet);
% pcolor(axisxnm,axisxnm,LocMicrovilli);
imagesc(axispixel,axispixel,dilateLocMicrovilli);

set(gca,'YDir','normal')
% xlim(range_x)
% ylim(range_y)
axis equal 
%pause
hold on
xyLocMinZ(xyLocMinZ==0)=nan;
% xyLocMinZzero=xyLocMinZ.*0;
% colormap(jet(num+1))


for mol=1:size(LocTipsTable,2)
    Posy= LocTipsTable(2,mol); %% 
    Posx= LocTipsTable(3,mol);   

 scatter(Posx,Posy,20,'+','MarkerEdgeColor','b');
% text(blobCentroid(1) + labelShiftX, blobCentroid(2), num2str(mol), 'FontSize', fontSize, 'FontWeight', 'Bold');
%text(Posx+2, Posy +0.5, num2str(mol), 'FontSize', 5, 'FontWeight', 'Bold','Color',[0 0 0]); 
end
set(gca,'LooseInset',get(gca,'TightInset'))
title ('LocTips (+ min pixel)','Fontsize',18)

% xlim([range_x]);
% ylim([range_y]);
%set(gca,'YDir','normal')
             set(gca,'FontSize',14)
             xlabel('x(pixel)','FontSize',16)
             ylabel('y(pixel)','FontSize',16)
             if centerofmass==1
             title ('LocTips Centroid','FontSize',16)
             else 
                 title ('LocTips minpixel','FontSize',16)
             end
             colorbar
        save(['xyLocTip_' SaveFileName,'.mat'],'xyLocMinZ','-mat');%tip min
        saveas(gcf,['xyLocTipArea_' SaveFileName],'jpg'); 
        saveas(gcf,['xyLocTipArea_' SaveFileName],'fig');     

 hold off       
            %% create meanDeltaZ and TipLoc figure (surfc_YM)    
            
LocMicrovilli2=1*(dilateLocMicrovilli>0);
LocMicrovilli2(LocMicrovilli2==0)=nan;


figure
colormap(jet);
%     axes1 = axes('Parent',gcf);
%     view(axes1,[83 12]);
    surf(axispixel,axispixel,meanLocDeltaZ,meanLocDeltaZ) %% surf=AveZi; contour=STDZi

  %  surfc_YM2(axisxnm,axisxnm,meanLocDeltaZ,meanLocDeltaZ,meanLocDeltaZ,xyLocMinZ) %% surf=AveZi; contour=STDZi
   % axis([1 xnm(end) 1 ynm(end) 0 500]);
             xlim([range_x]);ylim([range_y]);
             set(gca,'FontSize',12)
             title( ['Reconstructed microvilli'],'FontSize',16)
             xlabel('x(pixel)','FontSize',16)
             ylabel('y(pixel)','FontSize',16)
             zlabel('z(nm) Mean','FontSize',16)
    saveas(gcf,['ReconMicrovilli_' SaveFileName],'jpg'); 
    saveas(gcf,['ReconMicrovilli_' SaveFileName],'fig');        

    save(['LocTipsTable' SaveFileName,'.mat'],'LocTipsTable','-mat');


    
    
    

end
```
