## Supplementary material for "Microvillar cartography: a super-resolution single-molecule imaging method to map the positions of membrane proteins with respect to cellular surface topography": Analysis MATLAB codes: Modnlparci_ver19.m

```
function [Delta] = Modnlparci_ver19(beta,resid,varargin)
%NLPARCI Confidence intervals for parameters in nonlinear regression.
%   CI = NLPARCI(BETA,RESID,'covar',SIGMA) returns 95% confidence intervals
%   CI for the nonlinear least squares parameter estimates BETA.   Before
%   calling NLPARCI, use NLINFIT to fit a nonlinear regression model
%   and get the coefficient estimates BETA, residuals RESID, and estimated
%   coefficient covariance matrix SIGMA.
%
%   CI = NLPARCI(BETA,RESID,'jacobian',J) is an alternative syntax that
%   also computes 95% confidence intervals.  J is the Jacobian computed by
%   NLINFIT.  You should use the 'covar' input rather than the 'jacobian'
%   input if you use a robust option with NLINFIT, because the SIGMA
%   parameter is required to take the robust fitting into account.
%
%   CI = NLPARCI(...,'alpha',ALPHA) returns 100(1-ALPHA) percent
%   confidence intervals.
%
%   NLPARCI treats NaNs in RESID or J as missing values, and ignores the
%   corresponding observations.
%
%   The confidence interval calculation is valid when the length of RESID
%   exceeds the length of BETA, and J has full column rank.  When J is
%   ill-conditioned, confidence intervals may be inaccurate.
%
%   Example:
%      load reaction;
%      [beta,resid,J,Sigma] = nlinfit(reactants,rate,@hougen,beta);
%      ci = nlparci(beta,resid,'covar',Sigma);
%
%   See also NLINFIT, NLPREDCI, NLINTOOL.

% Old syntax still supported:
%    CI = NLPARCI(BETA,RESID,J,ALPHA)
    
%   To compute confidence intervals when the parameters or data are complex,
%   you will need to split the problem into its real and imaginary parts.
%   First, define your parameter vector BETA as the concatenation of the real
%   and imaginary parts of the original parameter vector.  Then concatenate the
%   real and imaginary parts of the response vector Y as a single vector.
%   Finally, modify your model function MODELFUN to accept X and the purely
%   real parameter vector, and return a concatenation of the real and
%   imaginary parts of the fitted values.  Given this formulation of the
%   problem, NLINFIT will compute purely real estimates, and confidence
%   intervals are feasible.

%   References:
%      [1] Seber, G.A.F, and Wild, C.J. (1989) Nonlinear Regression, Wiley.

%   Copyright 1993-2009 The MathWorks, Inc. 
%   $Revision: 1.1.8.1 $  $Date: 2010/03/16 00:16:24 $

J = [];
Sigma = [];
alpha = 0.05;
if nargin>=3 && ischar(varargin{1})
   % Calling sequence with named arguments
   okargs =   {'jacobian' 'covariance' 'alpha'};
   defaults = {[]         []           0.05};
   [eid emsg J Sigma alpha] = internal.stats.getargs(okargs,defaults,varargin{:});
   if ~isempty(eid)
      error(sprintf('stats:nlparci:%s',eid),emsg);
   end
else
   % CI = NLPARCI(BETA,RESID,J,ALPHA)
   if nargin>=3, J = varargin{1}; end
   if nargin>=4, alpha = varargin{2}; end
end
if nargin<=2 || isempty(resid) || (isempty(J) && isempty(Sigma))
   error('stats:nlparci:TooFewInputs',...
         'Requires BETA, RESID, and either J or SIGMA.');
end;
if ~isreal(beta) || ~isreal(J)
    error('stats:nlparci:ComplexParams',...
         ['Cannot compute confidence intervals for complex parameters.  You must\n' ...
          'reparameterize the model into its real and imaginary parts.']);
end
if isempty(alpha)
    alpha = 0.05;
elseif ~isscalar(alpha) || ~isnumeric(alpha) || alpha<=0 || alpha >= 1
    error('stats:nlparci:BadAlpha',...
          'ALPHA must be a scalar between 0 and 1.');
end

% Remove missing values.
resid = resid(:);
missing = isnan(resid);
if ~isempty(missing)
    resid(missing) = [];
end
n = length(resid);
p = numel(beta);
v = n-p;

if ~isempty(Sigma)
   se = sqrt(diag(Sigma));
else
   % Estimate covariance from J and residuals
   J(missing,:) = [];
   if size(J,1)~=n || size(J,2)~=p
      error('stats:nlparci:InputSizeMismatch',...
            'The size of J does not match the sizes of BETA and RESID.');
   end

   % Approximation when a column is zero vector
   temp = find(max(abs(J)) == 0);
   if ~isempty(temp)
      J(:,temp) = sqrt(eps(class(J)));
   end

   % Calculate covariance matrix
   [emp,R] = qr(J,0);   % modify from ~ to emp for matlab7
   Rinv = R\eye(size(R));
   diag_info = sum(Rinv.*Rinv,2);

   rmse = norm(resid) / sqrt(v);
   se = sqrt(diag_info) * rmse;
end

% Calculate confidence interval
Delta = se * tinv(1-alpha/2,v);
% ci = [(beta(:) - delta) (beta(:) + delta)];
```
