## Supplementary material for "Microvillar cartography: a super-resolution single-molecule imaging method to map the positions of membrane proteins with respect to cellular surface topography": Analysis MATLAB codes: mygauss2D_v19.m

```
function[H]= mygauss2D_v19(p,xy_dat) 

H = p(1)*(exp(-0.5*((xy_dat(:,2)-p(2))).^2./(p(4)^2)-0.5*((xy_dat(:,1)-p(3))).^2./(p(5)^2))) + p(6);

% Creates a 2D gaussian.
% Inputs 
% X,Y  - x and y values created using the meshgrid function
% p - a list of parameter 
%amp = p(1);
%cx = p(2); 
%cy = p(3);
%wx = p(4);
%wy = p(5);
%bg = p(6)


%H =(p(1)*exp(-0.5*((xy_dat(:,1)-p(2))/w1)^2-0.5*((y-yc)/w2)^2))+p(6);

%ztmp = p(5)*(exp(-0.5*(X-p(1)).^2./(p(3)^2)-0.5*(Y-p(2)).^2./(p(4)^2))) - m;

%z = sum(sum(ztmp.^2));
```
