## Supplementary material for "Microvillar cartography: a super-resolution single-molecule imaging method to map the positions of membrane proteins with respect to cellular surface topography": Analysis MATLAB codes: mygauss3.m

```
function H = mygauss3(p,xy_dat)
% Creates a 2D gaussian.
% Inputs - 
% X,Y  - x and y values created using the meshgrid function
% p - a list of parameter : amplitude, x-center, y-center, width,
% background

H = p(1)*exp( - ((xy_dat(:,1)-p(2)).^2 + (xy_dat(:,2)-p(3)).^2)./(2*p(4)^2))+p(5) ;
end
```
