## Supplementary material for "Microvillar cartography: a super-resolution single-molecule imaging method to map the positions of membrane proteins with respect to cellular surface topography": Analysis MATLAB codes: NumPho_2plot_color.m

```
%% Display the reconstruction images with 3D or 2D color map for number of photons. 
%% Choose the "Particle Table" (usually FITsetPT)
%%%% choose the range to make color figure by setting "on=1 off=0) by Number of photons of each molecules
%%%% change colormap type
%%%% save the sorted particle table as a cell type. 
%%%% Right range of intensity 
function[ColorFigure]=NumPho_2plot_color(myColorMap,StormData1,StormData2,StormFileName,miniPN,maxiPN,range_x,range_y,markersize)
% Resolsize=5;
% Framesize=[256,256]; %% frame demention

% %Colortype='jet';%%% Hot, Jet, Autumn, Cool, Gray %% single color = red, blue
% RangeSetting=[1 1 1 1 1 1 1 1 1 1];% [1 1 1 1 1 1 1 1 1 1] for Low threshold to 10000
% % miniPN =1500;
% % maxiPN =10000;

%     FileName=StormFileName1;
%     SaveColorFileName = ['Color',Colortype ,num2str(maxiPN),FileName(1:end-4)]; 
   
%     if length(SaveColorFileName)>60 || length(SaveColorFileName)>60 
%     SaveColorFileName = [SaveColorFileName(1:40),SaveColorFileName(end-10:end-4)];% type char
%  
%     end

StormData1(:,10)=roundn(StormData1(:,10),2);
StormData2(:,10)=roundn(StormData2(:,10),2);

for kk=1:size(StormData1,1)
if StormData1(kk,10)<= miniPN;
StormData1(kk,10)=miniPN;
elseif StormData1(kk,10)>= maxiPN;
StormData1(kk,10)=maxiPN;
end
end


for kk=1:size(StormData2,1)
if StormData2(kk,10)<= miniPN;
StormData2(kk,10)=miniPN;
elseif StormData2(kk,10)>= maxiPN;
StormData2(kk,10)=maxiPN;
end
end


ColorFigure = figure;
set(gcf, 'Position', [10 20 2000 900]);
colormap(myColorMap)

axis equal square
hold all

subplot(1, 2, 1);
scatter(StormData1(:,3), StormData1(:,2), 0.5, StormData1(:,10),'*')%,filled);
axis equal square
title(['focalplane: 0nm'],'Fontsize',18);
set(gca,'LooseInset',get(gca,'TightInset'))
addscale3_um_range_label(range_x,range_y,markersize,0)
colorbar
caxis ([miniPN,maxiPN]) 

subplot(1, 2, 2);
scatter(StormData2(:,3), StormData2(:,2), 0.5, StormData2(:,10),'*')%,filled);
axis equal square
title(['focalplane: -400nm'],'Fontsize',18);
set(gca,'LooseInset',get(gca,'TightInset'))
addscale3_um_range_label(range_x,range_y,markersize,0)
colorbar
caxis ([miniPN,maxiPN]) 
%caxis ([miniPN,maxiPN]) 

SaveName=[num2str(miniPN),'-',num2str(maxiPN),StormFileName(1:end-4)];
% 
if length(SaveName)>60 
    SaveName = [SaveName(1:60)];% type char
end
saveas(gcf,[SaveName]);%,'jpg');    
%set(gcf, 'Position', get(0, 'ScreenSize'));
saveas(gcf,[SaveName],'jpg');    
% % 

%close 
 end
```
