## Supplementary material for "Microvillar cartography: a super-resolution single-molecule imaging method to map the positions of membrane proteins with respect to cellular surface topography": Analysis MATLAB codes: OpenVAfigures_Roi.m

```
%% Open fits file or Tif file for VA calibration 
%ver4: option added for bacground substraction (edge intensity) 
%Make a stack of files of images at different angle;  
% input : FNlist, Path,ws,xmax,ymax, allws
%         read a file ==> makes 1 frame (mean value)
%       
% output: igstack; GetVA (eg:897 get the scale of micrometer of filename)
% option: #allws=0--> stacks of (wsXws)size of series of movies 
%                     and the centerpixel(xmax,ymax)of original movie
%    
%         #allws =1--> ws= size(igstack)  
function[igstack,filename,NumofFiles,Bacground]= OpenVAfigures_Roi(FNlist,Path,ws,subbac,subbacmaxangle,ch,range_x,range_y)

ws=range_x(2)-range_x(1)+1;%% must be odd number

if   iscell(FNlist)
     NumofFiles=size(FNlist,2);
else
     NumofFiles=1;
end


for ff = 1:NumofFiles;

    if  iscell(FNlist);    
        filename=FNlist{ff};
    else 
        filename=FNlist;
    end


    if strcmp(filename(end-3:end),'fits')
        AA = fitsread([Path, filename]);
    else
        strcmp(filename(end-2:end),'tif')
        AA = imread([Path, filename]);
    end

       %% All frames (make single frame from a movie) 
    AA=double(AA); 

    if length(size(AA))<3
        Ig=double(AA); 
    else 
        Ig=mean(AA,3);
    end
% choose channel

    
    if ch==2
    Ig=Ig(1:(size(Ig,1))/2,1:end);
    elseif ch==1
    Ig=Ig(size(Ig,1)/2+1:end,1:end);
    end

    %%%# Read an image    
    imagesc(Ig)

    % figure
    % pcolor(Ig);
    %h=pcolor(Ig);
    % set(h,'EdgeColor','none')
    axis equal
    xlim(range_x);
    ylim(range_y);
    set(gca,'YDir','normal')
    % surf(Ig(1:size(Ig,1),1:size(Ig,2)));
  %  pause
    close

 %   if allws==1
    igstack(:,:,ff)=Ig;
%    ws=size(igstack,1);

%     else
%     pn = (ws-1)/2;
%    igstack(:,:,ff)=Ig(range_x(1):range_x(2),range_y(1):range_y(2));%=1;
%     % %Q= AA.*Q; 
%                %Qw=Qw(xmax-pn:xmax+pn,ymax-pn:ymax+pn); %size changed
%     %            figure
%     %            contour(Qw,'DisplayName','Qw');
%     %            colorbar;axis equal;
% %     ws=ws;
%     end
%     
            
    %% GetVA
   % getparts = strread(filename,'%s','delimiter','_');
   % getva=getparts(end-1);
   % token = strtok(getva, 'va');
   % getva=str2double(token);
   % getVA(ff)=getva;    
    
end
    %% background substraction option 
if subbac==0
     for nn=1:size(igstack,3)
            
            Qb=igstack(:,:,nn);
            Qb(2:ws-1,2:ws-1)=0;% extanded a logical matrix as a size of ws (13x13)
            ind=find(Qb>0);
            Bacground=mean(Qb(ind));
                       %    contour(Qb,'DisplayName','Qb');figure(gcf)
     end
    
elseif subbac==1 && subbacmaxangle==0   
        
        for nn=1:size(igstack,3)
            
            Qb=igstack(:,:,nn);
            Qb(2:ws-1,2:ws-1)=0;% extanded a logical matrix as a size of ws (13x13)
            ind=find(Qb>0);
            Bacground=mean(Qb(ind));
                       %    contour(Qb,'DisplayName','Qb');figure(gcf)
            igstack(:,:,nn)=igstack(:,:,nn)-Bacground;
        end
elseif subbac==1 && subbacmaxangle==1     
         if getVA(1)<getVA(end)
            Qb=igstack(:,:,end);
         else 
             error('bagmax')
         end
            Qb(2:ws-1,2:ws-1)=0;% extanded a logical matrix as a size of ws (13x13)
            ind=find(Qb>0);
            Bacground=mean(Qb(ind))
            
        for nn=1:size(igstack,3)
                               %    contour(Qb,'DisplayName','Qb');figure(gcf)
           igstack(:,:,nn)=igstack(:,:,nn)-Bacground;
        end
end
           


end
```
