## Supplementary material for "Microvillar cartography: a super-resolution single-molecule imaging method to map the positions of membrane proteins with respect to cellular surface topography": Analysis MATLAB codes: Reconstruction_ver9.m

```
function [CMfig,Fitfig,MerFig]= Reconstruction_ver9(CMPT,FitPT,Parameter,SaveName,RecPix)
%Reconstruction_v27(RecPix,CMoutPT,FitoutPT,FileDimention,ReconCM,ReconFit,SaveFileName)
% v27, CM is blue, Fit is red

if  nargin <6
    RecPix   = Parameter.RecPix;
elseif nargin <5
    
    if Parameter.ApplyFilter==0
        SaveName = Parameter.SaveFileName;
    elseif Parameter.ApplyFilter==1
    SaveName = [Parameter.FilterType, '_', Parameter.SaveFileName];
    else 
        error('Set ApplyFilter option')
    end
    
    RecPix   = Parameter.RecPix;
  
    
end

if isempty(SaveName)
    SaveName = input ('save as: ');
end

if isempty(RecPix)
    RecPix = input('Rendering size (nm) for Reconstruction :');
end
    
FileDimention = Parameter.FileDimention;
Magnify       = 240   ;    % Magnification of system : 240x 
Pixsize       = 16000  ;    % Camera CCD pixel size in nm
Pixelnm       = round((Pixsize/Magnify)*100)/100  ; %66.6700nm  
Rend          = (round(RecPix/Pixelnm*100))/100;     % Rend: rendering pixels for desired size (example 1pixel is 66.67nm, 66,67/Rend=desired pixel size(nm))  

if size (CMPT,1)>0 
            CMfig=figure('Name','CM_Reconstruction');
            CM=CMPT(:,2);
            CN=CMPT(:,3);
            %axis equal
            plot(CN,CM,'.','MarkerSize',Rend);  %% Don't know why 
            axis equal
            xlim([0 FileDimention(2)]);
            ylim([0 FileDimention(1)]); 
            %xlim([1 FileDimention(1)]);
            %ylim([1 FileDimention(2)]);
            
            if length(SaveName)>51
                dd =(length(SaveName)-50);
                SaveName=SaveName(dd:end);       
            end
            
                figuresavenamecm= ['CMrec',num2str(RecPix),'nm_',SaveName];
                saveas(CMfig,figuresavenamecm);%,'jpg');
                saveas(CMfig,figuresavenamecm,'tif');            

            %  saveas(CMfig,['CMrec',num2str(RecPix),'nm_',SaveName],'tif'); 

        
end

if size (FitPT,1)>0 
             Fitfig=figure('Name','Fit_Reconstruction');
             FM=FitPT(:,2);
             FN=FitPT(:,3);
             %axis equal
             plot(FN,FM,'.','MarkerSize',Rend,'Color','red');   %% Don't know why 
             axis equal
             xlim([0 FileDimention(2)]);
             ylim([0 FileDimention(1)]); 
%             xlim([1 FileDimention(1)]);
%             ylim([1 FileDimention(2)]);   
             
             if length(SaveName)>51
                dd =(length(SaveName)-50);
                SaveName= SaveName(dd:end); 
             end
             figuresavenamefit= ['Fitrec',num2str(RecPix),'nm_',SaveName];
             saveas(Fitfig,figuresavenamefit);%,'jpg');
             saveas(Fitfig,['Fitrec',num2str(RecPix),'nm_',SaveName],'tif'); 
             %  %            saveas(figure(gcf),figuresavenamefit);%,'jpg');
%              saveas(figure(gcf),['Fitrec',num2str(RecPix),'nm_',SaveName],'tif'); 
             % saveas(figure(gcf),['Fitrec',num2str(RecPix),'nm_',SaveName,]);%,'jpg');
end

if size (FitPT,1)>0 && size(CMPT,1)>0 
 MerFig =figure('Name','CMFit_mergeReconstruction');
             CM=CMPT(:,2);
             CN=CMPT(:,3);
             plot(CN,CM,'.','MarkerSize',Rend); 
             axis equal
             xlim([0 FileDimention(2)]);
             ylim([0 FileDimention(1)]);              
            % xlim([1 FileDimention(1)]);
            % ylim([1 FileDimention(2)]); 
             hold on   
             FM=FitPT(:,2);
             FN=FitPT(:,3);
             %axis equal
             plot(FN,FM,'.','MarkerSize',Rend,'Color','red');   
%              xlim([1 FileDimention(1)]);
%              ylim([1 FileDimention(2)]);   
             
             if length(SaveName)>51
                dd =(length(SaveName)-50);
                SaveName=SaveName(dd:end);  
             end
             figuresavenamefit= ['Merge',num2str(RecPix),'nm_',SaveName];
             saveas(MerFig,figuresavenamefit);%,'jpg');
             saveas(MerFig,['Merge',num2str(RecPix),'nm_',SaveName],'tif'); 
    
end 

end


    
%         if size (CMoutPT,1)>0 && ReconFit==1 && size (FitoutPT,1)>0
%     figure('Name','CM_Reconstruction')
%     M=CMoutPT(:,2);
%     N=CMoutPT(:,3);
%     %axis equal
%     CMrec=plot(N,M,'.','MarkerSize',Rend);  %% Don't know why 
%     xlim([1 FileDimention(1)]);
%     ylim([1 FileDimention(2)]);

% elseif ReconCM==1 &&size (CMoutPT,1)>0 && ReconFit==0 && size(FitoutPT,1)==0
%     figure('Name','CM_Reconstruction')
%     M=CMoutPT(:,2);
%     N=CMoutPT(:,3);
%     %axis equal
%     CMrec=plot(N,M,'.','MarkerSize',Rend);  %% Don't know why 
%     xlim([1 FileDimention(1)]);
%     ylim([1 FileDimention(2)]);
%     
% elseif ReconCM==0 && size (CMoutPT,1)==0 && ReconFit==1 &&size(FitoutPT,1)>0
% 
%     % Rend: rendering pixels for desired size (example 1pixel is 66.67nm, 66,67/Rend=desired pixel size(nm))  
%      figure('Name','2DFit_Reconstruction')
%      M=FitoutPT(:,2);
%      N=FitoutPT(:,3);
%      %axis equal
%      Fitrec=plot(N,M,'.','MarkerSize',Rend);  %% Don't know why 
%      xlim([1 FileDimention(1)]);
%      ylim([1 FileDimention(2)]);   
%       
% 
%     end
```
