## Supplementary material for "Microvillar cartography: a super-resolution single-molecule imaging method to map the positions of membrane proteins with respect to cellular surface topography": Analysis MATLAB codes: Result_NormXcorr2_shift_v5_Range.m

```
function [ShiftResult,cc_Qws]=Result_NormXcorr2_shift_v5_Range(Template,RefImage_Range,Filefolder,FileNameHead,lastframe,ApplyfilterCC,kernelsizeCC,getfocalplane,x_range,y_range)
%%% input (ParticleNum in one frame,framex,framey,nframes)
%   eg: SimulateSM_YM_celltype(10,100,100,5)
%%% output (Original_Loc Table)
%%% input  (SimImage: cell type simmulate images)
%%% v1: change the result file order; normxcoor2%% %% example [Original_Loc,SimImage] = SimulateSM_YM_celltype_shift(1,11,11,5,0.2,0);
%MovieName=input('Saving FileName??');
%function [Original_Loc,SimImage,Result] = SimulateSM_YM_celltype_shift(PhotonNum,ParticleNum,framex,framey,nframes,offset_yshift,offset_xshift,ws)
%option=2;% 1: use the center position from the max of template
%option=2;% 2: use the center position from the max of offset

%option_newpeak=0; %% newpeak==1: use newfind peak, 0== use ori_peaks
%sizepixel=66.67;
% v3: normxcorr2 template is larger size
%close all; clear all; 
% v4: normxcorr2: template and offset_Template is the same size


%% Pick DTable
% [FileName,Motherfolder] = uigetfile( ...
% {  '*.mat','MAT-files (*.mat)'; ...
%    '*.*',  'All Files (*.*)'}, ...
%    'Pick a file')%, ...
%   % 'MultiSelect', 'on');
% 
% cd(Motherfolder)
% 
% SimImage=importdata(FileName);


%% v2: xcoor2
%% load SimImage ;
%SimImage=()
%clearvars -except SimImage

% TestMovie=SimImage; 

%makeSimImagetoTif=0;
%getfocalplane=num2str(getfocalplane);
makeCCImagetoTif=1;
Qshow=1;
saveQ=1;
analysistypename=['NormXcorr2fit'];
%Applyfilter=1   ;      
filtertype='MeanFilter' ;% 'MedianFilter' 'LogFilter';'GaussFilter'
%kernelsize=5; 
filtersigma=10;

%Motherfolder = uigetdir;
%MotherPath=[Motherfolder '\'];

%cd (Motherfolder)

OptionNewpeak=0;

if OptionNewpeak==1
SaveName=([analysistypename]);
elseif OptionNewpeak==0
SaveName=([analysistypename]);
end

if ApplyfilterCC==1
SaveName=([SaveName,'_',filtertype,'_', num2str(kernelsizeCC)]);
elseif OptionNewpeak==0
SaveName=([SaveName,'_',filtertype,'_', num2str(kernelsizeCC)]);
end


if iscell(RefImage_Range)
nframes=size(RefImage_Range,2);
else
nframes=size(RefImage_Range,3);    
end

offset_xshift=0;
offset_yshift=0;
ws=11; %% window size for fitting 


SaveFolderName=[FileNameHead,SaveName,'_ws',num2str(ws),'_',getfocalplane];
if (exist (SaveFolderName)==0);
    mkdir(pwd,SaveFolderName)
else (exist(SaveFolderName)==1);
    SaveNewName=input('NewName??(Y==>1,N==>0):')
    if SaveNewName==1;
    SaveFolderName=input('SaveFolderName?? : ','s'); 
    end
mkdir(pwd,SaveFolderName)
end
cd (SaveFolderName);  


% draw a figure
% if makeSimImagetoTif ==1
% for k=1:nframes
%     h=(imshow(SimImage{k}));
%     Simimagetest = getimage(h);
%     cmap = colormap('gray');
%     Simimagetest=uint16(Simimagetest);
%     imwrite(Simimagetest,cmap,['FrameImage_',num2str(k),'.tif']);
% end

% 
if lastframe==0
Ref_Frame=RefImage_Range{1};
elseif lastframe==1
Ref_Frame=RefImage_Range{end};
else
    error('check the lastframe option')
end
%autocorrelation find the original peak (Reference position) 
%ca = normxcorr2(Template,Template); 

  ca = normxcorr2(Ref_Frame,Template); 
  %maxvalue =  max(abs(cc(:)))
%  [max_ca, imax] = max(abs(ca(:)));
% range_x=size(Ref_Frame,1);
% range_y=size(Ref_Frame,2);
 [max_ca]= max(ca(:));
% [max_ca]= max(max(ca(y_range(2)-20:y_range(2)+20,x_range(2)-20:x_range(2)+20)));

 % [ypeak, xpeak] = ind2sub(size(cc),imax(1));
 [ori_ypeak, ori_xpeak] =find(ca==max_ca);
  

 
 %[ori_ypeak, ori_xpeak] = ind2sub(size(ca),imax(1));


for k=1:nframes 

Off_Template=RefImage_Range{k};

%Off_Template=Off_Template(15:end-15,15:end-15);
% CC{k-1}=xcorr2(Templete,Off_Templete);

%   cc = normxcorr2(Off_Template,LargeTemplate); 
   cc = normxcorr2(Off_Template,Template); 
%cc = xcorr2(Off_Template,Template); %% for xcoor2 the order is different
  %maxvalue =  max(abs(cc(:)))
   %% Meanfilter substract 
         if ApplyfilterCC==1
             
            cc=FilterFrameOut_ws(cc,filtertype,kernelsizeCC,filtersigma);
         end 
  
 % [max_cc, imax] = max(abs(cc(:)));
 
 %[max_cc, imax] = max(abs(cc(:))) %find max from all frames
  [max_cc]= max(cc(:));

% [max_cc]= max(max(cc(y_range(2)-20:y_range(2)+20,x_range(2)-20:x_range(2)+20)));

 % [ypeak, xpeak] = ind2sub(size(cc),imax(1));
  [ypeak, xpeak] =find(cc==max_cc);
%   
%   xpeak
%   ypeak
  reference_y=-(k-1)*offset_yshift;
  reference_x=-(k-1)*offset_xshift;

%auto_c= normxcorr2(Off_Template,Off_Template);


% Off_LargeTemplate=PoissNoise;
% 
% %LargeTemplate= random('poisson',TemplateBgd);
% 
% xrange=[(size(LargeTemplate,1)-size(Template,1)-1)/2+1,((size(LargeTemplate,1)-size(Template,1)-1)/2)+size(Template,1)];
% yrange=[(size(LargeTemplate,2)-size(Template,2)-1)/2+1,((size(LargeTemplate,2)-size(Template,2)-1)/2)+size(Template,2)];
% 
% 
% 
% Off_LargeTemplate(xrange(1):xrange(2),yrange(1):yrange(2))=Off_Template(1:end,1:end);
% %imagesc(Off_LargeTemplate)
% %pause

auto_c= normxcorr2(Off_Template,Off_Template); 
% auto_c= xcorr2(Off_Template,Off_Template);
 [max_auto, imaxauto] = max(abs(auto_c(:)));
 [auto_ypeak, auto_xpeak] = ind2sub(size(auto_c),imaxauto(1));
 
 
 
 

%% generate figure for cc
if makeCCImagetoTif==1;
figure
%colormap(gray)
%h=imagesc(cc);
%axis equal tight 
CCFrame=flipud(cc);
h = imshow(CCFrame,[]);
%cmap = colormap('gray');
CCtest = getimage(h);
%imwrite(CCtest,cmap,['Image_frame',num2str(k),'.tif']);
%saveas(h,['CC_Frame',num2str(k)],'tif')
imwrite(CCtest,['CC_Frame',num2str(k),'.tif']);
%pause

%  h=(imshow(SimImage{k},[]));
%     Simimagetest = getimage(h);
%     cmap = colormap('gray');
%     Simimagetest=uint16(Simimagetest);
%     imwrite(Simimagetest,cmap,['SimImage_frame',num2str(k),'.tif']);
close
end

%% calculate the Peak nearby the center position;
if OptionNewpeak==0 % original use

Ypeak=ori_ypeak;
Xpeak=ori_xpeak;

elseif OptionNewpeak==1
    if abs(ypeak-ori_ypeak)<2
       Ypeak=ori_ypeak;
%     elseif abs(ypeak-ori_ypeak)>10 || abs(xpeak-ori_xpeak)>10
%         
%        [Ypeak Xpeak]=max(Ypeak-10:Ypeak+10,:);
    else 
       Ypeak=ypeak; 
    end

    if abs(xpeak-ori_xpeak)<2
       Xpeak=ori_xpeak;
    else
       Xpeak=xpeak;
    end
end
%cc=cc';
pn=(ws-1)/2;
%Q=cc(Ypeak-pn:Ypeak+pn,Xpeak-pn:Xpeak+pn);version 2
Q=cc(Ypeak-pn:Ypeak+pn,Xpeak-pn:Xpeak+pn);%version 2
cc_Qws{k}=Q;
if Qshow==1
colormap(hot);imagesc(Q)
set(gca,'YDir','normal');
%pause; %%%%%%%%%%%%%%%%%%%%%%%%%%%%%%editted 150825
close 
end
%Q'=Q;
[bb,ci] = fit2dgaussian_4YM(Q,[(ws+1)/2;(ws+1)/2],ws);
% yshift= Ypeak-bb(3)%-Ypeak; % shifted as :0.5
% xshift= Xpeak-bb(2)%-Xpeak;  % shifted as : +0.2
%bb
%pause
% x_fit=bb(2)-((ws+1)/2)+Xpeak -xrange(1)+1;%-ori_xpeak)+ori; % fitting Result after center pixel correction
% y_fit=bb(3)-((ws+1)/2)+Ypeak -yrange(1)+1;%-ori_ypeak); % fitting Result after center pixel correction

 x_fit=bb(2)-((ws+1)/2)+Xpeak %-ori_xpeak)+ori; % fitting Result after center pixel correction
 y_fit=bb(3)-((ws+1)/2)+Ypeak %-ori_ypeak); % fitting Result after center pixel correction


CCall{k}=cc;
BBall{k}=bb;

%pause

delta_xshift=reference_x+ori_xpeak-x_fit;
delta_yshift=reference_y+ori_ypeak-y_fit;
%pause

%auto_c=auto_c;
pn=(ws-1)/2;
autoQ=auto_c(auto_ypeak-pn:auto_ypeak+pn,auto_xpeak-pn:auto_xpeak+pn);
auto_Qws{k}=autoQ;
if Qshow==1
colormap(jet);imagesc(autoQ);
set(gca,'YDir','normal');
%pause;
close;
end

[autobb,ci] = fit2dgaussian_4YM(autoQ,[(ws+1)/2;(ws+1)/2],ws);
auto_x=autobb(2)-((ws+1)/2);
auto_y=autobb(3)-((ws+1)/2);

%Result(k,:)=[ypeak,xpeak,max_cc,reference_x,reference_y,xshift,yshift,reference_x-xshift,reference_y-yshift];
%Result(k,:)=[ypeak,xpeak,max_cc,reference_y,reference_x,yshift,xshift,delta_yshift,delta_xshift,auto_y,auto_x];
Result_Fit(k,:)=[reference_x,reference_y,x_fit,y_fit,(reference_x+ori_xpeak),(reference_y+ori_ypeak),delta_xshift,delta_yshift,auto_x,auto_y];

end
%Result_Fit=-Result_Fit;
ShiftResult=Result_Fit;
if saveQ==1
[h]=makeimagefile(cc_Qws,'gray','cc_Qws');
%[h]=makeimagefile(ccQ_all,'gray','RefImage_Range');
end

%% Result Figure
figure;
subplot(2,1,1)
plot(1:nframes,Result_Fit(:,7),'--*b'); hold on
%plot(1:nframes,Result_Fit(:,8)+ori_xpeak,'b'); hold on
ylabel('X (pixel)','fontsize',16);
%xlabel('frame(nth)','fontsize',16);
title([analysistypename]);
subplot(2,1,2)
plot(1:nframes,Result_Fit(:,8),'--*r');hold on
%plot(1:nframes,Result_Fit(:,6)+ori_ypeak,'r'); hold on
ylabel('Y (pixel)','fontsize',16);
xlabel('Movie(nth)','fontsize',16);
title([analysistypename]);%,' shifty:',num2str(offset_yshift)]);
save(['NormXcorr2_Result2D_Fit'],'Result_Fit');

saveas(gcf,['NormXcorr2_Result2D_Fit'])
saveas(gcf,['NormXcorr2_Result2D_Fit'],'bmp')
%analysistypename='Xcorr2 2DFit';
%deltashift(Result_Fit,shiftx,shifty,sx,sy,amp,snr,analysistypename)
%deltashift_fit(Result_Fit,offset_xshift,offset_yshift,analysistypename)

%% compare autocorrelation

figure;
% subplot(1,2,1)
AutoXnm=Result_Fit(:,9)*66.67;
AutoYnm=Result_Fit(:,10)*66.67;
plot(1:nframes,AutoXnm,'-*k'); hold on
plot(1:nframes,AutoYnm,'-ok'); hold on
% legend('X','Y')
% legend('boxoff')
hline = refline([0 0]);
set(hline,'Color','k')
ylabel('shift(nm)','fontsize',16);
xlabel('frame(nth)','fontsize',16);
title(['Autoxcoor2 2Dfit']);%; shiftx:',num2str(offset_xshift),' shiftx:',num2str(offset_yshift)]);
% subplot(1,2,2)
% plot(Result_Fit_nm(:,1),Result_Fit_nm(:,10),'-*k');hold on
% plot(Result_Fit_nm(:,2),Result_Fit_nm(:,6),'k'); hold on
% ylabel('nm','fontsize',16);
% xlabel('shift(pixel)','fontsize',16);
% title(['NormXcorr2_2DFit shifty:',num2str(offset_yshift)]);
% save(['Xcorr2_Result2D_Fit'],'Result_Fit');
saveas(gcf,['NormXcorr2_Result2D_Auto'])
saveas(gcf,['NormXcorr2_Result2D_Auto'],'bmp')
close; 

save(['NormXcoor2_Result_Fit_pix',SaveName],'Result_Fit');
save(['Xcorr2_Result_Fit_pix_',SaveName],'Result_Fit','-ascii');
Result_Fit_nm=Result_Fit*66.67;
save(['NormXcorr2_Result_Fit_nm_',SaveName],'Result_Fit_nm');
save(['NormXcorr2_Result_Fit_nm_',SaveName],'Result_Fit_nm','-ascii');
%elseif option==2
%save(['SimResultSummary2_',num2str(PhotonNum),'_',num2str(ParticleNum),'_',num2str(framex),'_',num2str(framey),'_',num2str(nframes)],'Result2');
%end
save(['CCall_',SaveName],'CCall');
save(['BBall_',SaveName],'BBall'); 
save(['cc_Qws_',SaveName],'cc_Qws');
save(['auto_Qws_',SaveName],'auto_Qws');


%% ParameterSummary
% Sim_Parameter.MoiveName=SaveName;
% Sim_Parameter.SaveFolderName=SaveFolderName;
% Sim_Parameter.PhotonNum=PhotonNum;
% Sim_Parameter.ParticleNum=ParticleNum;
% Sim_Parameter.Sigma_x=Sigma_x;
% Sim_Parameter.Sigma_y=Sigma_y;
% Sim_Parameter.framex=framex;
% Sim_Parameter.framey=framey;
% Sim_Parameter.offset_yshift=offset_yshift;
% Sim_Parameter.offset_xshift=offset_xshift;
% Sim_Parameter.ws=ws;
% Sim_Parameter.AddNoise=AddNoise;
% Sim_Parameter.nmovie=nmovie;
% Sim_Parameter.fixedcenterposition=fixedcenterposition;
% Sim_Parameter.MeanPixelBgd=MeanPixelBgd;
% Sim_Parameter.EMgainofCCD=EMgainofCCD;
% Sim_Parameter.QE=QE;
% Sim_Parameter.Sensitivity=Sensitivity;
% % Sim_Parameter.pixelsizenm=pixelsizenm;
% 
% save(['ParameterSummary'],'Sim_Parameter');
% save(['CCall_',num2str(PhotonNum),'_',num2str(ParticleNum),'_',num2str(framex),'_',num2str(framey),'_',num2str(nframes)],'CCall_2');
% save(['BBall_',num2str(PhotonNum),'_',num2str(ParticleNum),'_',num2str(framex),'_',num2str(framey),'_',num2str(nframes)],'BBall_2')
%save(['SimOriginalLoc_',num2str(PhotonNum),'_',num2str(ParticleNum),'_',num2str(framex),'_',num2str(framey),'_',num2str(nframes)],'ccAll');
cd (Filefolder)
%close all

end
 %end
%end
%%% How to use
%%% Particle_Loc is the list of the x,y corrdiates of the particle 
%%% try to change the relevant background if you want to simulate the more
%%% similar situation. Random is for the possion random background
%%% %%%[Images,Particle_Loc] = SimulateParticleField(10,5)
%%%surf(Images{1})
```
