## Supplementary material for "Microvillar cartography: a super-resolution single-molecule imaging method to map the positions of membrane proteins with respect to cellular surface topography": Analysis MATLAB codes: Result_NormXcorr2_shift_v6_Range_ChannelComp_2MeanMovie.m

```
% modified on 14 March 2016 by YM 
% original Mfile name: Result_NormXcorr2_shift_v6_Range_ChannelComp_2MeanMovie.m


function [ShiftResult,cc_Qws]=Result_NormXcorr2_shift_v6_Range_ChannelComp_2MeanMovie(FrameA,RefImage_Range,Filefolder,FileNameHead,ApplyfilterCC,kernelsizeCC,getfocalplane,x_range,y_range,ws,OptionNewpeak)

makeCCImagetoTif=1;
Qshow=1;
saveQ=1;
analysistypename=['Mean_NormXcorr2fit'];
%Applyfilter=1   ;      
filtertype='MeanFilter' ;% 'MedianFilter' 'LogFilter';'GaussFilter'
%kernelsize=5; 
filtersigma=10;

%Motherfolder = uigetdir;
%MotherPath=[Motherfolder '\'];

%cd (Motherfolder)

%OptionNewpeak=0;

if OptionNewpeak==1
SaveName=([analysistypename]);
elseif OptionNewpeak==0
SaveName=([analysistypename]);
end

if ApplyfilterCC==1
SaveName=([SaveName,'_',filtertype,'_', num2str(kernelsizeCC)]);
elseif OptionNewpeak==0
SaveName=([SaveName,'_',filtertype,'_', num2str(kernelsizeCC)]);
end


if iscell(RefImage_Range)
nframes=size(RefImage_Range,2);
elseif length(size(RefImage_Range))==3
nframes=size(RefImage_Range,3);    
elseif length(size(RefImage_Range))==2
nframes=1;
end


if iscell(FrameA)
nframesA=size(FrameA,2);
elseif length(size(FrameA))==3
nframesA=size(RefImage_Range,3);    
elseif length(size(FrameA))==2
nframesA=1;
end


offset_xshift=0;
offset_yshift=0;
%ws=11; %% window size for fitting 


Ref_Frame=zeros(size(RefImage_Range{1}));
Template=zeros(size(FrameA{1}));

for k=1:nframes
Ref_Frame=Ref_Frame+RefImage_Range{k};

%autocorrelation find the original peak (Reference position) 
%ca = normxcorr2(Template,Template); 
end

for k=1:nframesA

Template=Template+FrameA{k};

end


Ref_Frame=Ref_Frame/nframes;
Template=Template/nframesA;

figure
subplot(1,3,1)
imagesc(Template);
set(gca,'YDir','normal');
%titie='movieA'
axis equal tight
xlim([x_range(1) x_range(2)]);
ylim([y_range(1) y_range(2)]);

subplot(1,3,2)
imagesc(Ref_Frame);

set(gca,'YDir','normal');
axis equal tight
%titie='movieB'
hold on
cmean = normxcorr2(Ref_Frame,Template); 


 ori_ypeak = y_range(2);
 ori_xpeak = x_range(2);


%cc = xcorr2(Off_Template,Template); %% for xcoor2 the order is different
  %maxvalue =  max(abs(cc(:)))
   %% Meanfilter substract 
         if ApplyfilterCC==1
             
            cmean=FilterFrameOut_ws(cmean,filtertype,kernelsizeCC,filtersigma);
         end 
  
 % [max_cc, imax] = max(abs(cc(:)));
 
 %[max_cc, imax] = max(abs(cc(:))) %find max from all frames
 cmeancenter=cmean(y_range(2)-10:y_range(2)+10,x_range(2)-10:x_range(2)+10);
 
  [max_cmean]= max(cmeancenter(:));

 %[max_cmean, imax] = max(abs(cmean(:))) %find max from all frames
 
 [ypeak, xpeak] =find(cmean==max_cmean);
%   
%   xpeak
%   ypeak
  reference_y=-(k-1)*offset_yshift;
  reference_x=-(k-1)*offset_xshift;

%auto_c= normxcorr2(Off_Template,Off_Template);


% Off_LargeTemplate=PoissNoise;
% 
% %LargeTemplate= random('poisson',TemplateBgd);
% 
% xrange=[(size(LargeTemplate,1)-size(Template,1)-1)/2+1,((size(LargeTemplate,1)-size(Template,1)-1)/2)+size(Template,1)];
% yrange=[(size(LargeTemplate,2)-size(Template,2)-1)/2+1,((size(LargeTemplate,2)-size(Template,2)-1)/2)+size(Template,2)];
% 
% 
% 
% Off_LargeTemplate(xrange(1):xrange(2),yrange(1):yrange(2))=Off_Template(1:end,1:end);
% %imagesc(Off_LargeTemplate)
% %pause

% auto_c= normxcorr2(Off_Template,Off_Template); 
% % auto_c= xcorr2(Off_Template,Off_Template);
%  [max_auto, imaxauto] = max(abs(auto_c(:)));
%  [auto_ypeak, auto_xpeak] = ind2sub(size(auto_c),imaxauto(1));
 


%% calculate the Peak nearby the center position;
if OptionNewpeak==0 % original use

Ypeak=ori_ypeak;
Xpeak=ori_xpeak;

elseif OptionNewpeak==1
    if abs(ypeak-ori_ypeak)<1
       Ypeak=ori_ypeak;
    else
       Ypeak=ypeak;
    end

    if abs(xpeak-ori_xpeak)<1
       Xpeak=ori_xpeak;
    else
       Xpeak=xpeak;
    end
end
%cc=cc';
pn=(ws-1)/2;
%Q=cc(Ypeak-pn:Ypeak+pn,Xpeak-pn:Xpeak+pn);version 2
Q=cmean(Ypeak-pn:Ypeak+pn,Xpeak-pn:Xpeak+pn);%version 2
cc_Qws=Q;
if Qshow==1
subplot (1,3,3)
colormap(hot);imagesc(Q)
set(gca,'YDir','normal');
axis equal tight
hold off


saveas(gcf,['ComptwoMovie'])
saveas(gcf,['ComptwoMovie'],'tif')

%pause;
close 
%% generate figure for cc
if makeCCImagetoTif==1;
figure
%colormap(gray)
%h=imagesc(cc);
%axis equal tight 
CCFrame=flipud(cmean);
h = imshow(CCFrame,[]);
%cmap = colormap('gray');
CCtest = getimage(h);
%imwrite(CCtest,cmap,['Image_frame',num2str(k),'.tif']);
%saveas(h,['CC_Frame',num2str(k)],'tif')
imwrite(CCtest,['CCmean_Frame','.tif']);
%pause

%  h=(imshow(SimImage{k},[]));
%     Simimagetest = getimage(h);
%     cmap = colormap('gray');
%     Simimagetest=uint16(Simimagetest);
%     imwrite(Simimagetest,cmap,['SimImage_frame',num2str(k),'.tif']);
%close
end


%end
%Q'=Q;
[bb,ci] = fit2dgaussian_4YM(Q,[(ws+1)/2;(ws+1)/2],ws);
% yshift= Ypeak-bb(3)%-Ypeak; % shifted as :0.5
% xshift= Xpeak-bb(2)%-Xpeak;  % shifted as : +0.2
%bb
%pause
% x_fit=bb(2)-((ws+1)/2)+Xpeak -xrange(1)+1;%-ori_xpeak)+ori; % fitting Result after center pixel correction
% y_fit=bb(3)-((ws+1)/2)+Ypeak -yrange(1)+1;%-ori_ypeak); % fitting Result after center pixel correction

 x_fit=bb(2)-((ws+1)/2)+Xpeak; %-ori_xpeak)+ori; % fitting Result after center pixel correction
 y_fit=bb(3)-((ws+1)/2)+Ypeak ;%-ori_ypeak); % fitting Result after center pixel correction


CCall=cmean;
BBall=bb;

%pause

delta_xshift=reference_x+ori_xpeak-x_fit;
delta_yshift=reference_y+ori_ypeak-y_fit;
%pause

%auto_c=auto_c;
% pn=(ws-1)/2;
% autoQ=auto_c(auto_ypeak-pn:auto_ypeak+pn,auto_xpeak-pn:auto_xpeak+pn);
% auto_Qws{k}=autoQ;
% if Qshow==1
% colormap(jet);imagesc(autoQ);
% set(gca,'YDir','normal');
% %pause;
% close;
% end

% [autobb,ci] = fit2dgaussian_4YM(autoQ,[(ws+1)/2;(ws+1)/2],ws);
% auto_x=autobb(2)-((ws+1)/2);
% auto_y=autobb(3)-((ws+1)/2);

%Result(k,:)=[ypeak,xpeak,max_cc,reference_x,reference_y,xshift,yshift,reference_x-xshift,reference_y-yshift];
%Result(k,:)=[ypeak,xpeak,max_cc,reference_y,reference_x,yshift,xshift,delta_yshift,delta_xshift,auto_y,auto_x];
Result_Fit(1,:)=[reference_x,reference_y,x_fit,y_fit,(reference_x+ori_xpeak),(reference_y+ori_ypeak),delta_xshift,delta_yshift];
%disp(Result_Fit);
%Result_Fit=-Result_Fit;
ShiftResult=Result_Fit;
if saveQ==1
[h]=makeimagefile(cc_Qws,'gray','cc_Qws');
%[h]=makeimagefile(ccQ_all,'gray','RefImage_Range');
end

%% Result Figure
figure;
subplot(2,1,1)
plot(1:nframes,Result_Fit(:,7),'--*b'); hold on
%plot(1:nframes,Result_Fit(:,8)+ori_xpeak,'b'); hold on
ylabel('X (pixel)','fontsize',16);
%xlabel('frame(nth)','fontsize',16);
title([analysistypename]);
subplot(2,1,2)
plot(1:nframes,Result_Fit(:,8),'--*r');hold on
%plot(1:nframes,Result_Fit(:,6)+ori_ypeak,'r'); hold on
ylabel('Y (pixel)','fontsize',16);
xlabel('Movie(nth)','fontsize',16);
title([analysistypename]);%,' shifty:',num2str(offset_yshift)]);
save(['NormXcorr2_Result2D_Fit'],'Result_Fit');

saveas(gcf,['NormXcorr2_Result2D_Fit'])
saveas(gcf,['NormXcorr2_Result2D_Fit'],'bmp')
%analysistypename='Xcorr2 2DFit';
%deltashift(Result_Fit,shiftx,shifty,sx,sy,amp,snr,analysistypename)
%deltashift_fit(Result_Fit,offset_xshift,offset_yshift,analysistypename)

% %% compare autocorrelation
% 
% figure;
% % subplot(1,2,1)
% AutoXnm=Result_Fit(:,9)*66.67;
% AutoYnm=Result_Fit(:,10)*66.67;
% plot(1:nframes,AutoXnm,'-*k'); hold on
% plot(1:nframes,AutoYnm,'-ok'); hold on
% % legend('X','Y')
% % legend('boxoff')
% hline = refline([0 0]);
% set(hline,'Color','k')
% ylabel('shift(nm)','fontsize',16);
% xlabel('frame(nth)','fontsize',16);
% title(['Autoxcoor2 2Dfit']);%; shiftx:',num2str(offset_xshift),' shiftx:',num2str(offset_yshift)]);
% % subplot(1,2,2)
% % plot(Result_Fit_nm(:,1),Result_Fit_nm(:,10),'-*k');hold on
% % plot(Result_Fit_nm(:,2),Result_Fit_nm(:,6),'k'); hold on
% % ylabel('nm','fontsize',16);
% % xlabel('shift(pixel)','fontsize',16);
% % title(['NormXcorr2_2DFit shifty:',num2str(offset_yshift)]);
% % save(['Xcorr2_Result2D_Fit'],'Result_Fit');
% saveas(gcf,['NormXcorr2_Result2D_Auto'])
% saveas(gcf,['NormXcorr2_Result2D_Auto'],'bmp')
% close; 
% save(['NormXcoor2_Result_Fit_pix',SaveName],'Result_Fit');
% save(['Xcorr2_Result_Fit_pix_',SaveName],'Result_Fit','-ascii');
Result_Fit_nm=Result_Fit*66.67;
save(['NormXcorr2_Result_Fit_nm_',SaveName],'Result_Fit_nm');
save(['NormXcorr2_Result_Fit_nm_',SaveName],'Result_Fit_nm','-ascii');
%elseif option==2
%save(['SimResultSummary2_',num2str(PhotonNum),'_',num2str(ParticleNum),'_',num2str(framex),'_',num2str(framey),'_',num2str(nframes)],'Result2');
%end
save(['CCall_',SaveName],'CCall');
save(['BBall_',SaveName],'BBall'); 
save(['cc_Qws_',SaveName],'cc_Qws');
% save(['auto_Qws_',SaveName],'auto_Qws');

%% ParameterSummary
% Sim_Parameter.MoiveName=SaveName;
% Sim_Parameter.SaveFolderName=SaveFolderName;
% Sim_Parameter.PhotonNum=PhotonNum;
% Sim_Parameter.ParticleNum=ParticleNum;
% Sim_Parameter.Sigma_x=Sigma_x;
% Sim_Parameter.Sigma_y=Sigma_y;
% Sim_Parameter.framex=framex;
% Sim_Parameter.framey=framey;
% Sim_Parameter.offset_yshift=offset_yshift;
% Sim_Parameter.offset_xshift=offset_xshift;
% Sim_Parameter.ws=ws;
% Sim_Parameter.AddNoise=AddNoise;
% Sim_Parameter.nmovie=nmovie;
% Sim_Parameter.fixedcenterposition=fixedcenterposition;
% Sim_Parameter.MeanPixelBgd=MeanPixelBgd;
% Sim_Parameter.EMgainofCCD=EMgainofCCD;
% Sim_Parameter.QE=QE;
% Sim_Parameter.Sensitivity=Sensitivity;
% % Sim_Parameter.pixelsizenm=pixelsizenm;
% 
% save(['ParameterSummary'],'Sim_Parameter');
% save(['CCall_',num2str(PhotonNum),'_',num2str(ParticleNum),'_',num2str(framex),'_',num2str(framey),'_',num2str(nframes)],'CCall_2');
% save(['BBall_',num2str(PhotonNum),'_',num2str(ParticleNum),'_',num2str(framex),'_',num2str(framey),'_',num2str(nframes)],'BBall_2')
%save(['SimOriginalLoc_',num2str(PhotonNum),'_',num2str(ParticleNum),'_',num2str(framex),'_',num2str(framey),'_',num2str(nframes)],'ccAll');

%close all

end
 %end
%end
%%% How to use
%%% Particle_Loc is the list of the x,y corrdiates of the particle 
%%% try to change the relevant background if you want to simulate the more
%%% similar situation. Random is for the possion random background
%%% %%%[Images,Particle_Loc] = SimulateParticleField(10,5)
%%%surf(Images{1})
```
