## Supplementary material for "Microvillar cartography: a super-resolution single-molecule imaging method to map the positions of membrane proteins with respect to cellular surface topography": Analysis MATLAB codes: VATipinROI_sort.m

```
%% Deleter LocTips outside of the region of interests;
function[ROILocTipsTable]=VATipinROI_sort(LocTipsTable,range_x,range_y)
% ROIws=101; % analysis area limit to a cell region
% ws=9; % window for a individual tip rigion
% xcnt=135;
% ycnt=130;
% ROIpn=(ROIws-1)/2;
% pw=(ws-1)/2;

TipData=LocTipsTable';

xROIlim=[range_x];
yROIlim=[range_y];


cnt=0;
    for k=1:size(TipData,1);

        if TipData(k,2)>=yROIlim(1) && TipData(k,2)<=yROIlim(2) && ...
                TipData(k,3)>=xROIlim(1) && TipData(k,3)<=xROIlim(2) 

            cnt=cnt+1;  
            ROILocTipsTable(cnt,:)=TipData(k,:);

        end

    end
ROILocTipsTable=ROILocTipsTable';% for saving as the same format..
end
```
