## Supplementary material for "Microvillar cartography: a super-resolution single-molecule imaging method to map the positions of membrane proteins with respect to cellular surface topography": Analysis MATLAB codes: VAtipRIO_contour_Map.m

```
%% Limit Tips within ROI;
function[ret,ROILocTipsTable]=VAtipRIO_contour_Map(SaveName,VARec,myColorMap,MyMarkerColor,LocTipsTable,range_x,range_y,markersize,minoption,zMax)

[newLocTipsTable]=VATipinROI_sort(LocTipsTable,range_x,range_y);
newLocTipsTable=newLocTipsTable';

ROILocTipsTable=zeros(size(newLocTipsTable,1),11);
cnt=0;

[B,IX]=sort(newLocTipsTable(:,4),'ascend');

for jj = 1:size(newLocTipsTable,1)
    cnt=cnt+1;
     A = newLocTipsTable(IX(cnt),:); 
     A(1,11)=A(1);
     A(1,1) = cnt;
     %A(1,11) = newLocTipsTable(IX(cnt),1);
     ROILocTipsTable(cnt,:)= A;
end

%ROILocTipsTable= A;
%% Draw figure
% myColorMap = colormap('gray(16)'); %set the colormap used  %autumn
% %myColorMap =(flipud(myColorMap));

ret=figure
colormap(myColorMap)
resultImage = contour(VARec);

set(gca,'YDir','normal')
%colorbar
caxis([0 zMax]) 
% set(resultImage,'EdgeColor','none')
% view ([0 90])
%axis resultImage;
set(gca,'LooseInset',get(gca,'TightInset'))
axis equal square
hold all;

for mol=1:size(ROILocTipsTable,1)
    if minoption==0
    Posy= ROILocTipsTable(mol,2); %% 
    Posx= ROILocTipsTable(mol,3);   
    elseif minoption==1
    Posy= ROILocTipsTable(mol,6); %% 
    Posx= ROILocTipsTable(mol,7);   
    end
%CM=Fit2SingleMol(:,2)
%CN=Fit2SingleMol(:,3)
 scatter(Posx,Posy,20,'+','MarkerEdgeColor',MyMarkerColor);
% text(blobCentroid(1) + labelShiftX, blobCentroid(2), num2str(mol), 'FontSize', fontSize, 'FontWeight', 'Bold');
text(Posx+2, Posy +0.5, num2str(mol), 'FontSize', 16, 'FontWeight', 'Bold','Color',[0 0 0]); 
set(gca,'LooseInset',get(gca,'TightInset'))
if minoption==0
 title ('LocTips (+ min pixel)','Fontsize',18)
elseif minoption==1
 title ('LocTips (+ centroid)','Fontsize',18)
end
end
addscale3_um_range_label(range_x,range_y,markersize,0)


    saveas(gcf,SaveName);%,'jpg'); 
    set(gcf, 'Position', get(0, 'ScreenSize')); 
    saveas(gcf,['scale' SaveName],'jpg'); 

save(['Table_' SaveName],'ROILocTipsTable');


hold off
end
```
