## Supplementary material for "Microvillar cartography: a super-resolution single-molecule imaging method to map the positions of membrane proteins with respect to cellular surface topography": Analysis MATLAB codes: zSTDlimit_sort.m

```
function[zSTD_LocTipsTable]=zSTDlimit_sort(ROILocTipsTable,zSTDlimit)
%zSTD_LocTipsTable=newLocTipsTable';
cnt=0;
for kk=1:size(ROILocTipsTable,1)
if ROILocTipsTable(kk,5)<zSTDlimit
    cnt=cnt+1;
newLocTipsTable(cnt,:)=ROILocTipsTable(kk,:);
end
end


zSTD_LocTipsTable=zeros(size(newLocTipsTable,1),11);

cnt=0;
[B,IX]=sort(newLocTipsTable(:,4),'ascend');


for jj = 1:size(newLocTipsTable,1)

    cnt=cnt+1;
     A = newLocTipsTable(IX(cnt),:); 
     A(1,11)=A(11); % orignal label number
     A(1,1) = cnt;
   
     %A(1,11) = newLocTipsTable(IX(cnt),1);
     zSTD_LocTipsTable(cnt,:)= A;

end

zSTD_LocTipsTable=zSTD_LocTipsTable';


end
```
